# microRNA-218-5p Coordinates Scaling of Excitatory and Inhibitory Synapses during Homeostatic Synaptic Plasticity

**DOI:** 10.1101/2023.10.20.563224

**Authors:** David Colameo, Sara M. Maley, Waleed ElGrawani, Carlotta Gilardi, Simon Galkin, Steven A. Brown, Gerhard Schratt

## Abstract

Homeostatic synaptic plasticity (HSP) is a fundamental neuronal mechanism that allows networks to compensate for prolonged changes in activity by adjusting synaptic strength. This process is crucial for maintaining stable brain function and has been implicated in memory consolidation during sleep. While scaling of both excitatory and inhibitory synapses plays an important role during homeostatic synaptic plasticity, molecules coordinating both of these processes are unknown.

In this study, we investigate the role of miR-218-5p as a regulator of inhibitory and excitatory synapses in the context of picrotoxin (PTX)-induced homeostatic synaptic downscaling (HSD) in rat hippocampal neurons. Using enrichment analysis of miRNA-binding sites in differentially expressed genes changing upon PTX-induced HSD, we bioinformatically predicted and experimentally validated increased miR-218-5p activity upon PTX-treatment in the process compartment. By monitoring synapse structure *in vitro* with confocal microscopy, we demonstrate that inhibiting miR-218-5p activity exerts a dual effect during HSD: it prevents the downscaling of excitatory synapses and dendritic spines, while at the same time blocking inhibitory synapse upscaling. Furthermore, we identify the Neuroligin2 interacting molecule Mdga1 as a crucial target of miR-218-5p in the context of homeostatic upscaling of inhibitory synapses. By performing long-term electroencephalographic (EEG) recordings, we further revealed that local inhibition of miR-218-5p in the somatosensory cortex reduced local slow-wave activity (SWA) during non-rapid-eye-movement (NREM) sleep.

In summary, this study uncovers miR-218-5p as a key player in coordinating inhibitory and excitatory synapses during homeostatic plasticity and sleep. Our findings contribute to a deeper understanding of how neural circuits maintain stability in the face of activity-induced perturbations, with potential implications for both physiological and pathological conditions.

**Significance Statement:** Homeostatic synaptic plasticity mechanisms evolved to keep neuronal firing rates within a physiological range in response to alterations in neural network activity. It has been proposed that similar mechanisms take place during sleep in a process that promotes memory consolidation and synaptic renormalization. In this study, posttranscriptional regulation of synaptic proteins by miR-218-5p has been identified to coordinate both excitatory and inhibitory synaptic scaling during activity-dependent homeostatic synaptic plasticity. Intriguingly, local inhibition of miR-218-5p in the cortex of mice resulted in reduced slow-wave activity, an EEG-signature of synchronous firing during non-rapid eye movement sleep and a hallmark correlate of sleep pressure. Overall, these findings propose a convergent, posttranscriptional mechanism to coordinate both excitatory and inhibitory synaptic strength in response to alterations in neuronal activity with potential implications for sleep.

## Introduction

In response to prolonged changes in neural activity, homeostatic synaptic plasticity (HSP) emerges as a fundamental regulatory process that maintains equilibrium within neuronal networks (1, 2). While Hebbian plasticity has been associated with input-specific adaptations that change in the direction of activation, HSP is considered a global-weight normalizer that scales synaptic strength within a dynamic range of plasticity opposite to the direction of stimulation. In fact, it is well established that hyperactive neuronal networks will scale down excitatory synapses in both number and size in a mechanism that has been coined as homeostatic synaptic downscaling. In contrast, silencing neuronal activity will result in long-term strengthening of excitatory synapses, also known as homeostatic synaptic upscaling (3). These distinct forms of homeostatic synaptic plasticity play a critical role in maintaining networks that are not only flexible but also stable.

So far, in vitro studies on picrotoxin or bicuculine-induced HSD or TTX-induced HSU typically focused on changes at excitatory synapses. However, it has been shown that inhibitory GABAergic synapses similarly undergo scaling during extended periods of neural network suppression or hyperactivity (4–7). Intriguingly, it appears that the scaling at inhibitory synapses occurs in the opposite direction compared to their excitatory counterparts. Therefore, we will refer in the rest of this study to the reciprocal upscaling at inhibitory and downscaling at excitatory synapses as activity– or PTX-induced homeostatic synaptic plasticity (HSP).

Changes in gene expression play a pivotal role in mediating HSP, occurring both neuron-wide and in a compartment-specific manner along neuronal processes (8–10). Emerging evidence underscores the significance of activity-dependent microRNAs (miRNAs) in post-transcriptionally regulating various pathways in both homeostatic synaptic upscaling (11–13) and downscaling (14–17) of excitatory synapses. While extensive research has unveiled the molecular mechanisms underlying excitatory synapse homeostatic plasticity (3), the molecular control of synaptic homeostasis at inhibitory synapses remains relatively unexplored, specifically with regard to a potential involvement of miRNAs in this process.

From a physiological perspective, visual sensory deprivation results in homeostatic upscaling of excitatory synapses in the visual cortex during development (18, 19). In contrast, homeostatic synaptic downscaling has emerged as an important component of sleep-dependent memory consolidation (20–22). Analogous to human EEG recordings, distinct frequency patterns characterize brain activity in rodents during different sleep stages. Rapid eye movement (REM) sleep is characterized by dominance of EEG theta band frequencies (6-10Hz), while non-rapid eye movement (NREM) sleep exhibits slow waves within the delta band frequencies (0.5-4Hz) (23). Notably, the power of these slow waves, also known as slow-wave activity (SWA), monotonically build up during wakefulness, reaches highest levels at first episodes of sleep, and dissipates across the course of sleep (24). Following prolonged wakefulness, as experimentally induced via sleep deprivation and/or restrictions, NREM SWA is further increased in a manner that is proportional to the sleep loss. Therefore, SWA during NREM sleep is a reliable correlate of sleep pressure in various species including humans (25) and rodents (23).

Additionally, sleep plays a critical role in memory consolidation. This is evident in the impairment of memory recall that is associated with the lack of sleep (26, 27). The synaptic homeostasis hypothesis proposes that SWA during sleep mediates synaptic downscaling (20) and undergoes local, use-dependent alterations after learning tasks (28).

Recent investigations have unveiled dynamic fluctuations in excitatory-inhibitory (E-I) balance that are dependent on the time of day (29), as observed in the visual cortex of mice. During sleep phases, the frequency of miniature excitatory postsynaptic current events (mEPSCs) decreases, while miniature inhibitory postsynaptic current frequency exhibits an opposing increase. This study unveils E-I balance oscillations tied to sleep patterns, reminiscent of the bidirectional scaling of inhibitory and excitatory synapses observed during activity-driven HSP *in vitro*.

Furthermore, recent comprehensive explorations into the synaptic transcriptome, proteome, and phosphoproteome shed light on the circadian regulation of synapse-localized transcript abundance, whereby protein phosphorylation and accumulation respond to sleep-wake cycles (30, 31). Notably, genes associated with synaptic plasticity and memory peak during the light phase (the sleep phase in rodents), while genes linked to metabolism and oxidative stress peak during the dark phase. It has been concluded that circadian accumulation of transcripts prepares specific genes at synapses, while sleep pressure drives activity-dependent protein phosphorylation and protein expression. Investigating whether microRNAs contribute to the posttranscriptional regulation of synapse-localized transcripts and their activity-driven expression holds promise for further insights.

Here, we identify miR-218-5p as a novel coordinator of activity-induced homeostatic scaling of inhibitory and excitatory synapses in response to prolonged increase in neuronal activity. miR-218-5p is upregulated in the synaptic compartment of primary rat hippocampal neurons during scaling. Using confocal microscopy, we revealed that miR-218-5p is essential for the downscaling of structural parameters associated with excitatory synaptic strength (spine volume, GluA1/GluA2 co-cluster size), as well as those associated with the upscaling of inhibitory synaptic strength (gamma2-clusters and Nlgn2-vGAT co-clusters). RNA-seq analysis of cortical regions injected with miR-218-5p inhibitors further allowed us to identify the inhibitory synapse regulator Mdga1 as a novel direct target of miR-218-5p. Moreover, by conducting long-term electroencephalographic (EEG) recordings in the mouse cortex, we observed a specific reduction in slow-wave-activity-associated frequency bands in hemispheres injected with miR-218-5p inhibitors compared to control hemispheres. This was accompanied by an increase in NREM sleep duration during later phases of the dark phase. Collectively, our study suggests that the coordinated control of excitatory and inhibitory synaptic function by microRNAs could play an important role in physiological processes involving homeostatic plasticity mechanisms.

## Results

### miR-218-5p Activity Predicted to Increase During Homeostatic Synaptic Downscaling

In a previous study, we conducted an extensive multi-omics analysis to characterize compartmentalized gene expression during picrotoxin (PTX)-induced homeostatic synaptic downscaling (HSD) (10). In this follow-up investigation, we aimed to identify novel compartment-specific, post-transcriptional regulators of HSD, focusing on miRNAs. We took advantage of the previously published RNA-sequencing (RNA-seq) dataset from PTX or mock-treated compartmentalized rat hippocampal neurons cultures, which allowed to differentiate between RNAs regulated in the somatic and process (dendritic and axonal) compartment. Upon updating the RNA-seq dataset from Rno6 to Rno7, we uncovered marked differences in gene expression when testing for compartment– and PTX-treatment specific effects (Suppl. Figure1A-C, Suppl. Table 1). We identified 3656 genes enriched in the somatic and 3591 genes enriched in the process compartment (Suppl. Figure1C, Suppl. Figure2B,D) and 1060 downregulated genes and 1301 upregulated genes that reacted to PTX-treatment irrespective of their compartment-localization (Suppl. Figure 1C, Suppl. Figure 2A,C).

Furthermore, through surrogate variable analysis, we achieved improved correction of batch effects in our dataset, enhancing our ability to perform interaction analysis between compartments and PTX-treatment effects (Suppl. Figure 1C). This approach enabled us to capture compartment-specific effects in response to PTX treatment statistically accurately. Specifically, 118 genes exhibited specific differential expression within the process compartment compared to the overall PTX effect (Figure 1A-B).

**Figure 1:**
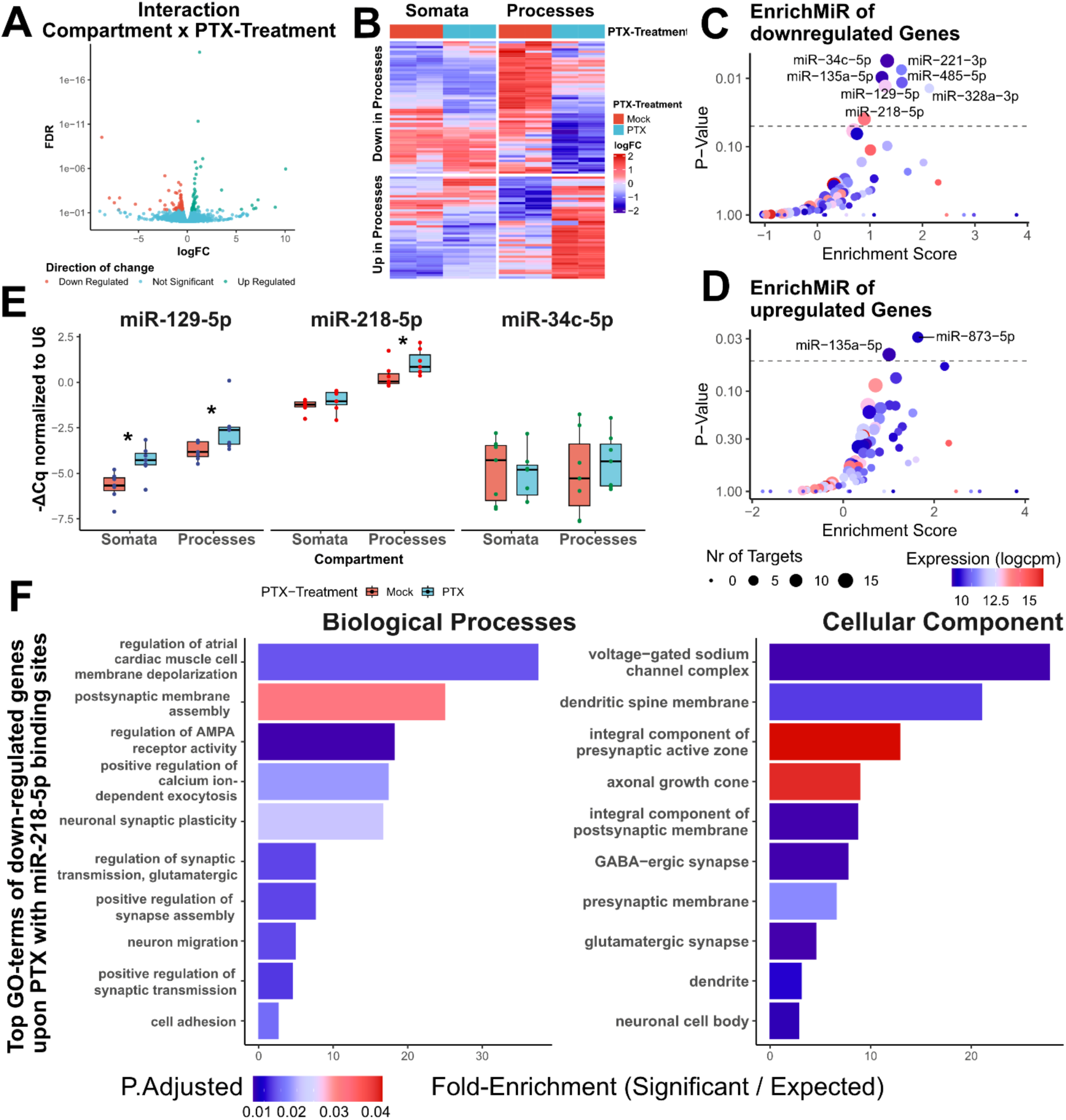
miR-218-5p activity is upregulated during homeostatic synaptic downscaling. (A) Volcano of differentially expressed genes (DEG) changing upon PTX in the process compartment (Interaction-term: Compartment x PTX-Treatment (B) Heatmap of log-fold changes (logFC, relative to mock-conditions) of all significantly changing DEGs from A. (C-D) EnrichMiR of down– and upregulated genes in the process-compartment upon PTX-treatment, respectively. Figure 1 continued on the next page. (E) Taqman RT-qPCR of three candidate miRNA (miR-129-5p, miR-218-5p and miR-34c-5p) upon PTX-treatment in compartmentalized cultures; n=7 independent biological replicates, data was fit to a linear-mixed model with fixed effects for PTX-treatment and compartment-localization and their interaction and random effect of independent biological replicates. Dunn’s post-hoc analysis was used to estimate PTX-effect in somatic and process-compartments *p<0.05 (F) Top 20 Gene Ontologies associated with predicted targets of miR-218-5p during PTX-treatment in the somatic and process-compartment.

To identify potential microRNA-binding site (MBS) enrichment in differentially expressed genes, we employed *enrichMiR* analysis using their dedicated toolbox (32). *enrichMiR* allows to statistically test for overrepresentation of MBS in sets of genes from differential expression analyses. Consistent with our previous findings (10), we observed an increased enrichment of MBSs in downregulated genes following PTX treatment compared to upregulated genes, indicating increased functional activity of miRNAs during HSD (Suppl. Figure 2E, 2G). MBS enrichment in downregulated genes by PTX-treatment included activity-dependent miRNAs involved in synaptic plasticity such as miR-132-3p, miR-191a-5p, members of the miR-34 family and miR-218-5p (33–36) (Suppl. Figure 2E). Similarly, we identified an increased enrichment of MBSs in genes enriched in the process-compartment compared to genes enriched in the somatic compartment (Suppl. Figure 2F,H), supporting the notion of an increased necessity in local, post-transcriptional control of process-enriched genes (10).

The interpretation of differentially expressed genes (DEGs) under the interaction term (compartment × PTX treatment) requires careful consideration. To calculate the adjusted log-fold change for genes exhibiting distinct responses to PTX in the process compartment, we summed up the log-fold changes of the interaction term with those of the main PTX effect. Subsequently, we conducted *enrichMiR* analysis using a log-fold cutoff of log2(1.2) (equivalent to a 20% change upon PTX treatment) for significant genes. This approach identified the enrichment of miR-129-5p, a previously described regulator of HSD (16), as well as miR-218-5p and miR-34-5p in genes specifically downregulated in the process-compartment (Figure 1C).

We next wanted to assess whether changes in miRNA activity predicted by *enrichMiR* translated into corresponding alterations in miRNA levels. Therefore, we conducted RT-qPCR measurements for three miRNA candidates, miR-218-5p, miR-34c-5p, and miR-129-5p, the latter serving as a positive control based on a prior study (16). For the miR-34-5p family, we focused on the most abundant isoform in primary hippocampal cells, namely miR-34c-5p (16).

Consistent with our previous results (16), miR-129-5p was upregulated by PTX in both compartments. Notably, miR-218-5p demonstrated significant upregulation exclusively in the process compartment, with an increase of around 1.67-fold after PTX treatment (Figure 1E). Conversely, miR-34c-5p expression remained unchanged in both compartments (Figure 1E). Based on these results, we decided to focus on miR-218-5p as a potential compartment-specific regulator of HSD for our further analysis.

Gene ontology (GO) analysis of downregulated DEGs containing binding sites for miR-218-5p revealed an overrepresentation of GO terms associated with excitatory synapse function, synaptic plasticity, and memory (Figure 1F). Particularly noteworthy, miR-218-5p-regulated genes were also associated with GABAergic synapses and the regulation of AMPA receptor activity.

Collectively, these findings suggest that we have successfully captured the gene signature of a potential new regulator of HSD, which may preferentially exert its regulatory effects in the synapto-dendritic compartment of hippocampal neurons.

### Blocking miR-218-5p Activity Impedes PTX-induced Excitatory Synaptic Downscaling

We proceeded to investigate the functional impact of miRNA manipulation on spine morphology to evaluate structural alterations induced by HSD. Measuring activity-dependent changes in spine head volume is a well-validated correlate of synaptic strength *in vitro* (15, 16, 37). To suppress miRNA activity, we transfected locked nucleic acid (LNA) modified antisense oligonucleotides that sequester endogenous miRNAs. Spine volume was assessed in PTX or mock-treated cultures after transfection by confocal microscopy based on the co-transfected GFP (Figure 2A).

**Figure 2:**
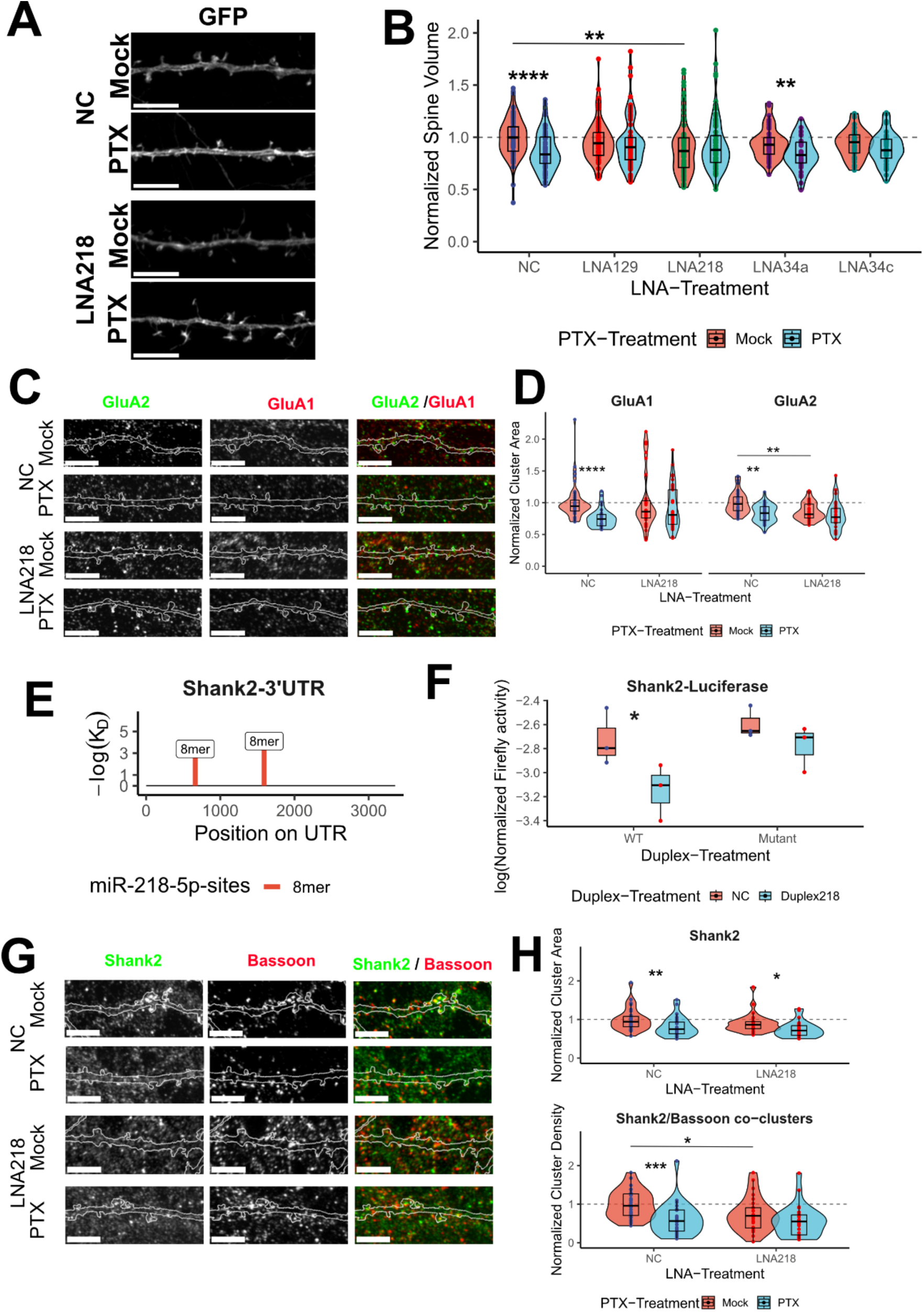
miR-218-5p occludes homeostatic downscaling of excitatory synapses in rat hippocampal neurons in vitro. (A) Example pictures of dendrites of primary hippocampal neurons transfected with either negative control LNA or LNA against miR-218-5p (LNA218) and treated with either PTX or vehicle (Mock). White tool bars represent 5µm. (B) Quantification of spine head volume normalized to the mean of the mock-treated, negative control (NC) group of the corresponding independent biological replicate. 8-14 cells per condition and per independent biological replicate (n=8). Data was fit to a linear-mixed model with fixed effects for PTX-treatment and LNA-treatment and their interaction and random effect of independent biological replicates. Dunn’s post-hoc analysis was used to estimate PTX-effect in different LNA-condition and to estimate LNA-effect compared to negative control within PTX-treatment *p<0.05, **p<0.01, ***p<0.001, ****p<0.0001. (C) Example pictures of dendrites of primary hippocampal neurons transfected with GFP (white outline) and either NC or LNA218 in combination with PTX– and mock-treatment stained for AMPA-R subunits GluA2 and GluA1. White tool bars represent 5µm. (D) Quantification of cluster area of GluA2 and GluA1 normalized to the mean of the mock-treated, negative control (NC) group of the corresponding independent biological replicate of primary hippocampal preparations. 8-12 images per condition and per independent biological replicate n=3. Data was fit to a linear-mixed model with fixed effects for PTX-treatment and LNA-treatment and their interaction and random effect of independent biological replicates. Dunn’s post-hoc analysis was used to estimate PTX-effect in NC and LNA218-conditions and to estimate LNA218-effect compared to negative control within PTX-treatment *p<0.05, **p<0.01, ***p<0.001, ****p<0.0001. (E) 3’ untranslated regions (3’UTRs) of Shank2 containing miR-218-5p binding sites as predicted by *scanMiR* (40) denoted as 8mer seed regions. (F) Luciferase assay of a fragment of the wildtype and mutant 3’UTR of Shank2 upon miR-218-5p overexpression by Duplex218 or negative control duplex (NC). Values are ratio between Firefly-bioluminescence and transfection-control of Renilla bioluminescence, log2-transformed. n=3 independent biological replicates in primary cortical preparations. Data was fit to a linear-mixed model with fixed effects for Genotype (WT vs Mutant) and Duplex-treatment and their interaction and random effect of independent biological replicates. Dunn’s post-hoc analysis was used to estimate Duplex-effect in WT and mutant-conditions and to estimate Duplex218-effect compared to negative control within Genotype *p<0.05. (G) Example pictures of dendrites of primary hippocampal neurons transfected with GFP (white outline) and either NC or LNA218 in combination with PTX– and mock-treatment stained for Shank2 and Bassoon. White tool bars represent 5µm. (H) Quantification of cluster area of Shank2 (upper plot) and Shank2-Bassoon co-cluster densities (lower plot) normalized to the mean of the mock-treated, negative control (NC) group of the corresponding independent biological replicate of primary hippocampal preparations. 8-12 images per condition and per independent biological replicate n=3. Data was fit to a linear-mixed model with fixed effects for PTX-treatment and LNA-treatment and their interaction and random effect of independent biological replicates. Dunn’s post-hoc analysis was used to estimate PTX-effect in NC and LNA218-conditions and to estimate LNA218-effect compared to negative control within PTX-treatment *p<0.05, **p<0.01, ***p<0.001, ****p<0.0001.

Under scrambled LNA control conditions (negative control, NC), spine volume exhibited a significant reduction of about 13% (Figure 2B) following PTX-induced HSD, in agreement with previous results (15, 16). As a positive control, we included inhibition of miR-129-5p (16), which effectively prevented PTX-induced spine volume reduction (Figure 2B). Inhibition of miR-218-5p activity (LNA218) consistently resulted in smaller spines, displaying a reduction of approximately 0.9-fold compared to mock-treated NC. This suggests a fundamental role for miR-218-5p in maintaining baseline spine size. PTX treatment failed to further decrease spine volume in LNA218 transfected neurons, indicating that miR-218-5p inhibition occludes HSD. Spines in this condition rather exhibited a non-significant trend towards an increased size (observed in 7 of 8 replicates, with a 1.07-fold change compared to mock-treated LNA218 condition). In contrast, LNA-mediated inhibition of two members of the miR-34 family (miR-34a, miR-34c) did neither have a significant effect on baseline spine size nor interfere with PTX-dependent spine size reduction, demonstrating specificity of our approach.

To extend our findings from dendritic spine morphology, we explored the requirement for miR-218-5p activity in mediating alterations in post-synaptic proteins after HSD induction. The reduction in cluster number and size of α-amino-3-hydroxy-5-methyl-4-isoxazolepropionic acid receptor (AMPA-R) subunits, GluA1 and GluA2, is a well-established mechanism in HSD (20, 22). To investigate the potential direct or indirect role of miR-218-5p in downregulating GluA2 and GluA1 clusters during HSD, we conducted surface immunostaining in hippocampal neurons co-transfected with GFP and LNA218 (Figure 2C).

PTX treatment led to smaller surface GluA1 and GluA2 clusters (0.77-fold and 0.83-fold reduction, respectively; Figure 2D), accompanied by a decrease in the number of co-localized clusters (Suppl. Figure 3D; 0.71-fold change), demonstrating efficient downscaling of surface AMPA-R with our protocol. Blocking miR-218-5p activity resulted in reduced GluA2, but not GluA1 cluster area under baseline conditions (0.85-fold change; Figure 2D), mirroring our spine results (Figure 2B). Importantly, neurons transfected with LNA218 failed to downscale GluA1 and GluA2 cluster area following PTX treatment (Figure 2D), providing independent support for the requirement of miR-218-5p in HSD. GluA1/2 co-cluster density was unaffected by LNA treatment (Suppl. Figure 3D).

Furthermore, beyond AMPA-R subunit dynamics during HSD, we examined structural alterations in Shank2, a postsynaptic density scaffolding protein, and Bassoon, a general presynaptic marker essential for active zone organization. Previous studies have indicated the regulation of members of the Shank-family during homeostatic synaptic plasticity (38). Interestingly, the 3’UTR of the Shank2 gene contains two strong 8-mer binding sites for miR-218-5p (Figure 2E). We therefore conducted a dual luciferase assay to evaluate their functional relevance. Overexpressing miR-218-5p through mimic transfection reduced luciferase activity regulated by the wildtype (WT) 3’UTR of Shank2 at the post-transcriptional level (0.74-fold reduction in luciferase activity; Figure 2F). Notably, mutation of miR-218-5p binding sites attenuated the downregulation following miR-218-5p overexpression (0.88-fold compared to the mutant control condition, Figure 2F), confirming the responsiveness of Shank2 3’UTR to changes in miR-218-5p levels.

To explore the relationship between miR-218-5p regulation and Shank2 in a more physiological context, we performed immunostainings of Shank2 and Bassoon in GFP-transfected hippocampal neurons in combination with LNA218 and PTX treatment (Figure 2G). In control conditions, we observed a reduction in co-localized Shank2-Bassoon co-cluster density (0.61-fold change, Figure 2H, lower graph) and decreased Shank2 cluster size following HSD induction (0.81-fold change, Figure 2H, upper graph). Additionally, we noted a non-significant trend toward smaller Shank2 clusters and significantly fewer Shank2-Bassoon co-clusters under baseline conditions when miR-218-5p was blocked compared to the negative control (Figure 2H, lower graph).

Interestingly, in combination with PTX-treatment, while inhibiting miR-218-5p activity attenuated but not completely abolished downscaling of Shank2 cluster size (Figure 2H, upper graph), downscaling of Shank2-Bassoon co-clusters was occluded in LNA218 conditions (Figure 2H, lower graph). In summary, our findings suggest that miR-218-5p is a positive regulator of excitatory synapse structure under basal activity conditions and occludes PTX-induced HSD.

### Stereotactic injections of miR-218-5p inhibitors into the somatosensory cortex I upregulates negative regulator of inhibitory synapses Mdga1

Our subsequent goal was to investigate potential targets of miR-218-5p through a more direct and unbiased approach. Recognizing the challenges of achieving high transfection efficiency in primary neuronal cultures, we leveraged gymnosis-based delivery of short nucleotides by directly injecting LNAs into cortical regions of mice (Figure 3A, cartoon) (16, 39).

**Figure 3:**
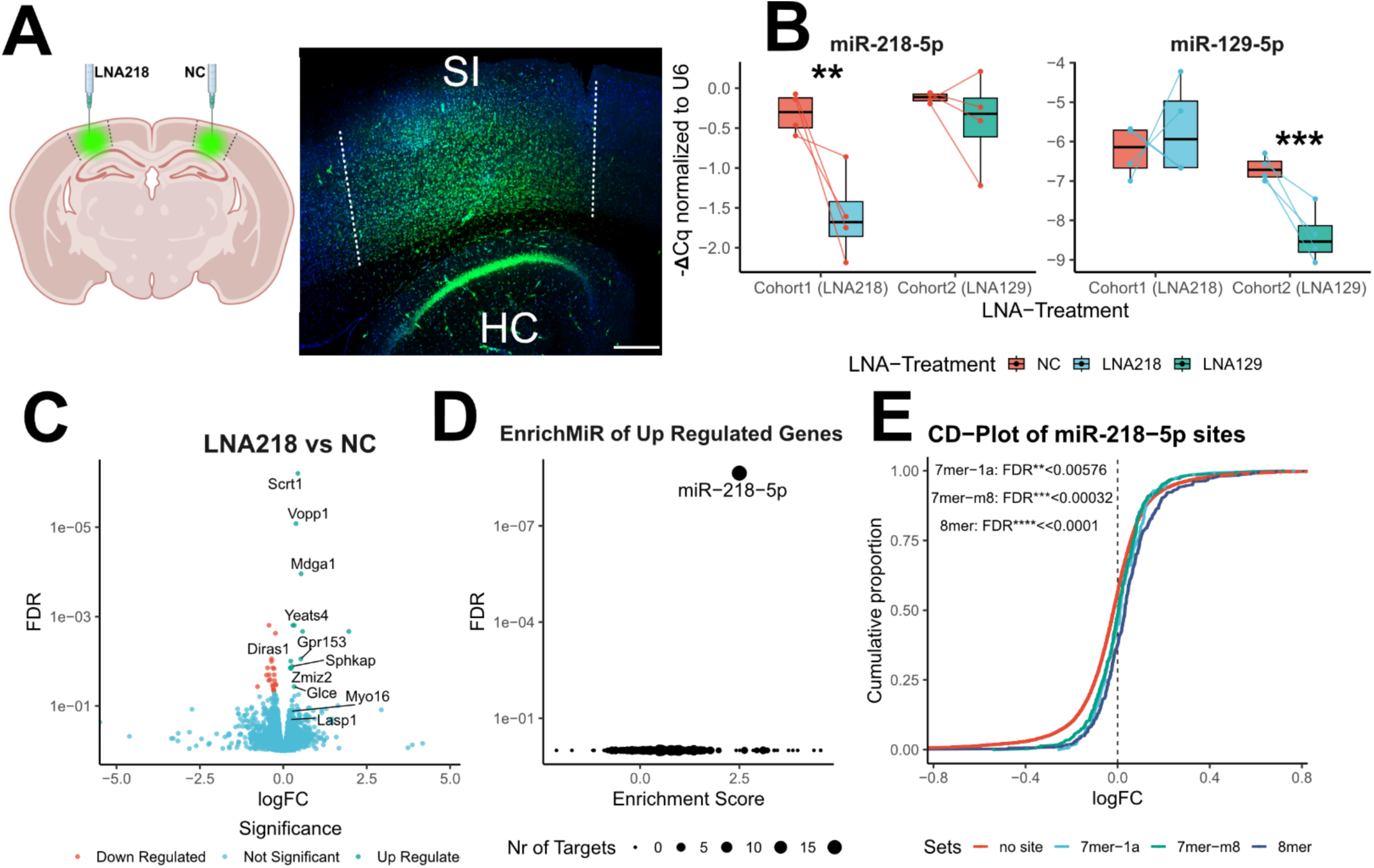
Identification of miR-218-5p target mRNAs in the mouse cortex *in vivo*. (A) Schematic of stereotactic injections (left) and example picture of fluorescently labelled LNAs (right) in the somatosensory cortex I of mouse brains. White bar represents 150µm. (B) Validation of LNA-mediated knockdown by Taqman RT-qPCR. – delta Cq-values of miR-218-5p (right plot) or miR-129-5p (left plot) normalized to U6 comparing hemispheres injected with LNA-negative control (NC) or LNA against miR-218-5p (LNA218) or against miR-129-5p (LNA129). n=4 mice. Data was fit to a linear-mixed model per miRNA with fixed effects for LNA-treatment and random effect of independent biological replicates (mice). Dunn’s post-hoc analysis was used to estimate LNA-effect on hemispheres injected with either LNA218 or LNA129 compared to corresponding NC-injected hemisphere within mouse. **p<0.01, ***p<0.001 (C) Volcano plot showing log-fold-changes (logFC) to LNA218 treatment vs NC and significance level by FDR. Labelled genes contain predicted miR-218-5p binding sites. Significance threshold: logFC<0 and FDR < 0.05 as downregulated: logFC<0 and FDR < 0.05 as upregulated. (D) enrichMiR-analysis of top 50 upregulated genes (FDR <0.1) (E) Cumulative distribution plot of all genes containing predicted miR-218-5p binding sites divided by seed-region (8mer, 7mer-m8 and 7mer-1a). Multiple testing corrected Kolmogorov-Smirnov test was used to test for shift in cumulative distribution compared to genes that do not contain miR-218-5p binding sites.

To validate the successful uptake of LNAs via gymnosis, we injected fluorescently labeled LNAs (FAM-LNAs) into the somatosensory cortex I (SI) (Figure 3A). While we observed strong labeling of the SI and some spill-over into the hippocampal areas beneath the targeted SI region, fluorescence was not detected in more dorsal or ventral regions of the hippocampus and cortex (Suppl. Figure 4A). Higher magnification images confirmed the intracellular uptake of labeled LNAs in both cortical and hippocampal neurons (Suppl. Figure 4B).

For subsequent functional experiments, we stereotactically injected unlabeled scrambled negative control LNAs into the ipsilateral SI and unlabeled LNAs targeting either miR-129-5p or miR-218-5p (LNA129 and LNA218) into the contralateral SI region. Comparing miRNA levels using RT-qPCR in hemispheres injected with negative control LNAs to those injected with LNAs targeting miR-218-5p and miR-129-5p, respectively, confirmed miRNA-specific knockdown (0.38-fold change for miR-218-5p and 0.26-fold change for miR-129-5p; Figure 3B). In contrast, untargeted miRNAs remained unaffected by both LNA218 and LNA129 injections (Figure 3B).

Conducting bulk RNA-sequencing on hemispheres injected with LNA218 or negative control LNAs led to the identification of 35 differentially expressed genes under an FDR-cutoff of 0.05, with 13 upregulated and 22 downregulated genes (Figure 3C). *enrichMiR* analysis for miRNA-binding sites within the top 50 differentially expressed genes revealed a significant overrepresentation of miR-218-5p binding sites among upregulated genes, confirming the specific inhibition of miR-218-5p activity (Figure 3D). This conclusion was supported by the comparison of the cumulative distributions of log-fold changes of genes with predicted miR-218-5p binding sites versus those without. A pronounced rightward shift, pointing to upregulation, was observed in the distribution of genes containing miR-218-5p binding sites (Figure 3E).

In agreement, among the 13 most significantly upregulated genes, 9 possessed at least one strong binding site for miR-218-5p (Suppl. Figure 4C). In summary, in vivo LNA injections enabled us to identify strongly regulated genes targeted by miR-218-5p in cortical neurons.

Next, we focused on the 23 genes, among the top 50 upregulated ones, that contained miR-218-5p binding site predicted by *scanMiR* (40). We intersected this list with the predicted genes regulated by miR-218-5p during homeostatic synaptic downscaling (n=154) and identified an overlap of 6 genes (Diras1, Kcnh1, Lasp1, Mdga1, Sphkap, Zmiz2, Figure 4A). Notably, Mdga1, Lasp1, Kcnh1, and Diras1 emerged as promising candidates, as they have previously been implicated in various synaptic or neuronal activity functions. For instance, Mdga1 is associated with inhibitory synapse stability (41, 42), Lasp1 contributes to cytoskeletal organization at dendritic spines (43), Kcnh1 is a voltage-gated potassium channel involved in potassium channelopathies (44), and the GTPase Diras1 is involved in cholinergic signaling, neurodevelopment, and epilepsy (45).

**Figure 4:**
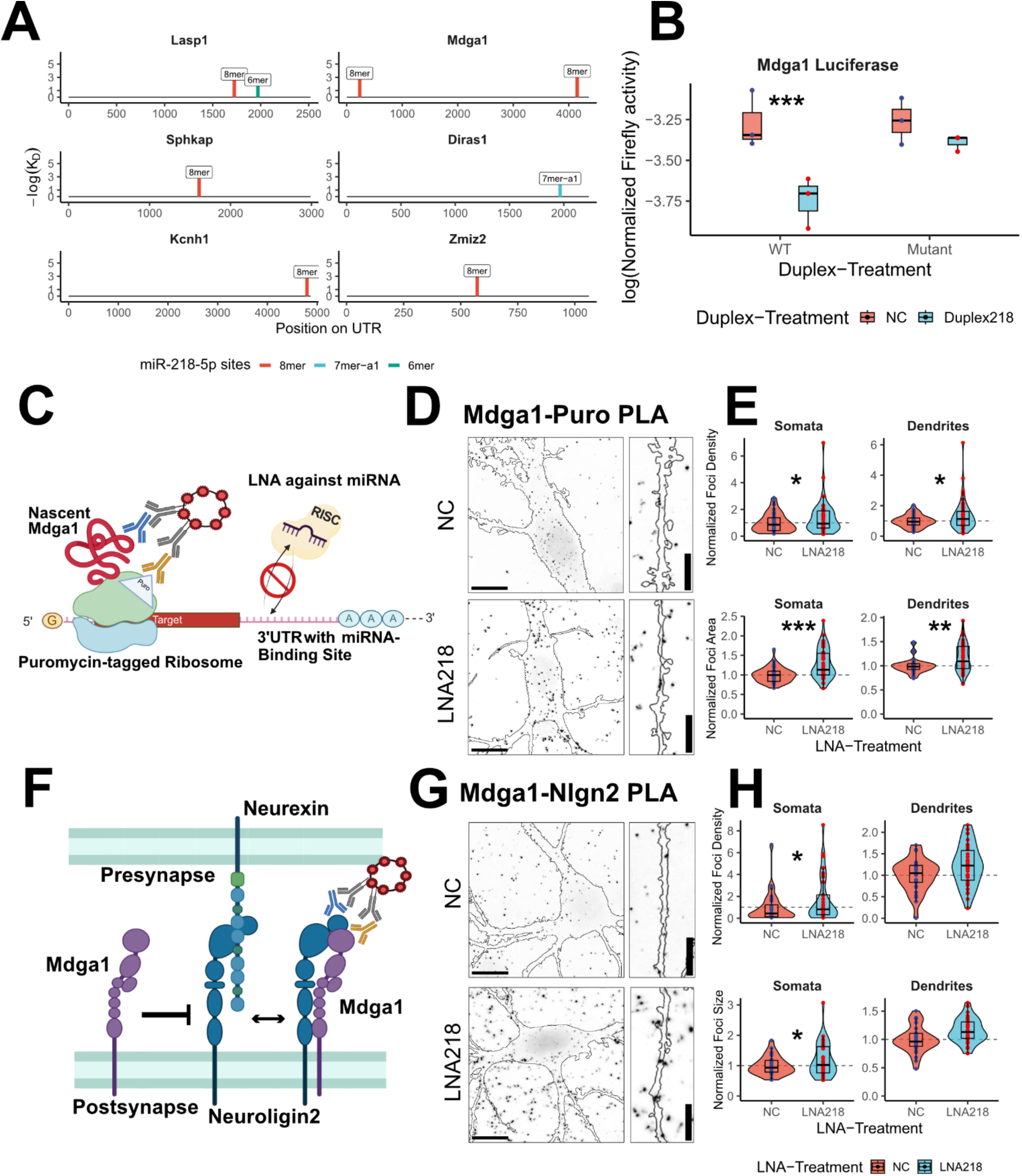
Negative regulator of inhibitory synapse stability Mdga1 is a direct target of miR-218-5p. (A) 3’ untranslated regions (3’UTRs) of genes that were upregulated by LNA218-treatment and downregulated by PTX-treatment containing miR-218-5p binding sites as predicted by *scanMiR* (denoted as 8mer, 7mer-m8 and 6mer seed regions). Figure 4 continued on next page (B) Luciferase assay of fragment of wildtype and mutant 3’UTR of Mdga1 upon miR-218-5p overexpression by Duplex218 or negative control duplex (NC). Values are ratio between Firefly-bioluminescence and transfection-control of Renilla bioluminescence, log2-transformed. n=3 independent biological replicates in primary cortical preparations. Data was fit to a linear-mixed model with fixed effects for Genotype (WT vs Mutant) and Duplex-treatment and their interaction and random effect of independent biological replicates. Dunn’s post-hoc analysis was used to estimate Duplex-effect in WT and mutant-conditions and to estimate Duplex218-effect compared to negative control within Genotype ***p<0.001. (C) Schematic of the puromycin proximity ligation assay (Puro-PLA) for the identification of Mdga1-translation foci (D) Example pictures of soma and dendrites of primary hippocampal neurons transfected with GFP (white outline) and either NC or LNA218 and stained for Mdga1-puromycin proximity ligation assay (PLA). White tool bars represent 5µm. (E) Quantification of translation foci cluster area (upper plot) and densities (lower plot) normalized to the mean of negative control (NC) group of the corresponding independent biological replicate of primary hippocampal preparations. 8-12 images per condition and per independent biological replicate n=4. Data was fit to a linear-mixed model with fixed effects for PTX-treatment and LNA-treatment and their interaction and random effect of independent biological replicates. Dunn’s post-hoc analysis was used to estimate LNA218-effect compared to negative control *p<0.05, **p<0.01, ***p<0.001. (F) Schematic of the Mdga1 competition with Neuroligin2 (Nlgn2)/Neurexin complexes. (G) Example pictures of soma and dendrites of primary hippocampal neurons transfected with GFP (white outline) and either NC or LNA218 and stained for Mdga1-Nlgn2 proximity ligation assay (PLA). White tool bars represent 5µm. (H) Quantification of translation foci cluster area (upper plot) and densities (lower plot) normalized to the mean of negative control (NC) group of the corresponding independent biological replicate of primary hippocampal preparations. 8-12 images per condition and per independent biological replicate n=4. Data was fit to a linear-mixed model with fixed effects for PTX-treatment and LNA-treatment and their interaction and random effect of independent biological replicates. Dunn’s post-hoc analysis was used to estimate LNA218-effect compared to negative control *p<0.05, **p<0.01, ***p<0.001.

We focused on Mdga1 for our further studies since it exhibited the most robust upregulation upon miR-218-5p inhibition and contained two notably strong binding sites at either end of its 3’UTR (Figure 4A). We assessed the direct interaction between these binding sites and miR-218-5p through a dual luciferase assay and observed that a fragment containing the first binding site (at position 246 bp of the 3’UTR) was sufficient to mediate a consistent downregulation in luciferase activity upon miR-218-5p overexpression (0.72-fold change in WT condition). Mutation of this binding site completely abolished miR-218-5p mediated downregulation of the reporter gene (Figure 4B), confirming that it was responsible for the inhibitory effect of miR-218-5p.

To validate the Mdga1 upregulation following miR-218-5p inhibition at the protein level, we employed *in situ* puromycin proximity ligation assay (Puro-PLA, Figure 4C (10, 46)) to assess the number and size of translation foci of nascent Mdga1 peptides (Figure 4D and E). We observed a higher number of translation foci in both the soma and dendrites of LNA218-transfected rat primary hippocampal neurons, along with larger translation foci indicating a potentially enhanced translation efficiency by polysomes (Figure 4E). Thus, miR-218-5p inhibits the translation of endogenous Mdga1 mRNA under basal neuronal activity conditions.

Mdga1 is a glycosylphosphatidylinositol (GPI)-anchored cell surface glycoprotein implicated in controlling inhibitory synapse stability via its interaction with the transmembrane protein Neuroligin2 (Nlgn2) (41, 47). Mdga1-mediated sequestration of Nlgn2 from its trans-synaptic partner Neurexin1 (Nrxn1) results in inhibitory synapse destabilization (Figure 4F). To ascertain whether the upregulation of Mdga1 through miR-218-5p inhibition leads to increased interaction with Nlgn2, we employed the PLA approach to assess the proximity of the two proteins (Figure 4F). Indeed, somatic coincidence foci were more abundant in number and size (Figure 4G-H). Although a similar trend was observed, no significant changes were detected in the number and size of PLA foci in dendrites, suggesting that miR-218-5p-dependent Mdga1 upregulation primarily impacts perisomatic inhibitory synapses containing Nlgn2.

### miR-218-5p is necessary for upscaling inhibitory synapses during prolonged network activity

Given Mdga1’s function as a negative modulator of inhibitory synapse stability, we hypothesized that miR-218-5p might be involved in the PTX-dependent upscaling of inhibitory synapses.

Considering the diverse subunit composition of GABAa receptors (GABAa-R), we initially monitored changes in the surface expression of clusters containing the gamma2-subunit, a key constituent of mature inhibitory synapses (Figure 5A, (48)). In alignment with existing literature (4, 5), our results demonstrated an increase in the size of somatic and dendritic gamma2 clusters following PTX-induced homeostatic synaptic scaling (1.20-fold and 1.39-fold increase, respectively, Figure 5B). Notably, inhibition of miR-218-5p efficiently abolished the upscaling of inhibitory synapses induced by PTX. Additionally, in analogy to our prior observations in the context of excitatory synapses, we detected a miR-218 dependent increase in cluster size in mock-treated neurons, implying that miR-218 negatively regulates inhibitory synapse size already under basal neuronal activity levels (1.24-fold and 1.38-fold change, Figure 5B) and that miR-218-5p inhibition occludes PTX-induced upscaling of inhibitory synapses.

Having shown that miR-218-5p occludes PTX-dependent upscaling of inhibitory synapses, we wanted to further explore whether this is also reflected at the level of Nlgn2, which directly interacts with the miR-218-5p target Mdga1. Thus, we conducted co-immunostainings of Nlgn2 in conjunction with vesicular GABA transporter (vGAT), a presynaptic marker specific to inhibitory synapses (Figure 5C). Our results show an enlargement in the somatic Nlgn2-cluster size during PTX-induced homeostatic synaptic plasticity (1.28-fold increase compared to mock-treated conditions, Figure 5D), while no significant alteration was observed in dendritic Nlgn2 clusters.

This observation might suggest that changes in Nlgn2-clusters predominantly occur in perisomatic inhibitory synapses during homeostatic synaptic plasticity. Consistent with the gamma2 findings, inhibition of miR-218-5p activity prevented PTX-induced upscaling of Nlgn2 clusters in the soma and resulted in larger somatic and dendritic Nlgn2 clusters under basal activity levels compared to the mock-treated negative control (1.30-fold and 1.16-fold changes in dendritic and somatic Nlgn2 clusters, respectively). Interestingly, a significant decrease in somatic Nlgn2-vGAT-co-cluster density was observed when miR-218-5p was inhibited under basal conditions (Suppl. Figure 5C). Somatic vGAT puncta size exhibited a moderate increase upon PTX treatment, while dendritic vGAT puncta displayed a trend towards upscaling (1.19-fold change in somatic and 1.14-fold change in dendritic vGAT puncta, Figure 5D). Interestingly, both somatic and dendritic vGAT cluster size upscaling was prevented upon inhibiting miR-218-5p activity.

Additionally, we extended our investigation to co-immunostainings of Gephyrin, a crucial postsynaptic scaffolding protein at inhibitory synapses, in combination with vGAT. Intriguingly, we observed PTX-induced upscaling of both dendritic and somatic Gephyrin and vGAT cluster sizes and co-cluster densities (Suppl. Figure 6A-C). However, inhibition of miR-218-5p activity only blocked the PTX-mediated upscaling of vGAT cluster size, but not Gephyrin. This suggests that miR-218-5p dependent regulation of inhibitory synaptic proteins, such as Mdga1, affects a specific subset of inhibitory synapses, e.g., those containing Nlgn2 and gamma2.

In summary, our comprehensive morphological investigations emphasize a significant involvement of miR-218-5p in homeostatic upscaling of inhibitory synapses, likely mediated through the modulation of Mdga1.

### Enrichment of miR-218-5p binding sites in oscillating Synaptic Transcripts

Given the evident involvement of miR-218-5p in the modulation of excitatory and inhibitory homeostatic synaptic scaling, we wanted to study potential neurophysiological implications. We decided to focus on sleep, since homeostatic synaptic scaling has been shown to contribute to memory consolidation during sleep (21, 49). To explore possible post-transcriptional regulation by miRNA during sleep-wake-cycles, we harnessed the extensive dataset provided by Noya et al. (30), which encompasses the synaptic transcriptome across a 24-hour cycle. This dataset involves synaptosomal RNA samples collected at six distinct timepoints (Zeitgeber time (ZT): 0, 4, 8, 12, 16, and 20).

From this dataset, we computed gene sets that were either specifically enriched in the light-or dark-phase (ZT0-4, 4-8 and 8-12 as light phase vs ZT12-16, ZT16-20 and ZT 20-0/24 as dark-phase) (Fig. 5A). Using this type of comparison, we caught rhythmic synaptic transcripts similarly described by Noya et al. (Figure 5A, Suppl. Figure 7A, B).

Next, we performed *enrichMiR* analysis specifically on the light-phase and dark-phase enriched gene sets (Figure 5B). Thereby, we detected an enrichment of binding sites for three miRNAs (miR-218-5p, miR-34-5p and miR-135-5p) in the light-phase and two miRNAs (miR-200b-ep, miR-6540-5p) in the dark-phase. Strikingly, all three of the miRNAs (miR-218-5p, miR-34-5p, miR-135-5p) whose binding sites are overrepresented in the light-phase gene set overlap with miRNAs that we previously identified in the context of PTX-mediated HSD *in vitro* (Figure 1C). Thus, miRNA-dependent regulation of HSP appears to be implicated in the control of oscillating synaptic transcripts, particularly at the light-dark phase transition.

### Altered Slow-Wave Activity during NREM Sleep Following miR-218-5p Inhibition

Since *enrichMiR* analysis suggested an important role of miR-218-5p-dependent regulation of oscillating synaptic transcripts, we decided to explore the functional impact of miR-218-5p inhibition on neuronal networks through the utilization of electroencephalography (EEG) measurements. EEG implantations in mice provide a means for extended recordings that capture putative local and/or global sleep EEG features in response to miR-218-5p inhibition during wake and sleep phases.

A long-term experiment was executed in mice, where we recorded local field potential (LFP) signals from the somatosensory cortex of both brain hemispheres: One hemisphere functioned as an internal control, while the other incorporated a cannula to allow for post-surgery miRNA-inhibitor delivery (Figure 6C). Leveraging the efficient gymnosis-based delivery of LNAs, we aimed to discern the effects of miR-218-5p inhibition on sleep EEG patterns. We first recorded a baseline day where mice were undisturbed in their sleeping chambers before we microinfused LNAs and continuously recorded EEG for 9 days to track changes of local EEG patterns over time. To validate effective miRNA inhibition, miRNA and target mRNA levels were quantified post-recordings within the SI cortical region using RT-qPCR. In all cases, efficient and specific miRNA knockdown was confirmed (0.48-fold expression of miR-218-5p in LNA injected hemisphere relative to the control hemisphere, Figure 6D). Accordingly, the miR-218-5p target mRNA Mdga1 was significantly upregulated (approximately 1.78-fold in LNA-injected hemisphere compared to the control hemisphere), while mRNA levels of the miR-218 target Shank2 showed a non-significant trend towards upregulation (approximately 1.29-fold, Figure 6D).

**Figure 5:**
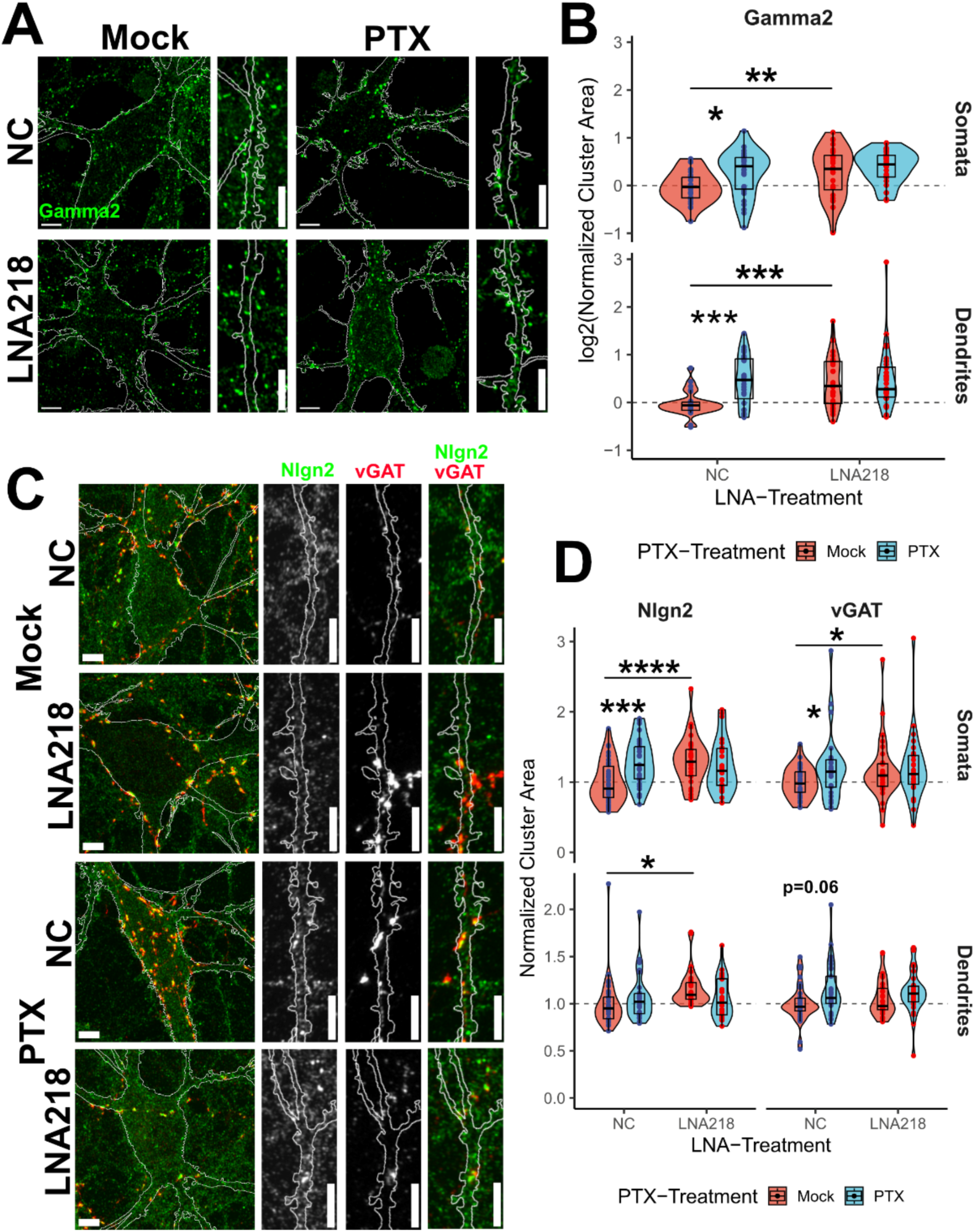
miR-218-5p inhibition occludes PTX-dependent upscaling of inhibitory synapses. (A) Example pictures of soma and dendrites of primary hippocampal neurons transfected with GFP (white outline) and either NC or LNA218 in combination with PTX– and mock-treatment and stained for surface GABA-receptor subunit Gamma2. White tool bar in cellbody inset represents 10µm and in the dendrite inset represents 10µm. (B) Quantification of cluster area of Gamma2 normalized to the mean of the mock-treated, negative control (NC) group of the corresponding independent biological replicate of primary hippocampal preparations. 8-12 images per condition and per independent biological replicate n=3. Data was fit to a linear-mixed model per compartment (soma or dendrites) with fixed effects for PTX-treatment and LNA-treatment and their interaction and random effect of independent biological replicates. Dunn’s post-hoc analysis was used to estimate PTX-effect in NC and LNA218-conditions and to estimate LNA218-effect compared to negative control within PTX-treatment *p<0.05, **p<0.01, ***p<0.001, ****p<0.0001. (C) Example pictures of soma and dendrites of primary hippocampal neurons transfected with GFP (white outline) and either NC or LNA218 in combination with PTX– and mock-treatment and stained for postsynaptic Nlgn2 and presynaptic vGAT. White tool bar in cellbody inset represents 10µm and in the dendrite inset represents 10µm. (D) Quantification of cluster area of Nlgn2 and vGAT normalized to the mean of the mock-treated, negative control (NC) group of the corresponding independent biological replicate of primary hippocampal preparations. 8-12 images per condition and per independent biological replicate n=3. Data was fit to a linear-mixed model per compartment (soma or dendrites) and per staining (Nlgn2 and vGAT) with fixed effects for PTX-treatment and LNA-treatment and their interaction and random effect of independent biological replicates. Dunn’s post-hoc analysis was used to estimate PTX-effect in NC and LNA218-conditions and to estimate LNA218-effect compared to negative control within PTX-treatment *p<0.05, **p<0.01, ***p<0.001, ****p<0.0001.

**Figure 6:**
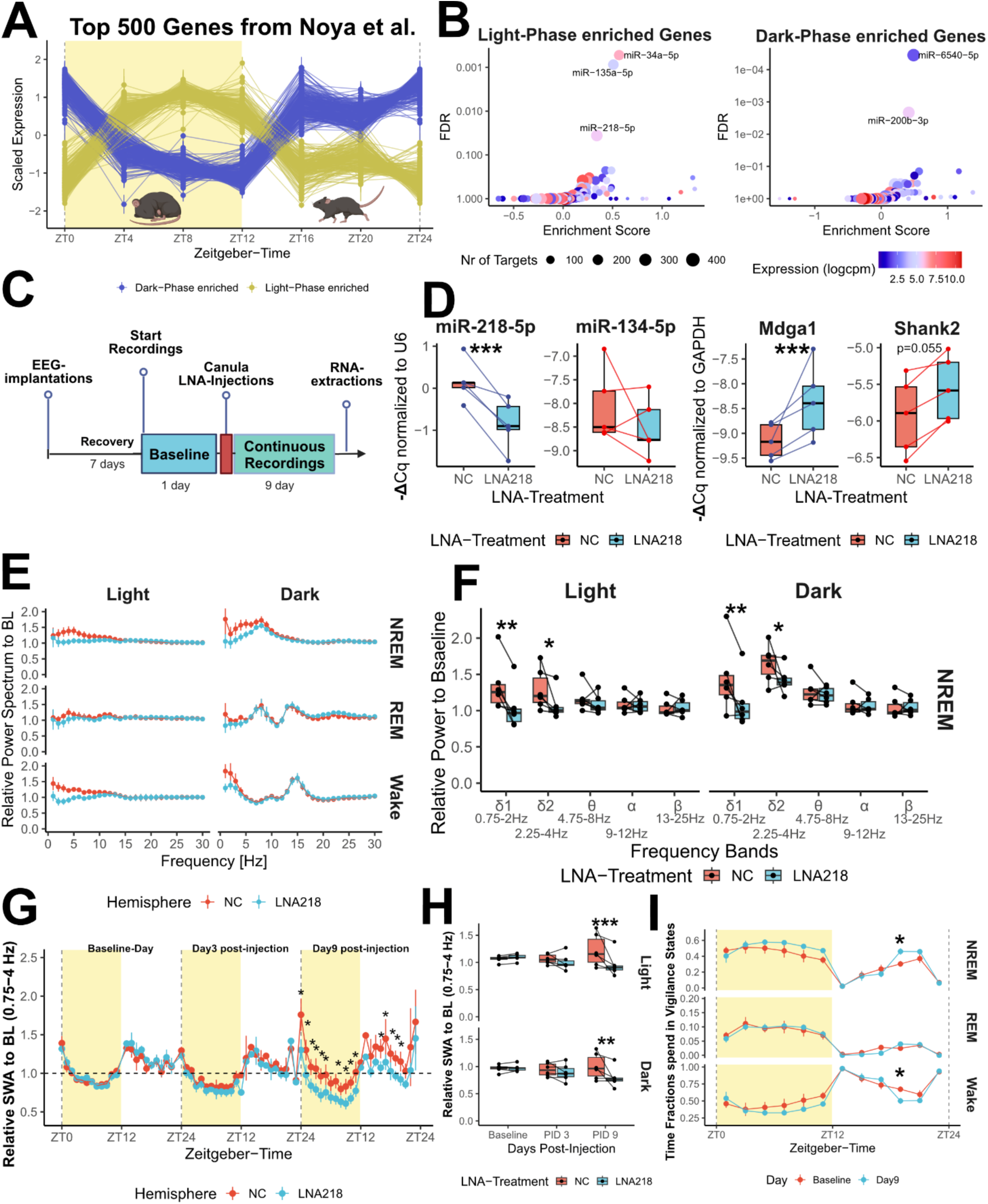
Slow-wave activity during NREM-sleep is reduced by miR-218-5p inhibition. (A) Top 500 Genes in synaptosomal preparations that are enriched either in the light (yellow shading) or dark phase in the dataset from Noya et al. (B) enrichMiR-analysis of light-phase (left) or dark-phase (right) enriched genes (C) Experimental workflow for the study of sleep EEG changes in response to local microinfusion of LNA218. (D) Validation of LNA-mediated knockdown by Taqman RT-qPCR in mice that underwent EEG-recordings (left plot). –ΔCq-values of miR-218-5p or miR-134-5p normalized to U6 comparing control hemispheres injected with LNA-negative control (NC) or LNA against miR-218-5p (LNA218). n=5 mice. Data was fit to a linear-mixed model per miRNA with fixed effects for LNA-treatment and random effect of independent biological replicates (mice). Dunn’s post-hoc analysis was used to estimate LNA-effect on hemispheres injected with LNA218 compared to corresponding control hemisphere within mouse. Validation of LNA-mediated upregulation of target genes Mdga1 and Shank2 by RT-qPCR in mice that underwent EEG-recordings (right plot). –ΔCq-values of Mdga1 or Shank2 normalized to GAPDH comparing control hemispheres with LNA against miR-218-5p (LNA218). n=5 mice. Data was fit to a linear-mixed model per target-mRNA with fixed effects for LNA-treatment and random effect of independent biological replicates (mice). Dunn’s post-hoc analysis was used to estimate LNA-effect on hemispheres injected with LNA218 compared to corresponding control hemisphere within mouse. **p<0.01, ***p<0.001 (E) Power spectrum (0.5-25 Hz) in different vigilance states in the light and dark phase of postinjection-day 9, relative to baseline day. (F) Frequency-binned relative power in NREM sleep at postinjection-day 9, relative to the average NREM power in baseline day (ZT0-12). Data was fit to a linear-mixed model with fixed effects for LNA-treatment and Frequency-bands and random effect of independent biological replicates (n=6 mice). Dunn’s post-hoc analysis was used to estimate LNA-effect on hemispheres injected with LNA218 compared to corresponding control hemisphere within mouse. *p<0.05, **p<0.01 (G) Relative NREM sleep SWA time course at baseline, postinjection-day 3, and postinjection-day 9. Data was fit to a linear-mixed model with fixed effects for LNA-treatment and Zeitgeber-time and random effect of independent biological replicates (n=6 mice). Dunn’s post-hoc analysis was used to estimate LNA-effect within time-points on hemispheres injected with LNA218 compared to corresponding control hemisphere within mouse. *p<0.05 (H) Time-course of relative SWA (0.75-4Hz) in the light and dark phase of postinjection-day 9. Data was fit to a linear-mixed model with fixed effects for LNA-treatment and Days Post-Injection and random effect of independent biological replicates (n=6 mice). Dunn’s post-hoc analysis was used to estimate LNA-effect within Days Post-Injection on hemispheres injected with LNA218 compared to corresponding control hemisphere within mouse. **p<0.01, ***p<0.001 (I) Time course of time spent in vigilance states (wake, NREM– and REM-sleep) with 2 hours bins at postinjection day 9. Data was fit to a linear-mixed model with fixed effects for LNA-treatment and Zeitgeber-time and random effect of independent biological replicates (n=6 mice). Dunn’s post-hoc analysis was used to estimate LNA-effect within time-points on hemispheres injected with LNA218 compared to corresponding control hemisphere within mouse. *p<0.05

Regarding EEG power spectral analysis, local inhibition of cortical miR-218-5p resulted in a significant reduction in slower frequencies (0.75-4Hz) during NREM and wake states, whereas higher frequencies (4.75-25Hz) showed no difference in comparison to contralateral hemispheres (Figure 6E and Figure 6F, Suppl. Figure 7C). In contrast, frequencies during REM-sleep remained unchanged between hemispheres (Figure 6E and Suppl. Figure 7C).

A more time-resolved comparison showed a gradual transition if we compare time courses of normalized SWA (0.75-4Hz) at baseline-day, post-injection day 3 (PID3) and postinjection day 9 (PID9; Figure 6G and H). While we identified a marked reduction in SWA at PID9, we could not detect any changes at PID3 between LNA218-injected hemispheres. At a systemic level, local unihemispheric cortical inhibition of miR-218-5p was sufficient to increase time that mice spent in NREM sleep during the last hours of the dark phase (ZT18-20, Figure 6I).

In conclusion, this study enabled real-time monitoring of sleep pattern alterations following local miR-218-5p inhibition. During NREM-sleep, we identified marked reduction in local SWA in hemispheres that were injected with miR-218-5p inhibitors as well as increased sleep duration during the active phase (dark phase). These results suggest that molecular regulation by miR-218-5p has a pivotal impact on the coordination of synchronous neuronal firing during sleep.

## Discussion

Our study provides multiple lines of evidence which suggest a role of miR-218-5p in orchestrating both excitatory and inhibitory homeostatic synaptic plasticity. Upregulated miR-218-5p mediates downregulation of direct targets like Shank2, which are involved in the control of excitatory synaptic strength. Concurrently, miR-218-5p dependent downregulation of Mdga1, a negative regulator of inhibitory synapses, leads to the disinhibition of inhibitory synapses, resulting in an overall rise in inhibitory drive. To our knowledge, miR-218-5p represents the first example of a miRNA regulating homeostatic inhibitory upscaling during PTX-mediated HSP. Furthermore, the simultaneous control exerted by miR-218-5p over excitatory and inhibitory synaptic strengths proposes a concerted mechanism through which neurons adapt dynamically to augmented neuronal activity, selectively diminishing excitation while strengthening inhibition. Additionally, we propose that this mechanism holds physiological significance in coordinating sleep-associated homeostatic synaptic plasticity.

Utilizing confocal microscopy, we have established that miR-218-5p is essential for mediating the downscaling of pivotal glutamatergic AMPA-R subunits, GluA1 and GluA2, at postsynaptic densities (50). Intriguingly, inhibition of miR-218-5p did not influence the densities of GluA2/GluA1 clusters, implying that miR-218-5p specifically modulates the strength of individual synapses rather than the formation/elimination of synapses during HSP. This distinction is reflected by our observation that the suppression of miR-218-5p activity occludes the reduction of spine volume upon PTX treatment.

It has been previously reported that overexpressing miR-218-5p activity results in upregulation of luciferase reporters containing the 3’UTR of GluA2 (35). While the study did not assess whether this regulation was mediated by activity, it also reported an increase in mEPSC when miR-218-5p is upregulated, which is in line with our observations that miR-218-5p is a positive regulator of excitatory synaptic strength under basal conditions.

However, given that several of the predicted targets of miR-218-5p during HSP engage diverse facets of excitatory synapse function and dendritic spine morphology, we favor a model whereby miR-218-5p downregulates AMPA-Rs via the control of interacting proteins. For example, predicted miR-218-5p targets participate in intracellular signaling (Prkar2b, Srcin1, Camkk2, mGluR1, etc., see Table S2), cytoskeletal reorganization at dendritic spines (Add2, Lasp1, Myo16), and orchestration of AMPA-R subunit scaffolding at the postsynaptic density (Shank2). Given the intricate interplay among these diverse targets, it is plausible that miR-218-5p-mediated co-regulation of dendritic spine morphology and AMPA-R subunit surface expression mutually impact each other during activity-driven homeostatic synaptic downscaling.

Furthermore, we investigated the impact of miR-218-5p inhibition on PTX-induced homeostatic inhibitory synaptic upscaling. Our findings substantiate the necessity of miR-218-5p activity for the upscaling of two independent structural markers of inhibitory synapses (Gamma2 and Nlgn2-vGAT clusters). Analogous to their excitatory counterparts, alterations in cluster size rather than density were observed, reinforcing the notion that miR-218-5p regulates synaptic strength rather than stability of inhibitory synapses. Notably, the dispensability of miR-218-5p for Gephyrin-cluster upscaling indicates its importance for specific aspects of inhibitory synapse strength.

Mechanistically, miR-218-5p-mediated downregulation of the direct target Mdga1 might lead to a release of Nlgn2, thereby enabling its interaction with transsynaptic neurexins and reinforcing pre– and postsynaptic clustering of vGAT and Nlgn2-positive terminals. Given the predominant upscaling of inhibitory synapses onto neuronal somas, we hypothesize that Nlgn2-positive synapses favor a perisomatic localization. Recent studies suggest that Mdga1 plays also an important in regulating excitatory synaptic strength (42, 51). Knocking down Mdga1 results in a decrease in AMPA-R mediated synaptic currents, which suggests a dual role of Mdga1 during HSP (51). Since Mdga1 is a fast-diffusing membrane-bound protein, it might be interesting to study how its synaptic localization dynamics are changing during HSP.

The sleep-induced synaptic changes that restore synaptic strength share common mechanistic characteristics with homeostatic synaptic plasticity *in vitro*. A prominent example represents Homer1a-mediated GluA1 internalization, which occurs during sleep (21) and bicuculline-induced homeostatic synaptic downscaling (38, 52). Our findings of a common regulatory gene signature enriched in miR-218-5p binding sites among genes downregulated by PTX treatment and transcripts enriched during the light phase lend further support for the relevance of homeostatic plasticity mechanisms during sleep. Since miRNAs typically exerts negative regulatory effects on gene expression, one way to interpret the enrichment of binding sites in up-regulated genes is a decrease in miRNA-activity and the enrichment in down-regulated genes as an increase in miRNA-activity. Conversely, we can also take into account the buffering and fine-tuning effect of miRNA in regulating gene expression in interaction with their target mRNAs (53, 54). In this respect, we can also interpret the enrichment of miRNAs as an increased need for fine-tuning regulation during these crucial phases of the day.

Our EEG experiments revealed depleted SWA levels in cortical networks in which miR-218-5p activity was consistently suppressed. Given that sleep SWA, predominant during NREM sleep, entails synchronous cortical neuron firing in the thalamocortical ensembles (55), it has been speculated on the role of this firing pattern in promoting downscaling of excitatory and possibly upscaling of inhibitory synapses (20, 29, 56). While miR-218-5p inhibition disrupts activity-driven homeostatic synaptic plasticity *in vitro* and adjusted baseline set-points of inhibitory and excitatory synaptic strength, we can hypothesize that similar synaptic alterations *in vivo* result in disturbed coordination of neuronal firing during sleep, leading to the observed reduction of SWA. Electrophysiological recordings using multi-electrode array recordings of cortical neuronal assemblies *in vivo* and whole-cell patch clamp to probe for synaptic strength *ex vivo* and around the clock could provide further insights into the exact mechanism of how miR-218-5p regulates NREM-sleep SWA. Furthermore, we report increased time spent in NREM-sleep towards the end of the dark phase (ZT18-20), which is in line with recent reports indicating a role of cortex in sleep-wake regulation (57). Since wildtype mice show an increased propensity to sleep towards the end of the dark phase, a behavior coined as afternoon nap or siesta (58), this finding could be interpreted as a compensatory mechanism to extend sleep and recover from disrupted local NREM-sleep SWA across sleep bouts.

Recently, it has been reported that low expression of miR-218-5p in the medial prefrontal cortex (mPFC) of mice correlated with susceptibility to stress in a chronic social defeat model of depression (59). Furthermore, inhibition of miR-218-5p resulted in increased susceptibility to stress, whereas upregulation of miR-218-5p conferred resilience. These findings suggest a potential involvement of miR-218-dependent homeostatic synaptic plasticity in depression.

Indeed, it has been proposed that lithium, a mood stabilizer used for the treatment of bipolar disorder, might confer its therapeutic effects by promoting homeostatic synaptic downscaling through Gsk-3β activation and elevation of BDNF-levels (60). BDNF, a neurotrophic factor that has been implicated in many aspects of synaptic plasticity, sleep and depression (61), is consistently upregulated by PTX-induced homeostatic synaptic plasticity in our datasets (10). Furthermore, it has been reported that BDNF stimulation of primary hippocampal neurons promotes miR-218-5p as well as miR-132-5p packaging into extracellular vesicles (62). These findings position miR-218-5p as an ideal candidate regulator of homeostatic synaptic plasticity in the context of neuropsychiatric disorders.

Furthermore, a recent study highlights how miR-218-5p is important during neurodevelopment for proper hippocampal assembly of interneuron and pyramidal neuron networks (63). Prenatal inhibition of miR-218-5p leads to heightened network activity as well as a predisposition for epileptic seizures later in life. These findings are supported by miRNA *in situ* hybridization experiments in human post-mortem tissue of patients suffering from mesial temporal lobe epilepsy which show decreased levels of miR-218-5p in hippocampal neurons (64). Since homeostatic mechanisms could become maladaptive during epilepsy (65), future experiments are needed to address whether aberrant miR-218-5p regulation during HSP is linked to epileptogenic neuronal activity.

In conclusion, our study identified miR-218 as a coordinator of inhibitory and excitatory synaptic strength during homeostatic synaptic plasticity, with potential implications for sleep, as well as neurological and neuropsychiatric disorders.

## Materials and Methods

### DNA Constructs and Plasmids

Wild-type and mutant Shank2 and Mdga1 3′ UTR luciferase constructs were generated by standard cloning procedures into pmiRGLO dual-luciferase expression vector (Promega). First, fragments of the 3’ UTRs were PCR-amplified from mouse complementary DNA (cDNA) using primers described in the table below into pBluescript(+)-plasmid. Mutagenesis was then performed using site-directed mutagenesis using Phusion Hotstart High-Fidelity Polimerase II (ThermoFisher) using primers described in the table below. WT and mutagenized 3’UTR were then cut and inserted into pmiRGLO using the restriction sites NheI and SalI. Bacterial clones were screened for mutations by Sanger sequencing.

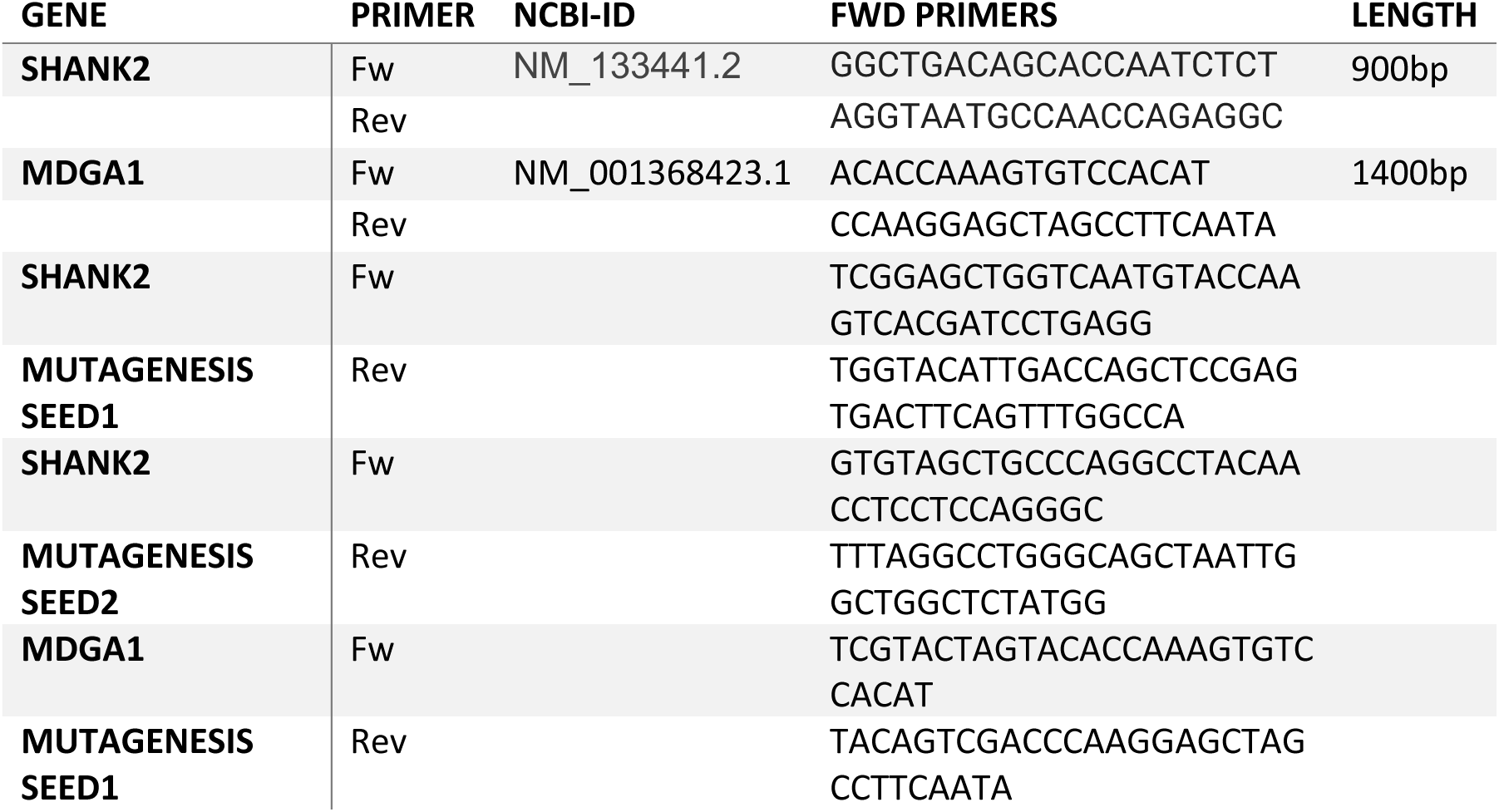

### Primary Hippocampal Neurons and Pharmacological Treatments

Cultures of primary cortical and hippocampal neurons were established using Sprague-Dawley rat embryos at embryonic day 18 (E18), following previously published protocols (61). The ethical use of pregnant rats for embryo brain collection was authorized by the Veterinary Office of the Canton Zurich, Switzerland, under license ZH027/2021. Dissociated cortical neurons were directly plated on 24-well coverslips coated with poly-L-ornithine for luciferase assay experiments, while hippocampal neurons were seeded on poly-L-lysine/laminin-coated coverslips in 24-well plates for immunofluorescence and in-situ proximity ligation assays.

For creating compartmentalized cell cultures, dissociated hippocampal cells were plated on 1-μm pore and 30-mm diameter polyethylene tetra-phthalate (PET) membrane filter inserts (Millipore). These inserts were matrix-coated with poly-L-lysine and laminin on both the upper and lower surfaces, as previously described (10, 62). All neuronal cultures were maintained in Neurobasal-plus medium supplemented with 2% B27, 2 mM GlutaMAX, 100 μg/ml streptomycin, and 100 U/ml penicillin (all ThermoFisher) in a 37°C incubator with 5% CO2.

Primary neuronal cultures were fed 60% fresh supplemented Neurobasal-plus medium every 3-4 days. In order to induce activity-dependent homeostatic synaptic plasticity, primary hippocampal neuronal cultures were treated with either 100µM picrotoxin (Sigma-Aldrich) dissolved in ethanol or equivalent volume of absolute ethanol (mock-treatment) for 48 hours at DIV19 to 21, according to established protocols (10, 15–17)

### Transfections

Transfections were performed using Lipofectamine2000 (Invitrogen) on duplicate or triplicate coverslips at DIV9 or 12. A total of 1 µg DNA was applied per coverslip, with experimental plasmids (100-150 ng) supplemented by an empty pcDNA3.1 carrier plasmid to reach the total 1 µg DNA quantity.

In brief, Lipofectamine2000 was preincubated for 5-10 minutes in plain Neurobasal-plus medium without supplements. This mixture was then combined with DNA constructs and incubated for 25 minutes to form DNA-lipofectamine complexes. After incubation, the complexes were diluted 1:5 in plain Neurobasal-plus medium and introduced to primary neurons for a 2-hour incubation.

Following two washes, neurons were treated with APV-supplemented Neurobasal-plus medium (1:1000) for 45 minutes, subsequently washed, and replaced with conditioned medium composed of 60% old and 40% fresh medium.

### Dual Luciferase Assay

Primary cortical neurons were transfected in triplicates with 100ng of pmiR-Glo-plasmids containing wildtype or mutant 3’UTRs of Mdga1 or Shank2 together with pre-miR-218-5p duplex (0.3125pmol for Mdga1 and 1.25pmol for Shank2, Ambion Pre-miR-218-5p Precursor) or respective quantities of pre-miR negative control duplex (Ambion) at DIV9. At DIV12 cells are lysed in Passive lysis buffer (diluted 1:5 in MilliQ-water, Promega) for 15min and dual luciferase assay performed using home-made reagents according to Baker and Boyce (63) on the GloMax Discover GM3000 (Promega).

For analysis, bioluminescence readings of the Firefly luciferase are normalized to control bioluminescence readings of the Renilla luciferase as an internal transfection control. Resulting ratios are then log-transformed.

### Spine Analysis and Immunofluorescence

For spine analysis, primary hippocampal neurons were transfected in duplicates or triplicates using 150ng of a plasmid carrying enhanced GFP-gene in combination with 5nmol Power Lock-Nucleic Acid inhibitors (Qiagen) against miR-218-5p, miR-129-5p, miR-34a-5p or miR-34c-5p at DIV12. After PTX-treatment from DIV19 to 21, cells were fixed in 4% PFA dissolved in 4%-sucrose PBS (fixation solution) solution for 30min at room temperature, rinsed and mounted using Aqua-Poly/Mount (Polysciences) mounting medium onto microscopy glass slides.

For immunofluorescence, cells are fixed in fixation solution for 15min, permeabilized in 0.1%-Triton-X dissolved in 10% normal goat serum (NGS, Invitrogen) and PBS for 10min and incubated in primary antibody solution for 1 hour (antibodies diluted in 10% NGS-PBS solution, see table of antibodies for dilutions and brand). After 3 times 7minutes washing steps in PBS, cells are incubated in fluorescently labelled secondary antibody solution (antibodies diluted in 10% NGS-PBS solution, see table of antibodies for dilutions and brand). After 3 times 7minutes washing steps in PBS and Hoechst counterstaining (1:10000) in the first washing step, coverslips were mounted using Aqua-Poly/Mount (Polysciences) mounting medium onto microscopy glass slides. Stained and fixed coverslips were kept at 4°Celsius until further use for confocal imaging. For surface stainings of Gamma2 or GluA2 / GluA1 co-stainings, primary hippocampal neurons were incubated in neuronal medium containing primary antibodies for 1 hour at 37°C in the incubator. After 3×5min washing steps with pre-warmed, fresh supplemented Neurobasal-plus medium and 2 quick rinses with room-temperature PBS, neurons were fixated in fixation buffer for 15min. After fixation and rinsing with PBS, cells were treated for immunofluorescence as described earlier except by omission of the permeabilization step.

### Duolink Proximity Ligation Assay

Translation foci detection involved using an antibody against puromycin after ribosomal metabolic labeling with puromycin and an antibody against the target nascent peptide, following established methods (tom Dieck et al, 2015). Utilizing the Duolink in situ orange PLA mouse/rabbit kit (Sigma), these two antibodies were spatially coincided with high precision (< 40 nm) through rolling circle amplification with fluorescently labeled oligonucleotides.

For the procedure, primary hippocampal neurons were cultured on coverslips and treated with 100 nM PTX for 48 hours. Before fixation, cells were exposed to puromycin (3 μM final concentration, Invivogen) for 5 minutes or left untreated for control conditions. After brief rinses with PBS, the cells were fixed in 4% sucrose-PFA for 10 minutes at room temperature, followed by permeabilization using a buffer containing 0.05% Triton X and 10% normal goat serum in PBS for 5 minutes. The Duolink Proximity Assay (Sigma) was executed as per the manufacturer’s protocol. The coverslips were positioned on parafilm in a wet chamber. Cells underwent blocking with the provided buffer at 37°C for 1 hour, followed by incubation with the primary antibody solution (mouse anti-puromycin, rabbit anti-peptide of interest, and chicken anti-MAP2 in the supplied antibody diluent) at room temperature for 1 hour. Subsequent steps included three 5-minute washes in washing buffer A (Duolink kit’s supplied), followed by incubation in secondary antibody solutions (anti-rabbit Plus-probe, 1:10, anti-mouse Minus-probe, 1:10, and anti-chicken Alexa Plus 647-secondary antibody, 1:1,000, Thermo Fisher Scientific) at 37°C for 1 hour. This was followed by a ligation step (30 minutes at 37°C) and amplification step (100 minutes at 37°C) as per the manufacturer’s protocol. The final washing steps involved three 10-minute washes with the supplied washing buffer B, followed by brief nuclear counterstaining using Hoechst (1:10,000) in PBS and mounting on a glass slide with Aqua-Poly-Mount (Polysciences).

For co-localization assay between Mdga1 and Nlgn2, proximity ligation assay was performed as described above with omission of puromycin treatment. Instead, primary mouse antibody against Nlgn2 (Synaptic Systems) and rabbit anti-Mdga1 (Synaptic System) was used.

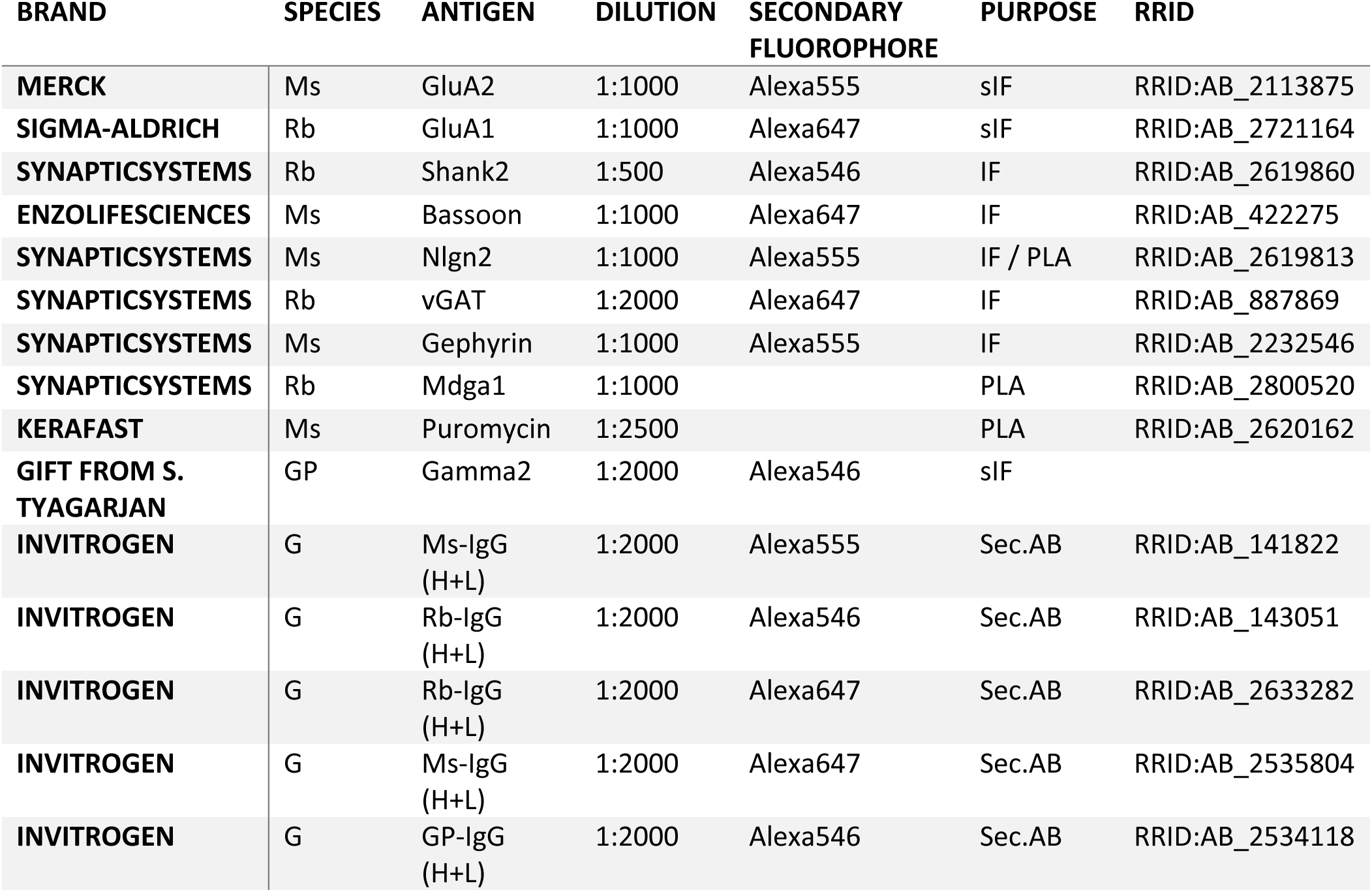

### Antibodies

Table 2: Antibodies used in the study. MS = Mouse; Rb = Rabbit; GP = Guinea Pig; G=Goat; sIF= surface immunofluorescence; IF = immunofluorescence; PLA = Promoximity Ligation Assay; Sec.AB = Secondary antibody

### Image Acquisition and Image Analysis

All images were acquired on a AiryScan Zeiss LSM 800 in Fast Airyscan optimal high-resolution acquisition mode (11.73 pixels / µm in the xy dimensions and Z-stacking at an interval of 0.18 µm per slice) using a 63× 1.4 NA oil objective (Plan-Apochromat 63×/1.4 Oil DIC M27). GFP signal was used to assess morphology and health of cells and selected according to closest resemblance to pyramidal neurons (developed apical and basal dendritic arbor containing dendritic spines). Laser power and gain was optimized on mock treated negative control conditions and kept constant across experimental replicate.

The openly available plugin Cluster Analysis containing custom python-scripts written in the Fiji-imaging framework (64, 65) were used to quantify confocal microscopy pictures. Installation instructions are available on the github repository of the plugin (https://github.com/dcolam/Cluster-Analysis-Plugin).

Spine Analysis: Our automated spine analysis workflow builds upon established manual methods (18, 19, 36). Consistency was ensured by applying the following empirically determined parameters across all experiments and conditions. In brief, the channel containing the GFP signal is transformed into a binary representation using the minimum cross-entropy thresholding method (also known as Li threshold in ImageJ, (41)), outlining the entire neuron. The resulting mask undergoes skeletonization, and spines branches are identified using a length filter ranging from 0.1 to 5µm. Spine heads are localized by creating a region of interest (ROI) with a 0.5 µm radius at the tip of the skeletonized spine branch. The spine head outlines are segmented by overlaying the resulting circular ROIs onto the original binarized GFP image and applying a size filter of 0.1 to 5µm². The resultant outlines of the spine heads are used to calculate the mean pixel intensity by superimposing them onto the original unbinarized GFP channel. The mean pixel intensity of the spine heads is then normalized against the mean intensity of the entire segmented neuron. Synaptic Cluster Analysis: Co-localization analysis of pre– and postsynaptic (or in the case of GluA1/Glu2 immunostainings only postsynaptic) stainings was accomplished by the automatic segmentation tools provided by the plugin utilizing different neuronal compartments visualized in the GFP channel. The segmentation of the soma was accomplished by taking advantage of the brighter somatic GFP-signal. The minimum thresholding method in combination with a Gaussian Blurring ensured the proper segmentation of the soma with a size filter of 50-500µm². Inverting the resulting somatic mask allowed to segment dendrites. In addition to the resulting somatic and dendritic ROIs, a segmentation of the whole neuron was used to quantify pre– and postsynaptic puncta. Within these ROIs, channels containing the pre– or postsynaptic markers are binarized using the Otsu or Triangle thresholding method implemented on ImageJ and clusters filtered and segmented according to an area filter of 0.25-2.5µm². Due to generally better signal-to-noise ratio in the presynaptic marker channels (vGAT or Bassoon, GluA2 for surface AMPA-receptor stainings), segmented presynaptic clusters were superimposed onto the postsynaptic channel and coincidences of signals counted and segmented. Parameters like area and mean-intensity are then measured in the pre– and postsynaptic channels within the co-localized segmentations. Furthermore, co-localized synaptic densities were estimated by dividing the number of co-localized particles by the area in the corresponding segmentation.

### Mouse Husbandry

Animals were housed in groups of 3–5 per cage, with food and water ad libitum. All experiments were performed on adult male mice (3 months old). The animal house had an inverted light-dark cycle, all testing was done during the dark phase (9 a.m. to 9 p.m.). All animal experiments were performed in accordance with the animal protection law of Switzerland and were approved by the local cantonal authorities (ZH194/21).

### Stereotactic Injections

Stereotactic brain injections were performed on 3-month-old C57BL/6JRj wild type mice. Isoflurane anesthetized mice received subcutaneous injection of 5 mg/kg of analgesic Meloxicam and subsequently injected bilaterally with 0.5 µl of 300µM of miRCURY Power LNA miRNA Inhibitors (negative control or against miR-218-5p, test injections with 5’FAM-labeling) into trunk primary somatosensory cortex SI (coordinates from bregma: anterior/posterior −1.6 mm, medial/lateral ± 1.6 mm, and dorsal/ventral −0.9 mm). Lidocain (<2mg/kg)/Bupivacain (2mg/kg) solution was distributed on wound site before suturing the skin. For analgesia treatment, the animals received another subcutaneous injection of 5 mg/kg of Meloxicam 6-8 hours after surgery and paracetamol were added to the drinking water for 48 hours post-surgery. Postoperative health checks were carried on over the 3 days after surgery.

### RNA extractions

RNA from brain tissue was extracted using the RNA-Solv reagent kit (Omega Bio-tek) according to manufacturer protocol. RNA from compartmentalized cultures was extracted using the mirVana™ total RNA Isolation Kit. Removal of genomic DNA was accomplished by Turbo DNase treatment (ThermoFisher). RNA samples were stored at –80°C until further use.

### Quantitative Real Time PCR

RNA was reverse transcribed using the iScript cDNA Synthesis Kit (Bio-Rad) for mRNA-detection or Taqman MicroRNA Reverse Transcription Kit (Thermo Fisher Scientific) for miRNA-detection according to manufacturer protocol. Quantitative real-time PCR was performed on the Step One Plus Real-Time PCR System (Applied Biosystems) using iTaq SYBR Green Supermix with ROX (Bio-Rad) for mRNA detection or Taqman Universal PCR Master Mix (Thermo Fisher Scientific). Quantification cycle (Cq) was normalized to either Cq-values of U6 for miRNA or GAPDH for mRNA. Statistics and plotting were kept in the log-scale and directly performed from the delta Cq values.

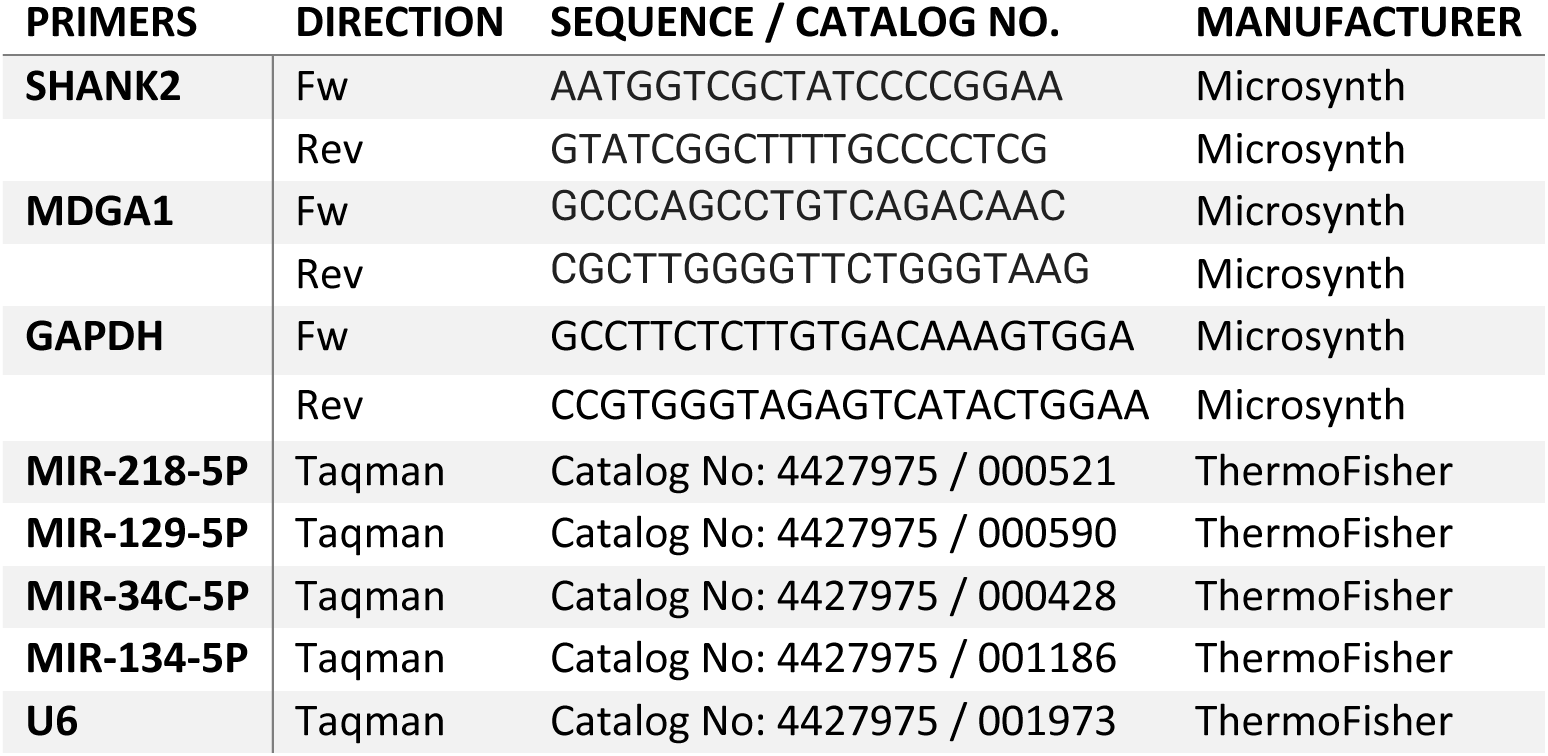

### PolyA RNA-sequencing

200ng of DNase-treated RNA per sample from mouse cortical regions were sent for sequencing to Novogene (London) who performed quality control, mRNA library preparation by PolyA enrichment and paired-end sequencing on an Illumina Novaseq 6000 sequencer at a depth of 6 million reads per sample.

### Bioinformatics Analysis

General workflow of differential expression analysis consisted of genome alignment using Salmon (66), count-filtering, surrogate variable analysis using the R-package sva (67), normalization and model fitting using edgeR (68). Custom scripts were written using R-markdown, which are available upon request. Below we described the specific models for the different RNA sequencing datasets used in the study:

LNA-Sequencing: Paired-end reads were aligned to the mouse genome assembly GRCm39 and quantified using Salmon (66). Prior to differential expression analysis, surrogate variable analysis (sva) was used to estimate and correct technical batch effects (67). After count filtering and normalisation, edgeR (68) was used to fit a gene wise negative binomial generalized linear model (glm-fit) in the form of ∼ LNA-Treatment + SV1 + SV2 (where SV stands for surrogate variable) and contrasted for differences between LNA218-treated and negative control treated conditions. PTX-Sequencing: Reads generated by our previous study (10) was re-aligned to the newest rat genome assembly mRatBN7.2 using Salmon. After sva, filtering and normalization, a glm-fit model in the form of ∼ PTX-Treatment X Compartment + SV1, where mock-treated, somatically localized conditions constituted the intercept, was fitted to contrast gene wise responses to main-effects of PTX-treatment, main-effect of Compartment and their interaction-term. We calculated the adjusted log-fold-change of process-compartment specific PTX-effects by summing up the log-fold-change of the main-effect of the PTX-treatment with the log-fold change of the interaction-term. Adjusted log-fold changes were used for downstream analyses such as enrichMiR-analysis.

Synaptic transcriptome: The dataset from Noya et al. (30) was aligned to the mouse genome assembly GRCm39. The dataset is comprised of conditions described by localization (Synaptosomal or homogenate origin), time of collection (Zeitgeber-time (ZT) 4, 8, 12, 16, 20, 0/24) and sleep deprivation (control mice or sleep-deprived 4 hours prior brain collection). For simplicity reasons, we dealt for our analyses only with synaptosomal– and sleep deprivation conditions.

After standard workflow described above, we fitted a glm-model in the form of ∼ ZT-effect x Sleep-deprivation-effect + SV1 + SV2 + SV3. In addition to standard contrasting for every variable, we constructed a contrast in the form of (Control-ZT4 + Control-ZT8 + Control-ZT12) – (Control-ZT16 + Control-ZT20) to catch differences between light-phase time-points (ZT4-ZT12) and dark-phase time-points (ZT16-ZT24, note that ZT24 is the intercept and does not need to be specified in the contrast). The same contrast design has been specified using sleep-deprived conditions.

### EnrichMiR

The BioConductor R-package enrichMiR (32) enables enrichment analysis of miRNA-binding sites using sets of genes or down– and upregulated genes calculated by differential expression analysis. In short, a collection of miRNA-binding sites (MBS) within 3’UTRs of genes such as Targetscan8 (69, 70) or scanMiR (39) is used to identify genes that contain specific MBS. Exact Fisher-test allows to identify overrepresentation of specific MBS in sets of genes that are up– or downregulated upon a specific treatment compared to genes that are not changing. For our purposes, we always used the Targetscan8 collection for mouse or rat, depending on dataset, and while the package supports different enrichment tests, we employed the sideoverlap-test for up– or downregulated genes.

### Gene Ontology Analysis

Gene Ontology Analysis was implemented using the R-package TopGo and always using the dataset-specific gene background.

### EEG measurement and analysis

Surgery: EEG: Adult mutant or WT mice were used for surgery (8-10 weeks old at surgery). In general, methods used here mirrored those in Muheim et al. (71). In brief, 4 gold-plated miniature screws (0.9 mm diameter) were positioned as following: 2 screws on the frontal left and right hemispheres (±1.5 ML, 1.2 AP from bregma) for implant fixation purposes, 1 screw in the parietal left hemisphere (–1.7 ML, –1.7 AP from bregma) for EEG recordings, and a screw positioned above the cerebellum (0 ML, –2 AP from lambda) as a reference.

Contralateral to the parietal EEG screw (i.e., (1.7 ML, –1.7 AP from bregma), we implanted a cannula (Plastics One Inc., C315GA/Spc, cut 0.6 mm below pedestal) attached to an insulated wire (0.127 mm), protruding 0.2 mm below cannula tip, to obtain local field potential (LFP) from the injection site. Additionally, 2 gold wires (0.2 mm diameter) were bilaterally inserted in the neck muscle to obtain EMG signals. EEG, LFP, and EMG electrodes were then connected to stainless steel wires (0.07 mm thickness) wires that is further soldered to a custom-made fine cable. The implants were fixed to the skull with dental cement (Paladur 2-component system). Following surgery, mice were single housed in insulated recording chambers and allowed to recover and habituate to chambers for at least 7 days before recordings.

Experimental design: After recovery, we continuously collected EEG/EMG signals over the following 11 days. The first 24h, we obtained a baseline recording which was followed by microinfusion of 1µl of LNA against miR-218-5p (300µM) through cannula under isofluorane anesthesia. The first 24h of recording post-injection were excluded from analysis to avoid the confounding effects of anesthesia on sleep EEG patterns. Post recordings, mice were sacrificed, and the injected cortical area was extracted for RNA-processing.

Sleep data analysis: Both EEG and EMG signals were amplified (factor 2000), analogue filtered (high-pass filter: –3 dB at 0.016 Hz; low-pass filter: –3 dB at 40 Hz), sampled with 1024 Hz, then decimated and stored with a 128 Hz resolution in European Data Format (EDF) file. Signals were then digitally band-pass filtered using Chebyshev Type II filter (between 0.1 to 40 Hz for EEG and 10 to 30 Hz for EMG). EEG power spectra were computed for 4-s epochs by a Fast Fourier Transform routine within the frequency range of 0.5 – 25 Hz. Between 0.5 Hz and 5 Hz, 0.5 Hz bins were used, and between 5 and 25 Hz 1 Hz bins were used. Vigilance states (NREM, REM and wake) were automatically scored by SPINDLE (72) and scoring was visually validated by the experimenter. Epochs containing artifacts were identified and excluded from the spectral analysis. Spectral power was normalized to the average spectral power of the same vigilance state in the light phase (ZT0-12) of the baseline day. Slow wave activity (SWA) time course during NREM sleep, was calculated as the average delta power (0.75-4 Hz) within one-hour intervals relative to the average NREM delta power in the light phase (ZT0-12) of the baseline day. Data analysis was carried out using custom-made scripts in MATLAB (MathWorks R2020b).

### Statistics

All statistical modelling was performed using R (R 4.0.3, 10.10.2020) in custom R-markdown scripts. To correctly correct for batch effects, we implemented linear mixed models using the lmer-function from the R-packages lme4 and lmerTest. Post-hoc analysis was used using the emmeans R-package using Dunn’s test against a reference control. Linear mixed models were comprised of fixed effects such as pharmacological and LNA-treatments and their interaction as well as random effects. Random effects were independent biological replicates such as in vitro different primary hippocampal preparations or number of mice in vivo experiments. Data was always represented with a violin-plot showing the distribution density of the individual data-points as well as boxplots denoting median as a central line, 25th to 75th percentile as a box and whiskers as data points within 1.5× interquartile range (IQR). R-markdown of all employed statistical tests are available as in supplements.

## Supporting information

Statistics details

## Acknowledgments

We greatly acknowledge the technical support in the preparation of primary hippocampal cultures by Cristina Furler, Tatjana Wüst and Dr. Roberto Fiore. Furthermore, we would like to thank Dr. Roberto Fiore for support in acquisition of microscopy images as well as inputs on the manuscript. We also thank Dr. Brunno Rocha Levone for support in stereotactic injections and inputs on the manuscript, as well as Dr. Pierre-Luc Germain and Emanuel Sonder for inputs on statistical analyses and the manuscript. Furthermore, we would like to thank Prof. Shiva Tyagarajan for the generous gift of the gamma2-antibody. This work was in part funded by an ETH Research Grant (NeuroSno) to GS.

## References

1. G. Turrigiano, Homeostatic Synaptic Plasticity: Local and Global Mechanisms for Stabilizing Neuronal Function. Cold Spring Harb Perspect Biol 4, a005736 (2012).

2. J. Li, E. Park, L. R. Zhong, L. Chen, Homeostatic synaptic plasticity as a metaplasticity mechanism — a molecular and cellular perspective. Current Opinion in Neurobiology 54, 44–53 (2019).

3. D. Fernandes, A. L. Carvalho, Mechanisms of homeostatic plasticity in the excitatory synapse. Journal of Neurochemistry 139, 973–996 (2016).

4. M. D. Rannals, J. Kapur, Homeostatic Strengthening of Inhibitory Synapses Is Mediated by the Accumulation of GABAA Receptors. J Neurosci 31, 17701–17712 (2011).

5. H. Pribiag, H. Peng, W. A. Shah, D. Stellwagen, S. Carbonetto, Dystroglycan mediates homeostatic synaptic plasticity at GABAergic synapses. Proc Natl Acad Sci U S A 111, 6810– 6815 (2014).

6. L. C. Rutherford, S. B. Nelson, G. G. Turrigiano, BDNF Has Opposite Effects on the Quantal Amplitude of Pyramidal Neuron and Interneuron Excitatory Synapses. Neuron 21, 521–530 (1998).

7. D. Stellwagen, R. C. Malenka, Synaptic scaling mediated by glial TNF-α. Nature 440, 1054– 1059 (2006).

8. C. T. Schanzenbächer, S. Sambandan, J. D. Langer, E. M. Schuman, Nascent Proteome Remodeling following Homeostatic Scaling at Hippocampal Synapses. Neuron 92, 358–371 (2016).

9. A. R. Dörrbaum, B. Alvarez-Castelao, B. Nassim-Assir, J. D. Langer, E. M. Schuman, Proteome dynamics during homeostatic scaling in cultured neurons. eLife 9, e52939 (2020).

10. D. Colameo, et al., Pervasive compartment-specific regulation of gene expression during homeostatic synaptic scaling. EMBO reports 22, e52094 (2021).

11. M. Letellier, et al., miR-92a regulates expression of synaptic GluA1-containing AMPA receptors during homeostatic scaling. Nat Neurosci 17, 1040–1042 (2014).

12. S. Dubes, et al., miR-124-dependent tagging of synapses by synaptopodin enables input-specific homeostatic plasticity. EMBO J 41, e109012 (2022).

13. Q. Hou, et al., MicroRNA miR124 is required for the expression of homeostatic synaptic plasticity. Nat Commun 6, 10045 (2015).

14. J. E. Cohen, P. R. Lee, S. Chen, W. Li, R. D. Fields, MicroRNA regulation of homeostatic synaptic plasticity. Proceedings of the National Academy of Sciences 108, 11650–11655 (2011).

15. R. Fiore, et al., MiR-134-dependent regulation of Pumilio-2 is necessary for homeostatic synaptic depression. EMBO J 33, 2231–2246 (2014).

16. M. Rajman, et al., A microRNA-129-5p/Rbfox crosstalk coordinates homeostatic downscaling of excitatory synapses. The EMBO Journal 36, 1770–1787 (2017).

17. M. O. Inouye, D. Colameo, I. Ammann, J. Winterer, G. Schratt, miR-329– and miR-495– mediated Prr7 down-regulation is required for homeostatic synaptic depression in rat hippocampal neurons. Life Science Alliance 5 (2022).

18. N. S. Desai, R. H. Cudmore, S. B. Nelson, G. G. Turrigiano, Critical periods for experience-dependent synaptic scaling in visual cortex. Nat Neurosci 5, 783–789 (2002).

19. S. Glazewski, S. Greenhill, K. Fox, Time-course and mechanisms of homeostatic plasticity in layers 2/3 and 5 of the barrel cortex. Philos Trans R Soc Lond B Biol Sci 372, 20160150 (2017).

20. G. Tononi, C. Cirelli, Sleep and the Price of Plasticity: From Synaptic and Cellular Homeostasis to Memory Consolidation and Integration. Neuron 81, 12–34 (2014).

21. G. H. Diering, et al., Homer1a drives homeostatic scaling-down of excitatory synapses during sleep. Science 355, 511–515 (2017).

22. G. H. Diering, Remembering and forgetting in sleep: Selective synaptic plasticity during sleep driven by scaling factors Homer1a and Arc. Neurobiol Stress 22, 100512 (2022).

23. S. Maret, et al., Homer1a is a core brain molecular correlate of sleep loss. Proceedings of the National Academy of Sciences 104, 20090–20095 (2007).

24. I. Tobler, A. A. Borbély, The effect of 3-h and 6-h sleep deprivation on sleep and EEG spectra of the rat. Behavioural Brain Research 36, 73–78 (1990).

25. D.-J. Dijk, Regulation and Functional Correlates of Slow Wave Sleep. J Clin Sleep Med 5, S6– S15 (2009).

26. M. W. Chee, L. Y. Chuah, Functional neuroimaging insights into how sleep and sleep deprivation affect memory and cognition. Current Opinion in Neurology 21, 417 (2008).

27. Y. Chai, et al., Two nights of recovery sleep restores hippocampal connectivity but not episodic memory after total sleep deprivation. Sci Rep 10, 8774 (2020).

28. R. Huber, M. Felice Ghilardi, M. Massimini, G. Tononi, Local sleep and learning. Nature 430, 78–81 (2004).

29. M. C. D. Bridi, et al., Daily Oscillation of the Excitation-Inhibition Balance in Visual Cortical Circuits. Neuron 105, 621–629.e4 (2020).

30. S. B. Noya, et al., The forebrain synaptic transcriptome is organized by clocks but its proteome is driven by sleep. Science 366 (2019).

31. F. Brüning, et al., Sleep-wake cycles drive daily dynamics of synaptic phosphorylation. Science 366, eaav3617 (2019).

32. M. Soutschek, T. Germade, P.-L. Germain, G. Schratt, enrichMiR predicts functionally relevant microRNAs based on target collections. Nucleic Acids Research 50, W280–W289 (2022).

33. G. A. Wayman, et al., An activity-regulated microRNA controls dendritic plasticity by down-regulating p250GAP. Proceedings of the National Academy of Sciences 105, 9093–9098 (2008).

34. E. M. McNeill, et al., The conserved microRNA miR-34 regulates synaptogenesis via coordination of distinct mechanisms in presynaptic and postsynaptic cells. Nat Commun 11, 1092 (2020).

35. A. Rocchi, et al., Neurite-Enriched MicroRNA-218 Stimulates Translation of the GluA2 Subunit and Increases Excitatory Synaptic Strength. Mol Neurobiol 56, 5701–5714 (2019).

36. L. Mohammadipoor-Ghasemabad, M. H. Sangtarash, V. Sheibani, H. A. Sasan, S. Esmaeili-Mahani, Hippocampal microRNA-191a-5p Regulates BDNF Expression and Shows Correlation with Cognitive Impairment Induced by Paradoxical Sleep Deprivation. Neuroscience 414, 49–59 (2019).

37. D. Meyer, T. Bonhoeffer, V. Scheuss, Balance and Stability of Synaptic Structures during Synaptic Plasticity. Neuron 82, 430–443 (2014).

38. W. E. Heavner, et al., Remodeling of the Homer-Shank interactome mediates homeostatic plasticity. Science Signaling (2021) https:/doi.org/10.1126/scisignal.abd7325 (February 1, 2022).

39. C. A. Stein, et al., Efficient gene silencing by delivery of locked nucleic acid antisense oligonucleotides, unassisted by transfection reagents. Nucleic Acids Res 38, e3 (2010).

40. M. Soutschek, F. Gross, G. Schratt, P.-L. Germain, scanMiR: a biochemically based toolkit for versatile and efficient microRNA target prediction. Bioinformatics 38, 2466–2473 (2022).

41. S. A. Connor, et al., Loss of Synapse Repressor MDGA1 Enhances Perisomatic Inhibition, Confers Resistance to Network Excitation, and Impairs Cognitive Function. Cell Reports 21, 3637–3645 (2017).

42. A. Toledo, et al., MDGAs are fast-diffusing molecules that delay excitatory synapse development by altering neuroligin behavior. eLife 11, e75233 (2022).

43. K. R. Myers, K. Yu, J. Kremerskothen, E. Butt, J. Q. Zheng, The Nebulin Family LIM and SH3 Proteins Regulate Postsynaptic Development and Function. J. Neurosci. 40, 526–541 (2020).

44. K. W. Gripp, et al., Syndromic disorders caused by gain-of-function variants in KCNH1, KCNK4, and KCNN3—a subgroup of K+ channelopathies. Eur J Hum Genet 29, 1384–1395 (2021).

45. F. Wielaender, et al., Generalized myoclonic epilepsy with photosensitivity in juvenile dogs caused by a defective DIRAS family GTPase 1. Proc Natl Acad Sci U S A 114, 2669–2674 (2017).

46. S. tom Dieck, et al., Direct visualization of newly synthesized target proteins in situ. Nat Methods 12, 411–414 (2015).

47. H. Ali, L. Marth, D. Krueger-Burg, Neuroligin-2 as a central organizer of inhibitory synapses in health and disease. Science Signaling 13, eabd8379 (2020).

48. C. Schweizer, et al., The gamma 2 subunit of GABA(A) receptors is required for maintenance of receptors at mature synapses. Mol Cell Neurosci 24, 442–450 (2003).

49. L. de Vivo, et al., Ultrastructural evidence for synaptic scaling across the wake/sleep cycle. Science 355, 507–510 (2017).

50. G. H. Diering, R. L. Huganir, The AMPA Receptor Code of Synaptic Plasticity. Neuron 100, 314–329 (2018).

51. M. A. Bemben, et al., Contrasting synaptic roles of MDGA1 and MDGA2. 2023.05.25.542333 (2023).

52. J.-H. Hu, et al., Homeostatic scaling requires group I mGluR activation mediated by Homer1a. Neuron 68, 1128–1142 (2010).

53. G. Schratt, Fine-tuning neural gene expression with microRNAs. Current Opinion in Neurobiology 19, 213–219 (2009).

54. M. S. Ebert, P. A. Sharp, Roles for microRNAs in conferring robustness to biological processes. Cell 149, 515–524 (2012).

55. J. Durkin, et al., Cortically coordinated NREM thalamocortical oscillations play an essential, instructive role in visual system plasticity. Proceedings of the National Academy of Sciences 114, 10485–10490 (2017).

56. G. Tononi, C. Cirelli, “Sleep and Synaptic Down-Selection” in Micro-, Meso– and Macro-Dynamics of the Brain, G. Buzsáki, Y. Christen, Eds. (Springer, 2016) (December 17, 2018).

57. L. B. Krone, et al., A role for the cortex in sleep–wake regulation. Nat Neurosci 24, 1210– 1215 (2021).

58. B. Collins, et al., Circadian VIPergic Neurons of the Suprachiasmatic Nuclei Sculpt the Sleep-Wake Cycle. Neuron 108, 486–499.e5 (2020).

59. A. Torres-Berrío, et al., MiR-218: A Molecular Switch and Potential Biomarker of Susceptibility to Stress. Mol Psychiatry 25, 951–964 (2020).

60. E. T. Kavalali, L. M. Monteggia, Targeting Homeostatic Synaptic Plasticity for Treatment of Mood Disorders. Neuron 106, 715–726 (2020).

61. B. C. Monteiro, et al., Relationship Between Brain-Derived Neurotrofic Factor (Bdnf) and Sleep on Depression: A Critical Review. Clin Pract Epidemiol Ment Health 13, 213–219 (2017).

62. A. Antoniou, L. Auderset, L. Kaurani, A. Fischer, A. Schneider, Neuronal extracellular vesicles mediate BDNF-dependent dendritogenesis and synapse maturation via microRNAs. 2021.05.11.443606 (2021).

63. Seth R. Taylor, et al., MicroRNA-218 instructs proper assembly of hippocampal networks. bioRxiv, 2022.08.24.505085 (2022).

64. S. S. Kaalund, et al., Aberrant expression of miR-218 and miR-204 in human mesial temporal lobe epilepsy and hippocampal sclerosis-convergence on axonal guidance. Epilepsia 55, 2017–2027 (2014).

65. G. Lignani, P. Baldelli, V. Marra, Homeostatic Plasticity in Epilepsy. Frontiers in Cellular Neuroscience 14 (2020).

