## Supplemental Figures for "microRNA-218-5p Coordinates Scaling of Excitatory and Inhibitory Synapses during Homeostatic Synaptic Plasticity"

*Gerhard Schratt

PNAS strongly encourages authors to supply an ORCID identifier for each author. Do not include ORCIDs in the manuscript file; individual authors must link their ORCID account to their PNAS account at www.pnascentral.org. For proper authentication, authors must provide their ORCID at submission and are not permitted to add ORCIDs on proofs.

**Author Contributions:** DC designed project, prepared RNA-samples, performed RNA-seq analyses, microscopy experiments, RT-qPCR-validations, prepared the figures and wrote the original draft of the manuscript. SMM cloned luciferase constructs and performed EEG-cannula implantations, sleep recordings, sleep deprivation experiments and initial data analysis under the supervision of WEG and SAB. WEG analyzed and conceptualized sleep experiments together with SAB. GC performed stereotactic injections and helped with sample preparations. SG performed luciferase assays. GS supervised the project, edited and reviewed the manuscript and supplied funding.

**Competing Interest Statement:** The authors declare no competing interests.

**Classification:** BIOLOGICAL SCIENCES - Neuroscience

**Keywords:** homeostatic synaptic plasticity, excitatory and inhibitory synaptic scaling, microRNA, non-REM sleep, sleep-dependent synaptic plasticity

**This PDF file includes:**

Supplemental Figures 1 – 7

Supplemental Table 1

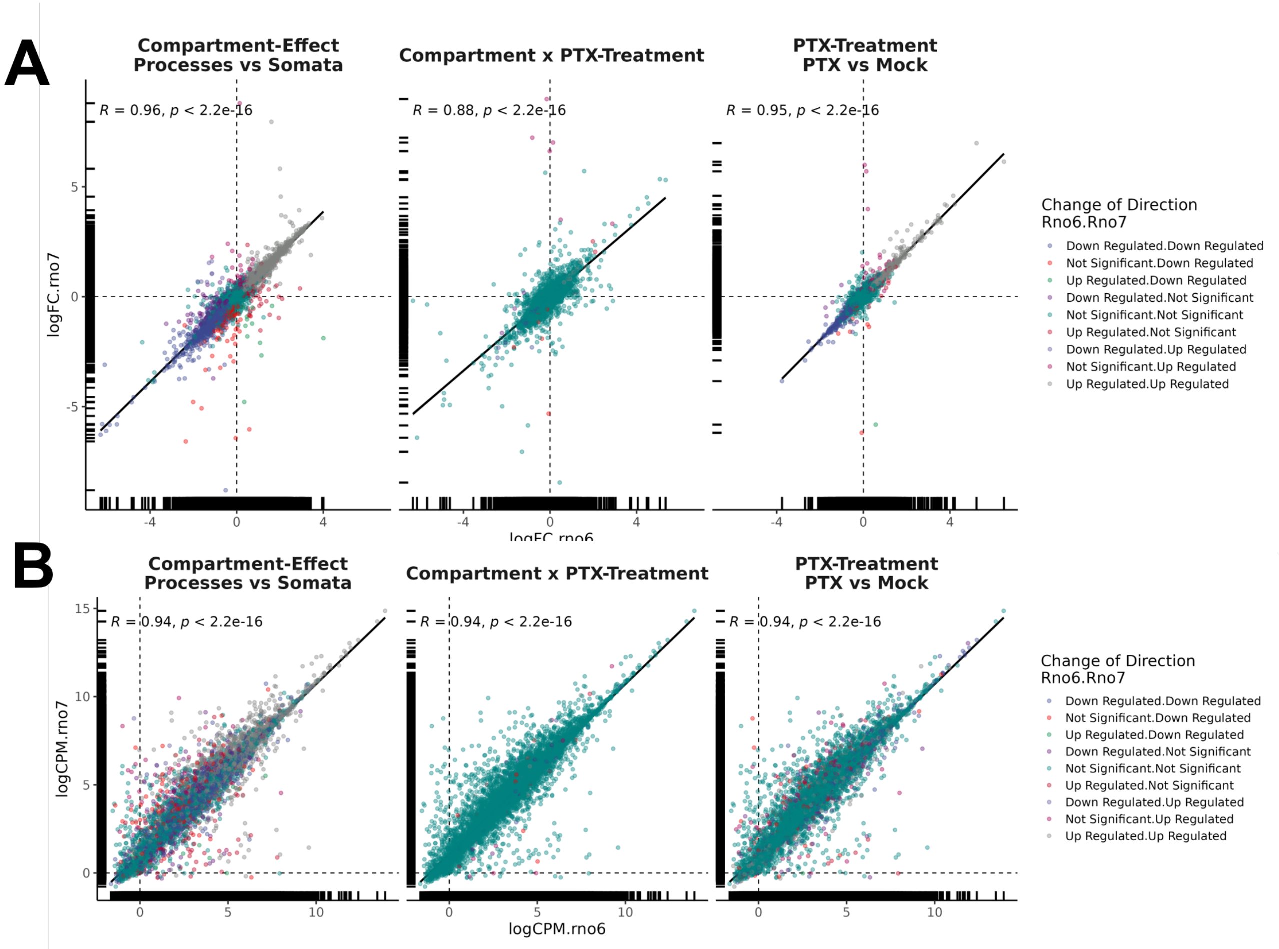


**Supplemental Figure 1: Comparison between alignment of PTX-dataset to Rno7 or Rno6**

The same models for differential expression analysis was used for both models in the form of ~PTX-Treatment x Compartment + SV1 (surrogate variable analysis)

(A) Correlation plots of log-fold-changes (logFC) changes between Rno6 and Rno7 in the different contrasts. R- and p-value was calculated by Spearman's correlation test. Colors were generated by combination of change of direction in Rno6 and Rno7.

(B) Correlation plots of log counts per millions (logCPM) changes between Rno6 and Rno7 in the different contrasts. R- and p-value were calculated by Spearman's correlation test. Colors were generated by combination of change of direction in Rno6 and Rno7.

| Effect-Term | Significant in Rno6 | Significant in Rno7 | total Genes |
|---|---|---|---|
| DEA.CompartmentProcesses | Down Regulated | Down Regulated | 3234 |
| DEA.CompartmentProcesses | Down Regulated | Not Significant | 283 |
| DEA.CompartmentProcesses | Down Regulated | Up Regulated | 9 |
| DEA.CompartmentProcesses | Not Significant | Down Regulated | 318 |
| DEA.CompartmentProcesses | Not Significant | Not Significant | 4638 |
| DEA.CompartmentProcesses | Not Significant | Up Regulated | 201 |
| DEA.CompartmentProcesses | Up Regulated | Down Regulated | 13 |
| DEA.CompartmentProcesses | Up Regulated | Not Significant | 195 |
| DEA.CompartmentProcesses | Up Regulated | Up Regulated | 3381 |
| DEA.CompartmentProcesses:PTXTreatmentPTX | Down Regulated | Down Regulated | 42 |
| DEA.CompartmentProcesses:PTXTreatmentPTX | Down Regulated | Not Significant | 9 |
| DEA.CompartmentProcesses:PTXTreatmentPTX | Not Significant | Down Regulated | 28 |
| DEA.CompartmentProcesses:PTXTreatmentPTX | Not Significant | Not Significant | 12137 |
| DEA.CompartmentProcesses:PTXTreatmentPTX | Not Significant | Up Regulated | 16 |
| DEA.CompartmentProcesses:PTXTreatmentPTX | Up Regulated | Not Significant | 8 |
| DEA.CompartmentProcesses:PTXTreatmentPTX | Up Regulated | Up Regulated | 32 |
| DEA.PTXTreatmentPTX | Down Regulated | Down Regulated | 946 |
| DEA.PTXTreatmentPTX | Down Regulated | Not Significant | 94 |
| DEA.PTXTreatmentPTX | Not Significant | Down Regulated | 113 |
| DEA.PTXTreatmentPTX | Not Significant | Not Significant | 9757 |
| DEA.PTXTreatmentPTX | Not Significant | Up Regulated | 266 |
| DEA.PTXTreatmentPTX | Up Regulated | Down Regulated | 1 |
| DEA.PTXTreatmentPTX | Up Regulated | Not Significant | 60 |
| DEA.PTXTreatmentPTX | Up Regulated | Up Regulated | 1035 |

**Supplementary Table 1: Table showing difference in DEGs between Rno6 and Rno7 alignments.**

Same model was used for both alignments.

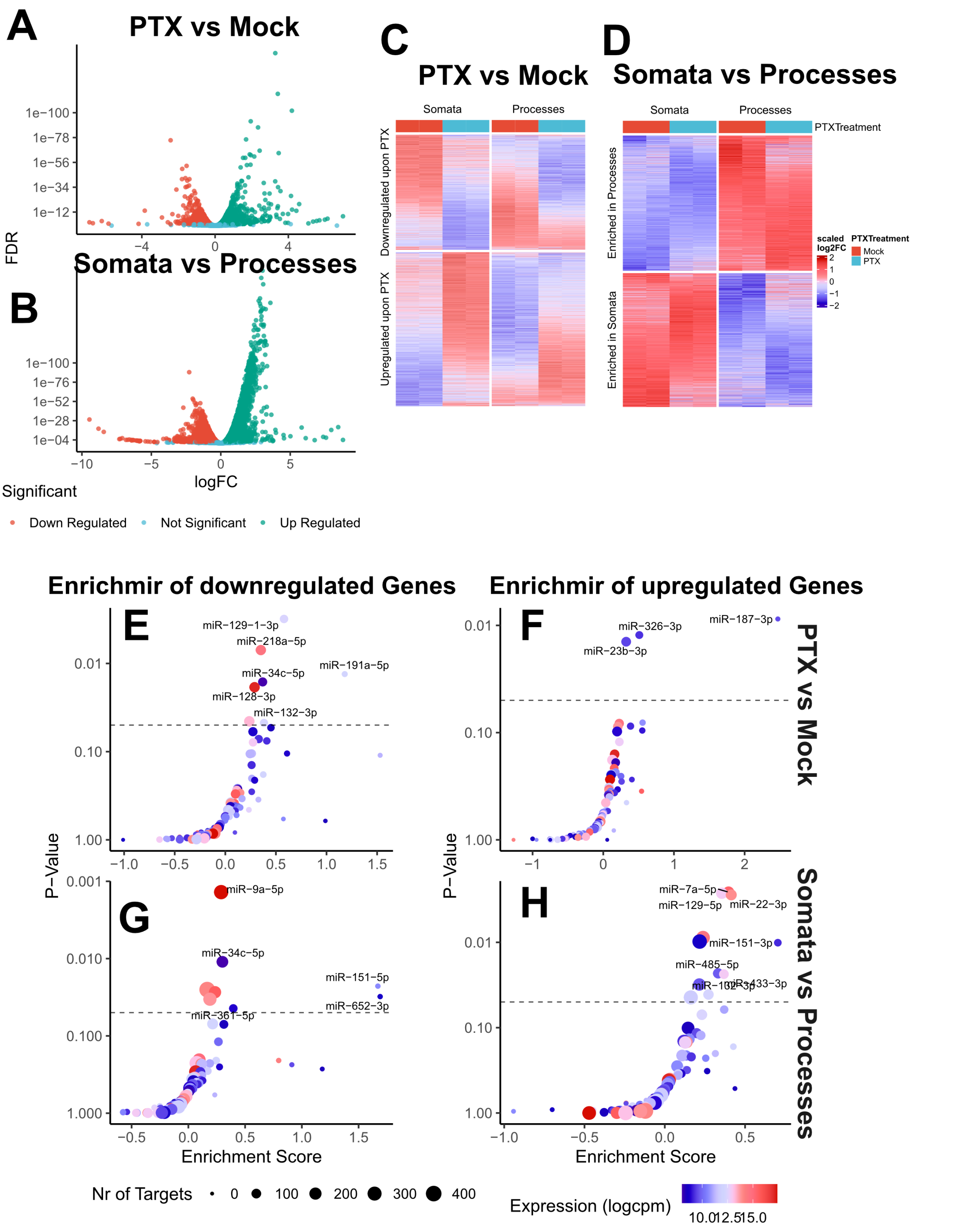


**Supplementary *Figure* 2: Differential Expression Analysis of main-effects of PTX-Treatment and Compartment and enrichMiR-analysis**

(A) and (B) Volcano plots of differentially expressed genes in the main effects of PTX-Treatment and Compartment (logFC(PTX/Mock) and log(Processes/Somata)

(C) and (D) Heatmaps of all significantly changing genes in the main effects of PTX-Treatment and Compartment. Scaled logFC relative to mock-treated Somata logCPM

(E) – (H) enrichMiR-plots of miRNA-binding sites enriched in up- or downregulated genes in the main effects of PTX-Treatment and Compartment

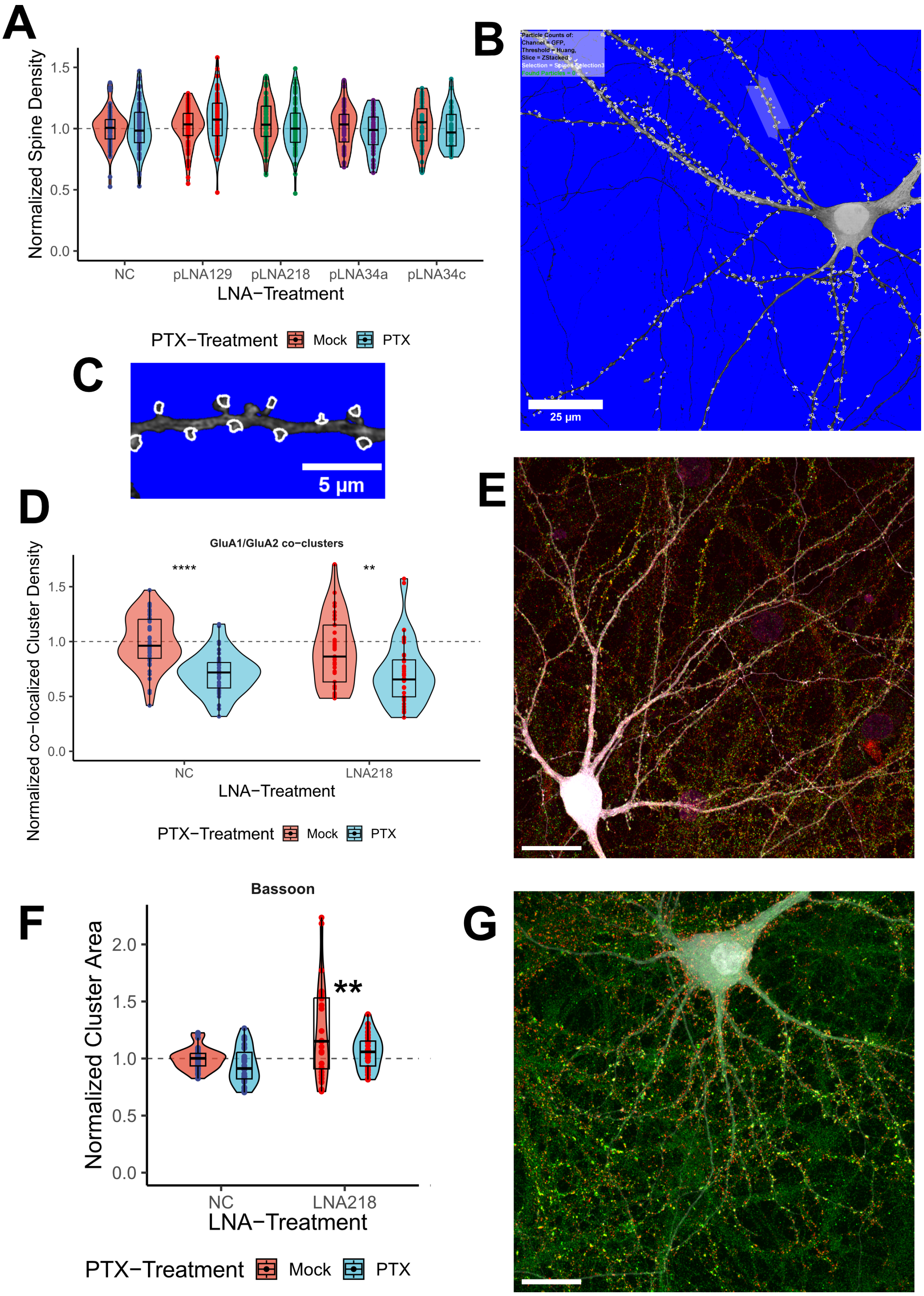


**Supplementary Figure 3: Spine and Cluster Densities**

(A) Quantification of spine densities (number of spines / area of cell) normalized to the mean of the mock-treated, negative control (NC) group of the corresponding independent biological replicate. 8-14 cells per condition and per independent biological replicate (n=8). Data was fit to a linear-mixed model with fixed effects for PTX-treatment and LNA-treatment and their interaction and random effect of independent biological replicates. Dunn's post-hoc analysis was used to estimate PTX-effect in different LNA-condition and to estimate LNA-effect compared to negative control within PTX-treatment *p<0.05, **p<0.01, ***p<0.001, ****p<0.0001. Continued on next page

(B) and (C) Example output of the automated spine analysis

(D) Quantification of GluA2/GluA1 co-cluster densities (number of clusters / area of cell) normalized to the mean of the mock-treated, negative control (NC) group of the corresponding independent biological replicate of primary hippocampal preparations. 8-12 images per condition and per independent biological replicate n=3. Data was fit to a linear-mixed model with fixed effects for PTX-treatment and LNA-treatment and their interaction and random effect of independent biological replicates. Dunn's post-hoc analysis was used to estimate PTX-effect in NC and LNA218-conditions and to estimate LNA218-effect compared to negative control within PTX-treatment *p<0.05, **p<0.01, ***p<0.001, ****p<0.0001.

(E) Example picture of a whole primary hippocampal neuron immunostained for GluA2 and GluA1 and transfected with GFP. White bar represents 25µm

(F) Quantification of cluster area of Basson normalized to the mean of the mock-treated, negative control (NC) group of the corresponding independent biological replicate of primary hippocampal preparations. 8-12 images per condition and per independent biological replicate n=3. Data was fit to a linear-mixed model with fixed effects for PTX-treatment and LNA-treatment and their interaction and random effect of independent biological replicates. Dunn's post-hoc analysis was used to estimate PTX-effect in NC and LNA218-conditions and to estimate LNA218-effect compared to negative control within PTX-treatment *p<0.05, **p<0.01, ***p<0.001, ****p<0.0001.

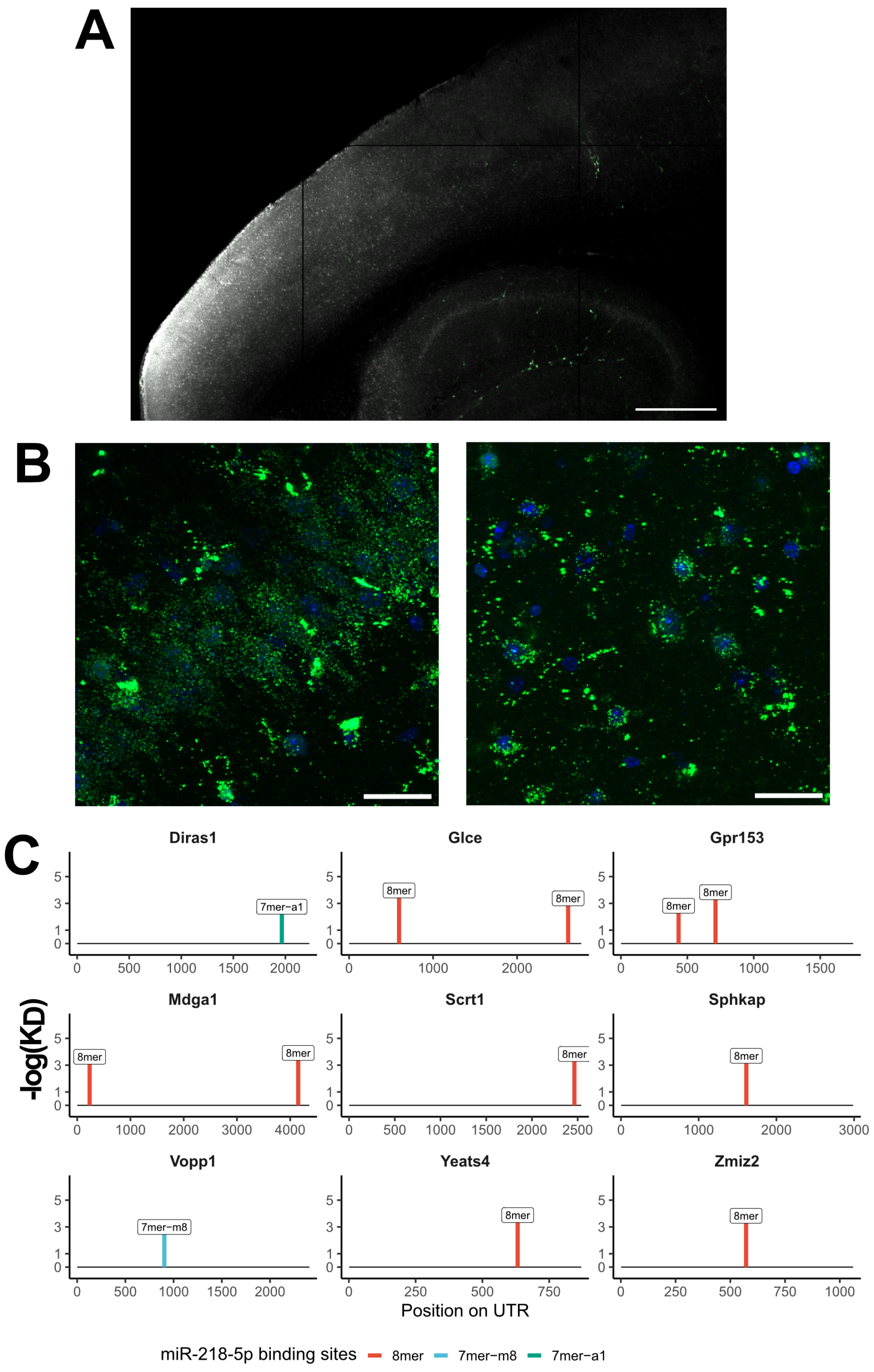


**Supplementary Figure 4: Examples of non-Injection sites and 3'UTRs of upregulated genes upon LNA-treatment**

(A) Ventral slice of an injected brain (same as in Figure 3A) with lack of fluorescence except in blood vessels (grey=DAPI, green = FAM-labelled LNA). White bar represents 500µm

(B) Higher magnification of CA1 pyramidal cell layer (left) and SI-cortical area (right) (Blue=DAPI, green = FAM-labelled LNA). White bar represents 250µm.

(C) 3’UTRs of all significantly upregulated genes containing miR-218-5p binding sites predicted by *scanMiR.*

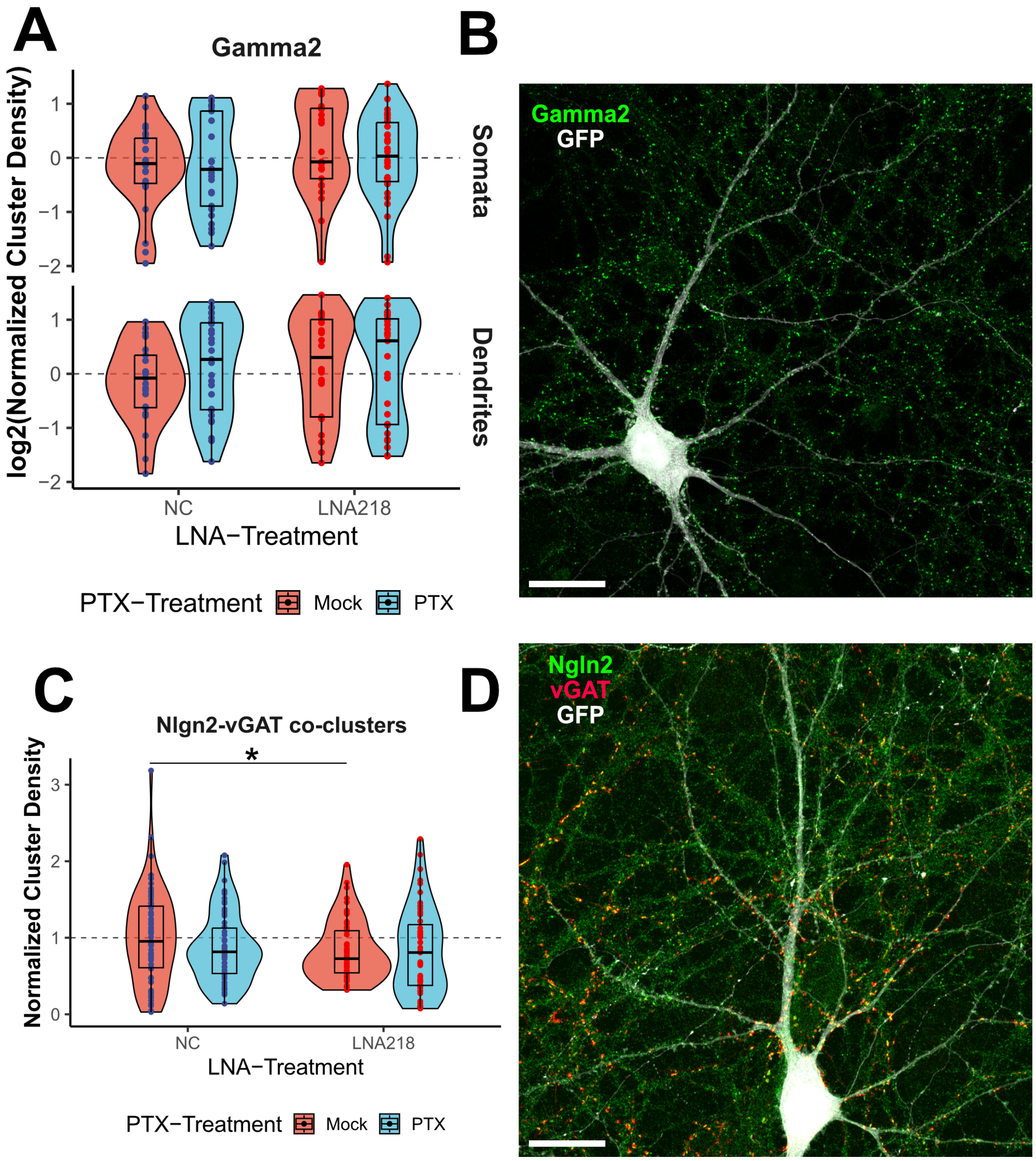


**Supplementary Figure 5: Cluster Densities of Gamma2 clusters and Nlgn2-vGAT co-clusters**

(A) Quantification of Gamma2-cluster densities (number of clusters / area of cell) normalized to the mean of the mock-treated, negative control (NC) group of the corresponding independent biological replicate of primary hippocampal preparations. 8-12 images per condition and per independent biological replicate n=3. Data was fit to a linear-mixed model with fixed effects for PTX-treatment and LNA-treatment and their interaction and random effect of independent biological replicates. Dunn's post-hoc analysis was used to estimate PTX-effect in NC and LNA218-conditions and to estimate LNA218-effect compared to negative control within PTX-treatment

(B) Example picture of a whole primary hippocampal neuron immunostained for Gamma2 and transfected with GFP. White bar represents 25µm

(A) Quantification of Nlgn2-vGAT co-cluster densities (number of clusters / area of cell) normalized to the mean of the mock-treated, negative control (NC) group of the corresponding independent biological replicate of primary hippocampal preparations. 8-12 images per condition and per independent biological replicate n=3. Data was fit to a linear-mixed model with fixed effects for PTX-treatment and LNA-treatment and their interaction and random effect of independent biological replicates. Dunn's post-hoc analysis was used to estimate PTX-effect in NC and LNA218-conditions and to estimate LNA218-effect compared to negative control within PTX-treatment *p<0.05

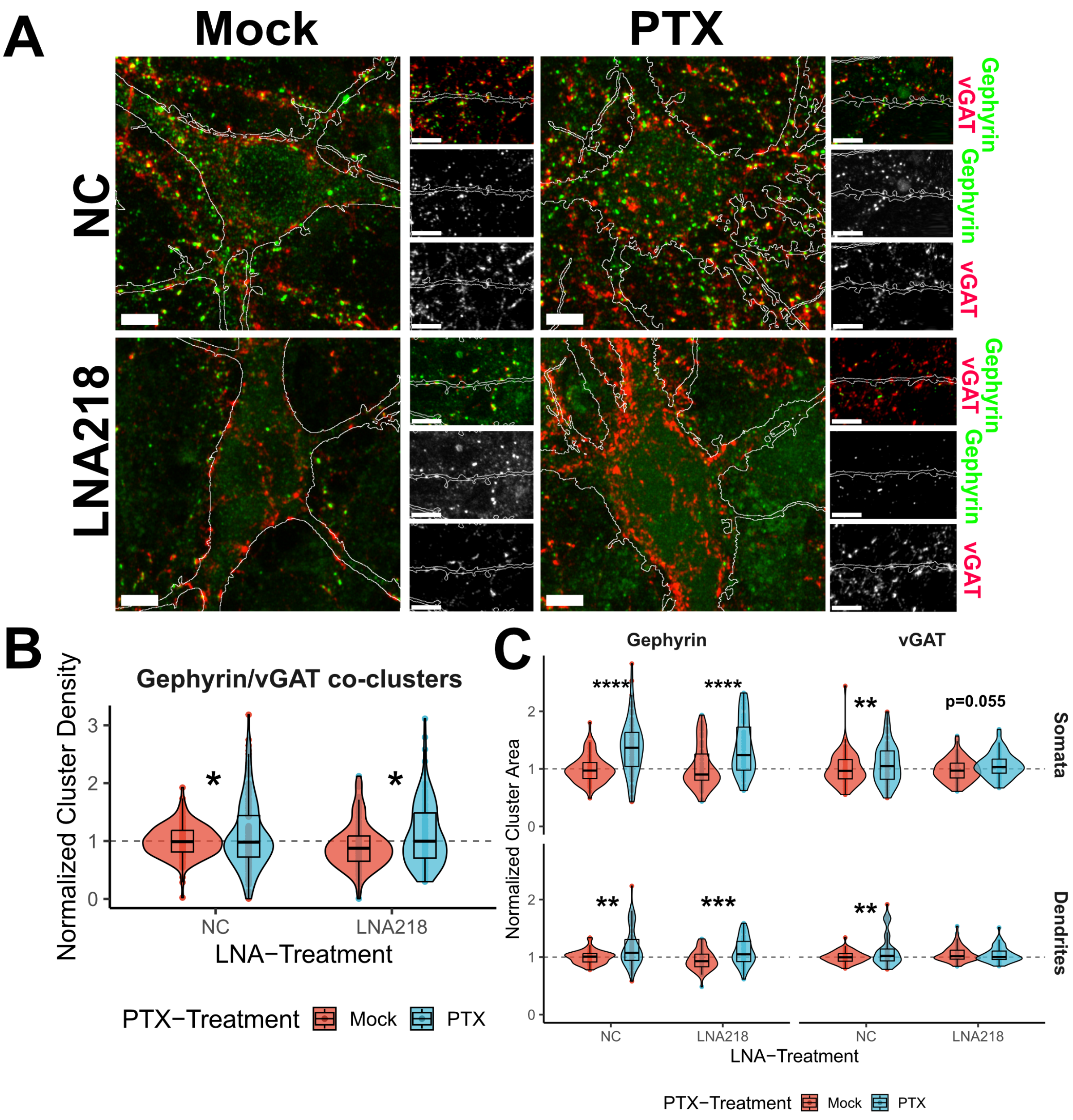


**Supplementary *Figure 6:* Gephyrin and vGAT cluster changes upon PTX-treatment and miR-218-5p inhibition**

(A) Example pictures of soma and dendrites of primary hippocampal neurons transfected with GFP (white outline) and either NC or LNA218 in combination with PTX- and mock-treatment and stained for postsynaptic Gephyrin and presynaptic vGAT. White tool bar in cellbody inset represents 10µm and in the dendrite inset represents 10µm.

(B) and (C) Quantification of cluster area (B) and co-cluster density (number of clusters / area of the cell) (C) of Gephyrin and vGAT normalized to the mean of the mock-treated, negative control (NC) group of the corresponding independent biological replicate of primary hippocampal preparations. 8-12 images per condition and per independent biological replicate n=3. Data was fit to a linear-mixed model per compartment (soma or dendrites) and per staining (Nlgn2 and vGAT) with fixed effects for PTX-treatment and LNA-treatment and their interaction and random effect of independent biological replicates. Dunn's post-hoc analysis was used to estimate PTX-effect in NC and LNA218-conditions and to estimate LNA218-effect compared to negative control within PTX-treatment *p<0.05, **p<0.01, ***p<0.001, ****p<0.0001.

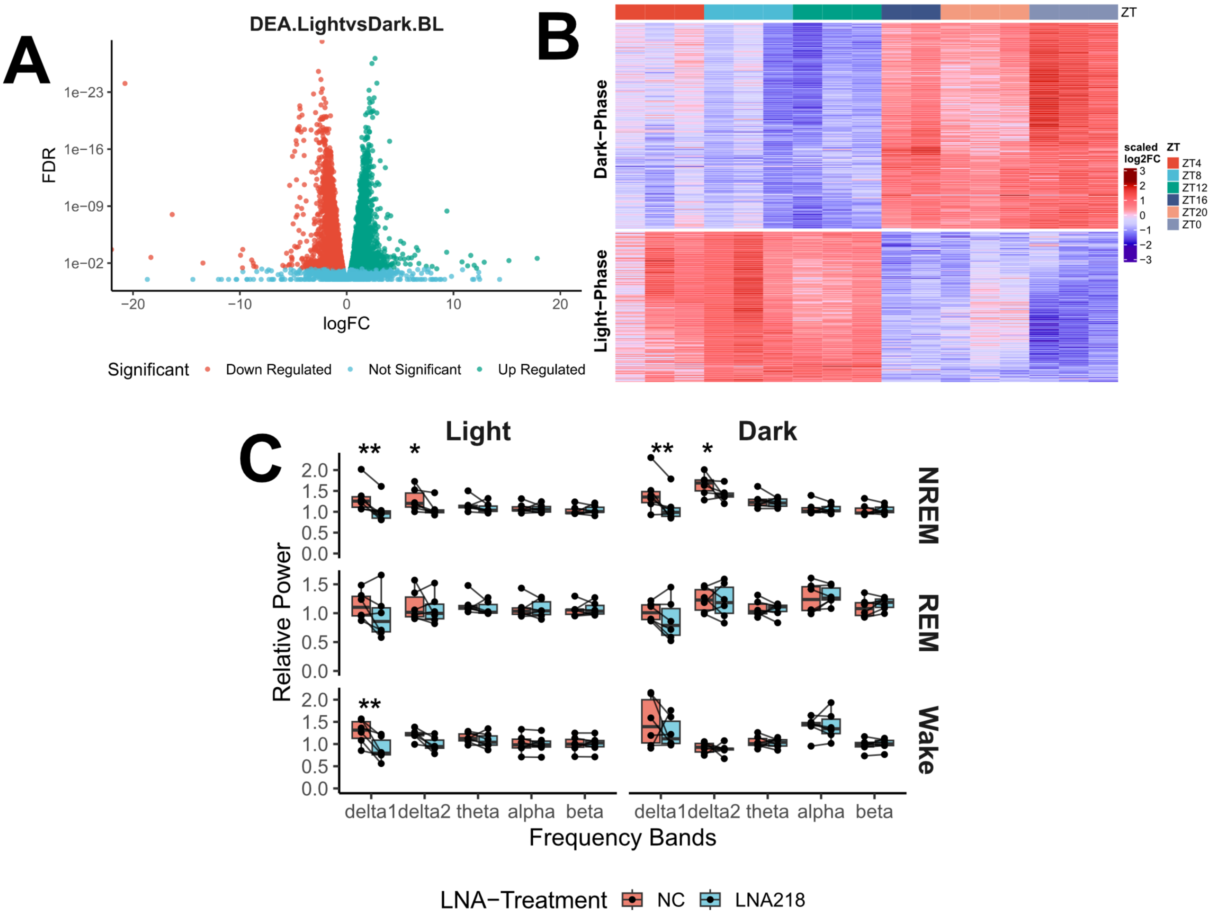


**Supplementary Figure 5: Genes enriched in the Light- or Dark-Phase using the Dataset from Noya et al., 2019 and Relative Power during Vigilance state**

(A) Volcano plot of DEGs enriched in either the light-phase (Up Regulated, ZT4, ZT8, ZT12) or the dark-phase (Down Regulated, ZT16, ZT16, ZT20), FDR-cutoff < 0.05, DEA = Differential Expression Analysis

(B) Heatmap showing top 500 differentially expressed genes. Scaled logFC relative to ZT0.

(C) Frequency-binned relative power in three vigilance stages (NREM, REM and wake) at postinjection-day 9, relative to the average power of corresponding vigilance state in baseline day (ZT0-12). Data was fit to a linear-mixed model per vigilance state with fixed effects for LNA-treatment and Frequency-bands and random effect of independent biological replicates (n=6 mice). Dunn's post-hoc analysis was used to estimate LNA-effect on hemispheres injected with LNA218 compared to corresponding control hemisphere within mouse and vigilance. *p<0.05, **p<0.01
