## Supplementary material for "microRNA-218-5p Coordinates Scaling of Excitatory and Inhibitory Synapses during Homeostatic Synaptic Plasticity": Statistics details

David Colameo

9 10 2023

#### Statistics in all presented Plots

##### Load Libraries

```
pkgTest <- function(x)
  {
    if (!require(x,character.only = TRUE))
    {
      install.packages(x,dep=TRUE)
        if(!require(x,character.only = TRUE)) stop("Package not found")
    }
  }

packages <- c("ggplot2", "ggsci", "ggpubr",
              "SummarizedExperiment", "plgINS", "SEtools", "scales",
              "ggrepel", "stringr", "sechm", "lme4", "lmerTest", "emmeans",
              "topGO","org.Rn.eg.db", "rstatix", "xlsx", "readxl")



for(package in packages){
  pkgTest(package)
}

ag <- function(df, cols, fun=mean) {
  c <- colnames(df[,unlist(lapply(df, is.numeric))])
  print(c)
  df <- aggregate(df[,c], by=as.list(df[,cols]), FUN=fun)
  return(df[, colSums(is.na(df)) != nrow(df)])
}

reverselog_trans <- function(base = exp(1)) {
  trans <- function(x) -log(x, base)
  inv <- function(x) base^(-x)
  trans_new(paste0("reverselog-", format(base)), trans, inv,
            log_breaks(base = base),
            domain = c(1e-1000, Inf))
}


allData <- readRDS("~/projects/Manuscript_miR218/R-Markdowns/allData.rds")
```

### Figure 1:

##### Figure 1E: Statistics on qPCR

- Measure miRNA-levels of miR-129-5p, miR-218-5p and miR-34c-5p in RNA from compartmentalized and PTX-treated primary hippocampal cultures
- Usage of a linear-mixed model with fixed effects of PTX-treatment and Compartment localisation and random effect of independent-biological replicate (n=8, number of hippocampal preparations)
- -dCq normalized to U6, in the log-scale

Three linear mixed model per miRNA followed by Dunn's post-hoc test and multiple testing correction

miR-129-5p Model

```
qpcr.data <- allData$qpcr.data
mod <- lmer(-dCq ~ Compartment*PTXTreatment + (1|Experiment),
            subset(qpcr.data, Gene %in% c("miR129")))
summary(mod)
```

```
## Linear mixed model fit by REML. t-tests use Satterthwaite's method [
## lmerModLmerTest]
## Formula: -dCq ~ Compartment * PTXTreatment + (1 | Experiment)
##    Data: subset(qpcr.data, Gene %in% c("miR129"))
##
## REML criterion at convergence: 67.3
##
## Scaled residuals:
##      Min       1Q   Median       3Q      Max
## -1.61148 -0.55051  0.01075  0.30756  2.91911
##
## Random effects:
##  Groups     Name        Variance Std.Dev.
##  Experiment (Intercept) 0.2582   0.5081
##  Residual               0.5329   0.7300
## Number of obs: 28, groups:  Experiment, 7
##
## Fixed effects:
##                                      Estimate Std. Error      df t value
## (Intercept)                           -5.7085     0.3362 18.1887 -16.981
## CompartmentProcesses                   1.9668     0.3902 18.0000   5.040
## PTXTreatmentPTX                        1.3936     0.3902 18.0000   3.572
## CompartmentProcesses:PTXTreatmentPTX  -0.2123     0.5518 18.0000  -0.385
##                                      Pr(>|t|)
## (Intercept)                          1.32e-12 ***
## CompartmentProcesses                 8.51e-05 ***
## PTXTreatmentPTX                       0.00218 **
## CompartmentProcesses:PTXTreatmentPTX  0.70497
## ---
## Signif. codes:  0 '***' 0.001 '**' 0.01 '*' 0.05 '.' 0.1 ' ' 1
##
## Correlation of Fixed Effects:
##             (Intr) CmprtP PTXTPT
## CmprtmntPrc -0.580
## PTXTrtmnPTX -0.580  0.500
```

```
## CmP:PTXTPTX  0.410 -0.707 -0.707
```

```
emmeans(mod, trt.vs.ctrl ~ PTXTreatment | Compartment, ref = "Mock")
```

```
## $emmeans
## Compartment = Somata:
##  PTXTreatment emmean    SE   df lower.CL upper.CL
##  Mock          -5.71 0.336 18.2    -6.41    -5.00
##  PTX           -4.31 0.336 18.2    -5.02    -3.61
##
## Compartment = Processes:
##  PTXTreatment emmean    SE   df lower.CL upper.CL
##  Mock          -3.74 0.336 18.2    -4.45    -3.04
##  PTX           -2.56 0.336 18.2    -3.27    -1.85
##
## Degrees-of-freedom method: kenward-roger
## Confidence level used: 0.95
##
## $contrasts
## Compartment = Somata:
##  contrast   estimate   SE df t.ratio p.value
##  PTX - Mock     1.39 0.39 18   3.572  0.0022
##
## Compartment = Processes:
##  contrast   estimate   SE df t.ratio p.value
##  PTX - Mock     1.18 0.39 18   3.027  0.0072
##
## Degrees-of-freedom method: kenward-roger
```

```
contr1 <- as.data.frame(emmeans(mod, trt.vs.ctrl ~ PTXTreatment | Compartment, ref = "Mock")$contrasts)
contr1$Gene <- "miR129"
```

miR-218-5p Model

```
mod <- lmer(-dCq ~ Compartment*PTXTreatment + (1|Experiment),
            subset(qpcr.data, Gene %in% c("miR218")))
summary(mod)
```

```
## Linear mixed model fit by REML. t-tests use Satterthwaite's method [
## lmerModLmerTest]
## Formula: -dCq ~ Compartment * PTXTreatment + (1 | Experiment)
##    Data: subset(qpcr.data, Gene %in% c("miR218"))
##
## REML criterion at convergence: 45.9
##
## Scaled residuals:
##      Min       1Q   Median       3Q      Max
## -1.60463 -0.58056  0.08945  0.52680  1.86623
##
## Random effects:
##  Groups     Name        Variance Std.Dev.
##  Experiment (Intercept) 0.1418   0.3765
##  Residual               0.2062   0.4541
## Number of obs: 28, groups:  Experiment, 7
##
## Fixed effects:
```

```
##                                     Estimate Std. Error      df t value
## (Intercept)                          -1.2969     0.2229 16.0215  -5.817
## CompartmentProcesses                  1.6371     0.2427 18.0000   6.745
## PTXTreatmentPTX                       0.2729     0.2427 18.0000   1.124
## CompartmentProcesses:PTXTreatmentPTX  0.4649     0.3432 18.0000   1.354
##                                     Pr(>|t|)
## (Intercept)                         2.61e-05 ***
## CompartmentProcesses                2.54e-06 ***
## PTXTreatmentPTX                        0.276
## CompartmentProcesses:PTXTreatmentPTX   0.192
## ---
## Signif. codes:  0 '***' 0.001 '**' 0.01 '*' 0.05 '.' 0.1 ' ' 1
##
## Correlation of Fixed Effects:
##              (Intr) CmprtP PTXTPT
## CmprtmntPrc -0.544
## PTXTrtmnPTX -0.544  0.500
## CmP:PTXTPTX  0.385 -0.707 -0.707
```

```
emmeans(mod, trt.vs.ctrl ~ PTXTreatment | Compartment, ref = "Mock")
```

```
## $emmeans
## Compartment = Somata:
##  PTXTreatment emmean    SE df lower.CL upper.CL
##  Mock          -1.30 0.223 16   -1.769   -0.824
##  PTX           -1.02 0.223 16   -1.497   -0.551
##
## Compartment = Processes:
##  PTXTreatment emmean    SE df lower.CL upper.CL
##  Mock           0.34 0.223 16   -0.132    0.813
##  PTX            1.08 0.223 16    0.605    1.551
##
## Degrees-of-freedom method: kenward-roger
## Confidence level used: 0.95
##
## $contrasts
## Compartment = Somata:
##  contrast   estimate    SE df t.ratio p.value
##  PTX - Mock    0.273 0.243 18   1.124  0.2757
##
## Compartment = Processes:
##  contrast   estimate    SE df t.ratio p.value
##  PTX - Mock    0.738 0.243 18   3.040  0.0071
##
## Degrees-of-freedom method: kenward-roger
```

```
contr2 <- as.data.frame(emmeans(mod, trt.vs.ctrl ~ PTXTreatment | Compartment, ref = "Mock")$contrasts)
contr2$Gene <- "miR218"
```

mir-34c-5p Model

```
mod <- lmer(-dCq ~ Compartment*PTXTreatment + (1|Experiment),
            subset(qpcr.data, Gene %in% c("miR34c")))
summary(mod)
```

```
## Linear mixed model fit by REML. t-tests use Satterthwaite's method [
```

```
## lmerModLmerTest]
## Formula: -dCq ~ Compartment * PTXTreatment + (1 | Experiment)
##    Data: subset(qpcr.data, Gene %in% c("miR34c"))
##
## REML criterion at convergence: 93.4
##
## Scaled residuals:
##      Min       1Q   Median       3Q      Max
## -2.25652 -0.51224 -0.09161  0.57494  1.84926
##
## Random effects:
##  Groups     Name        Variance Std.Dev.
##  Experiment (Intercept) 1.854    1.362
##  Residual               1.282    1.132
## Number of obs: 28, groups:  Experiment, 7
##
## Fixed effects:
##                                      Estimate Std. Error      df t value
## (Intercept)                           -4.8514     0.6694 11.7160  -7.247
## CompartmentProcesses                  -0.1508     0.6053 18.0000  -0.249
## PTXTreatmentPTX                       -0.2479     0.6053 18.0000  -0.410
## CompartmentProcesses:PTXTreatmentPTX   0.9155     0.8560 18.0000   1.069
##                                      Pr(>|t|)
## (Intercept)                          1.16e-05 ***
## CompartmentProcesses                    0.806
## PTXTreatmentPTX                         0.687
## CompartmentProcesses:PTXTreatmentPTX    0.299
## ---
## Signif. codes:  0 '***' 0.001 '**' 0.01 '*' 0.05 '.' 0.1 ' ' 1
##
## Correlation of Fixed Effects:
##             (Intr) CmprtP PTXTPT
## CmprtmntPrc -0.452
## PTXTrtmnPTX -0.452  0.500
## CmP:PTXTPTX  0.320 -0.707 -0.707
```

```
emmeans(mod, trt.vs.ctrl ~ PTXTreatment | Compartment, ref = "Mock")
```

```
## $emmeans
## Compartment = Somata:
##  PTXTreatment emmean    SE   df lower.CL upper.CL
##  Mock          -4.85 0.669 11.7    -6.31    -3.39
##  PTX           -5.10 0.669 11.7    -6.56    -3.64
##
## Compartment = Processes:
##  PTXTreatment emmean    SE   df lower.CL upper.CL
##  Mock          -5.00 0.669 11.7    -6.46    -3.54
##  PTX           -4.33 0.669 11.7    -5.80    -2.87
##
## Degrees-of-freedom method: kenward-roger
## Confidence level used: 0.95
##
## $contrasts
## Compartment = Somata:
##  contrast   estimate    SE df t.ratio p.value
```

```
##  PTX - Mock   -0.248 0.605 18  -0.410  0.6869
##
## Compartment = Processes:
##  contrast   estimate    SE df t.ratio p.value
##  PTX - Mock    0.668 0.605 18   1.103  0.2846
##
## Degrees-of-freedom method: kenward-roger
```

```
contr3 <- as.data.frame(emmeans(mod, trt.vs.ctrl ~ PTXTreatment | Compartment, ref = "Mock")$contrasts)
contr3$Gene <- "miR34c"
```

- Multiple testing correction of individual miRNA-contrasts and Fold-change estimation

```
##     contrast Compartment   estimate        SE df    t.ratio     p.value   Gene
## 1 PTX - Mock      Somata  1.3936302 0.3902047 18  3.5715362 0.002180983 miR129
## 2 PTX - Mock   Processes  1.1813405 0.3902047 18  3.0274892 0.007239085 miR129
## 3 PTX - Mock      Somata  0.2728587 0.2427112 18  1.1242113 0.275692842 miR218
## 4 PTX - Mock   Processes  0.7377404 0.2427112 18  3.0395809 0.007050832 miR218
## 5 PTX - Mock      Somata -0.2479441 0.6053136 18 -0.4096127 0.686923512 miR34c
## 6 PTX - Mock   Processes  0.6675913 0.6053136 18  1.1028849 0.284604841 miR34c
##   p.adjusted p.signif Fold.Change
## 1 0.01308590        *   2.6273897
## 2 0.01447817        *   2.2678740
## 3 0.34152581       ns   1.2081995
## 4 0.01447817        *   1.6675620
## 5 0.68692351       ns   0.8420956
## 6 0.34152581       ns   1.5884187
```

### Figure2:

##### Figure2A: Spine Volume upon PTX-treatment and inhibition of different microRNAs

- Normalized Spine Volume of GFP and LNA (miRNA-inhibitor transfected) neurons that were treated with PTX or vehicle
- The value of every neuron was normalized to the mean of the corresponding negative control (NC) and mock-treated condition of every indepedent biological replicate
- Interaction of fixed-effects of LNA-treatment and PTX-treatment and random effect of independent-biological replicate (n=7, number of hippocampal preparations, 8-12 neurons per condition and experiment)
- Followed by Dunn's-Test where every condition was contrasted to:
  - First, the Mock condition within LNA-treatment
  - Second, the negative control (NC) within PTX-treatment

```
dat.spine <- allData$dat.spine
mod <- lmer(SpineHead ~ LNATreatment*PTXTreatment + (1|Experiment), dat.spine)
summary(mod)
```

```
## Linear mixed model fit by REML. t-tests use Satterthwaite's method [
## lmerModLmerTest]
## Formula: SpineHead ~ LNATreatment * PTXTreatment + (1 | Experiment)
##    Data: dat.spine
##
```

```
## REML criterion at convergence: -266
##
## Scaled residuals:
##     Min      1Q  Median      3Q     Max
## -4.8331 -0.6802 -0.0346  0.5192  4.5049
##
## Random effects:
##  Groups     Name        Variance Std.Dev.
##  Experiment (Intercept) 0.01419  0.1191
##  Residual               0.03372  0.1836
## Number of obs: 609, groups:  Experiment, 8
##
## Fixed effects:
##                                    Estimate Std. Error        df t value
## (Intercept)                         0.99142    0.04725  10.24252  20.983
## LNATreatmentpLNA129                -0.03109    0.03002 592.05241  -1.036
## LNATreatmentpLNA218                -0.09333    0.02951 592.14750  -3.162
## LNATreatmentpLNA34a                 0.01543    0.03498 594.31034   0.441
## LNATreatmentpLNA34c                 0.01522    0.03647 594.24887   0.417
## PTXTreatmentPTX                    -0.13204    0.03097 592.35320  -4.263
## LNATreatmentpLNA129:PTXTreatmentPTX  0.11529    0.04358 592.14377   2.646
## LNATreatmentpLNA218:PTXTreatmentPTX  0.16315    0.04256 592.25400   3.833
## LNATreatmentpLNA34a:PTXTreatmentPTX  0.03366    0.05125 592.11291   0.657
## LNATreatmentpLNA34c:PTXTreatmentPTX  0.08123    0.05130 592.15567   1.584
##                                    Pr(>|t|)
## (Intercept)                        9.43e-10 ***
## LNATreatmentpLNA129                 0.30080
## LNATreatmentpLNA218                 0.00165 **
## LNATreatmentpLNA34a                 0.65934
## LNATreatmentpLNA34c                 0.67664
## PTXTreatmentPTX                    2.35e-05 ***
## LNATreatmentpLNA129:PTXTreatmentPTX  0.00837 **
## LNATreatmentpLNA218:PTXTreatmentPTX  0.00014 ***
## LNATreatmentpLNA34a:PTXTreatmentPTX  0.51159
## LNATreatmentpLNA34c:PTXTreatmentPTX  0.11382
## ---
## Signif. codes:  0 '***' 0.001 '**' 0.01 '*' 0.05 '.' 0.1 ' ' 1
##
## Correlation of Fixed Effects:
##                          (Intr) LNATrLNA129 LNATrLNA218 LNATretmntpLNA34
## LNATrLNA129              -0.322
## LNATrLNA218              -0.327  0.516
## LNATretmntpLNA34         -0.273  0.435       0.446
## LNATrtmntpLNA34c         -0.262  0.417       0.425       0.405
## PTXTrtmnPTX              -0.314  0.493       0.501       0.409
## LNATLNA129:               0.223 -0.690      -0.357      -0.297
## LNATLNA218:               0.227 -0.359      -0.694      -0.301
## LNATrtmntpLNA34:PTXTrPTX  0.190 -0.298      -0.303      -0.639
## LNATrtmntpLNA34c:PTXTPTX  0.189 -0.298      -0.303      -0.246
##                          LNATrtmntpLNA34c PTXTPT LNATLNA129: LNATLNA218:
## LNATrLNA129
## LNATrLNA218
## LNATretmntpLNA34
## LNATrtmntpLNA34c
```

```
## PTXTrtmnPTX                0.391
## LNATLNA129:               -0.284            -0.710
## LNATLNA218:               -0.287            -0.727  0.517
## LNATrtmntpLNA34:PTXTrPTX -0.235            -0.604  0.429       0.439
## LNATrtmntpLNA34c:PTXTPTX -0.669            -0.604  0.428       0.438
##                          LNATrtmntpLNA34:PTXTrPTX
## LNATrLNA129
## LNATrLNA218
## LNATretmntpLNA34
## LNATrtmntpLNA34c
## PTXTrtmnPTX
## LNATLNA129:
## LNATLNA218:
## LNATrtmntpLNA34:PTXTrPTX
## LNATrtmntpLNA34c:PTXTPTX  0.364
```

```
emmeans(mod, trt.vs.ctrl ~ PTXTreatment | LNATreatment, ref="Mock")
```

```
## $emmeans
## LNATreatment = NC:
##  PTXTreatment emmean     SE    df lower.CL upper.CL
##  Mock          0.991 0.0472 10.32    0.887    1.096
##  PTX           0.859 0.0477 10.70    0.754    0.965
##
## LNATreatment = pLNA129:
##  PTXTreatment emmean     SE    df lower.CL upper.CL
##  Mock          0.960 0.0471 10.20    0.856    1.065
##  PTX           0.944 0.0477 10.69    0.838    1.049
##
## LNATreatment = pLNA218:
##  PTXTreatment emmean     SE    df lower.CL upper.CL
##  Mock          0.898 0.0468  9.94    0.794    1.002
##  PTX           0.929 0.0471 10.17    0.825    1.034
##
## LNATreatment = pLNA34a:
##  PTXTreatment emmean     SE    df lower.CL upper.CL
##  Mock          1.007 0.0505 13.44    0.898    1.116
##  PTX           0.908 0.0531 16.30    0.796    1.021
##
## LNATreatment = pLNA34c:
##  PTXTreatment emmean     SE    df lower.CL upper.CL
##  Mock          1.007 0.0516 14.58    0.896    1.117
##  PTX           0.956 0.0521 15.16    0.845    1.067
##
## Degrees-of-freedom method: kenward-roger
## Confidence level used: 0.95
##
## $contrasts
## LNATreatment = NC:
##  contrast   estimate     SE  df t.ratio p.value
##  PTX - Mock  -0.1320 0.0310 592  -4.263  <.0001
##
## LNATreatment = pLNA129:
##  contrast   estimate     SE  df t.ratio p.value
##  PTX - Mock  -0.0167 0.0307 592  -0.546  0.5856
```

```
##
## LNATreatment = pLNA218:
##  contrast   estimate     SE  df t.ratio p.value
##  PTX - Mock   0.0311 0.0292 592   1.064  0.2880
##
## LNATreatment = pLNA34a:
##  contrast   estimate     SE  df t.ratio p.value
##  PTX - Mock  -0.0984 0.0408 592  -2.409  0.0163
##
## LNATreatment = pLNA34c:
##  contrast   estimate     SE  df t.ratio p.value
##  PTX - Mock  -0.0508 0.0409 592  -1.242  0.2146
##
## Degrees-of-freedom method: kenward-roger
```

```
emmeans(mod, trt.vs.ctrl ~ LNATreatment | PTXTreatment, ref="NC")
```

```
## $emmeans
## PTXTreatment = Mock:
##  LNATreatment emmean     SE    df lower.CL upper.CL
##  NC            0.991 0.0472 10.32    0.887    1.096
##  pLNA129       0.960 0.0471 10.20    0.856    1.065
##  pLNA218       0.898 0.0468  9.94    0.794    1.002
##  pLNA34a       1.007 0.0505 13.44    0.898    1.116
##  pLNA34c       1.007 0.0516 14.58    0.896    1.117
##
## PTXTreatment = PTX:
##  LNATreatment emmean     SE    df lower.CL upper.CL
##  NC            0.859 0.0477 10.70    0.754    0.965
##  pLNA129       0.944 0.0477 10.69    0.838    1.049
##  pLNA218       0.929 0.0471 10.17    0.825    1.034
##  pLNA34a       0.908 0.0531 16.30    0.796    1.021
##  pLNA34c       0.956 0.0521 15.16    0.845    1.067
##
## Degrees-of-freedom method: kenward-roger
## Confidence level used: 0.95
##
## $contrasts
## PTXTreatment = Mock:
##  contrast      estimate     SE  df t.ratio p.value
##  pLNA129 - NC   -0.0311 0.0300 592  -1.036  0.6614
##  pLNA218 - NC   -0.0933 0.0295 592  -3.162  0.0062
##  pLNA34a - NC    0.0154 0.0350 594   0.441  0.9489
##  pLNA34c - NC    0.0152 0.0365 594   0.417  0.9551
##
## PTXTreatment = PTX:
##  contrast      estimate     SE  df t.ratio p.value
##  pLNA129 - NC    0.0842 0.0315 592   2.669  0.0283
##  pLNA218 - NC    0.0698 0.0307 592   2.277  0.0791
##  pLNA34a - NC    0.0491 0.0395 595   1.243  0.5281
##  pLNA34c - NC    0.0964 0.0382 595   2.523  0.0424
##
## Degrees-of-freedom method: kenward-roger
## P value adjustment: dunnettx method for 4 tests
```

Same Statistics for Spine Density found in Supplemental Figure 2A:

Spine Density: Number of Spines per Area of neuron

```
mod <- lmer(SpineDensity ~ LNATreatment*PTXTreatment + (1|Experiment), dat.spine)
summary(mod)
```

```
## Linear mixed model fit by REML. t-tests use Satterthwaite's method [
## lmerModLmerTest]
## Formula: SpineDensity ~ LNATreatment * PTXTreatment + (1 | Experiment)
##    Data: dat.spine
##
## REML criterion at convergence: -389
##
## Scaled residuals:
##     Min      1Q  Median      3Q     Max
## -3.6911 -0.6486  0.0205  0.6538  2.9158
##
## Random effects:
##  Groups     Name        Variance Std.Dev.
##  Experiment (Intercept) 0.002821 0.05311
##  Residual               0.027891 0.16701
## Number of obs: 609, groups:  Experiment, 8
##
## Fixed effects:
##                                      Estimate Std. Error         df t value
## (Intercept)                          1.005863   0.027051  24.584639  37.185
## LNATreatmentpLNA129                  0.020691   0.027301 592.927696   0.758
## LNATreatmentpLNA218                  0.053525   0.026834 593.247221   1.995
## LNATreatmentpLNA34a                  0.033788   0.031718 598.337444   1.065
## LNATreatmentpLNA34c                  0.033775   0.033068 598.264420   1.021
## PTXTreatmentPTX                      0.003464   0.028155 593.863443   0.123
## LNATreatmentpLNA129:PTXTreatmentPTX   0.055017   0.039621 593.218749   1.389
## LNATreatmentpLNA218:PTXTreatmentPTX  -0.040037   0.038696 593.558164  -1.035
## LNATreatmentpLNA34a:PTXTreatmentPTX  -0.050609   0.046603 593.122873  -1.086
## LNATreatmentpLNA34c:PTXTreatmentPTX  -0.033675   0.046639 593.263770  -0.722
##                                     Pr(>|t|)
## (Intercept)                           <2e-16 ***
## LNATreatmentpLNA129                   0.4488
## LNATreatmentpLNA218                   0.0465 *
## LNATreatmentpLNA34a                   0.2872
## LNATreatmentpLNA34c                   0.3075
## PTXTreatmentPTX                       0.9021
## LNATreatmentpLNA129:PTXTreatmentPTX   0.1655
## LNATreatmentpLNA218:PTXTreatmentPTX   0.3013
## LNATreatmentpLNA34a:PTXTreatmentPTX   0.2779
## LNATreatmentpLNA34c:PTXTreatmentPTX   0.4706
## ---
## Signif. codes:  0 '***' 0.001 '**' 0.01 '*' 0.05 '.' 0.1 ' ' 1
##
## Correlation of Fixed Effects:
##                          (Intr) LNATrLNA129 LNATrLNA218 LNATretmntpLNA34
## LNATrLNA129              -0.512
## LNATrLNA218              -0.520  0.516
## LNATretmntpLNA34         -0.436  0.436       0.447
```

```
## LNATrtmntpLNA34c          -0.417  0.418       0.426       0.402
## PTXTrtmnPTX               -0.498  0.493       0.501       0.412
## LNATLNA129:                0.354 -0.690      -0.357      -0.298
## LNATLNA218:                0.361 -0.359      -0.694      -0.303
## LNATrtmntpLNA34:PTXTrPTX   0.301 -0.298      -0.303      -0.642
## LNATrtmntpLNA34c:PTXTPTX   0.300 -0.298      -0.303      -0.248
##                           LNATrtmntpLNA34c PTXTPT LNATLNA129: LNATLNA218:
## LNATrLNA129
## LNATrLNA218
## LNATretmntpLNA34
## LNATrtmntpLNA34c
## PTXTrtmnPTX                0.394
## LNATLNA129:               -0.286           -0.710
## LNATLNA218:               -0.289           -0.727  0.517
## LNATrtmntpLNA34:PTXTrPTX -0.236           -0.604  0.428       0.439
## LNATrtmntpLNA34c:PTXTPTX -0.671           -0.604  0.428       0.438
##                           LNATrtmntpLNA34:PTXTrPTX
## LNATrLNA129
## LNATrLNA218
## LNATretmntpLNA34
## LNATrtmntpLNA34c
## PTXTrtmnPTX
## LNATLNA129:
## LNATLNA218:
## LNATrtmntpLNA34:PTXTrPTX
## LNATrtmntpLNA34c:PTXTPTX  0.364
```

```
emmeans(mod, trt.vs.ctrl ~ PTXTreatment | LNATreatment, ref="Mock")
```

```
## $emmeans
## LNATreatment = NC:
##  PTXTreatment emmean     SE   df lower.CL upper.CL
##  Mock          1.006 0.0271 22.6    0.950     1.06
##  PTX           1.009 0.0277 24.7    0.952     1.07
##
## LNATreatment = pLNA129:
##  PTXTreatment emmean     SE   df lower.CL upper.CL
##  Mock          1.027 0.0269 22.0    0.971     1.08
##  PTX           1.085 0.0276 24.7    1.028     1.14
##
## LNATreatment = pLNA218:
##  PTXTreatment emmean     SE   df lower.CL upper.CL
##  Mock          1.059 0.0264 20.6    1.004     1.11
##  PTX           1.023 0.0268 21.8    0.967     1.08
##
## LNATreatment = pLNA34a:
##  PTXTreatment emmean     SE   df lower.CL upper.CL
##  Mock          1.040 0.0315 39.1    0.976     1.10
##  PTX           0.993 0.0348 56.9    0.923     1.06
##
## LNATreatment = pLNA34c:
##  PTXTreatment emmean     SE   df lower.CL upper.CL
##  Mock          1.040 0.0329 45.9    0.973     1.11
##  PTX           1.009 0.0336 49.6    0.942     1.08
##
```

```
## Degrees-of-freedom method: kenward-roger
## Confidence level used: 0.95
##
## $contrasts
## LNATreatment = NC:
##  contrast   estimate     SE  df t.ratio p.value
##  PTX - Mock  0.00346 0.0282 593   0.123  0.9022
##
## LNATreatment = pLNA129:
##  contrast   estimate     SE  df t.ratio p.value
##  PTX - Mock  0.05848 0.0279 593   2.094  0.0366
##
## LNATreatment = pLNA218:
##  contrast   estimate     SE  df t.ratio p.value
##  PTX - Mock -0.03657 0.0266 593  -1.375  0.1696
##
## LNATreatment = pLNA34a:
##  contrast   estimate     SE  df t.ratio p.value
##  PTX - Mock -0.04714 0.0371 592  -1.269  0.2048
##
## LNATreatment = pLNA34c:
##  contrast   estimate     SE  df t.ratio p.value
##  PTX - Mock -0.03021 0.0372 592  -0.812  0.4169
##
## Degrees-of-freedom method: kenward-roger
```

```
emmeans(mod, trt.vs.ctrl ~ LNATreatment | PTXTreatment, ref="NC")
```

```
## $emmeans
## PTXTreatment = Mock:
##  LNATreatment emmean     SE   df lower.CL upper.CL
##  NC            1.006 0.0271 22.6    0.950     1.06
##  pLNA129       1.027 0.0269 22.0    0.971     1.08
##  pLNA218       1.059 0.0264 20.6    1.004     1.11
##  pLNA34a       1.040 0.0315 39.1    0.976     1.10
##  pLNA34c       1.040 0.0329 45.9    0.973     1.11
##
## PTXTreatment = PTX:
##  LNATreatment emmean     SE   df lower.CL upper.CL
##  NC            1.009 0.0277 24.7    0.952     1.07
##  pLNA129       1.085 0.0276 24.7    1.028     1.14
##  pLNA218       1.023 0.0268 21.8    0.967     1.08
##  pLNA34a       0.993 0.0348 56.9    0.923     1.06
##  pLNA34c       1.009 0.0336 49.6    0.942     1.08
##
## Degrees-of-freedom method: kenward-roger
## Confidence level used: 0.95
##
## $contrasts
## PTXTreatment = Mock:
##  contrast      estimate     SE  df t.ratio p.value
##  pLNA129 - NC  2.07e-02 0.0273 592   0.758  0.8231
##  pLNA218 - NC  5.35e-02 0.0268 593   1.994  0.1503
##  pLNA34a - NC  3.38e-02 0.0318 598   1.063  0.6441
##  pLNA34c - NC  3.38e-02 0.0331 598   1.019  0.6718
```

```
##
## PTXTreatment = PTX:
##  contrast      estimate     SE  df t.ratio p.value
##  pLNA129 - NC  7.57e-02 0.0287 593   2.639  0.0308
##  pLNA218 - NC  1.35e-02 0.0279 593   0.484  0.9364
##  pLNA34a - NC -1.68e-02 0.0359 599  -0.469  0.9409
##  pLNA34c - NC  9.94e-05 0.0347 599   0.003  1.0000
##
## Degrees-of-freedom method: kenward-roger
## P value adjustment: dunnettx method for 4 tests
```

**Figure 2B: GluA1/2 cluster size analysis upon miR-218-5p inhibition and PTX-treatment**

- GluA1 and GluA2 cluster size

```
dat.glua <- allData$dat.glua
mod <- lmer(NormArea ~ LNATreatment*Treatment*Second_Channel +  (1|Experiment),
            subset(dat.glua, Channel_Name == "GFP"))
summary(mod)
```

```
## Linear mixed model fit by REML. t-tests use Satterthwaite's method [
## lmerModLmerTest]
## Formula:
## NormArea ~ LNATreatment * Treatment * Second_Channel + (1 | Experiment)
##    Data: subset(dat.glua, Channel_Name == "GFP")
##
## REML criterion at convergence: 50.1
##
## Scaled residuals:
##     Min      1Q  Median      3Q     Max
## -2.2741 -0.6063 -0.1560  0.4487  5.4264
##
## Random effects:
##  Groups     Name        Variance Std.Dev.
##  Experiment (Intercept) 0.004982 0.07058
##  Residual               0.061248 0.24748
## Number of obs: 329, groups:  Experiment, 4
##
## Fixed effects:
##                                                 Estimate Std. Error
## (Intercept)                                    1.008e+00  4.981e-02
## LNATreatmentLNA218                             9.699e-03  5.376e-02
## TreatmentPTX                                  -2.291e-01  5.561e-02
## Second_ChannelGluA2                            1.642e-15  4.950e-02
## LNATreatmentLNA218:TreatmentPTX                1.239e-01  7.903e-02
## LNATreatmentLNA218:Second_ChannelGluA2        -1.539e-01  7.590e-02
## TreatmentPTX:Second_ChannelGluA2               6.317e-02  7.642e-02
## LNATreatmentLNA218:TreatmentPTX:Second_ChannelGluA2 -2.457e-02  1.102e-01
##                                                       df t value Pr(>|t|)
## (Intercept)                                    8.299e+00  20.239 2.34e-08
## LNATreatmentLNA218                             3.182e+02   0.180   0.8569
## TreatmentPTX                                   3.198e+02  -4.120 4.83e-05
## Second_ChannelGluA2                            3.178e+02   0.000   1.0000
## LNATreatmentLNA218:TreatmentPTX                3.190e+02   1.568   0.1179
```

```
## LNATreatmentLNA218:Second_ChannelGluA2              3.178e+02  -2.027   0.0435
## TreatmentPTX:Second_ChannelGluA2                    3.182e+02   0.827   0.4091
## LNATreatmentLNA218:TreatmentPTX:Second_ChannelGluA2  3.180e+02  -0.223   0.8237
##
## (Intercept)                                        ***
## LNATreatmentLNA218
## TreatmentPTX                                       ***
## Second_ChannelGluA2
## LNATreatmentLNA218:TreatmentPTX
## LNATreatmentLNA218:Second_ChannelGluA2             *
## TreatmentPTX:Second_ChannelGluA2
## LNATreatmentLNA218:TreatmentPTX:Second_ChannelGluA2
## ---
## Signif. codes:  0 '***' 0.001 '**' 0.01 '*' 0.05 '.' 0.1 ' ' 1
##
## Correlation of Fixed Effects:
##                  (Intr) LNATrLNA218 TrtPTX S_CGA2 LNATrLNA218:TPTX LNATLNA218:S
## LNATrLNA218      -0.461
## TreatmntPTX      -0.433  0.402
## Scnd_ChnGA2      -0.497  0.460       0.445
## LNATrLNA218:TPTX  0.305 -0.674      -0.703 -0.313
## LNATLNA218:S      0.324 -0.706      -0.290 -0.652  0.480
## TPTX:S_CGA2       0.319 -0.296      -0.719 -0.648  0.507            0.422
## LNATLNA218:TPTX: -0.221  0.485       0.499  0.449 -0.713           -0.689
##                  TPTX:S
## LNATrLNA218
## TreatmntPTX
## Scnd_ChnGA2
## LNATrLNA218:TPTX
## LNATLNA218:S
## TPTX:S_CGA2
## LNATLNA218:TPTX: -0.694
```

```
emmeans(mod, trt.vs.ctrl ~ Treatment | LNATreatment | Second_Channel, ref="Mock")
```

```
## $emmeans
## LNATreatment = NC, Second_Channel = GluA1:
##  Treatment emmean     SE    df lower.CL upper.CL
##  Mock       1.008 0.0498  8.74    0.895    1.121
##  PTX        0.779 0.0566 13.81    0.657    0.901
##
## LNATreatment = LNA218, Second_Channel = GluA1:
##  Treatment emmean     SE    df lower.CL upper.CL
##  Mock       1.018 0.0539 11.93    0.900    1.135
##  PTX        0.913 0.0524 10.67    0.797    1.028
##
## LNATreatment = NC, Second_Channel = GluA2:
##  Treatment emmean     SE    df lower.CL upper.CL
##  Mock       1.008 0.0498  8.74    0.895    1.121
##  PTX        0.842 0.0538 11.58    0.725    0.960
##
## LNATreatment = LNA218, Second_Channel = GluA2:
##  Treatment emmean     SE    df lower.CL upper.CL
##  Mock       0.864 0.0539 11.93    0.746    0.981
##  PTX        0.797 0.0524 10.67    0.682    0.913
```

```
##
## Degrees-of-freedom method: kenward-roger
## Confidence level used: 0.95
##
## $contrasts
## LNATreatment = NC, Second_Channel = GluA1:
##  contrast   estimate     SE  df t.ratio p.value
##  PTX - Mock  -0.2291 0.0557 320  -4.110  0.0001
##
## LNATreatment = LNA218, Second_Channel = GluA1:
##  contrast   estimate     SE  df t.ratio p.value
##  PTX - Mock  -0.1052 0.0562 318  -1.872  0.0622
##
## LNATreatment = NC, Second_Channel = GluA2:
##  contrast   estimate     SE  df t.ratio p.value
##  PTX - Mock  -0.1659 0.0532 319  -3.121  0.0020
##
## LNATreatment = LNA218, Second_Channel = GluA2:
##  contrast   estimate     SE  df t.ratio p.value
##  PTX - Mock  -0.0666 0.0562 318  -1.185  0.2369
##
## Degrees-of-freedom method: kenward-roger
```

```
emmeans(mod, trt.vs.ctrl ~ LNATreatment | Treatment | Second_Channel, ref="NC")
```

```
## $emmeans
## Treatment = Mock, Second_Channel = GluA1:
##  LNATreatment emmean     SE    df lower.CL upper.CL
##  NC            1.008 0.0498  8.74    0.895    1.121
##  LNA218        1.018 0.0539 11.93    0.900    1.135
##
## Treatment = PTX, Second_Channel = GluA1:
##  LNATreatment emmean     SE    df lower.CL upper.CL
##  NC            0.779 0.0566 13.81    0.657    0.901
##  LNA218        0.913 0.0524 10.67    0.797    1.028
##
## Treatment = Mock, Second_Channel = GluA2:
##  LNATreatment emmean     SE    df lower.CL upper.CL
##  NC            1.008 0.0498  8.74    0.895    1.121
##  LNA218        0.864 0.0539 11.93    0.746    0.981
##
## Treatment = PTX, Second_Channel = GluA2:
##  LNATreatment emmean     SE    df lower.CL upper.CL
##  NC            0.842 0.0538 11.58    0.725    0.960
##  LNA218        0.797 0.0524 10.67    0.682    0.913
##
## Degrees-of-freedom method: kenward-roger
## Confidence level used: 0.95
##
## $contrasts
## Treatment = Mock, Second_Channel = GluA1:
##  contrast    estimate     SE  df t.ratio p.value
##  LNA218 - NC   0.0097 0.0538 318   0.180  0.8570
##
## Treatment = PTX, Second_Channel = GluA1:
```

```
##  contrast    estimate     SE  df t.ratio p.value
##  LNA218 - NC   0.1336 0.0586 321   2.278  0.0234
##
## Treatment = Mock, Second_Channel = GluA2:
##  contrast    estimate     SE  df t.ratio p.value
##  LNA218 - NC  -0.1442 0.0538 318  -2.681  0.0077
##
## Treatment = PTX, Second_Channel = GluA2:
##  contrast    estimate     SE  df t.ratio p.value
##  LNA218 - NC  -0.0449 0.0559 320  -0.802  0.4231
##
## Degrees-of-freedom method: kenward-roger
```

- GluA1/GluA2 co-cluster density

```
mod <- lmer(NormNorm ~ LNATreatment*Treatment +  (1|Experiment),
            subset(dat.glua, Channel_Name != "GFP" & Second_Channel == "GluA2"))
summary(mod)
```

```
## Linear mixed model fit by REML. t-tests use Satterthwaite's method [
## lmerModLmerTest]
## Formula: NormNorm ~ LNATreatment * Treatment + (1 | Experiment)
##    Data: subset(dat.glua, Channel_Name != "GFP" & Second_Channel == "GluA2")
##
## REML criterion at convergence: 4.6
##
## Scaled residuals:
##     Min      1Q  Median      3Q     Max
## -2.3388 -0.5818 -0.0828  0.5167  3.3945
##
## Random effects:
##  Groups     Name        Variance Std.Dev.
##  Experiment (Intercept) 0.02452  0.1566
##  Residual               0.05203  0.2281
## Number of obs: 167, groups:  Experiment, 4
##
## Fixed effects:
##                                Estimate Std. Error        df t value Pr(>|t|)
## (Intercept)                     0.99217    0.08479   3.64183  11.702 0.000509
## LNATreatmentLNA218             -0.08836    0.04965 160.08315  -1.780 0.077031
## TreatmentPTX                   -0.28770    0.04920 160.30170  -5.847 2.72e-08
## LNATreatmentLNA218:TreatmentPTX  0.10984    0.07129 160.06228   1.541 0.125356
##
## (Intercept)                    ***
## LNATreatmentLNA218             .
## TreatmentPTX                   ***
## LNATreatmentLNA218:TreatmentPTX
## ---
## Signif. codes:  0 '***' 0.001 '**' 0.01 '*' 0.05 '.' 0.1 ' ' 1
##
## Correlation of Fixed Effects:
##             (Intr) LNATrLNA218 TrtPTX
## LNATrLNA218 -0.251
## TreatmntPTX -0.243  0.414
## LNATLNA218:  0.168 -0.685      -0.686
```

```r
emmeans(mod, trt.vs.ctrl ~ Treatment | LNATreatment, ref="Mock")
```

```
## $emmeans
## LNATreatment = NC:
##  Treatment emmean     SE   df lower.CL upper.CL
##  Mock       0.992 0.0848 3.71    0.749    1.235
##  PTX        0.704 0.0871 4.11    0.465    0.944
##
## LNATreatment = LNA218:
##  Treatment emmean     SE   df lower.CL upper.CL
##  Mock       0.904 0.0868 4.08    0.665    1.143
##  PTX        0.726 0.0861 3.93    0.485    0.966
##
## Degrees-of-freedom method: kenward-roger
## Confidence level used: 0.95
##
## $contrasts
## LNATreatment = NC:
##  contrast   estimate     SE  df t.ratio p.value
##  PTX - Mock   -0.288 0.0492 160  -5.843  <.0001
##
## LNATreatment = LNA218:
##  contrast   estimate     SE  df t.ratio p.value
##  PTX - Mock   -0.178 0.0519 160  -3.429  0.0008
##
## Degrees-of-freedom method: kenward-roger
```

```r
emmeans(mod, trt.vs.ctrl ~ LNATreatment | Treatment, ref="NC")
```

```
## $emmeans
## Treatment = Mock:
##  LNATreatment emmean     SE   df lower.CL upper.CL
##  NC            0.992 0.0848 3.71    0.749    1.235
##  LNA218        0.904 0.0868 4.08    0.665    1.143
##
## Treatment = PTX:
##  LNATreatment emmean     SE   df lower.CL upper.CL
##  NC            0.704 0.0871 4.11    0.465    0.944
##  LNA218        0.726 0.0861 3.93    0.485    0.966
##
## Degrees-of-freedom method: kenward-roger
## Confidence level used: 0.95
##
## $contrasts
## Treatment = Mock:
##  contrast    estimate     SE  df t.ratio p.value
##  LNA218 - NC  -0.0884 0.0497 160  -1.779  0.0771
##
## Treatment = PTX:
##  contrast    estimate     SE  df t.ratio p.value
##  LNA218 - NC   0.0215 0.0520 161   0.413  0.6800
##
## Degrees-of-freedom method: kenward-roger
```

**Figure 2C: Shank2 and Bassoon clustering**

Shank2 Cluster size

```
dat.shank2 <- allData$dat.shank2
mod <- lmer(NormArea ~ LNATreatment*Treatment + (1|Experiment),
            subset(dat.shank2, Second_Channel == "Shank2" & Channel_Name == "GFP"))
```

```
## boundary (singular) fit: see help('isSingular')
```

```
summary(mod)
```

```
## Linear mixed model fit by REML. t-tests use Satterthwaite's method [
## lmerModLmerTest]
## Formula: NormArea ~ LNATreatment * Treatment + (1 | Experiment)
##    Data: subset(dat.shank2, Second_Channel == "Shank2" & Channel_Name ==
##     "GFP")
##
## REML criterion at convergence: 24.7
##
## Scaled residuals:
##     Min      1Q  Median      3Q     Max
## -1.6736 -0.6422 -0.2148  0.4152  3.7470
##
## Random effects:
##  Groups     Name        Variance Std.Dev.
##  Experiment (Intercept) 0.00000  0.0000
##  Residual               0.06533  0.2556
## Number of obs: 109, groups:  Experiment, 3
##
## Fixed effects:
##                                 Estimate Std. Error        df t value Pr(>|t|)
## (Intercept)                      1.00000    0.04830 105.00000  20.702  < 2e-16
## LNATreatmentLNA218              -0.09826    0.06831 105.00000  -1.438  0.15331
## TreatmentPTX                    -0.18632    0.06894 105.00000  -2.703  0.00803
## LNATreatmentLNA218:TreatmentPTX  0.02676    0.09798 105.00000   0.273  0.78529
##
## (Intercept)                     ***
## LNATreatmentLNA218
## TreatmentPTX                    **
## LNATreatmentLNA218:TreatmentPTX
## ---
## Signif. codes:  0 '***' 0.001 '**' 0.01 '*' 0.05 '.' 0.1 ' ' 1
##
## Correlation of Fixed Effects:
##             (Intr) LNATrLNA218 TrtPTX
## LNATrLNA218 -0.707
## TreatmntPTX -0.701  0.495
## LNATLNA218:  0.493 -0.697      -0.704
## optimizer (nloptwrap) convergence code: 0 (OK)
## boundary (singular) fit: see help('isSingular')
```

```
emmeans(mod, trt.vs.ctrl ~ Treatment | LNATreatment, ref="Mock")
```

```
## $emmeans
## LNATreatment = NC:
```

```
##  Treatment emmean     SE   df lower.CL upper.CL
##  Mock       1.000 0.0483 25.9    0.901    1.099
##  PTX        0.814 0.0492 27.6    0.713    0.915
##
## LNATreatment = LNA218:
##  Treatment emmean     SE   df lower.CL upper.CL
##  Mock       0.902 0.0483 25.9    0.802    1.001
##  PTX        0.742 0.0502 29.2    0.640    0.845
##
## Degrees-of-freedom method: kenward-roger
## Confidence level used: 0.95
##
## $contrasts
## LNATreatment = NC:
##  contrast   estimate     SE  df t.ratio p.value
##  PTX - Mock   -0.186 0.0690 103  -2.702  0.0081
##
## LNATreatment = LNA218:
##  contrast   estimate     SE  df t.ratio p.value
##  PTX - Mock   -0.160 0.0696 103  -2.291  0.0240
##
## Degrees-of-freedom method: kenward-roger
```

```
emmeans(mod, trt.vs.ctrl ~ LNATreatment | Treatment, ref="NC")
```

```
## $emmeans
## Treatment = Mock:
##  LNATreatment emmean     SE   df lower.CL upper.CL
##  NC            1.000 0.0483 25.9    0.901    1.099
##  LNA218        0.902 0.0483 25.9    0.802    1.001
##
## Treatment = PTX:
##  LNATreatment emmean     SE   df lower.CL upper.CL
##  NC            0.814 0.0492 27.6    0.713    0.915
##  LNA218        0.742 0.0502 29.2    0.640    0.845
##
## Degrees-of-freedom method: kenward-roger
## Confidence level used: 0.95
##
## $contrasts
## Treatment = Mock:
##  contrast    estimate     SE  df t.ratio p.value
##  LNA218 - NC  -0.0983 0.0683 103  -1.438  0.1534
##
## Treatment = PTX:
##  contrast    estimate     SE  df t.ratio p.value
##  LNA218 - NC  -0.0715 0.0703 103  -1.018  0.3112
##
## Degrees-of-freedom method: kenward-roger
```

Bassoon Cluster Size

```
mod <- lmer(NormArea ~ LNATreatment*Treatment + (1|Experiment),
            subset(dat.shank2, Second_Channel == "Bassoon" & Channel_Name == "GFP"))
summary(mod)
```

```
## Linear mixed model fit by REML. t-tests use Satterthwaite's method [
## lmerModLmerTest]
## Formula: NormArea ~ LNATreatment * Treatment + (1 | Experiment)
##    Data: subset(dat.shank2, Second_Channel == "Bassoon" & Channel_Name ==
##     "GFP")
##
## REML criterion at convergence: -22.6
##
## Scaled residuals:
##     Min      1Q  Median      3Q     Max
## -1.8281 -0.6386 -0.1137  0.4019  4.2690
##
## Random effects:
##  Groups     Name        Variance Std.Dev.
##  Experiment (Intercept) 0.02634  0.1623
##  Residual               0.03914  0.1978
## Number of obs: 109, groups:  Experiment, 3
##
## Fixed effects:
##                                Estimate Std. Error        df t value Pr(>|t|)
## (Intercept)                     1.00004    0.10089   2.47570   9.913  0.00471
## LNATreatmentLNA218              0.23139    0.05288 102.99647   4.376  2.9e-05
## TreatmentPTX                   -0.06027    0.05337 102.99897  -1.129  0.26145
## LNATreatmentLNA218:TreatmentPTX  -0.11521    0.07584 102.99802  -1.519  0.13180
##
## (Intercept)                    **
## LNATreatmentLNA218             ***
## TreatmentPTX
## LNATreatmentLNA218:TreatmentPTX
## ---
## Signif. codes:  0 '***' 0.001 '**' 0.01 '*' 0.05 '.' 0.1 ' ' 1
##
## Correlation of Fixed Effects:
##             (Intr) LNATrLNA218 TrtPTX
## LNATrLNA218 -0.262
## TreatmntPTX -0.260  0.495
## LNATLNA218:  0.183 -0.697      -0.704
```

```
emmeans(mod, trt.vs.ctrl ~ Treatment | LNATreatment, ref="Mock")
```

```
## $emmeans
## LNATreatment = NC:
##  Treatment emmean    SE   df lower.CL upper.CL
##  Mock        1.00 0.101 2.48    0.637     1.36
##  PTX         0.94 0.101 2.50    0.579     1.30
##
## LNATreatment = LNA218:
##  Treatment emmean    SE   df lower.CL upper.CL
##  Mock        1.23 0.101 2.48    0.869     1.59
##  PTX         1.06 0.101 2.53    0.697     1.42
##
## Degrees-of-freedom method: kenward-roger
## Confidence level used: 0.95
##
## $contrasts
```

```
## LNATreatment = NC:
##  contrast   estimate     SE  df t.ratio p.value
##  PTX - Mock  -0.0603 0.0534 103  -1.129  0.2615
##
## LNATreatment = LNA218:
##  contrast   estimate     SE  df t.ratio p.value
##  PTX - Mock  -0.1755 0.0539 103  -3.256  0.0015
##
## Degrees-of-freedom method: kenward-roger
```

```
emmeans(mod, trt.vs.ctrl ~ LNATreatment | Treatment, ref="NC")
```

```
## $emmeans
## Treatment = Mock:
##  LNATreatment emmean    SE   df lower.CL upper.CL
##  NC             1.00 0.101 2.48    0.637     1.36
##  LNA218         1.23 0.101 2.48    0.869     1.59
##
## Treatment = PTX:
##  LNATreatment emmean    SE   df lower.CL upper.CL
##  NC             0.94 0.101 2.50    0.579     1.30
##  LNA218         1.06 0.101 2.53    0.697     1.42
##
## Degrees-of-freedom method: kenward-roger
## Confidence level used: 0.95
##
## $contrasts
## Treatment = Mock:
##  contrast    estimate     SE  df t.ratio p.value
##  LNA218 - NC    0.231 0.0529 103   4.376  <.0001
##
## Treatment = PTX:
##  contrast    estimate     SE  df t.ratio p.value
##  LNA218 - NC    0.116 0.0544 103   2.137  0.0350
##
## Degrees-of-freedom method: kenward-roger
```

Shank2-Bassoon co-cluster density

```
dat.shank2.density <- allData$dat.shank2.density
mod <- lmer(NormNorm ~ LNATreatment*Treatment + (1|Experiment), dat.shank2.density)
summary(mod)
```

```
## Linear mixed model fit by REML. t-tests use Satterthwaite's method [
## lmerModLmerTest]
## Formula: NormNorm ~ LNATreatment * Treatment + (1 | Experiment)
##    Data: dat.shank2.density
##
## REML criterion at convergence: 118.9
##
## Scaled residuals:
##     Min      1Q  Median      3Q     Max
## -1.6633 -0.7531 -0.1016  0.4972  3.6218
##
## Random effects:
##  Groups     Name        Variance Std.Dev.
```

```
##  Experiment (Intercept) 0.005376 0.07332
##  Residual               0.159430 0.39929
## Number of obs: 108, groups:  Experiment, 3
##
## Fixed effects:
##                                Estimate Std. Error        df t value Pr(>|t|)
## (Intercept)                     1.00117    0.08653  10.03415  11.570 3.99e-07
## LNATreatmentLNA218             -0.27485    0.10771 102.00469  -2.552 0.012198
## TreatmentPTX                   -0.39090    0.10771 102.00469  -3.629 0.000446
## LNATreatmentLNA218:TreatmentPTX  0.22180    0.15376 102.02287   1.443 0.152224
##
## (Intercept)                    ***
## LNATreatmentLNA218             *
## TreatmentPTX                   ***
## LNATreatmentLNA218:TreatmentPTX
## ---
## Signif. codes:  0 '***' 0.001 '**' 0.01 '*' 0.05 '.' 0.1 ' ' 1
##
## Correlation of Fixed Effects:
##             (Intr) LNATrLNA218 TrtPTX
## LNATrLNA218 -0.611
## TreatmntPTX -0.611  0.491
## LNATLNA218:  0.428 -0.701      -0.701
```

```
emmeans(mod, trt.vs.ctrl ~ Treatment | LNATreatment, ref="Mock")
```

```
## $emmeans
## LNATreatment = NC:
##  Treatment emmean     SE   df lower.CL upper.CL
##  Mock       1.001 0.0866 10.2    0.809    1.194
##  PTX        0.610 0.0877 10.7    0.417    0.804
##
## LNATreatment = LNA218:
##  Treatment emmean     SE   df lower.CL upper.CL
##  Mock       0.726 0.0877 10.7    0.533    0.920
##  PTX        0.557 0.0891 11.3    0.362    0.753
##
## Degrees-of-freedom method: kenward-roger
## Confidence level used: 0.95
##
## $contrasts
## LNATreatment = NC:
##  contrast   estimate    SE  df t.ratio p.value
##  PTX - Mock   -0.391 0.108 102  -3.629  0.0004
##
## LNATreatment = LNA218:
##  contrast   estimate    SE  df t.ratio p.value
##  PTX - Mock   -0.169 0.110 102  -1.541  0.1264
##
## Degrees-of-freedom method: kenward-roger
```

```
emmeans(mod, trt.vs.ctrl ~ LNATreatment | Treatment, ref="NC")
```

```
## $emmeans
## Treatment = Mock:
```

```
##  LNATreatment emmean     SE   df lower.CL upper.CL
##  NC            1.001 0.0866 10.2    0.809    1.194
##  LNA218        0.726 0.0877 10.7    0.533    0.920
##
## Treatment = PTX:
##  LNATreatment emmean     SE   df lower.CL upper.CL
##  NC            0.610 0.0877 10.7    0.417    0.804
##  LNA218        0.557 0.0891 11.3    0.362    0.753
##
## Degrees-of-freedom method: kenward-roger
## Confidence level used: 0.95
##
## $contrasts
## Treatment = Mock:
##  contrast    estimate    SE  df t.ratio p.value
##  LNA218 - NC  -0.2749 0.108 102  -2.551  0.0122
##
## Treatment = PTX:
##  contrast    estimate    SE  df t.ratio p.value
##  LNA218 - NC  -0.0531 0.110 102  -0.483  0.6298
##
## Degrees-of-freedom method: kenward-roger
```

**Figure 2E: Luciferase Assay of Shank2-UTR**

```
luc <- allData$luc
mod <- lmer(log2(Value) ~ Genotype*DuplexTreatment + (1|Experiment),
            subset(luc, Construct == "Shank2"))
summary(mod)
```

```
## Linear mixed model fit by REML. t-tests use Satterthwaite's method [
## lmerModLmerTest]
## Formula: log2(Value) ~ Genotype * DuplexTreatment + (1 | Experiment)
##    Data: subset(luc, Construct == "Shank2")
##
## REML criterion at convergence: -0.4
##
## Scaled residuals:
##      Min       1Q   Median       3Q      Max
## -1.07773 -0.57921 -0.07564  0.53406  1.30038
##
## Random effects:
##  Groups     Name        Variance Std.Dev.
##  Experiment (Intercept) 0.01874  0.1369
##  Residual               0.02239  0.1496
## Number of obs: 12, groups:  Experiment, 3
##
## Fixed effects:
##                                          Estimate Std. Error      df t value
## (Intercept)                               -2.7240     0.1171  4.9290 -23.264
## GenotypeMutant                             0.1309     0.1222  6.0000   1.072
## DuplexTreatmentDuplex218                  -0.4238     0.1222  6.0000  -3.469
## GenotypeMutant:DuplexTreatmentDuplex218    0.2372     0.1728  6.0000   1.373
```

```
##                                          Pr(>|t|)
## (Intercept)                              3.14e-06 ***
## GenotypeMutant                             0.3251
## DuplexTreatmentDuplex218                   0.0133 *
## GenotypeMutant:DuplexTreatmentDuplex218    0.2189
## ---
## Signif. codes:  0 '***' 0.001 '**' 0.01 '*' 0.05 '.' 0.1 ' ' 1
##
## Correlation of Fixed Effects:
##             (Intr) GntypM DTD218
## GenotypMtnt -0.522
## DplxTrtD218 -0.522  0.500
## GntM:DTD218  0.369 -0.707 -0.707
```

```
emmeans(mod, trt.vs.ctrl ~ DuplexTreatment | Genotype, ref="NC")
```

```
## $emmeans
## Genotype = WT:
##  DuplexTreatment emmean    SE   df lower.CL upper.CL
##  NC               -2.72 0.117 4.93    -3.03    -2.42
##  Duplex218        -3.15 0.117 4.93    -3.45    -2.85
##
## Genotype = Mutant:
##  DuplexTreatment emmean    SE   df lower.CL upper.CL
##  NC               -2.59 0.117 4.93    -2.90    -2.29
##  Duplex218        -2.78 0.117 4.93    -3.08    -2.48
##
## Degrees-of-freedom method: kenward-roger
## Results are given on the log2 (not the response) scale.
## Confidence level used: 0.95
##
## $contrasts
## Genotype = WT:
##  contrast       estimate    SE df t.ratio p.value
##  Duplex218 - NC   -0.424 0.122  6  -3.469  0.0133
##
## Genotype = Mutant:
##  contrast       estimate    SE df t.ratio p.value
##  Duplex218 - NC   -0.187 0.122  6  -1.528  0.1774
##
## Degrees-of-freedom method: kenward-roger
## Results are given on the log2 (not the response) scale.
```

```
emmeans(mod, trt.vs.ctrl ~ Genotype | DuplexTreatment, ref="WT")
```

```
## $emmeans
## DuplexTreatment = NC:
##  Genotype emmean    SE   df lower.CL upper.CL
##  WT        -2.72 0.117 4.93    -3.03    -2.42
##  Mutant    -2.59 0.117 4.93    -2.90    -2.29
##
## DuplexTreatment = Duplex218:
##  Genotype emmean    SE   df lower.CL upper.CL
##  WT        -3.15 0.117 4.93    -3.45    -2.85
##  Mutant    -2.78 0.117 4.93    -3.08    -2.48
```

```
##
## Degrees-of-freedom method: kenward-roger
## Results are given on the log2 (not the response) scale.
## Confidence level used: 0.95
##
## $contrasts
## DuplexTreatment = NC:
##  contrast    estimate    SE df t.ratio p.value
##  Mutant - WT    0.131 0.122  6   1.072  0.3251
##
## DuplexTreatment = Duplex218:
##  contrast    estimate    SE df t.ratio p.value
##  Mutant - WT    0.368 0.122  6   3.013  0.0236
##
## Degrees-of-freedom method: kenward-roger
## Results are given on the log2 (not the response) scale.
```

### Figure 3

##### Figure 3E: qPCR of miR-218-5p and miR-129-5p in LNA-injected cortex

```
qpcr <- allData$qpcr

mod <- lmer(-dCq ~ LNATreatment*miR.Expression + (1|MouseID), qpcr )
```

```
## boundary (singular) fit: see help('isSingular')
```

```
summary(mod)
```

```
## Linear mixed model fit by REML. t-tests use Satterthwaite's method [
## lmerModLmerTest]
## Formula: -dCq ~ LNATreatment * miR.Expression + (1 | MouseID)
##    Data: qpcr
##
## REML criterion at convergence: 58.5
##
## Scaled residuals:
##      Min       1Q   Median       3Q      Max
## -1.58250 -0.62236  0.02442  0.27255  2.38184
##
## Random effects:
##  Groups   Name        Variance  Std.Dev.
##  MouseID  (Intercept) 2.634e-22 1.623e-11
##  Residual             3.823e-01 6.183e-01
## Number of obs: 32, groups:  MouseID, 8
##
## Fixed effects:
##                                         Estimate Std. Error      df t value
## (Intercept)                              -0.2182     0.2186 26.0000  -0.998
## LNATreatmentLNA218                       -1.3826     0.3786 26.0000  -3.652
## LNATreatmentLNA129                       -0.1954     0.3786 26.0000  -0.516
## miR.ExpressionmiR129                     -6.2428     0.3091 26.0000 -20.194
## LNATreatmentLNA218:miR.ExpressionmiR129   2.1503     0.5355 26.0000   4.016
```

```
## LNATreatmentLNA129:miR.ExpressionmiR129  -1.7451     0.5355 26.0000  -3.259
##                                          Pr(>|t|)
## (Intercept)                              0.327415
## LNATreatmentLNA218                       0.001151 **
## LNATreatmentLNA129                       0.610186
## miR.ExpressionmiR129                      < 2e-16 ***
## LNATreatmentLNA218:miR.ExpressionmiR129 0.000449 ***
## LNATreatmentLNA129:miR.ExpressionmiR129 0.003111 **
## ---
## Signif. codes:  0 '***' 0.001 '**' 0.01 '*' 0.05 '.' 0.1 ' ' 1
##
## Correlation of Fixed Effects:
##             (Intr) LNATrLNA218 LNATrLNA129 mR.ER1 LNATLNA218:
## LNATrLNA218 -0.577
## LNATrLNA129 -0.577  0.333
## mR.ExprR129 -0.707  0.408       0.408
## LNATLNA218:  0.408 -0.707      -0.236      -0.577
## LNATLNA129:  0.408 -0.236      -0.707      -0.577  0.333
## optimizer (nloptwrap) convergence code: 0 (OK)
## boundary (singular) fit: see help('isSingular')
```

```
emmeans(mod, trt.vs.ctrl ~ LNATreatment | miR.Expression, ref="NC")
```

```
## $emmeans
## miR.Expression = miR218:
##  LNATreatment emmean    SE df lower.CL upper.CL
##  NC           -0.218 0.219 26   -0.668    0.231
##  LNA218       -1.601 0.317 26   -2.253   -0.948
##  LNA129       -0.414 0.317 26   -1.066    0.239
##
## miR.Expression = miR129:
##  LNATreatment emmean    SE df lower.CL upper.CL
##  NC           -6.461 0.219 26   -6.910   -6.012
##  LNA218       -5.693 0.317 26   -6.346   -5.041
##  LNA129       -8.401 0.317 26   -9.054   -7.749
##
## Degrees-of-freedom method: kenward-roger
## Confidence level used: 0.95
##
## $contrasts
## miR.Expression = miR218:
##  contrast    estimate    SE   df t.ratio p.value
##  LNA218 - NC   -1.383 0.385 22.5  -3.588  0.0031
##  LNA129 - NC   -0.195 0.385 22.5  -0.507  0.8210
##
## miR.Expression = miR129:
##  contrast    estimate    SE   df t.ratio p.value
##  LNA218 - NC    0.768 0.385 22.5   1.992  0.1078
##  LNA129 - NC   -1.940 0.385 22.5  -5.035  0.0001
##
## Degrees-of-freedom method: kenward-roger
## P value adjustment: dunnettx method for 2 tests
```

### Figure 4

##### Figure 4B: Luciferase Assay of Mdga1 3'-UTR

```
luc.mdga1 <- allData$luc.mdga1

mod <- lmer(log2(Value) ~ Genotype*DuplexTreatment + (1|Experiment),
            subset(luc.mdga1, Construct == "Mdga1"))
summary(mod)
```

```
## Linear mixed model fit by REML. t-tests use Satterthwaite's method [
## lmerModLmerTest]
## Formula: log2(Value) ~ Genotype * DuplexTreatment + (1 | Experiment)
##    Data: subset(luc.mdga1, Construct == "Mdga1")
##
## REML criterion at convergence: -7.6
##
## Scaled residuals:
##     Min      1Q  Median      3Q     Max
## -1.0230 -0.5392 -0.2024  0.5570  1.2510
##
## Random effects:
##  Groups     Name        Variance Std.Dev.
##  Experiment (Intercept) 0.011276 0.10619
##  Residual               0.008221 0.09067
## Number of obs: 12, groups:  Experiment, 3
##
## Fixed effects:
##                                          Estimate Std. Error       df t value
## (Intercept)                              -3.27047    0.08062  3.99319 -40.568
## GenotypeMutant                            0.01157    0.07403  6.00000   0.156
## DuplexTreatmentDuplex218                 -0.47437    0.07403  6.00000  -6.407
## GenotypeMutant:DuplexTreatmentDuplex218  0.34353    0.10470  6.00000   3.281
##                                          Pr(>|t|)
## (Intercept)                              2.25e-06 ***
## GenotypeMutant                           0.880982
## DuplexTreatmentDuplex218                 0.000681 ***
## GenotypeMutant:DuplexTreatmentDuplex218 0.016799 *
## ---
## Signif. codes:  0 '***' 0.001 '**' 0.01 '*' 0.05 '.' 0.1 ' ' 1
##
## Correlation of Fixed Effects:
##             (Intr) GntypM DTD218
## GenotypMtnt -0.459
## DplxTrtD218 -0.459  0.500
## GntM:DTD218  0.325 -0.707 -0.707
```

```
emmeans(mod, trt.vs.ctrl ~ DuplexTreatment | Genotype, ref="NC")
```

```
## $emmeans
## Genotype = WT:
##  DuplexTreatment emmean     SE   df lower.CL upper.CL
##  NC               -3.27 0.0806 3.99    -3.49    -3.05
##  Duplex218        -3.74 0.0806 3.99    -3.97    -3.52
```

```
##
## Genotype = Mutant:
##  DuplexTreatment emmean     SE   df lower.CL upper.CL
##  NC                -3.26 0.0806 3.99    -3.48    -3.03
##  Duplex218         -3.39 0.0806 3.99    -3.61    -3.17
##
## Degrees-of-freedom method: kenward-roger
## Results are given on the log2 (not the response) scale.
## Confidence level used: 0.95
##
## $contrasts
## Genotype = WT:
##  contrast        estimate    SE df t.ratio p.value
##  Duplex218 - NC   -0.474 0.074  6  -6.407  0.0007
##
## Genotype = Mutant:
##  contrast        estimate    SE df t.ratio p.value
##  Duplex218 - NC   -0.131 0.074  6  -1.767  0.1276
##
## Degrees-of-freedom method: kenward-roger
## Results are given on the log2 (not the response) scale.
```

```
emmeans(mod, trt.vs.ctrl ~ Genotype, ref="WT")
```

```
## NOTE: Results may be misleading due to involvement in interactions
```

```
## $emmeans
##  Genotype emmean     SE   df lower.CL upper.CL
##  WT        -3.51 0.0716 2.64    -3.75    -3.26
##  Mutant    -3.32 0.0716 2.64    -3.57    -3.08
##
## Results are averaged over the levels of: DuplexTreatment
## Degrees-of-freedom method: kenward-roger
## Results are given on the log2 (not the response) scale.
## Confidence level used: 0.95
##
## $contrasts
##  contrast    estimate     SE df t.ratio p.value
##  Mutant - WT    0.183 0.0523  6   3.502  0.0128
##
## Results are averaged over the levels of: DuplexTreatment
## Degrees-of-freedom method: kenward-roger
## Results are given on the log2 (not the response) scale.
```

##### Figure 4E: Puromycin-proximity ligation assay (PLA) of Mdga1 nascent peptide

-Foci Density

```
dat.duolink.Mdga1 <- allData$dat.duolink.Mdga1
mod <- lmer(NormNorm ~ LNATreatment*Selection+(1|Experiment), dat.duolink.Mdga1)
summary(mod)
```

```
## Linear mixed model fit by REML. t-tests use Satterthwaite's method [
## lmerModLmerTest]
## Formula: NormNorm ~ LNATreatment * Selection + (1 | Experiment)
```

```
##    Data: dat.duolink.Mdga1
##
## REML criterion at convergence: 427.1
##
## Scaled residuals:
##     Min      1Q  Median      3Q     Max
## -1.4531 -0.6115 -0.1131  0.2425  5.1916
##
## Random effects:
##  Groups     Name        Variance Std.Dev.
##  Experiment (Intercept) 0.06853  0.2618
##  Residual               1.61299  1.2700
## Number of obs: 128, groups:  Experiment, 3
##
## Fixed effects:
##                                          Estimate Std. Error         df t value
## (Intercept)                              0.973364   0.284112   9.780419   3.426
## LNATreatmentLNA218                       0.790254   0.327508 122.854407   2.413
## SelectionDendrites                       0.008885   0.336529 121.995079   0.026
## LNATreatmentLNA218:SelectionDendrites  -0.529017   0.452316 121.990711  -1.170
##                                        Pr(>|t|)
## (Intercept)                             0.00669 **
## LNATreatmentLNA218                      0.01730 *
## SelectionDendrites                      0.97898
## LNATreatmentLNA218:SelectionDendrites  0.24445
## ---
## Signif. codes:  0 '***' 0.001 '**' 0.01 '*' 0.05 '.' 0.1 ' ' 1
##
## Correlation of Fixed Effects:
##             (Intr) LNATrLNA218 SlctnD
## LNATrLNA218 -0.624
## SlctnDndrts -0.603  0.524
## LNATLNA218:  0.449 -0.720      -0.744
```

```
emmeans(mod, trt.vs.ctrl ~ LNATreatment | Selection, ref="NC")
```

```
## $emmeans
## Selection = Somata:
##  LNATreatment emmean    SE   df lower.CL upper.CL
##  NC            0.973 0.285 9.94    0.339     1.61
##  LNA218        1.764 0.269 7.98    1.144     2.38
##
## Selection = Dendrites:
##  LNATreatment emmean    SE   df lower.CL upper.CL
##  NC            0.982 0.281 9.49    0.352     1.61
##  LNA218        1.243 0.256 6.65    0.631     1.86
##
## Degrees-of-freedom method: kenward-roger
## Confidence level used: 0.95
##
## $contrasts
## Selection = Somata:
##  contrast    estimate    SE  df t.ratio p.value
##  LNA218 - NC    0.790 0.329 123   2.403  0.0178
##
```

```
## Selection = Dendrites:
##  contrast    estimate    SE  df t.ratio p.value
##  LNA218 - NC    0.261 0.315 123   0.830  0.4084
##
## Degrees-of-freedom method: kenward-roger
```

-Foci Area

```
mod <- lmer(NormArea ~ LNATreatment*Selection+(1|Experiment), dat.duolink.Mdga1)
```

```
## boundary (singular) fit: see help('isSingular')
```

```
summary(mod)
```

```
## Linear mixed model fit by REML. t-tests use Satterthwaite's method [
## lmerModLmerTest]
## Formula: NormArea ~ LNATreatment * Selection + (1 | Experiment)
##    Data: dat.duolink.Mdga1
##
## REML criterion at convergence: 106.3
##
## Scaled residuals:
##     Min      1Q  Median      3Q     Max
## -1.8996 -0.5898 -0.1575  0.4922  5.3552
##
## Random effects:
##  Groups     Name        Variance Std.Dev.
##  Experiment (Intercept) 0.0000   0.0000
##  Residual               0.1234   0.3513
## Number of obs: 128, groups:  Experiment, 3
##
## Fixed effects:
##                                          Estimate Std. Error         df t value
## (Intercept)                             1.000e+00  6.638e-02  1.240e+02  15.064
## LNATreatmentLNA218                      1.889e-01  9.025e-02  1.240e+02   2.093
## SelectionDendrites                      1.381e-15  9.307e-02  1.240e+02   0.000
## LNATreatmentLNA218:SelectionDendrites -3.713e-02  1.251e-01  1.240e+02  -0.297
##                                       Pr(>|t|)
## (Intercept)                             <2e-16 ***
## LNATreatmentLNA218                      0.0384 *
## SelectionDendrites                      1.0000
## LNATreatmentLNA218:SelectionDendrites   0.7671
## ---
## Signif. codes:  0 '***' 0.001 '**' 0.01 '*' 0.05 '.' 0.1 ' ' 1
##
## Correlation of Fixed Effects:
##             (Intr) LNATrLNA218 SlctnD
## LNATrLNA218 -0.736
## SlctnDndrts -0.713  0.525
## LNATLNA218:  0.531 -0.722      -0.744
## optimizer (nloptwrap) convergence code: 0 (OK)
## boundary (singular) fit: see help('isSingular')
```

```
emmeans(mod, trt.vs.ctrl ~ LNATreatment | Selection, ref="NC")
```

```
## $emmeans
## Selection = Somata:
```

```
##  LNATreatment emmean     SE   df lower.CL upper.CL
##  NC              1.00 0.0669 32.3    0.864     1.14
##  LNA218          1.19 0.0615 25.1    1.062     1.32
##
## Selection = Dendrites:
##  LNATreatment emmean     SE   df lower.CL upper.CL
##  NC              1.00 0.0655 31.3    0.867     1.13
##  LNA218          1.15 0.0573 19.8    1.032     1.27
##
## Degrees-of-freedom method: kenward-roger
## Confidence level used: 0.95
##
## $contrasts
## Selection = Somata:
##  contrast    estimate     SE  df t.ratio p.value
##  LNA218 - NC    0.189 0.0913 124   2.070  0.0405
##
## Selection = Dendrites:
##  contrast    estimate     SE  df t.ratio p.value
##  LNA218 - NC    0.152 0.0873 124   1.738  0.0847
##
## Degrees-of-freedom method: kenward-roger
```

##### Figure 4H: Proximity ligation assay (PLA) of Mdga1 and Nlgn2 proteins

- Foci Density

```
dat.duolink.Mdga1xNlgn2 <- allData$dat.duolink.Mdga1xNlgn2

mod <- lmer(NormNorm ~ LNATreatment*Selection+(1|Experiment), dat.duolink.Mdga1xNlgn2)
summary(mod)
```

```
## Linear mixed model fit by REML. t-tests use Satterthwaite's method [
## lmerModLmerTest]
## Formula: NormNorm ~ LNATreatment * Selection + (1 | Experiment)
##    Data: dat.duolink.Mdga1xNlgn2
##
## REML criterion at convergence: 427.1
##
## Scaled residuals:
##     Min      1Q  Median      3Q     Max
## -1.4531 -0.6115 -0.1131  0.2425  5.1916
##
## Random effects:
##  Groups     Name        Variance Std.Dev.
##  Experiment (Intercept) 0.06853  0.2618
##  Residual               1.61299  1.2700
## Number of obs: 128, groups:  Experiment, 3
##
## Fixed effects:
##                                         Estimate Std. Error         df t value
## (Intercept)                             0.973364   0.284112   9.780419   3.426
## LNATreatmentLNA218                      0.790254   0.327508 122.854407   2.413
## SelectionDendrites                      0.008885   0.336529 121.995079   0.026
```

```
## LNATreatmentLNA218:SelectionDendrites  -0.529017   0.452316 121.990711  -1.170
##                                        Pr(>|t|)
## (Intercept)                             0.00669 **
## LNATreatmentLNA218                      0.01730 *
## SelectionDendrites                      0.97898
## LNATreatmentLNA218:SelectionDendrites  0.24445
## ---
## Signif. codes:  0 '***' 0.001 '**' 0.01 '*' 0.05 '.' 0.1 ' ' 1
##
## Correlation of Fixed Effects:
##             (Intr) LNATrLNA218 SlctnD
## LNATrLNA218 -0.624
## SlctnDndrts -0.603  0.524
## LNATLNA218:  0.449 -0.720      -0.744
```

```
emmeans(mod, trt.vs.ctrl ~ LNATreatment | Selection, ref="NC")
```

```
## $emmeans
## Selection = Somata:
##  LNATreatment emmean    SE   df lower.CL upper.CL
##  NC            0.973 0.285 9.94    0.339     1.61
##  LNA218        1.764 0.269 7.98    1.144     2.38
##
## Selection = Dendrites:
##  LNATreatment emmean    SE   df lower.CL upper.CL
##  NC            0.982 0.281 9.49    0.352     1.61
##  LNA218        1.243 0.256 6.65    0.631     1.86
##
## Degrees-of-freedom method: kenward-roger
## Confidence level used: 0.95
##
## $contrasts
## Selection = Somata:
##  contrast    estimate    SE  df t.ratio p.value
##  LNA218 - NC    0.790 0.329 123   2.403  0.0178
##
## Selection = Dendrites:
##  contrast    estimate    SE  df t.ratio p.value
##  LNA218 - NC    0.261 0.315 123   0.830  0.4084
##
## Degrees-of-freedom method: kenward-roger
```

- Foci Area

```
mod <- lmer(NormArea ~ LNATreatment*Selection +(1|Experiment), dat.duolink.Mdga1xNlgn2)
```

```
## boundary (singular) fit: see help('isSingular')
```

```
summary(mod)
```

```
## Linear mixed model fit by REML. t-tests use Satterthwaite's method [
## lmerModLmerTest]
## Formula: NormArea ~ LNATreatment * Selection + (1 | Experiment)
##    Data: dat.duolink.Mdga1xNlgn2
##
## REML criterion at convergence: 106.3
##
```

```
## Scaled residuals:
##     Min      1Q  Median      3Q     Max
## -1.8996 -0.5898 -0.1575  0.4922  5.3552
##
## Random effects:
##  Groups     Name        Variance Std.Dev.
##  Experiment (Intercept) 0.0000   0.0000
##  Residual               0.1234   0.3513
## Number of obs: 128, groups:  Experiment, 3
##
## Fixed effects:
##                                       Estimate Std. Error         df t value
## (Intercept)                          1.000e+00  6.638e-02  1.240e+02  15.064
## LNATreatmentLNA218                   1.889e-01  9.025e-02  1.240e+02   2.093
## SelectionDendrites                   1.381e-15  9.307e-02  1.240e+02   0.000
## LNATreatmentLNA218:SelectionDendrites -3.713e-02  1.251e-01  1.240e+02  -0.297
##                                      Pr(>|t|)
## (Intercept)                            <2e-16 ***
## LNATreatmentLNA218                     0.0384 *
## SelectionDendrites                     1.0000
## LNATreatmentLNA218:SelectionDendrites   0.7671
## ---
## Signif. codes:  0 '***' 0.001 '**' 0.01 '*' 0.05 '.' 0.1 ' ' 1
##
## Correlation of Fixed Effects:
##             (Intr) LNATrLNA218 SlctnD
## LNATrLNA218 -0.736
## SlctnDndrts -0.713  0.525
## LNATLNA218:  0.531 -0.722      -0.744
## optimizer (nloptwrap) convergence code: 0 (OK)
## boundary (singular) fit: see help('isSingular')
```

```
emmeans(mod, trt.vs.ctrl ~ LNATreatment | Selection, ref="NC")
```

```
## $emmeans
## Selection = Somata:
##  LNATreatment emmean     SE   df lower.CL upper.CL
##  NC             1.00 0.0669 32.3    0.864     1.14
##  LNA218         1.19 0.0615 25.1    1.062     1.32
##
## Selection = Dendrites:
##  LNATreatment emmean     SE   df lower.CL upper.CL
##  NC             1.00 0.0655 31.3    0.867     1.13
##  LNA218         1.15 0.0573 19.8    1.032     1.27
##
## Degrees-of-freedom method: kenward-roger
## Confidence level used: 0.95
##
## $contrasts
## Selection = Somata:
##  contrast    estimate     SE  df t.ratio p.value
##  LNA218 - NC    0.189 0.0913 124   2.070  0.0405
##
## Selection = Dendrites:
##  contrast    estimate     SE  df t.ratio p.value
```

```
##  LNA218 - NC    0.152 0.0873 124   1.738  0.0847
##
## Degrees-of-freedom method: kenward-roger
```

#### Figure 5:

##### Figure 5B: Gamma2-Cluster size upon PTX-Treatment and miR-218-5p inhibition

- Gamma2- Cluster Area

```
dat.gamma2 <- allData$dat.gamma2

mod <- lmer(log2(NormArea) ~ LNATreatment * Treatment * Selection + (1|Experiment), dat.gamma2)
summary(mod)
```

```
## Linear mixed model fit by REML. t-tests use Satterthwaite's method [
## lmerModLmerTest]
## Formula: log2(NormArea) ~ LNATreatment * Treatment * Selection + (1 |
##     Experiment)
##    Data: dat.gamma2
##
## REML criterion at convergence: 283.6
##
## Scaled residuals:
##     Min      1Q  Median      3Q     Max
## -3.0141 -0.6924  0.0059  0.6401  5.2082
##
## Random effects:
##  Groups     Name        Variance Std.Dev.
##  Experiment (Intercept) 0.05106  0.2260
##  Residual               0.18811  0.4337
## Number of obs: 223, groups:  Experiment, 3
##
## Fixed effects:
##                                                    Estimate Std. Error
## (Intercept)                                        -0.04261    0.15679
## LNATreatmentLNA218                                  0.31480    0.11944
## TreatmentPTX                                        0.27053    0.12042
## SelectionDendrites                                  0.01186    0.12267
## LNATreatmentLNA218:TreatmentPTX                    -0.13233    0.16525
## LNATreatmentLNA218:SelectionDendrites               0.14915    0.16877
## TreatmentPTX:SelectionDendrites                     0.20402    0.17024
## LNATreatmentLNA218:TreatmentPTX:SelectionDendrites  -0.29284    0.23317
##                                                          df t value Pr(>|t|)
## (Intercept)                                         3.76142  -0.272  0.80008
## LNATreatmentLNA218                                213.02630   2.636  0.00902
## TreatmentPTX                                      213.00736   2.247  0.02570
## SelectionDendrites                                212.99157   0.097  0.92305
## LNATreatmentLNA218:TreatmentPTX                   213.01491  -0.801  0.42415
## LNATreatmentLNA218:SelectionDendrites             212.99157   0.884  0.37783
## TreatmentPTX:SelectionDendrites                   212.99157   1.198  0.23208
## LNATreatmentLNA218:TreatmentPTX:SelectionDendrites 212.99225  -1.256  0.21052
##
## (Intercept)
```

```
## LNATreatmentLNA218                                 **
## TreatmentPTX                                       *
## SelectionDendrites
## LNATreatmentLNA218:TreatmentPTX
## LNATreatmentLNA218:SelectionDendrites
## TreatmentPTX:SelectionDendrites
## LNATreatmentLNA218:TreatmentPTX:SelectionDendrites
## ---
## Signif. codes:  0 '***' 0.001 '**' 0.01 '*' 0.05 '.' 0.1 ' ' 1
##
## Correlation of Fixed Effects:
##                  (Intr) LNATrLNA218 TrtPTX SlctnD LNATrLNA218:TPTX LNATLNA218:S
## LNATrLNA218      -0.403
## TreatmntPTX      -0.399  0.524
## SlctnDndrts      -0.391  0.514       0.509
## LNATrLNA218:TPTX  0.290 -0.721      -0.729 -0.371
## LNATLNA218:S      0.284 -0.707      -0.370 -0.727  0.511
## TrtmnPTX:SD       0.282 -0.370      -0.707 -0.721  0.515            0.524
## LNATLNA218:TPTX: -0.206  0.511       0.516  0.526 -0.708           -0.724
##                  TPTX:S
## LNATrLNA218
## TreatmntPTX
## SlctnDndrts
## LNATrLNA218:TPTX
## LNATLNA218:S
## TrtmnPTX:SD
## LNATLNA218:TPTX: -0.730
```

```
emmeans(mod, trt.vs.ctrl ~ Treatment | LNATreatment | Selection, ref="Mock")
```

```
## $emmeans
## LNATreatment = NC, Selection = Somata:
##  Treatment  emmean    SE   df lower.CL upper.CL
##  Mock      -0.0426 0.157 3.78 -0.48829    0.403
##  PTX        0.2279 0.155 3.60 -0.22160    0.677
##
## LNATreatment = LNA218, Selection = Somata:
##  Treatment  emmean    SE   df lower.CL upper.CL
##  Mock       0.2722 0.154 3.52 -0.17938    0.724
##  PTX        0.4104 0.152 3.34 -0.04681    0.868
##
## LNATreatment = NC, Selection = Dendrites:
##  Treatment  emmean    SE   df lower.CL upper.CL
##  Mock      -0.0307 0.157 3.78 -0.47643    0.415
##  PTX        0.4438 0.155 3.60 -0.00572    0.893
##
## LNATreatment = LNA218, Selection = Dendrites:
##  Treatment  emmean    SE   df lower.CL upper.CL
##  Mock       0.4332 0.154 3.52 -0.01836    0.885
##  PTX        0.4826 0.151 3.28  0.02351    0.942
##
## Degrees-of-freedom method: kenward-roger
## Results are given on the log2 (not the response) scale.
## Confidence level used: 0.95
##
```

```
## $contrasts
## LNATreatment = NC, Selection = Somata:
##  contrast   estimate    SE  df t.ratio p.value
##  PTX - Mock   0.2705 0.120 213   2.246  0.0257
##
## LNATreatment = LNA218, Selection = Somata:
##  contrast   estimate    SE  df t.ratio p.value
##  PTX - Mock   0.1382 0.113 213   1.221  0.2234
##
## LNATreatment = NC, Selection = Dendrites:
##  contrast   estimate    SE  df t.ratio p.value
##  PTX - Mock   0.4746 0.120 213   3.941  0.0001
##
## LNATreatment = LNA218, Selection = Dendrites:
##  contrast   estimate    SE  df t.ratio p.value
##  PTX - Mock   0.0494 0.112 213   0.440  0.6606
##
## Degrees-of-freedom method: kenward-roger
## Results are given on the log2 (not the response) scale.
```

```
emmeans(mod, trt.vs.ctrl ~ LNATreatment | Treatment | Selection, ref="NC")
```

```
## $emmeans
## Treatment = Mock, Selection = Somata:
##  LNATreatment  emmean    SE   df lower.CL upper.CL
##  NC           -0.0426 0.157 3.78 -0.48829    0.403
##  LNA218        0.2722 0.154 3.52 -0.17938    0.724
##
## Treatment = PTX, Selection = Somata:
##  LNATreatment  emmean    SE   df lower.CL upper.CL
##  NC            0.2279 0.155 3.60 -0.22160    0.677
##  LNA218        0.4104 0.152 3.34 -0.04681    0.868
##
## Treatment = Mock, Selection = Dendrites:
##  LNATreatment  emmean    SE   df lower.CL upper.CL
##  NC           -0.0307 0.157 3.78 -0.47643    0.415
##  LNA218        0.4332 0.154 3.52 -0.01836    0.885
##
## Treatment = PTX, Selection = Dendrites:
##  LNATreatment  emmean    SE   df lower.CL upper.CL
##  NC            0.4438 0.155 3.60 -0.00572    0.893
##  LNA218        0.4826 0.151 3.28  0.02351    0.942
##
## Degrees-of-freedom method: kenward-roger
## Results are given on the log2 (not the response) scale.
## Confidence level used: 0.95
##
## $contrasts
## Treatment = Mock, Selection = Somata:
##  contrast    estimate    SE  df t.ratio p.value
##  LNA218 - NC   0.3148 0.119 213   2.635  0.0090
##
## Treatment = PTX, Selection = Somata:
##  contrast    estimate    SE  df t.ratio p.value
##  LNA218 - NC   0.1825 0.115 213   1.593  0.1126
```

```
##
## Treatment = Mock, Selection = Dendrites:
##  contrast    estimate    SE  df t.ratio p.value
##  LNA218 - NC   0.4640 0.119 213   3.884  0.0001
##
## Treatment = PTX, Selection = Dendrites:
##  contrast    estimate    SE  df t.ratio p.value
##  LNA218 - NC   0.0388 0.114 213   0.341  0.7333
##
## Degrees-of-freedom method: kenward-roger
## Results are given on the log2 (not the response) scale.
```

- Gamma2-Cluster Density

```r
mod <- lmer( log2(NormNorm) ~ LNATreatment * Treatment * Selection + (1|Experiment), dat.gamma2)
summary(mod)
```

```
## Linear mixed model fit by REML. t-tests use Satterthwaite's method [
## lmerModLmerTest]
## Formula: log2(NormNorm) ~ LNATreatment * Treatment * Selection + (1 |
##     Experiment)
##    Data: dat.gamma2
##
## REML criterion at convergence: 705.5
##
## Scaled residuals:
##     Min      1Q  Median      3Q     Max
## -6.4476 -0.4534  0.1387  0.6695  1.4543
##
## Random effects:
##  Groups     Name        Variance Std.Dev.
##  Experiment (Intercept) 0.05808  0.241
##  Residual               1.35871  1.166
## Number of obs: 223, groups:  Experiment, 3
##
## Fixed effects:
##                                                     Estimate Std. Error
## (Intercept)                                         -0.20925    0.27192
## LNATreatmentLNA218                                  -0.00547    0.32096
## TreatmentPTX                                        -0.16114    0.32362
## SelectionDendrites                                   0.01394    0.32969
## LNATreatmentLNA218:TreatmentPTX                      0.19148    0.44406
## LNATreatmentLNA218:SelectionDendrites                0.34725    0.45359
## TreatmentPTX:SelectionDendrites                      0.47184    0.45754
## LNATreatmentLNA218:TreatmentPTX:SelectionDendrites  -0.74602    0.62665
##                                                           df t value Pr(>|t|)
## (Intercept)                                         16.21190  -0.770    0.453
## LNATreatmentLNA218                                 213.18410  -0.017    0.986
## TreatmentPTX                                       213.08859  -0.498    0.619
## SelectionDendrites                                 213.01030   0.042    0.966
## LNATreatmentLNA218:TreatmentPTX                    213.12424   0.431    0.667
## LNATreatmentLNA218:SelectionDendrites              213.01030   0.766    0.445
## TreatmentPTX:SelectionDendrites                    213.01030   1.031    0.304
## LNATreatmentLNA218:TreatmentPTX:SelectionDendrites 213.01368  -1.190    0.235
##
```

```
## Correlation of Fixed Effects:
##                   (Intr) LNATrLNA218 TrtPTX SlctnD LNATrLNA218:TPTX LNATLNA218:S
## LNATrLNA218       -0.625
## TreatmntPTX       -0.618  0.524
## SlctnDndrts       -0.606  0.514       0.509
## LNATrLNA218:TPTX  0.449 -0.721      -0.729 -0.371
## LNATLNA218:S      0.441 -0.707      -0.370 -0.727  0.511
## TrtmnPTX:SD       0.437 -0.370      -0.707 -0.721  0.515             0.524
## LNATLNA218:TPTX: -0.319  0.512       0.516  0.526 -0.708            -0.724
##                   TPTX:S
## LNATrLNA218
## TreatmntPTX
## SlctnDndrts
## LNATrLNA218:TPTX
## LNATLNA218:S
## TrtmnPTX:SD
## LNATLNA218:TPTX: -0.730
```

```
emmeans(mod, trt.vs.ctrl ~ Treatment | LNATreatment | Selection, ref="Mock")
```

```
## $emmeans
## LNATreatment = NC, Selection = Somata:
##  Treatment  emmean    SE   df lower.CL upper.CL
##  Mock      -0.2093 0.272 16.1   -0.786    0.367
##  PTX       -0.3704 0.264 14.5   -0.936    0.195
##
## LNATreatment = LNA218, Selection = Somata:
##  Treatment  emmean    SE   df lower.CL upper.CL
##  Mock      -0.2147 0.261 13.8   -0.774    0.345
##  PTX       -0.1844 0.252 12.0   -0.733    0.364
##
## LNATreatment = NC, Selection = Dendrites:
##  Treatment  emmean    SE   df lower.CL upper.CL
##  Mock      -0.1953 0.272 16.1   -0.772    0.381
##  PTX        0.1154 0.264 14.5   -0.450    0.681
##
## LNATreatment = LNA218, Selection = Dendrites:
##  Treatment  emmean    SE   df lower.CL upper.CL
##  Mock       0.1465 0.261 13.8   -0.413    0.706
##  PTX       -0.0974 0.249 11.5   -0.642    0.447
##
## Degrees-of-freedom method: kenward-roger
## Results are given on the log2 (not the response) scale.
## Confidence level used: 0.95
##
## $contrasts
## LNATreatment = NC, Selection = Somata:
##  contrast   estimate    SE  df t.ratio p.value
##  PTX - Mock  -0.1611 0.324 213  -0.498  0.6191
##
## LNATreatment = LNA218, Selection = Somata:
##  contrast   estimate    SE  df t.ratio p.value
##  PTX - Mock   0.0303 0.304 213   0.100  0.9206
##
## LNATreatment = NC, Selection = Dendrites:
```

```
##  contrast   estimate    SE  df t.ratio p.value
##  PTX - Mock   0.3107 0.324 213   0.960  0.3382
##
## LNATreatment = LNA218, Selection = Dendrites:
##  contrast   estimate    SE  df t.ratio p.value
##  PTX - Mock  -0.2438 0.302 213  -0.807  0.4203
##
## Degrees-of-freedom method: kenward-roger
## Results are given on the log2 (not the response) scale.
```

```
emmeans(mod, trt.vs.ctrl ~ LNATreatment | Treatment | Selection, ref="NC")
```

```
## $emmeans
## Treatment = Mock, Selection = Somata:
##  LNATreatment  emmean    SE   df lower.CL upper.CL
##  NC           -0.2093 0.272 16.1   -0.786    0.367
##  LNA218       -0.2147 0.261 13.8   -0.774    0.345
##
## Treatment = PTX, Selection = Somata:
##  LNATreatment  emmean    SE   df lower.CL upper.CL
##  NC           -0.3704 0.264 14.5   -0.936    0.195
##  LNA218       -0.1844 0.252 12.0   -0.733    0.364
##
## Treatment = Mock, Selection = Dendrites:
##  LNATreatment  emmean    SE   df lower.CL upper.CL
##  NC           -0.1953 0.272 16.1   -0.772    0.381
##  LNA218        0.1465 0.261 13.8   -0.413    0.706
##
## Treatment = PTX, Selection = Dendrites:
##  LNATreatment  emmean    SE   df lower.CL upper.CL
##  NC            0.1154 0.264 14.5   -0.450    0.681
##  LNA218       -0.0974 0.249 11.5   -0.642    0.447
##
## Degrees-of-freedom method: kenward-roger
## Results are given on the log2 (not the response) scale.
## Confidence level used: 0.95
##
## $contrasts
## Treatment = Mock, Selection = Somata:
##  contrast    estimate    SE  df t.ratio p.value
##  LNA218 - NC -0.00547 0.321 213  -0.017  0.9864
##
## Treatment = PTX, Selection = Somata:
##  contrast    estimate    SE  df t.ratio p.value
##  LNA218 - NC  0.18601 0.308 214   0.604  0.5465
##
## Treatment = Mock, Selection = Dendrites:
##  contrast    estimate    SE  df t.ratio p.value
##  LNA218 - NC  0.34178 0.321 213   1.064  0.2883
##
## Treatment = PTX, Selection = Dendrites:
##  contrast    estimate    SE  df t.ratio p.value
##  LNA218 - NC -0.21276 0.306 214  -0.696  0.4872
##
## Degrees-of-freedom method: kenward-roger
```

```
## Results are given on the log2 (not the response) scale.
```

**Figure 5D: Nlgn2-vGAT clusters upon PTX-Treatment and miR-218-5p inhibition**

- Nlgn2-cluster size

```r
dat.nlgn2 <- allData$dat.nlgn2

mod <- lmer(NormArea ~ LNATreatment * Treatment * Selection  + (1|Experiment),
            subset(dat.nlgn2, Channel_Name == "GFP" & Second_Channel == "Nlgn2"))

summary(mod)
```

```
## Linear mixed model fit by REML. t-tests use Satterthwaite's method [
## lmerModLmerTest]
## Formula: NormArea ~ LNATreatment * Treatment * Selection + (1 | Experiment)
##    Data: subset(dat.nlgn2, Channel_Name == "GFP" & Second_Channel == "Nlgn2")
##
## REML criterion at convergence: 109.2
##
## Scaled residuals:
##     Min      1Q  Median      3Q     Max
## -2.2073 -0.6107 -0.1328  0.4480  4.5170
##
## Random effects:
##  Groups     Name        Variance Std.Dev.
##  Experiment (Intercept) 0.001532 0.03915
##  Residual               0.082455 0.28715
## Number of obs: 242, groups:  Experiment, 3
##
## Fixed effects:
##                                                  Estimate Std. Error
## (Intercept)                                     1.002e+00  5.425e-02
## LNATreatmentLNA218                              2.999e-01  7.195e-02
## TreatmentPTX                                    2.819e-01  7.266e-02
## SelectionDendrites                             -4.853e-16  6.964e-02
## LNATreatmentLNA218:TreatmentPTX                -3.644e-01  1.047e-01
## LNATreatmentLNA218:SelectionDendrites          -1.420e-01  1.017e-01
## TreatmentPTX:SelectionDendrites                -2.139e-01  1.026e-01
## LNATreatmentLNA218:TreatmentPTX:SelectionDendrites  1.932e-01  1.481e-01
##                                                        df t value Pr(>|t|)
## (Intercept)                                     2.092e+01  18.474 1.98e-14
## LNATreatmentLNA218                              2.321e+02   4.167 4.35e-05
## TreatmentPTX                                    2.325e+02   3.879 0.000136
## SelectionDendrites                              2.319e+02   0.000 1.000000
## LNATreatmentLNA218:TreatmentPTX                 2.319e+02  -3.480 0.000599
## LNATreatmentLNA218:SelectionDendrites           2.319e+02  -1.396 0.164152
## TreatmentPTX:SelectionDendrites                 2.319e+02  -2.083 0.038315
## LNATreatmentLNA218:TreatmentPTX:SelectionDendrites  2.319e+02   1.305 0.193307
##
## (Intercept)                                    ***
## LNATreatmentLNA218                             ***
## TreatmentPTX                                   ***
## SelectionDendrites
```

```
## LNATreatmentLNA218:TreatmentPTX                    ***
## LNATreatmentLNA218:SelectionDendrites
## TreatmentPTX:SelectionDendrites                    *
## LNATreatmentLNA218:TreatmentPTX:SelectionDendrites
## ---
## Signif. codes:  0 '***' 0.001 '**' 0.01 '*' 0.05 '.' 0.1 ' ' 1
##
## Correlation of Fixed Effects:
##                  (Intr) LNATrLNA218 TrtPTX SlctnD LNATrLNA218:TPTX LNATLNA218:S
## LNATrLNA218      -0.621
## TreatmntPTX      -0.617  0.463
## SlctnDndrts      -0.642  0.484       0.479
## LNATrLNA218:TPTX  0.426 -0.687      -0.692 -0.333
## LNATLNA218:S      0.439 -0.707      -0.328 -0.685  0.486
## TrtmnPTX:SD       0.436 -0.328      -0.706 -0.678  0.490            0.465
## LNATLNA218:TPTX: -0.302  0.486       0.490  0.470 -0.707           -0.687
##                  TPTX:S
## LNATrLNA218
## TreatmntPTX
## SlctnDndrts
## LNATrLNA218:TPTX
## LNATLNA218:S
## TrtmnPTX:SD
## LNATLNA218:TPTX: -0.693
```

```
emmeans(mod, trt.vs.ctrl ~ Treatment | LNATreatment | Selection, ref="Mock")
```

```
## $emmeans
## LNATreatment = NC, Selection = Somata:
##  Treatment emmean     SE   df lower.CL upper.CL
##  Mock        1.00 0.0543 22.1    0.890     1.11
##  PTX         1.28 0.0580 28.3    1.165     1.40
##
## LNATreatment = LNA218, Selection = Somata:
##  Treatment emmean     SE   df lower.CL upper.CL
##  Mock        1.30 0.0574 26.6    1.184     1.42
##  PTX         1.22 0.0588 30.0    1.099     1.34
##
## LNATreatment = NC, Selection = Dendrites:
##  Treatment emmean     SE   df lower.CL upper.CL
##  Mock        1.00 0.0543 22.1    0.890     1.11
##  PTX         1.07 0.0580 28.3    0.951     1.19
##
## LNATreatment = LNA218, Selection = Dendrites:
##  Treatment emmean     SE   df lower.CL upper.CL
##  Mock        1.16 0.0574 26.6    1.042     1.28
##  PTX         1.06 0.0588 30.0    0.937     1.18
##
## Degrees-of-freedom method: kenward-roger
## Confidence level used: 0.95
##
## $contrasts
## LNATreatment = NC, Selection = Somata:
##  contrast   estimate     SE  df t.ratio p.value
##  PTX - Mock   0.2819 0.0728 233   3.874  0.0001
```

```
##
## LNATreatment = LNA218, Selection = Somata:
##  contrast   estimate     SE  df t.ratio p.value
##  PTX - Mock  -0.0825 0.0758 233  -1.089  0.2771
##
## LNATreatment = NC, Selection = Dendrites:
##  contrast   estimate     SE  df t.ratio p.value
##  PTX - Mock   0.0680 0.0728 233   0.935  0.3508
##
## LNATreatment = LNA218, Selection = Dendrites:
##  contrast   estimate     SE  df t.ratio p.value
##  PTX - Mock  -0.1032 0.0758 233  -1.362  0.1744
##
## Degrees-of-freedom method: kenward-roger
```

```
emmeans(mod, trt.vs.ctrl ~ LNATreatment | Treatment| Selection, ref="NC")
```

```
## $emmeans
## Treatment = Mock, Selection = Somata:
##  LNATreatment emmean     SE   df lower.CL upper.CL
##  NC             1.00 0.0543 22.1    0.890     1.11
##  LNA218         1.30 0.0574 26.6    1.184     1.42
##
## Treatment = PTX, Selection = Somata:
##  LNATreatment emmean     SE   df lower.CL upper.CL
##  NC             1.28 0.0580 28.3    1.165     1.40
##  LNA218         1.22 0.0588 30.0    1.099     1.34
##
## Treatment = Mock, Selection = Dendrites:
##  LNATreatment emmean     SE   df lower.CL upper.CL
##  NC             1.00 0.0543 22.1    0.890     1.11
##  LNA218         1.16 0.0574 26.6    1.042     1.28
##
## Treatment = PTX, Selection = Dendrites:
##  LNATreatment emmean     SE   df lower.CL upper.CL
##  NC             1.07 0.0580 28.3    0.951     1.19
##  LNA218         1.06 0.0588 30.0    0.937     1.18
##
## Degrees-of-freedom method: kenward-roger
## Confidence level used: 0.95
##
## $contrasts
## Treatment = Mock, Selection = Somata:
##  contrast    estimate     SE  df t.ratio p.value
##  LNA218 - NC   0.2999 0.0720 232   4.165  <.0001
##
## Treatment = PTX, Selection = Somata:
##  contrast    estimate     SE  df t.ratio p.value
##  LNA218 - NC  -0.0645 0.0761 232  -0.848  0.3975
##
## Treatment = Mock, Selection = Dendrites:
##  contrast    estimate     SE  df t.ratio p.value
##  LNA218 - NC   0.1579 0.0720 232   2.193  0.0293
##
## Treatment = PTX, Selection = Dendrites:
```

```
##  contrast    estimate     SE  df t.ratio p.value
##  LNA218 - NC  -0.0133 0.0761 232  -0.175  0.8611
##
## Degrees-of-freedom method: kenward-roger
```

- vGAT-cluster area

```r
mod <- lmer(NormArea ~ LNATreatment * Treatment * Selection  + (1|Experiment),
            subset(dat.nlgn2, Channel_Name == "GFP" & Second_Channel == "vGAT"))
```

```
## boundary (singular) fit: see help('isSingular')
```

```r
summary(mod)
```

```
## Linear mixed model fit by REML. t-tests use Satterthwaite's method [
## lmerModLmerTest]
## Formula: NormArea ~ LNATreatment * Treatment * Selection + (1 | Experiment)
##    Data: subset(dat.nlgn2, Channel_Name == "GFP" & Second_Channel == "vGAT")
##
## REML criterion at convergence: 173.2
##
## Scaled residuals:
##     Min      1Q  Median      3Q     Max
## -2.4560 -0.4659 -0.1208  0.3492  5.6001
##
## Random effects:
##  Groups     Name        Variance Std.Dev.
##  Experiment (Intercept) 0.0000   0.000
##  Residual               0.1096   0.331
## Number of obs: 241, groups:  Experiment, 3
##
## Fixed effects:
##                                                   Estimate Std. Error
## (Intercept)                                      1.000e+00  5.762e-02
## LNATreatmentLNA218                               1.600e-01  8.350e-02
## TreatmentPTX                                     1.920e-01  8.425e-02
## SelectionDendrites                              -1.016e-15  8.089e-02
## LNATreatmentLNA218:TreatmentPTX                 -1.560e-01  1.211e-01
## LNATreatmentLNA218:SelectionDendrites           -1.019e-01  1.177e-01
## TreatmentPTX:SelectionDendrites                 -4.979e-02  1.187e-01
## LNATreatmentLNA218:TreatmentPTX:SelectionDendrites  7.087e-02  1.710e-01
##                                                         df t value Pr(>|t|)
## (Intercept)                                      2.330e+02  17.355   <2e-16
## LNATreatmentLNA218                               2.330e+02   1.916   0.0566
## TreatmentPTX                                     2.330e+02   2.279   0.0236
## SelectionDendrites                               2.330e+02   0.000   1.0000
## LNATreatmentLNA218:TreatmentPTX                  2.330e+02  -1.288   0.1989
## LNATreatmentLNA218:SelectionDendrites            2.330e+02  -0.866   0.3876
## TreatmentPTX:SelectionDendrites                  2.330e+02  -0.419   0.6753
## LNATreatmentLNA218:TreatmentPTX:SelectionDendrites  2.330e+02   0.415   0.6789
##
## (Intercept)                                     ***
## LNATreatmentLNA218                              .
## TreatmentPTX                                    *
## SelectionDendrites
## LNATreatmentLNA218:TreatmentPTX
```

```
## LNATreatmentLNA218:SelectionDendrites
## TreatmentPTX:SelectionDendrites
## LNATreatmentLNA218:TreatmentPTX:SelectionDendrites
## ---
## Signif. codes:  0 '***' 0.001 '**' 0.01 '*' 0.05 '.' 0.1 ' ' 1
##
## Correlation of Fixed Effects:
##                   (Intr) LNATrLNA218 TrtPTX SlctnD LNATrLNA218:TPTX LNATLNA218:S
## LNATrLNA218      -0.690
## TreatmntPTX      -0.684  0.472
## SlctnDndrts      -0.712  0.492       0.487
## LNATrLNA218:TPTX  0.476 -0.690      -0.696 -0.339
## LNATLNA218:S      0.490 -0.710      -0.335 -0.687  0.489
## TrtmnPTX:SD       0.485 -0.335      -0.710 -0.681  0.494             0.468
## LNATLNA218:TPTX: -0.337  0.488       0.493  0.473 -0.708           -0.688
##                   TPTX:S
## LNATrLNA218
## TreatmntPTX
## SlctnDndrts
## LNATrLNA218:TPTX
## LNATLNA218:S
## TrtmnPTX:SD
## LNATLNA218:TPTX: -0.695
## optimizer (nloptwrap) convergence code: 0 (OK)
## boundary (singular) fit: see help('isSingular')
```

```
emmeans(mod, trt.vs.ctrl ~ Treatment | LNATreatment | Selection, ref="Mock")
```

```
## $emmeans
## LNATreatment = NC, Selection = Somata:
##  Treatment emmean     SE   df lower.CL upper.CL
##  Mock        1.00 0.0580 72.3    0.884     1.12
##  PTX         1.19 0.0617 88.8    1.069     1.31
##
## LNATreatment = LNA218, Selection = Somata:
##  Treatment emmean     SE   df lower.CL upper.CL
##  Mock        1.16 0.0611 79.6    1.038     1.28
##  PTX         1.20 0.0626 96.4    1.072     1.32
##
## LNATreatment = NC, Selection = Dendrites:
##  Treatment emmean     SE   df lower.CL upper.CL
##  Mock        1.00 0.0570 70.9    0.886     1.11
##  PTX         1.14 0.0617 88.8    1.020     1.26
##
## LNATreatment = LNA218, Selection = Dendrites:
##  Treatment emmean     SE   df lower.CL upper.CL
##  Mock        1.06 0.0611 79.6    0.937     1.18
##  PTX         1.12 0.0626 96.4    0.991     1.24
##
## Degrees-of-freedom method: kenward-roger
## Confidence level used: 0.95
##
## $contrasts
## LNATreatment = NC, Selection = Somata:
##  contrast   estimate     SE  df t.ratio p.value
```

```
##  PTX - Mock    0.192 0.0846 232   2.269  0.0242
##
## LNATreatment = LNA218, Selection = Somata:
##  contrast   estimate     SE  df t.ratio p.value
##  PTX - Mock    0.036 0.0875 233   0.411  0.6815
##
## LNATreatment = NC, Selection = Dendrites:
##  contrast   estimate     SE  df t.ratio p.value
##  PTX - Mock    0.142 0.0840 232   1.693  0.0919
##
## LNATreatment = LNA218, Selection = Dendrites:
##  contrast   estimate     SE  df t.ratio p.value
##  PTX - Mock    0.057 0.0875 233   0.652  0.5153
##
## Degrees-of-freedom method: kenward-roger
```

```
emmeans(mod, trt.vs.ctrl ~ LNATreatment | Treatment| Selection, ref="NC")
```

```
## $emmeans
## Treatment = Mock, Selection = Somata:
##  LNATreatment emmean     SE   df lower.CL upper.CL
##  NC             1.00 0.0580 72.3    0.884     1.12
##  LNA218         1.16 0.0611 79.6    1.038     1.28
##
## Treatment = PTX, Selection = Somata:
##  LNATreatment emmean     SE   df lower.CL upper.CL
##  NC             1.19 0.0617 88.8    1.069     1.31
##  LNA218         1.20 0.0626 96.4    1.072     1.32
##
## Treatment = Mock, Selection = Dendrites:
##  LNATreatment emmean     SE   df lower.CL upper.CL
##  NC             1.00 0.0570 70.9    0.886     1.11
##  LNA218         1.06 0.0611 79.6    0.937     1.18
##
## Treatment = PTX, Selection = Dendrites:
##  LNATreatment emmean     SE   df lower.CL upper.CL
##  NC             1.14 0.0617 88.8    1.020     1.26
##  LNA218         1.12 0.0626 96.4    0.991     1.24
##
## Degrees-of-freedom method: kenward-roger
## Confidence level used: 0.95
##
## $contrasts
## Treatment = Mock, Selection = Somata:
##  contrast     estimate     SE  df t.ratio p.value
##  LNA218 - NC  0.16000 0.0836 232   1.913  0.0569
##
## Treatment = PTX, Selection = Somata:
##  contrast     estimate     SE  df t.ratio p.value
##  LNA218 - NC  0.00399 0.0878 231   0.045  0.9638
##
## Treatment = Mock, Selection = Dendrites:
##  contrast     estimate     SE  df t.ratio p.value
##  LNA218 - NC  0.05815 0.0830 232   0.700  0.4844
##
```

```
## Treatment = PTX, Selection = Dendrites:
##  contrast    estimate     SE  df t.ratio p.value
##  LNA218 - NC -0.02700 0.0878 231  -0.308  0.7587
##
## Degrees-of-freedom method: kenward-roger
```

- vGAT-Ngln2 co-cluster density

```
dat.nlgn2.density <- allData$dat.nlgn2.density

mod <- lmer(NormNorm ~ LNATreatment * Treatment  + (1|Experiment), dat.nlgn2.density)
summary(mod)
```

```
## Linear mixed model fit by REML. t-tests use Satterthwaite's method [
## lmerModLmerTest]
## Formula: NormNorm ~ LNATreatment * Treatment + (1 | Experiment)
##    Data: dat.nlgn2.density
##
## REML criterion at convergence: 356.1
##
## Scaled residuals:
##     Min      1Q  Median      3Q     Max
## -2.1136 -0.6974 -0.0726  0.6502  4.0645
##
## Random effects:
##  Groups     Name        Variance Std.Dev.
##  Experiment (Intercept) 0.02027  0.1424
##  Residual               0.24737  0.4974
## Number of obs: 237, groups:  Experiment, 3
##
## Fixed effects:
##                                   Estimate Std. Error        df t value Pr(>|t|)
## (Intercept)                        1.01661    0.10297   3.58019   9.873  0.00101
## LNATreatmentLNA218                -0.17831    0.08954 231.11726  -1.991  0.04761
## TreatmentPTX                      -0.12871    0.09009 231.32716  -1.429  0.15445
## LNATreatmentLNA218:TreatmentPTX    0.12191    0.12949 230.98822   0.941  0.34747
##
## (Intercept)                      **
## LNATreatmentLNA218               *
## TreatmentPTX
## LNATreatmentLNA218:TreatmentPTX
## ---
## Signif. codes:  0 '***' 0.001 '**' 0.01 '*' 0.05 '.' 0.1 ' ' 1
##
## Correlation of Fixed Effects:
##             (Intr) LNATrLNA218 TrtPTX
## LNATrLNA218 -0.413
## TreatmntPTX -0.413  0.469
## LNATLNA218:  0.284 -0.690      -0.690
```

```
emmeans(mod, trt.vs.ctrl ~ Treatment | LNATreatment, ref="Mock")
```

```
## $emmeans
## LNATreatment = NC:
##  Treatment emmean    SE   df lower.CL upper.CL
##  Mock       1.017 0.103 3.66    0.720     1.31
```

```
##  PTX        0.888 0.105 3.98    0.595     1.18
##
## LNATreatment = LNA218:
##  Treatment emmean    SE   df lower.CL upper.CL
##  Mock       0.838 0.105 3.94    0.545     1.13
##  PTX        0.832 0.106 4.13    0.541     1.12
##
## Degrees-of-freedom method: kenward-roger
## Confidence level used: 0.95
##
## $contrasts
## LNATreatment = NC:
##  contrast   estimate     SE  df t.ratio p.value
##  PTX - Mock  -0.1287 0.0902 231  -1.427  0.1548
##
## LNATreatment = LNA218:
##  contrast   estimate     SE  df t.ratio p.value
##  PTX - Mock  -0.0068 0.0938 232  -0.072  0.9423
##
## Degrees-of-freedom method: kenward-roger
```

```
emmeans(mod, trt.vs.ctrl ~ LNATreatment | Treatment, ref="NC")
```

```
## $emmeans
## Treatment = Mock:
##  LNATreatment emmean    SE   df lower.CL upper.CL
##  NC            1.017 0.103 3.66    0.720     1.31
##  LNA218        0.838 0.105 3.94    0.545     1.13
##
## Treatment = PTX:
##  LNATreatment emmean    SE   df lower.CL upper.CL
##  NC            0.888 0.105 3.98    0.595     1.18
##  LNA218        0.832 0.106 4.13    0.541     1.12
##
## Degrees-of-freedom method: kenward-roger
## Confidence level used: 0.95
##
## $contrasts
## Treatment = Mock:
##  contrast    estimate     SE  df t.ratio p.value
##  LNA218 - NC  -0.1783 0.0896 231  -1.991  0.0477
##
## Treatment = PTX:
##  contrast    estimate     SE  df t.ratio p.value
##  LNA218 - NC  -0.0564 0.0937 231  -0.602  0.5479
##
## Degrees-of-freedom method: kenward-roger
```

#### Supplement-Figure 6: Gephyrin and vGAT-clustering

- Cluster Area

```
dat.gephyrin <- allData$dat.gephyrin
```

```r
mod <- lmer(NormArea ~ LNATreatment * Treatment* Selection * Second_Channel + (1| Experiment), dat.geph
summary(mod)
```

```
## Linear mixed model fit by REML. t-tests use Satterthwaite's method [
## lmerModLmerTest]
## Formula: NormArea ~ LNATreatment * Treatment * Selection * Second_Channel +
##     (1 | Experiment)
##    Data: dat.gephyrin
##
## REML criterion at convergence: 239.5
##
## Scaled residuals:
##     Min      1Q  Median      3Q     Max
## -3.5777 -0.5636 -0.0507  0.4488  5.6010
##
## Random effects:
##  Groups     Name        Variance Std.Dev.
##  Experiment (Intercept) 0.00944  0.09716
##  Residual               0.06846  0.26165
## Number of obs: 1051, groups:  Experiment, 4
##
## Fixed effects:
##                                                                  Estimate
## (Intercept)                                                     1.000e+00
## LNATreatmentLNA218                                              4.507e-02
## TreatmentPTX                                                    3.878e-01
## SelectionDendrites                                              5.479e-15
## Second_ChannelvGAT                                              7.404e-04
## LNATreatmentLNA218:TreatmentPTX                                -6.440e-02
## LNATreatmentLNA218:SelectionDendrites                          -8.877e-02
## TreatmentPTX:SelectionDendrites                                -2.111e-01
## LNATreatmentLNA218:Second_ChannelvGAT                          -5.473e-02
## TreatmentPTX:Second_ChannelvGAT                                -2.865e-01
## SelectionDendrites:Second_ChannelvGAT                          -9.754e-15
## LNATreatmentLNA218:TreatmentPTX:SelectionDendrites              2.094e-02
## LNATreatmentLNA218:TreatmentPTX:Second_ChannelvGAT              3.005e-02
## LNATreatmentLNA218:SelectionDendrites:Second_ChannelvGAT        1.586e-01
## TreatmentPTX:SelectionDendrites:Second_ChannelvGAT              2.263e-01
## LNATreatmentLNA218:TreatmentPTX:SelectionDendrites:Second_ChannelvGAT -1.253e-01
##                                                                Std. Error
## (Intercept)                                                     5.691e-02
## LNATreatmentLNA218                                              4.592e-02
## TreatmentPTX                                                    4.476e-02
## SelectionDendrites                                              4.190e-02
## Second_ChannelvGAT                                              4.177e-02
## LNATreatmentLNA218:TreatmentPTX                                 6.579e-02
## LNATreatmentLNA218:SelectionDendrites                           6.449e-02
## TreatmentPTX:SelectionDendrites                                 6.310e-02
## LNATreatmentLNA218:Second_ChannelvGAT                           6.424e-02
## TreatmentPTX:Second_ChannelvGAT                                 6.274e-02
## SelectionDendrites:Second_ChannelvGAT                           5.906e-02
## LNATreatmentLNA218:TreatmentPTX:SelectionDendrites              9.251e-02
## LNATreatmentLNA218:TreatmentPTX:Second_ChannelvGAT              9.207e-02
## LNATreatmentLNA218:SelectionDendrites:Second_ChannelvGAT        9.062e-02
```

```
## TreatmentPTX:SelectionDendrites:Second_ChannelvGAT                   8.863e-02
## LNATreatmentLNA218:TreatmentPTX:SelectionDendrites:Second_ChannelvGAT  1.300e-01
##                                                                              df
## (Intercept)                                                          5.330e+00
## LNATreatmentLNA218                                                   1.032e+03
## TreatmentPTX                                                         1.032e+03
## SelectionDendrites                                                   1.032e+03
## Second_ChannelvGAT                                                   1.032e+03
## LNATreatmentLNA218:TreatmentPTX                                      1.032e+03
## LNATreatmentLNA218:SelectionDendrites                                1.032e+03
## TreatmentPTX:SelectionDendrites                                      1.032e+03
## LNATreatmentLNA218:Second_ChannelvGAT                                1.032e+03
## TreatmentPTX:Second_ChannelvGAT                                      1.032e+03
## SelectionDendrites:Second_ChannelvGAT                                1.032e+03
## LNATreatmentLNA218:TreatmentPTX:SelectionDendrites                   1.032e+03
## LNATreatmentLNA218:TreatmentPTX:Second_ChannelvGAT                   1.032e+03
## LNATreatmentLNA218:SelectionDendrites:Second_ChannelvGAT             1.032e+03
## TreatmentPTX:SelectionDendrites:Second_ChannelvGAT                   1.032e+03
## LNATreatmentLNA218:TreatmentPTX:SelectionDendrites:Second_ChannelvGAT  1.032e+03
##                                                                     t value
## (Intercept)                                                          17.575
## LNATreatmentLNA218                                                    0.982
## TreatmentPTX                                                          8.664
## SelectionDendrites                                                    0.000
## Second_ChannelvGAT                                                    0.018
## LNATreatmentLNA218:TreatmentPTX                                      -0.979
## LNATreatmentLNA218:SelectionDendrites                                -1.377
## TreatmentPTX:SelectionDendrites                                      -3.346
## LNATreatmentLNA218:Second_ChannelvGAT                                -0.852
## TreatmentPTX:Second_ChannelvGAT                                      -4.566
## SelectionDendrites:Second_ChannelvGAT                                 0.000
## LNATreatmentLNA218:TreatmentPTX:SelectionDendrites                    0.226
## LNATreatmentLNA218:TreatmentPTX:Second_ChannelvGAT                    0.326
## LNATreatmentLNA218:SelectionDendrites:Second_ChannelvGAT              1.750
## TreatmentPTX:SelectionDendrites:Second_ChannelvGAT                    2.554
## LNATreatmentLNA218:TreatmentPTX:SelectionDendrites:Second_ChannelvGAT  -0.964
##                                                                     Pr(>|t|)
## (Intercept)                                                         6.35e-06
## LNATreatmentLNA218                                                   0.32655
## TreatmentPTX                                                         < 2e-16
## SelectionDendrites                                                   1.00000
## Second_ChannelvGAT                                                   0.98586
## LNATreatmentLNA218:TreatmentPTX                                      0.32785
## LNATreatmentLNA218:SelectionDendrites                                0.16893
## TreatmentPTX:SelectionDendrites                                      0.00085
## LNATreatmentLNA218:Second_ChannelvGAT                                0.39446
## TreatmentPTX:Second_ChannelvGAT                                     5.56e-06
## SelectionDendrites:Second_ChannelvGAT                                1.00000
## LNATreatmentLNA218:TreatmentPTX:SelectionDendrites                   0.82093
## LNATreatmentLNA218:TreatmentPTX:Second_ChannelvGAT                   0.74423
## LNATreatmentLNA218:SelectionDendrites:Second_ChannelvGAT             0.08046
## TreatmentPTX:SelectionDendrites:Second_ChannelvGAT                   0.01080
## LNATreatmentLNA218:TreatmentPTX:SelectionDendrites:Second_ChannelvGAT  0.33522
##
```

```
## (Intercept)                                                   ***
## LNATreatmentLNA218
## TreatmentPTX                                                  ***
## SelectionDendrites
## Second_ChannelvGAT
## LNATreatmentLNA218:TreatmentPTX
## LNATreatmentLNA218:SelectionDendrites
## TreatmentPTX:SelectionDendrites                               ***
## LNATreatmentLNA218:Second_ChannelvGAT
## TreatmentPTX:Second_ChannelvGAT                               ***
## SelectionDendrites:Second_ChannelvGAT
## LNATreatmentLNA218:TreatmentPTX:SelectionDendrites
## LNATreatmentLNA218:TreatmentPTX:Second_ChannelvGAT
## LNATreatmentLNA218:SelectionDendrites:Second_ChannelvGAT      .
## TreatmentPTX:SelectionDendrites:Second_ChannelvGAT            *
## LNATreatmentLNA218:TreatmentPTX:SelectionDendrites:Second_ChannelvGAT
## ---
## Signif. codes:  0 '***' 0.001 '**' 0.01 '*' 0.05 '.' 0.1 ' ' 1
##
## Correlation matrix not shown by default, as p = 16 > 12.
## Use print(x, correlation=TRUE)  or
##     vcov(x)        if you need it
```

```
emmeans(mod, trt.vs.ctrl ~ Treatment | LNATreatment | Selection | Second_Channel, ref="Mock")
```

```
## $emmeans
## LNATreatment = NC, Selection = Somata, Second_Channel = Gephyrin:
##  Treatment emmean     SE   df lower.CL upper.CL
##  Mock       1.000 0.0569 5.35    0.857     1.14
##  PTX        1.388 0.0590 6.19    1.245     1.53
##
## LNATreatment = LNA218, Selection = Somata, Second_Channel = Gephyrin:
##  Treatment emmean     SE   df lower.CL upper.CL
##  Mock       1.045 0.0599 6.55    0.902     1.19
##  PTX        1.369 0.0587 6.05    1.225     1.51
##
## LNATreatment = NC, Selection = Dendrites, Second_Channel = Gephyrin:
##  Treatment emmean     SE   df lower.CL upper.CL
##  Mock       1.000 0.0569 5.35    0.857     1.14
##  PTX        1.177 0.0589 6.12    1.034     1.32
##
## LNATreatment = LNA218, Selection = Dendrites, Second_Channel = Gephyrin:
##  Treatment emmean     SE   df lower.CL upper.CL
##  Mock       0.957 0.0595 6.40    0.813     1.10
##  PTX        1.090 0.0587 6.05    0.946     1.23
##
## LNATreatment = NC, Selection = Somata, Second_Channel = vGAT:
##  Treatment emmean     SE   df lower.CL upper.CL
##  Mock       1.001 0.0568 5.31    0.857     1.14
##  PTX        1.102 0.0586 6.00    0.959     1.25
##
## LNATreatment = LNA218, Selection = Somata, Second_Channel = vGAT:
##  Treatment emmean     SE   df lower.CL upper.CL
##  Mock       0.991 0.0594 6.33    0.848     1.13
```

```
##  PTX        1.058 0.0586 5.99    0.915     1.20
##
## LNATreatment = NC, Selection = Dendrites, Second_Channel = vGAT:
##  Treatment emmean     SE   df lower.CL upper.CL
##  Mock       1.001 0.0568 5.31    0.857     1.14
##  PTX        1.117 0.0586 6.00    0.974     1.26
##
## LNATreatment = LNA218, Selection = Dendrites, Second_Channel = vGAT:
##  Treatment emmean     SE   df lower.CL upper.CL
##  Mock       1.061 0.0594 6.33    0.918     1.20
##  PTX        1.039 0.0586 5.99    0.896     1.18
##
## Degrees-of-freedom method: kenward-roger
## Confidence level used: 0.95
##
## $contrasts
## LNATreatment = NC, Selection = Somata, Second_Channel = Gephyrin:
##  contrast   estimate     SE   df t.ratio p.value
##  PTX - Mock   0.3878 0.0448 1032   8.664  <.0001
##
## LNATreatment = LNA218, Selection = Somata, Second_Channel = Gephyrin:
##  contrast   estimate     SE   df t.ratio p.value
##  PTX - Mock   0.3234 0.0481 1032   6.720  <.0001
##
## LNATreatment = NC, Selection = Dendrites, Second_Channel = Gephyrin:
##  contrast   estimate     SE   df t.ratio p.value
##  PTX - Mock   0.1767 0.0446 1032   3.965  0.0001
##
## LNATreatment = LNA218, Selection = Dendrites, Second_Channel = Gephyrin:
##  contrast   estimate     SE   df t.ratio p.value
##  PTX - Mock   0.1332 0.0477 1032   2.794  0.0053
##
## LNATreatment = NC, Selection = Somata, Second_Channel = vGAT:
##  contrast   estimate     SE   df t.ratio p.value
##  PTX - Mock   0.1013 0.0440 1032   2.300  0.0216
##
## LNATreatment = LNA218, Selection = Somata, Second_Channel = vGAT:
##  contrast   estimate     SE   df t.ratio p.value
##  PTX - Mock   0.0670 0.0473 1032   1.416  0.1570
##
## LNATreatment = NC, Selection = Dendrites, Second_Channel = vGAT:
##  contrast   estimate     SE   df t.ratio p.value
##  PTX - Mock   0.1165 0.0440 1032   2.646  0.0083
##
## LNATreatment = LNA218, Selection = Dendrites, Second_Channel = vGAT:
##  contrast   estimate     SE   df t.ratio p.value
##  PTX - Mock  -0.0222 0.0473 1032  -0.470  0.6388
##
## Degrees-of-freedom method: kenward-roger
```

```
emmeans(mod, trt.vs.ctrl ~ LNATreatment | Treatment| Selection | Second_Channel, ref="NC")
```

```
## $emmeans
## Treatment = Mock, Selection = Somata, Second_Channel = Gephyrin:
##  LNATreatment emmean     SE   df lower.CL upper.CL
```

```
##  NC            1.000 0.0569 5.35    0.857     1.14
##  LNA218        1.045 0.0599 6.55    0.902     1.19
##
## Treatment = PTX, Selection = Somata, Second_Channel = Gephyrin:
##  LNATreatment emmean     SE   df lower.CL upper.CL
##  NC            1.388 0.0590 6.19    1.245     1.53
##  LNA218        1.369 0.0587 6.05    1.225     1.51
##
## Treatment = Mock, Selection = Dendrites, Second_Channel = Gephyrin:
##  LNATreatment emmean     SE   df lower.CL upper.CL
##  NC            1.000 0.0569 5.35    0.857     1.14
##  LNA218        0.957 0.0595 6.40    0.813     1.10
##
## Treatment = PTX, Selection = Dendrites, Second_Channel = Gephyrin:
##  LNATreatment emmean     SE   df lower.CL upper.CL
##  NC            1.177 0.0589 6.12    1.034     1.32
##  LNA218        1.090 0.0587 6.05    0.946     1.23
##
## Treatment = Mock, Selection = Somata, Second_Channel = vGAT:
##  LNATreatment emmean     SE   df lower.CL upper.CL
##  NC            1.001 0.0568 5.31    0.857     1.14
##  LNA218        0.991 0.0594 6.33    0.848     1.13
##
## Treatment = PTX, Selection = Somata, Second_Channel = vGAT:
##  LNATreatment emmean     SE   df lower.CL upper.CL
##  NC            1.102 0.0586 6.00    0.959     1.25
##  LNA218        1.058 0.0586 5.99    0.915     1.20
##
## Treatment = Mock, Selection = Dendrites, Second_Channel = vGAT:
##  LNATreatment emmean     SE   df lower.CL upper.CL
##  NC            1.001 0.0568 5.31    0.857     1.14
##  LNA218        1.061 0.0594 6.33    0.918     1.20
##
## Treatment = PTX, Selection = Dendrites, Second_Channel = vGAT:
##  LNATreatment emmean     SE   df lower.CL upper.CL
##  NC            1.117 0.0586 6.00    0.974     1.26
##  LNA218        1.039 0.0586 5.99    0.896     1.18
##
## Degrees-of-freedom method: kenward-roger
## Confidence level used: 0.95
##
## $contrasts
## Treatment = Mock, Selection = Somata, Second_Channel = Gephyrin:
##  contrast    estimate     SE   df t.ratio p.value
##  LNA218 - NC  0.04507 0.0459 1032   0.981  0.3266
##
## Treatment = PTX, Selection = Somata, Second_Channel = Gephyrin:
##  contrast    estimate     SE   df t.ratio p.value
##  LNA218 - NC -0.01933 0.0470 1032  -0.411  0.6813
##
## Treatment = Mock, Selection = Dendrites, Second_Channel = Gephyrin:
##  contrast    estimate     SE   df t.ratio p.value
##  LNA218 - NC -0.04370 0.0455 1032  -0.961  0.3366
##
```

```
## Treatment = PTX, Selection = Dendrites, Second_Channel = Gephyrin:
##  contrast    estimate     SE   df t.ratio p.value
##  LNA218 - NC -0.08716 0.0469 1032  -1.860  0.0631
##
## Treatment = Mock, Selection = Somata, Second_Channel = vGAT:
##  contrast    estimate     SE   df t.ratio p.value
##  LNA218 - NC -0.00966 0.0451 1032  -0.214  0.8304
##
## Treatment = PTX, Selection = Somata, Second_Channel = vGAT:
##  contrast    estimate     SE   df t.ratio p.value
##  LNA218 - NC -0.04401 0.0463 1032  -0.951  0.3419
##
## Treatment = Mock, Selection = Dendrites, Second_Channel = vGAT:
##  contrast    estimate     SE   df t.ratio p.value
##  LNA218 - NC  0.06013 0.0451 1032   1.333  0.1827
##
## Treatment = PTX, Selection = Dendrites, Second_Channel = vGAT:
##  contrast    estimate     SE   df t.ratio p.value
##  LNA218 - NC -0.07858 0.0463 1032  -1.698  0.0899
##
## Degrees-of-freedom method: kenward-roger
```

- vGAT-Gephyrin co-cluster density

```
dat.gephyrin.density <- allData$dat.gephyrin.density

mod <- lmer(NormNorm ~ LNATreatment * Treatment + (1| Experiment), dat.gephyrin.density)

summary(mod)
```

```
## Linear mixed model fit by REML. t-tests use Satterthwaite's method [
## lmerModLmerTest]
## Formula: NormNorm ~ LNATreatment * Treatment + (1 | Experiment)
##    Data: dat.gephyrin.density
##
## REML criterion at convergence: 594.2
##
## Scaled residuals:
##     Min      1Q  Median      3Q     Max
## -2.7608 -0.6333 -0.1136  0.5595  4.2305
##
## Random effects:
##  Groups     Name        Variance Std.Dev.
##  Experiment (Intercept) 0.04885  0.2210
##  Residual               0.17466  0.4179
## Number of obs: 520, groups:  Experiment, 4
##
## Fixed effects:
##                                Estimate Std. Error        df t value Pr(>|t|)
## (Intercept)                     0.99986    0.11549   3.38371   8.658  0.00201
## LNATreatmentLNA218             -0.05613    0.05178 513.43990  -1.084  0.27891
## TreatmentPTX                    0.13652    0.05058 513.20679   2.699  0.00718
## LNATreatmentLNA218:TreatmentPTX  0.01883    0.07445 513.48581   0.253  0.80040
##
## (Intercept)                    **
```

```
## LNATreatmentLNA218
## TreatmentPTX                     **
## LNATreatmentLNA218:TreatmentPTX
## ---
## Signif. codes:  0 '***' 0.001 '**' 0.01 '*' 0.05 '.' 0.1 ' ' 1
##
## Correlation of Fixed Effects:
##             (Intr) LNATrLNA218 TrtPTX
## LNATrLNA218 -0.189
## TreatmntPTX -0.193  0.437
## LNATLNA218:  0.132 -0.700      -0.686
```

```
emmeans(mod, trt.vs.ctrl ~ Treatment | LNATreatment, ref="Mock")
```

```
## $emmeans
## LNATreatment = NC:
##  Treatment emmean    SE   df lower.CL upper.CL
##  Mock       1.000 0.115 3.39    0.655     1.34
##  PTX        1.136 0.117 3.54    0.795     1.48
##
## LNATreatment = LNA218:
##  Treatment emmean    SE   df lower.CL upper.CL
##  Mock       0.944 0.117 3.60    0.603     1.28
##  PTX        1.099 0.117 3.52    0.757     1.44
##
## Degrees-of-freedom method: kenward-roger
## Confidence level used: 0.95
##
## $contrasts
## LNATreatment = NC:
##  contrast   estimate     SE  df t.ratio p.value
##  PTX - Mock    0.137 0.0506 513   2.699  0.0072
##
## LNATreatment = LNA218:
##  contrast   estimate     SE  df t.ratio p.value
##  PTX - Mock    0.155 0.0542 513   2.866  0.0043
##
## Degrees-of-freedom method: kenward-roger
```

```
emmeans(mod, trt.vs.ctrl ~ LNATreatment | Treatment, ref="NC")
```

```
## $emmeans
## Treatment = Mock:
##  LNATreatment emmean    SE   df lower.CL upper.CL
##  NC            1.000 0.115 3.39    0.655     1.34
##  LNA218        0.944 0.117 3.60    0.603     1.28
##
## Treatment = PTX:
##  LNATreatment emmean    SE   df lower.CL upper.CL
##  NC            1.136 0.117 3.54    0.795     1.48
##  LNA218        1.099 0.117 3.52    0.757     1.44
##
## Degrees-of-freedom method: kenward-roger
## Confidence level used: 0.95
##
```

```
## $contrasts
## Treatment = Mock:
##  contrast    estimate     SE  df t.ratio p.value
##  LNA218 - NC  -0.0561 0.0518 513  -1.084  0.2791
##
## Treatment = PTX:
##  contrast    estimate     SE  df t.ratio p.value
##  LNA218 - NC  -0.0373 0.0532 513  -0.701  0.4834
##
## Degrees-of-freedom method: kenward-roger
```

### Figure 6:

##### Figure 6B: qPCR of miR-218-5p and miR-134-5p & targets in LNA-injected cortical areas used for sleep experiments

```
taqman.sleep <- allData$taqman.sleep

mod <- lmer(-dCq ~ Hemisphere*miR.Expression + (1|Mouse), taqman.sleep)
summary(mod)
```

```
## Linear mixed model fit by REML. t-tests use Satterthwaite's method [
## lmerModLmerTest]
## Formula: -dCq ~ Hemisphere * miR.Expression + (1 | Mouse)
##    Data: taqman.sleep
##
## REML criterion at convergence: 48.2
##
## Scaled residuals:
##     Min      1Q  Median      3Q     Max
## -1.2340 -0.7062  0.0379  0.6231  1.4969
##
## Random effects:
##  Groups   Name        Variance Std.Dev.
##  Mouse    (Intercept) 0.07867  0.2805
##  Residual             0.39398  0.6277
## Number of obs: 24, groups:  Mouse, 6
##
## Fixed effects:
##                                     Estimate Std. Error       df t value
## (Intercept)                          0.03362    0.28067 18.46544   0.120
## HemisphereLNA218                    -1.02705    0.36239 15.00000  -2.834
## miR.ExpressionmiR134                -7.92550    0.36239 15.00000 -21.870
## HemisphereLNA218:miR.ExpressionmiR134  0.60871    0.51250 15.00000   1.188
##                                     Pr(>|t|)
## (Intercept)                           0.9059
## HemisphereLNA218                      0.0126 *
## miR.ExpressionmiR134                8.61e-13 ***
## HemisphereLNA218:miR.ExpressionmiR134   0.2534
## ---
## Signif. codes:  0 '***' 0.001 '**' 0.01 '*' 0.05 '.' 0.1 ' ' 1
##
```

```
## Correlation of Fixed Effects:
##              (Intr) HmLNA218 mR.ER1
## HmsphLNA218 -0.646
## mR.ExprR134 -0.646  0.500
## HLNA218:R.E  0.456 -0.707   -0.707
```

```
emmeans(mod, trt.vs.ctrl ~ Hemisphere | miR.Expression, ref="NC")
```

```
## $emmeans
## miR.Expression = miR218:
##  Hemisphere  emmean    SE   df lower.CL upper.CL
##  NC          0.0336 0.281 18.5   -0.555    0.622
##  LNA218     -0.9934 0.281 18.5   -1.582   -0.405
##
## miR.Expression = miR134:
##  Hemisphere  emmean    SE   df lower.CL upper.CL
##  NC         -7.8919 0.281 18.5   -8.480   -7.303
##  LNA218     -8.3102 0.281 18.5   -8.899   -7.722
##
## Degrees-of-freedom method: kenward-roger
## Confidence level used: 0.95
##
## $contrasts
## miR.Expression = miR218:
##  contrast    estimate    SE df t.ratio p.value
##  LNA218 - NC   -1.027 0.362 15  -2.834  0.0126
##
## miR.Expression = miR134:
##  contrast    estimate    SE df t.ratio p.value
##  LNA218 - NC   -0.418 0.362 15  -1.154  0.2664
##
## Degrees-of-freedom method: kenward-roger
```

Targets Mdga1 and Shank2

```
target.sleep <- allData$target.sleep

mod <- lmer(-dCq ~ Hemisphere*Gene + (1|Mouse), target.sleep)
summary(mod)
```

```
## Linear mixed model fit by REML. t-tests use Satterthwaite's method [
## lmerModLmerTest]
## Formula: -dCq ~ Hemisphere * Gene + (1 | Mouse)
##    Data: target.sleep
##
## REML criterion at convergence: 20.7
##
## Scaled residuals:
##      Min       1Q   Median       3Q      Max
## -1.69188 -0.34782 -0.05931  0.28688  2.21801
##
## Random effects:
##  Groups   Name        Variance Std.Dev.
##  Mouse    (Intercept) 0.2100   0.4583
##  Residual             0.0771   0.2777
## Number of obs: 20, groups:  Mouse, 5
```

```
##
## Fixed effects:
##                              Estimate Std. Error      df t value Pr(>|t|)
## (Intercept)                   -9.1561     0.2396  6.1416 -38.208 1.55e-08 ***
## HemisphereLNA218               0.7879     0.1756 12.0000   4.487 0.000744 ***
## GeneShank2                     3.2289     0.1756 12.0000  18.386 3.72e-10 ***
## HemisphereLNA218:GeneShank2   -0.4151     0.2484 12.0000  -1.671 0.120490
## ---
## Signif. codes:  0 '***' 0.001 '**' 0.01 '*' 0.05 '.' 0.1 ' ' 1
##
## Correlation of Fixed Effects:
##             (Intr) HmLNA218 GnShn2
## HmsphLNA218 -0.366
## GeneShank2  -0.366  0.500
## HLNA218:GS2  0.259 -0.707   -0.707
```

```
emmeans(mod, trt.vs.ctrl ~ Hemisphere|Gene, ref="NC")
```

```
## $emmeans
## Gene = MDGA1:
##  Hemisphere emmean   SE   df lower.CL upper.CL
##  NC          -9.16 0.24 6.14    -9.74    -8.57
##  LNA218      -8.37 0.24 6.14    -8.95    -7.79
##
## Gene = Shank2:
##  Hemisphere emmean   SE   df lower.CL upper.CL
##  NC          -5.93 0.24 6.14    -6.51    -5.34
##  LNA218      -5.55 0.24 6.14    -6.14    -4.97
##
## Degrees-of-freedom method: kenward-roger
## Confidence level used: 0.95
##
## $contrasts
## Gene = MDGA1:
##  contrast    estimate    SE df t.ratio p.value
##  LNA218 - NC    0.788 0.176 12   4.487  0.0007
##
## Gene = Shank2:
##  contrast    estimate    SE df t.ratio p.value
##  LNA218 - NC    0.373 0.176 12   2.123  0.0553
##
## Degrees-of-freedom method: kenward-roger
```

**Figure 6F:**

- Non-REM Sleep

```
binned.melt <- allData$binned.melt
mod <- lmer(power~ Hemisphere*Phase*Frequency_Bands + (1|Mouse),
            subset(binned.melt, Vigilance == "NREM"))
summary(mod)
```

```
## Linear mixed model fit by REML. t-tests use Satterthwaite's method [
## lmerModLmerTest]
## Formula: power ~ Hemisphere * Phase * Frequency_Bands + (1 | Mouse)
```

```
##    Data: subset(binned.melt, Vigilance == "NREM")
##
## REML criterion at convergence: -12.8
##
## Scaled residuals:
##     Min      1Q  Median      3Q     Max
## -2.4787 -0.3785 -0.0010  0.3650  3.7342
##
## Random effects:
##  Groups   Name        Variance Std.Dev.
##  Mouse    (Intercept) 0.01750  0.1323
##  Residual             0.03179  0.1783
## Number of obs: 120, groups:  Mouse, 6
##
## Fixed effects:
##                                              Estimate Std. Error        df
## (Intercept)                                  1.340987   0.090637 29.445562
## HemisphereLNA218                            -0.308433   0.102935 95.000000
## PhaseDark                                    0.098038   0.102935 95.000000
## Frequency_Bandsdelta2                       -0.049625   0.102935 95.000000
## Frequency_Bandstheta                        -0.173116   0.102935 95.000000
## Frequency_Bandsalpha                        -0.254473   0.102935 95.000000
## Frequency_Bandsbeta                         -0.310377   0.102935 95.000000
## HemisphereLNA218:PhaseDark                  -0.031761   0.145571 95.000000
## HemisphereLNA218:Frequency_Bandsdelta2       0.085755   0.145571 95.000000
## HemisphereLNA218:Frequency_Bandstheta        0.229428   0.145571 95.000000
## HemisphereLNA218:Frequency_Bandsalpha        0.297524   0.145571 95.000000
## HemisphereLNA218:Frequency_Bandsbeta         0.320193   0.145571 95.000000
## PhaseDark:Frequency_Bandsdelta2              0.259678   0.145571 95.000000
## PhaseDark:Frequency_Bandstheta              -0.001903   0.145571 95.000000
## PhaseDark:Frequency_Bandsalpha              -0.103285   0.145571 95.000000
## PhaseDark:Frequency_Bandsbeta               -0.084894   0.145571 95.000000
## HemisphereLNA218:PhaseDark:Frequency_Bandsdelta2  0.023837   0.205869 95.000000
## HemisphereLNA218:PhaseDark:Frequency_Bandstheta   0.062624   0.205869 95.000000
## HemisphereLNA218:PhaseDark:Frequency_Bandsalpha   0.026019   0.205869 95.000000
## HemisphereLNA218:PhaseDark:Frequency_Bandsbeta    0.022171   0.205869 95.000000
##                                             t value Pr(>|t|)
## (Intercept)                                  14.795 3.59e-15 ***
## HemisphereLNA218                             -2.996  0.00348 **
## PhaseDark                                     0.952  0.34330
## Frequency_Bandsdelta2                        -0.482  0.63084
## Frequency_Bandstheta                         -1.682  0.09589 .
## Frequency_Bandsalpha                         -2.472  0.01521 *
## Frequency_Bandsbeta                          -3.015  0.00329 **
## HemisphereLNA218:PhaseDark                   -0.218  0.82776
## HemisphereLNA218:Frequency_Bandsdelta2        0.589  0.55720
## HemisphereLNA218:Frequency_Bandstheta         1.576  0.11834
## HemisphereLNA218:Frequency_Bandsalpha         2.044  0.04374 *
## HemisphereLNA218:Frequency_Bandsbeta          2.200  0.03026 *
## PhaseDark:Frequency_Bandsdelta2               1.784  0.07764 .
## PhaseDark:Frequency_Bandstheta               -0.013  0.98960
## PhaseDark:Frequency_Bandsalpha               -0.710  0.47974
## PhaseDark:Frequency_Bandsbeta                -0.583  0.56115
## HemisphereLNA218:PhaseDark:Frequency_Bandsdelta2   0.116  0.90806
```

```
## HemisphereLNA218:PhaseDark:Frequency_Bandstheta    0.304  0.76165
## HemisphereLNA218:PhaseDark:Frequency_Bandsalpha    0.126  0.89969
## HemisphereLNA218:PhaseDark:Frequency_Bandsbeta     0.108  0.91447
## ---
## Signif. codes:  0 '***' 0.001 '**' 0.01 '*' 0.05 '.' 0.1 ' ' 1
##
## Correlation matrix not shown by default, as p = 20 > 12.
## Use print(x, correlation=TRUE)  or
##     vcov(x)        if you need it
```

```
emmeans(mod, trt.vs.ctrl ~ Hemisphere | Frequency_Bands | Phase, ref="NC")
```

```
## $emmeans
## Frequency_Bands = delta1, Phase = Light:
##  Hemisphere emmean     SE   df lower.CL upper.CL
##  NC           1.34 0.0906 29.4    1.156     1.53
##  LNA218       1.03 0.0906 29.4    0.847     1.22
##
## Frequency_Bands = delta2, Phase = Light:
##  Hemisphere emmean     SE   df lower.CL upper.CL
##  NC           1.29 0.0906 29.4    1.106     1.48
##  LNA218       1.07 0.0906 29.4    0.883     1.25
##
## Frequency_Bands = theta, Phase = Light:
##  Hemisphere emmean     SE   df lower.CL upper.CL
##  NC           1.17 0.0906 29.4    0.983     1.35
##  LNA218       1.09 0.0906 29.4    0.904     1.27
##
## Frequency_Bands = alpha, Phase = Light:
##  Hemisphere emmean     SE   df lower.CL upper.CL
##  NC           1.09 0.0906 29.4    0.901     1.27
##  LNA218       1.08 0.0906 29.4    0.890     1.26
##
## Frequency_Bands = beta, Phase = Light:
##  Hemisphere emmean     SE   df lower.CL upper.CL
##  NC           1.03 0.0906 29.4    0.845     1.22
##  LNA218       1.04 0.0906 29.4    0.857     1.23
##
## Frequency_Bands = delta1, Phase = Dark:
##  Hemisphere emmean     SE   df lower.CL upper.CL
##  NC           1.44 0.0906 29.4    1.254     1.62
##  LNA218       1.10 0.0906 29.4    0.914     1.28
##
## Frequency_Bands = delta2, Phase = Dark:
##  Hemisphere emmean     SE   df lower.CL upper.CL
##  NC           1.65 0.0906 29.4    1.464     1.83
##  LNA218       1.42 0.0906 29.4    1.233     1.60
##
## Frequency_Bands = theta, Phase = Dark:
##  Hemisphere emmean     SE   df lower.CL upper.CL
##  NC           1.26 0.0906 29.4    1.079     1.45
##  LNA218       1.22 0.0906 29.4    1.031     1.40
##
## Frequency_Bands = alpha, Phase = Dark:
```

```
##  Hemisphere emmean     SE   df lower.CL upper.CL
##  NC           1.08 0.0906 29.4    0.896     1.27
##  LNA218       1.06 0.0906 29.4    0.879     1.25
##
## Frequency_Bands = beta, Phase = Dark:
##  Hemisphere emmean     SE   df lower.CL upper.CL
##  NC           1.04 0.0906 29.4    0.859     1.23
##  LNA218       1.05 0.0906 29.4    0.861     1.23
##
## Degrees-of-freedom method: kenward-roger
## Confidence level used: 0.95
##
## $contrasts
## Frequency_Bands = delta1, Phase = Light:
##  contrast    estimate    SE df t.ratio p.value
##  LNA218 - NC -0.30843 0.103 95  -2.996  0.0035
##
## Frequency_Bands = delta2, Phase = Light:
##  contrast    estimate    SE df t.ratio p.value
##  LNA218 - NC -0.22268 0.103 95  -2.163  0.0330
##
## Frequency_Bands = theta, Phase = Light:
##  contrast    estimate    SE df t.ratio p.value
##  LNA218 - NC -0.07901 0.103 95  -0.768  0.4447
##
## Frequency_Bands = alpha, Phase = Light:
##  contrast    estimate    SE df t.ratio p.value
##  LNA218 - NC -0.01091 0.103 95  -0.106  0.9158
##
## Frequency_Bands = beta, Phase = Light:
##  contrast    estimate    SE df t.ratio p.value
##  LNA218 - NC  0.01176 0.103 95   0.114  0.9093
##
## Frequency_Bands = delta1, Phase = Dark:
##  contrast    estimate    SE df t.ratio p.value
##  LNA218 - NC -0.34019 0.103 95  -3.305  0.0013
##
## Frequency_Bands = delta2, Phase = Dark:
##  contrast    estimate    SE df t.ratio p.value
##  LNA218 - NC -0.23060 0.103 95  -2.240  0.0274
##
## Frequency_Bands = theta, Phase = Dark:
##  contrast    estimate    SE df t.ratio p.value
##  LNA218 - NC -0.04814 0.103 95  -0.468  0.6411
##
## Frequency_Bands = alpha, Phase = Dark:
##  contrast    estimate    SE df t.ratio p.value
##  LNA218 - NC -0.01665 0.103 95  -0.162  0.8718
##
## Frequency_Bands = beta, Phase = Dark:
##  contrast    estimate    SE df t.ratio p.value
##  LNA218 - NC  0.00217 0.103 95   0.021  0.9832
##
## Degrees-of-freedom method: kenward-roger
```

- REM-Sleep

```
mod <- lmer(power~ Hemisphere*Phase*Frequency_Bands + (1|Mouse),
            subset(binned.melt, Vigilance == "REM"))
summary(mod)
```

```
## Linear mixed model fit by REML. t-tests use Satterthwaite's method [
## lmerModLmerTest]
## Formula: power ~ Hemisphere * Phase * Frequency_Bands + (1 | Mouse)
##    Data: subset(binned.melt, Vigilance == "REM")
##
## REML criterion at convergence: -10.5
##
## Scaled residuals:
##     Min      1Q  Median      3Q     Max
## -2.2274 -0.5507 -0.0328  0.4597  3.1802
##
## Random effects:
##  Groups   Name        Variance Std.Dev.
##  Mouse    (Intercept) 0.01471  0.1213
##  Residual             0.03283  0.1812
## Number of obs: 120, groups:  Mouse, 6
##
## Fixed effects:
##                                                 Estimate Std. Error        df
## (Intercept)                                     1.136299   0.089006 35.477244
## HemisphereLNA218                               -0.179613   0.104604 95.000000
## PhaseDark                                      -0.113800   0.104604 95.000000
## Frequency_Bandsdelta2                          -0.005518   0.104604 95.000000
## Frequency_Bandstheta                            0.012507   0.104604 95.000000
## Frequency_Bandsalpha                           -0.049482   0.104604 95.000000
## Frequency_Bandsbeta                            -0.072686   0.104604 95.000000
## HemisphereLNA218:PhaseDark                      0.038576   0.147933 95.000000
## HemisphereLNA218:Frequency_Bandsdelta2          0.117873   0.147933 95.000000
## HemisphereLNA218:Frequency_Bandstheta           0.112306   0.147933 95.000000
## HemisphereLNA218:Frequency_Bandsalpha           0.168422   0.147933 95.000000
## HemisphereLNA218:Frequency_Bandsbeta            0.189593   0.147933 95.000000
## PhaseDark:Frequency_Bandsdelta2                 0.213126   0.147933 95.000000
## PhaseDark:Frequency_Bandstheta                  0.043998   0.147933 95.000000
## PhaseDark:Frequency_Bandsalpha                  0.290880   0.147933 95.000000
## PhaseDark:Frequency_Bandsbeta                   0.146418   0.147933 95.000000
## HemisphereLNA218:PhaseDark:Frequency_Bandsdelta2  0.002809   0.209209 95.000000
## HemisphereLNA218:PhaseDark:Frequency_Bandstheta   0.010235   0.209209 95.000000
## HemisphereLNA218:PhaseDark:Frequency_Bandsalpha   0.011970   0.209209 95.000000
## HemisphereLNA218:PhaseDark:Frequency_Bandsbeta    0.018166   0.209209 95.000000
##                                                t value Pr(>|t|)
## (Intercept)                                     12.767 7.95e-15 ***
## HemisphereLNA218                                -1.717   0.0892 .
## PhaseDark                                       -1.088   0.2794
## Frequency_Bandsdelta2                           -0.053   0.9580
## Frequency_Bandstheta                             0.120   0.9051
## Frequency_Bandsalpha                            -0.473   0.6373
## Frequency_Bandsbeta                             -0.695   0.4888
## HemisphereLNA218:PhaseDark                       0.261   0.7948
## HemisphereLNA218:Frequency_Bandsdelta2           0.797   0.4276
```

```
## HemisphereLNA218:Frequency_Bandstheta              0.759   0.4496
## HemisphereLNA218:Frequency_Bandsalpha              1.138   0.2578
## HemisphereLNA218:Frequency_Bandsbeta               1.282   0.2031
## PhaseDark:Frequency_Bandsdelta2                    1.441   0.1530
## PhaseDark:Frequency_Bandstheta                     0.297   0.7668
## PhaseDark:Frequency_Bandsalpha                     1.966   0.0522 .
## PhaseDark:Frequency_Bandsbeta                      0.990   0.3248
## HemisphereLNA218:PhaseDark:Frequency_Bandsdelta2   0.013   0.9893
## HemisphereLNA218:PhaseDark:Frequency_Bandstheta    0.049   0.9611
## HemisphereLNA218:PhaseDark:Frequency_Bandsalpha    0.057   0.9545
## HemisphereLNA218:PhaseDark:Frequency_Bandsbeta     0.087   0.9310
## ---
## Signif. codes:  0 '***' 0.001 '**' 0.01 '*' 0.05 '.' 0.1 ' ' 1
##
## Correlation matrix not shown by default, as p = 20 > 12.
## Use print(x, correlation=TRUE)  or
##     vcov(x)        if you need it
```

```
emmeans(mod, trt.vs.ctrl ~ Hemisphere | Frequency_Bands | Phase, ref="NC")
```

```
## $emmeans
## Frequency_Bands = delta1, Phase = Light:
##  Hemisphere emmean    SE   df lower.CL upper.CL
##  NC          1.136 0.089 35.5    0.956     1.32
##  LNA218      0.957 0.089 35.5    0.776     1.14
##
## Frequency_Bands = delta2, Phase = Light:
##  Hemisphere emmean    SE   df lower.CL upper.CL
##  NC          1.131 0.089 35.5    0.950     1.31
##  LNA218      1.069 0.089 35.5    0.888     1.25
##
## Frequency_Bands = theta, Phase = Light:
##  Hemisphere emmean    SE   df lower.CL upper.CL
##  NC          1.149 0.089 35.5    0.968     1.33
##  LNA218      1.081 0.089 35.5    0.901     1.26
##
## Frequency_Bands = alpha, Phase = Light:
##  Hemisphere emmean    SE   df lower.CL upper.CL
##  NC          1.087 0.089 35.5    0.906     1.27
##  LNA218      1.076 0.089 35.5    0.895     1.26
##
## Frequency_Bands = beta, Phase = Light:
##  Hemisphere emmean    SE   df lower.CL upper.CL
##  NC          1.064 0.089 35.5    0.883     1.24
##  LNA218      1.074 0.089 35.5    0.893     1.25
##
## Frequency_Bands = delta1, Phase = Dark:
##  Hemisphere emmean    SE   df lower.CL upper.CL
##  NC          1.022 0.089 35.5    0.842     1.20
##  LNA218      0.881 0.089 35.5    0.701     1.06
##
## Frequency_Bands = delta2, Phase = Dark:
##  Hemisphere emmean    SE   df lower.CL upper.CL
##  NC          1.230 0.089 35.5    1.050     1.41
```

```
##  LNA218      1.210 0.089 35.5    1.029     1.39
##
## Frequency_Bands = theta, Phase = Dark:
##  Hemisphere emmean    SE   df lower.CL upper.CL
##  NC          1.079 0.089 35.5    0.898     1.26
##  LNA218      1.061 0.089 35.5    0.880     1.24
##
## Frequency_Bands = alpha, Phase = Dark:
##  Hemisphere emmean    SE   df lower.CL upper.CL
##  NC          1.264 0.089 35.5    1.083     1.44
##  LNA218      1.303 0.089 35.5    1.123     1.48
##
## Frequency_Bands = beta, Phase = Dark:
##  Hemisphere emmean    SE   df lower.CL upper.CL
##  NC          1.096 0.089 35.5    0.916     1.28
##  LNA218      1.163 0.089 35.5    0.982     1.34
##
## Degrees-of-freedom method: kenward-roger
## Confidence level used: 0.95
##
## $contrasts
## Frequency_Bands = delta1, Phase = Light:
##  contrast    estimate    SE df t.ratio p.value
##  LNA218 - NC -0.17961 0.105 95  -1.717  0.0892
##
## Frequency_Bands = delta2, Phase = Light:
##  contrast    estimate    SE df t.ratio p.value
##  LNA218 - NC -0.06174 0.105 95  -0.590  0.5564
##
## Frequency_Bands = theta, Phase = Light:
##  contrast    estimate    SE df t.ratio p.value
##  LNA218 - NC -0.06731 0.105 95  -0.643  0.5215
##
## Frequency_Bands = alpha, Phase = Light:
##  contrast    estimate    SE df t.ratio p.value
##  LNA218 - NC -0.01119 0.105 95  -0.107  0.9150
##
## Frequency_Bands = beta, Phase = Light:
##  contrast    estimate    SE df t.ratio p.value
##  LNA218 - NC  0.00998 0.105 95   0.095  0.9242
##
## Frequency_Bands = delta1, Phase = Dark:
##  contrast    estimate    SE df t.ratio p.value
##  LNA218 - NC -0.14104 0.105 95  -1.348  0.1808
##
## Frequency_Bands = delta2, Phase = Dark:
##  contrast    estimate    SE df t.ratio p.value
##  LNA218 - NC -0.02036 0.105 95  -0.195  0.8461
##
## Frequency_Bands = theta, Phase = Dark:
##  contrast    estimate    SE df t.ratio p.value
##  LNA218 - NC -0.01850 0.105 95  -0.177  0.8600
##
## Frequency_Bands = alpha, Phase = Dark:
```

```
##  contrast    estimate    SE df t.ratio p.value
##  LNA218 - NC  0.03935 0.105 95   0.376  0.7076
##
## Frequency_Bands = beta, Phase = Dark:
##  contrast    estimate    SE df t.ratio p.value
##  LNA218 - NC  0.06672 0.105 95   0.638  0.5251
##
## Degrees-of-freedom method: kenward-roger
```

- Wake

```
mod <- lmer(power~ Hemisphere*Phase*Frequency_Bands + (1|Mouse),
            subset(binned.melt, Vigilance == "Wake"))
summary(mod)
```

```
## Linear mixed model fit by REML. t-tests use Satterthwaite's method [
## lmerModLmerTest]
## Formula: power ~ Hemisphere * Phase * Frequency_Bands + (1 | Mouse)
##    Data: subset(binned.melt, Vigilance == "Wake")
##
## REML criterion at convergence: 24.4
##
## Scaled residuals:
##     Min      1Q  Median      3Q     Max
## -2.8278 -0.4421 -0.0025  0.3243  3.3743
##
## Random effects:
##  Groups   Name        Variance Std.Dev.
##  Mouse    (Intercept) 0.004084 0.0639
##  Residual             0.049730 0.2230
## Number of obs: 120, groups:  Mouse, 6
##
## Fixed effects:
##                                                Estimate Std. Error       df
## (Intercept)                                     1.28014    0.09470 90.13810
## HemisphereLNA218                               -0.39661    0.12875 95.00000
## PhaseDark                                       0.21230    0.12875 95.00000
## Frequency_Bandsdelta2                          -0.07213    0.12875 95.00000
## Frequency_Bandstheta                           -0.14214    0.12875 95.00000
## Frequency_Bandsalpha                           -0.27233    0.12875 95.00000
## Frequency_Bandsbeta                            -0.27954    0.12875 95.00000
## HemisphereLNA218:PhaseDark                      0.16997    0.18208 95.00000
## HemisphereLNA218:Frequency_Bandsdelta2          0.17806    0.18208 95.00000
## HemisphereLNA218:Frequency_Bandstheta           0.33840    0.18208 95.00000
## HemisphereLNA218:Frequency_Bandsalpha           0.38674    0.18208 95.00000
## HemisphereLNA218:Frequency_Bandsbeta            0.39831    0.18208 95.00000
## PhaseDark:Frequency_Bandsdelta2                -0.50652    0.18208 95.00000
## PhaseDark:Frequency_Bandstheta                 -0.30279    0.18208 95.00000
## PhaseDark:Frequency_Bandsalpha                  0.17870    0.18208 95.00000
## PhaseDark:Frequency_Bandsbeta                  -0.23887    0.18208 95.00000
## HemisphereLNA218:PhaseDark:Frequency_Bandsdelta2  0.01678    0.25750 95.00000
## HemisphereLNA218:PhaseDark:Frequency_Bandstheta  -0.13754    0.25750 95.00000
## HemisphereLNA218:PhaseDark:Frequency_Bandsalpha  -0.14753    0.25750 95.00000
## HemisphereLNA218:PhaseDark:Frequency_Bandsbeta   -0.15136    0.25750 95.00000
##                                                t value Pr(>|t|)
```

```
## (Intercept)                                         13.517  < 2e-16 ***
## HemisphereLNA218                                    -3.080  0.00270 **
## PhaseDark                                            1.649  0.10247
## Frequency_Bandsdelta2                               -0.560  0.57665
## Frequency_Bandstheta                                -1.104  0.27240
## Frequency_Bandsalpha                                -2.115  0.03703 *
## Frequency_Bandsbeta                                 -2.171  0.03241 *
## HemisphereLNA218:PhaseDark                           0.933  0.35294
## HemisphereLNA218:Frequency_Bandsdelta2               0.978  0.33060
## HemisphereLNA218:Frequency_Bandstheta                1.859  0.06619 .
## HemisphereLNA218:Frequency_Bandsalpha                2.124  0.03627 *
## HemisphereLNA218:Frequency_Bandsbeta                 2.188  0.03116 *
## PhaseDark:Frequency_Bandsdelta2                     -2.782  0.00652 **
## PhaseDark:Frequency_Bandstheta                      -1.663  0.09962 .
## PhaseDark:Frequency_Bandsalpha                       0.981  0.32888
## PhaseDark:Frequency_Bandsbeta                       -1.312  0.19272
## HemisphereLNA218:PhaseDark:Frequency_Bandsdelta2     0.065  0.94819
## HemisphereLNA218:PhaseDark:Frequency_Bandstheta     -0.534  0.59451
## HemisphereLNA218:PhaseDark:Frequency_Bandsalpha     -0.573  0.56806
## HemisphereLNA218:PhaseDark:Frequency_Bandsbeta      -0.588  0.55806
## ---
## Signif. codes:  0 '***' 0.001 '**' 0.01 '*' 0.05 '.' 0.1 ' ' 1
##
## Correlation matrix not shown by default, as p = 20 > 12.
## Use print(x, correlation=TRUE)  or
##     vcov(x)        if you need it
```

```
emmeans(mod, trt.vs.ctrl ~ Hemisphere | Frequency_Bands | Phase, ref="NC")
```

```
## $emmeans
## Frequency_Bands = delta1, Phase = Light:
##  Hemisphere emmean     SE   df lower.CL upper.CL
##  NC          1.280 0.0947 90.1    1.092     1.47
##  LNA218      0.884 0.0947 90.1    0.695     1.07
##
## Frequency_Bands = delta2, Phase = Light:
##  Hemisphere emmean     SE   df lower.CL upper.CL
##  NC          1.208 0.0947 90.1    1.020     1.40
##  LNA218      0.989 0.0947 90.1    0.801     1.18
##
## Frequency_Bands = theta, Phase = Light:
##  Hemisphere emmean     SE   df lower.CL upper.CL
##  NC          1.138 0.0947 90.1    0.950     1.33
##  LNA218      1.080 0.0947 90.1    0.892     1.27
##
## Frequency_Bands = alpha, Phase = Light:
##  Hemisphere emmean     SE   df lower.CL upper.CL
##  NC          1.008 0.0947 90.1    0.820     1.20
##  LNA218      0.998 0.0947 90.1    0.810     1.19
##
## Frequency_Bands = beta, Phase = Light:
##  Hemisphere emmean     SE   df lower.CL upper.CL
##  NC          1.001 0.0947 90.1    0.812     1.19
##  LNA218      1.002 0.0947 90.1    0.814     1.19
```

```
##
## Frequency_Bands = delta1, Phase = Dark:
##  Hemisphere emmean     SE   df lower.CL upper.CL
##  NC          1.492 0.0947 90.1    1.304     1.68
##  LNA218      1.266 0.0947 90.1    1.078     1.45
##
## Frequency_Bands = delta2, Phase = Dark:
##  Hemisphere emmean     SE   df lower.CL upper.CL
##  NC          0.914 0.0947 90.1    0.726     1.10
##  LNA218      0.882 0.0947 90.1    0.694     1.07
##
## Frequency_Bands = theta, Phase = Dark:
##  Hemisphere emmean     SE   df lower.CL upper.CL
##  NC          1.048 0.0947 90.1    0.859     1.24
##  LNA218      1.022 0.0947 90.1    0.834     1.21
##
## Frequency_Bands = alpha, Phase = Dark:
##  Hemisphere emmean     SE   df lower.CL upper.CL
##  NC          1.399 0.0947 90.1    1.211     1.59
##  LNA218      1.411 0.0947 90.1    1.223     1.60
##
## Frequency_Bands = beta, Phase = Dark:
##  Hemisphere emmean     SE   df lower.CL upper.CL
##  NC          0.974 0.0947 90.1    0.786     1.16
##  LNA218      0.994 0.0947 90.1    0.806     1.18
##
## Degrees-of-freedom method: kenward-roger
## Confidence level used: 0.95
##
## $contrasts
## Frequency_Bands = delta1, Phase = Light:
##  contrast    estimate    SE df t.ratio p.value
##  LNA218 - NC -0.39661 0.129 95  -3.080  0.0027
##
## Frequency_Bands = delta2, Phase = Light:
##  contrast    estimate    SE df t.ratio p.value
##  LNA218 - NC -0.21855 0.129 95  -1.697  0.0929
##
## Frequency_Bands = theta, Phase = Light:
##  contrast    estimate    SE df t.ratio p.value
##  LNA218 - NC -0.05821 0.129 95  -0.452  0.6522
##
## Frequency_Bands = alpha, Phase = Light:
##  contrast    estimate    SE df t.ratio p.value
##  LNA218 - NC -0.00987 0.129 95  -0.077  0.9390
##
## Frequency_Bands = beta, Phase = Light:
##  contrast    estimate    SE df t.ratio p.value
##  LNA218 - NC  0.00169 0.129 95   0.013  0.9895
##
## Frequency_Bands = delta1, Phase = Dark:
##  contrast    estimate    SE df t.ratio p.value
##  LNA218 - NC -0.22664 0.129 95  -1.760  0.0816
##
```

```
## Frequency_Bands = delta2, Phase = Dark:
##  contrast    estimate    SE df t.ratio p.value
##  LNA218 - NC -0.03181 0.129 95  -0.247  0.8054
##
## Frequency_Bands = theta, Phase = Dark:
##  contrast    estimate    SE df t.ratio p.value
##  LNA218 - NC -0.02578 0.129 95  -0.200  0.8417
##
## Frequency_Bands = alpha, Phase = Dark:
##  contrast    estimate    SE df t.ratio p.value
##  LNA218 - NC  0.01257 0.129 95   0.098  0.9224
##
## Frequency_Bands = beta, Phase = Dark:
##  contrast    estimate    SE df t.ratio p.value
##  LNA218 - NC  0.02030 0.129 95   0.158  0.8750
##
## Degrees-of-freedom method: kenward-roger
```

**Figure 6G: Timecourse**

```
swa <- allData$swa

mod <- lmer(corrected ~ Hemisphere*as.factor(hours) + (1|Mouse), swa)
summary(mod)
```

```
## Linear mixed model fit by REML. t-tests use Satterthwaite's method [
## lmerModLmerTest]
## Formula: corrected ~ Hemisphere * as.factor(hours) + (1 | Mouse)
##    Data: swa
##
## REML criterion at convergence: -119.7
##
## Scaled residuals:
##     Min      1Q  Median      3Q     Max
## -2.8424 -0.5563 -0.0062  0.4752  4.0146
##
## Random effects:
##  Groups   Name        Variance Std.Dev.
##  Mouse    (Intercept) 0.009035 0.09505
##  Residual             0.030316 0.17412
## Number of obs: 654, groups:  Mouse, 6
##
## Fixed effects:
##                                      Estimate Std. Error         df t value
## (Intercept)                         1.377e+00  8.706e-02  1.035e+02  15.818
## HemisphereLNA218                   -7.346e-02  1.101e-01  5.090e+02  -0.667
## as.factor(hours)2                  -2.965e-01  1.055e-01  5.090e+02  -2.811
## as.factor(hours)3                  -3.720e-01  1.055e-01  5.090e+02  -3.527
## as.factor(hours)4                  -4.474e-01  1.102e-01  5.091e+02  -4.059
## as.factor(hours)5                  -4.540e-01  1.102e-01  5.091e+02  -4.119
## as.factor(hours)6                  -4.983e-01  1.055e-01  5.090e+02  -4.724
## as.factor(hours)7                  -4.697e-01  1.102e-01  5.091e+02  -4.261
## as.factor(hours)8                  -5.410e-01  1.055e-01  5.090e+02  -5.129
```

```
## as.factor(hours)9                  -5.113e-01  1.102e-01  5.091e+02  -4.638
## as.factor(hours)10                 -5.026e-01  1.102e-01  5.091e+02  -4.559
## as.factor(hours)11                 -3.914e-01  1.055e-01  5.090e+02  -3.710
## as.factor(hours)12                 -3.480e-01  1.055e-01  5.090e+02  -3.299
## as.factor(hours)14                 -5.258e-02  1.913e-01  5.092e+02  -0.275
## as.factor(hours)15                 -9.759e-02  1.169e-01  5.090e+02  -0.835
## as.factor(hours)16                 -1.508e-01  1.275e-01  5.092e+02  -1.183
## as.factor(hours)17                 -2.315e-01  1.460e-01  5.091e+02  -1.586
## as.factor(hours)18                 -3.229e-01  1.170e-01  5.091e+02  -2.760
## as.factor(hours)19                 -2.282e-01  1.275e-01  5.092e+02  -1.790
## as.factor(hours)20                 -2.196e-01  1.055e-01  5.090e+02  -2.081
## as.factor(hours)21                 -3.576e-01  1.055e-01  5.090e+02  -3.390
## as.factor(hours)22                 -2.385e-01  1.273e-01  5.091e+02  -1.873
## as.factor(hours)23                 -3.276e-01  1.273e-01  5.091e+02  -2.573
## as.factor(hours)25                 -8.270e-02  1.055e-01  5.090e+02  -0.784
## as.factor(hours)26                 -2.989e-01  1.055e-01  5.090e+02  -2.833
## as.factor(hours)27                 -3.373e-01  1.055e-01  5.090e+02  -3.198
## as.factor(hours)28                 -5.294e-01  1.055e-01  5.090e+02  -5.019
## as.factor(hours)29                 -4.506e-01  1.055e-01  5.090e+02  -4.272
## as.factor(hours)30                 -5.500e-01  1.055e-01  5.090e+02  -5.214
## as.factor(hours)31                 -5.423e-01  1.055e-01  5.090e+02  -5.141
## as.factor(hours)32                 -5.648e-01  1.055e-01  5.090e+02  -5.354
## as.factor(hours)33                 -5.436e-01  1.055e-01  5.090e+02  -5.154
## as.factor(hours)34                 -5.610e-01  1.055e-01  5.090e+02  -5.318
## as.factor(hours)35                 -5.494e-01  1.055e-01  5.090e+02  -5.209
## as.factor(hours)36                 -3.908e-01  1.102e-01  5.091e+02  -3.546
## as.factor(hours)37                 -3.404e-01  1.913e-01  5.092e+02  -1.779
## as.factor(hours)38                 -1.379e-01  1.460e-01  5.091e+02  -0.945
## as.factor(hours)39                 -2.960e-02  1.273e-01  5.091e+02  -0.232
## as.factor(hours)40                 -1.390e-01  1.273e-01  5.091e+02  -1.092
## as.factor(hours)41                 -2.335e-01  1.273e-01  5.091e+02  -1.834
## as.factor(hours)42                 -1.354e-01  1.170e-01  5.091e+02  -1.157
## as.factor(hours)43                 -3.237e-01  1.170e-01  5.091e+02  -2.767
## as.factor(hours)44                 -2.534e-01  1.055e-01  5.090e+02  -2.402
## as.factor(hours)45                 -3.402e-01  1.055e-01  5.090e+02  -3.225
## as.factor(hours)46                 -3.220e-01  1.055e-01  5.090e+02  -3.053
## as.factor(hours)47                 -1.077e-01  1.460e-01  5.091e+02  -0.738
## as.factor(hours)48                 -4.063e-01  1.913e-01  5.092e+02  -2.123
## as.factor(hours)49                  3.701e-01  1.102e-01  5.091e+02   3.357
## as.factor(hours)50                 -8.756e-02  1.102e-01  5.091e+02  -0.794
## as.factor(hours)51                 -2.956e-01  1.055e-01  5.090e+02  -2.802
## as.factor(hours)52                 -3.171e-01  1.055e-01  5.090e+02  -3.006
## as.factor(hours)53                 -4.154e-01  1.055e-01  5.090e+02  -3.938
## as.factor(hours)54                 -3.967e-01  1.055e-01  5.090e+02  -3.760
## as.factor(hours)55                 -5.082e-01  1.055e-01  5.090e+02  -4.818
## as.factor(hours)56                 -4.797e-01  1.055e-01  5.090e+02  -4.548
## as.factor(hours)57                 -5.805e-01  1.055e-01  5.090e+02  -5.504
## as.factor(hours)58                 -5.475e-01  1.055e-01  5.090e+02  -5.190
## as.factor(hours)59                 -4.979e-01  1.055e-01  5.090e+02  -4.721
## as.factor(hours)60                 -3.627e-01  1.055e-01  5.090e+02  -3.439
## as.factor(hours)61                 -2.405e-01  1.913e-01  5.092e+02  -1.257
## as.factor(hours)62                  1.105e-01  1.913e-01  5.092e+02   0.578
## as.factor(hours)63                 -1.463e-01  1.273e-01  5.091e+02  -1.149
## as.factor(hours)64                 -1.034e-02  1.169e-01  5.090e+02  -0.088
```

```
## as.factor(hours)65                  -3.617e-02  1.169e-01  5.090e+02  -0.309
## as.factor(hours)66                   1.135e-01  1.275e-01  5.092e+02   0.890
## as.factor(hours)67                  -1.231e-01  1.055e-01  5.090e+02  -1.167
## as.factor(hours)68                  -1.827e-01  1.055e-01  5.090e+02  -1.732
## as.factor(hours)69                  -2.258e-01  1.055e-01  5.090e+02  -2.141
## as.factor(hours)70                  -3.482e-01  1.055e-01  5.090e+02  -3.301
## as.factor(hours)71                  -2.296e-02  1.460e-01  5.091e+02  -0.157
## as.factor(hours)72                   2.095e-01  1.460e-01  5.091e+02   1.436
## HemisphereLNA218:as.factor(hours)2   1.331e-01  1.491e-01  5.090e+02   0.893
## HemisphereLNA218:as.factor(hours)3   9.969e-02  1.491e-01  5.090e+02   0.669
## HemisphereLNA218:as.factor(hours)4   9.226e-02  1.557e-01  5.090e+02   0.592
## HemisphereLNA218:as.factor(hours)5   8.531e-02  1.557e-01  5.090e+02   0.548
## HemisphereLNA218:as.factor(hours)6   1.378e-01  1.491e-01  5.090e+02   0.924
## HemisphereLNA218:as.factor(hours)7   6.790e-02  1.557e-01  5.090e+02   0.436
## HemisphereLNA218:as.factor(hours)8   6.840e-02  1.491e-01  5.090e+02   0.459
## HemisphereLNA218:as.factor(hours)9   5.038e-02  1.557e-01  5.090e+02   0.324
## HemisphereLNA218:as.factor(hours)10  5.123e-02  1.557e-01  5.090e+02   0.329
## HemisphereLNA218:as.factor(hours)11  4.755e-02  1.491e-01  5.090e+02   0.319
## HemisphereLNA218:as.factor(hours)12  2.385e-02  1.491e-01  5.090e+02   0.160
## HemisphereLNA218:as.factor(hours)14  1.320e-01  2.697e-01  5.090e+02   0.489
## HemisphereLNA218:as.factor(hours)15  1.519e-01  1.652e-01  5.090e+02   0.919
## HemisphereLNA218:as.factor(hours)16  1.682e-01  1.798e-01  5.090e+02   0.936
## HemisphereLNA218:as.factor(hours)17  1.535e-01  2.060e-01  5.090e+02   0.745
## HemisphereLNA218:as.factor(hours)18  1.830e-01  1.652e-01  5.090e+02   1.108
## HemisphereLNA218:as.factor(hours)19  7.216e-02  1.798e-01  5.090e+02   0.401
## HemisphereLNA218:as.factor(hours)20  8.126e-02  1.491e-01  5.090e+02   0.545
## HemisphereLNA218:as.factor(hours)21  1.227e-01  1.491e-01  5.090e+02   0.823
## HemisphereLNA218:as.factor(hours)22  1.177e-01  1.798e-01  5.090e+02   0.655
## HemisphereLNA218:as.factor(hours)23  8.642e-02  1.798e-01  5.090e+02   0.481
## HemisphereLNA218:as.factor(hours)25  9.665e-03  1.491e-01  5.090e+02   0.065
## HemisphereLNA218:as.factor(hours)26  4.058e-05  1.491e-01  5.090e+02   0.000
## HemisphereLNA218:as.factor(hours)27 -5.864e-03  1.491e-01  5.090e+02  -0.039
## HemisphereLNA218:as.factor(hours)28  5.770e-02  1.491e-01  5.090e+02   0.387
## HemisphereLNA218:as.factor(hours)29  5.686e-03  1.491e-01  5.090e+02   0.038
## HemisphereLNA218:as.factor(hours)30  1.818e-02  1.491e-01  5.090e+02   0.122
## HemisphereLNA218:as.factor(hours)31  3.538e-02  1.491e-01  5.090e+02   0.237
## HemisphereLNA218:as.factor(hours)32  2.300e-02  1.491e-01  5.090e+02   0.154
## HemisphereLNA218:as.factor(hours)33  2.265e-03  1.491e-01  5.090e+02   0.015
## HemisphereLNA218:as.factor(hours)34  1.165e-02  1.491e-01  5.090e+02   0.078
## HemisphereLNA218:as.factor(hours)35  1.458e-02  1.491e-01  5.090e+02   0.098
## HemisphereLNA218:as.factor(hours)36  1.313e-02  1.557e-01  5.090e+02   0.084
## HemisphereLNA218:as.factor(hours)37 -1.506e-01  2.697e-01  5.090e+02  -0.558
## HemisphereLNA218:as.factor(hours)38  2.938e-02  2.060e-01  5.090e+02   0.143
## HemisphereLNA218:as.factor(hours)39 -1.987e-02  1.798e-01  5.090e+02  -0.110
## HemisphereLNA218:as.factor(hours)40  1.240e-01  1.798e-01  5.090e+02   0.690
## HemisphereLNA218:as.factor(hours)41  2.686e-02  1.798e-01  5.090e+02   0.149
## HemisphereLNA218:as.factor(hours)42 -4.931e-02  1.652e-01  5.090e+02  -0.299
## HemisphereLNA218:as.factor(hours)43  4.896e-02  1.652e-01  5.090e+02   0.296
## HemisphereLNA218:as.factor(hours)44  1.257e-02  1.491e-01  5.090e+02   0.084
## HemisphereLNA218:as.factor(hours)45  2.499e-02  1.491e-01  5.090e+02   0.168
## HemisphereLNA218:as.factor(hours)46 -3.549e-04  1.491e-01  5.090e+02  -0.002
## HemisphereLNA218:as.factor(hours)47  1.746e-01  2.060e-01  5.090e+02   0.848
## HemisphereLNA218:as.factor(hours)48  4.846e-02  2.697e-01  5.090e+02   0.180
## HemisphereLNA218:as.factor(hours)49 -3.850e-01  1.557e-01  5.090e+02  -2.472
```

```
## HemisphereLNA218:as.factor(hours)50 -2.488e-01  1.557e-01  5.090e+02  -1.598
## HemisphereLNA218:as.factor(hours)51 -1.689e-01  1.491e-01  5.090e+02  -1.133
## HemisphereLNA218:as.factor(hours)52 -1.963e-01  1.491e-01  5.090e+02  -1.317
## HemisphereLNA218:as.factor(hours)53 -1.735e-01  1.491e-01  5.090e+02  -1.163
## HemisphereLNA218:as.factor(hours)54 -1.548e-01  1.491e-01  5.090e+02  -1.038
## HemisphereLNA218:as.factor(hours)55 -1.093e-01  1.491e-01  5.090e+02  -0.733
## HemisphereLNA218:as.factor(hours)56 -1.534e-01  1.491e-01  5.090e+02  -1.029
## HemisphereLNA218:as.factor(hours)57 -9.246e-02  1.491e-01  5.090e+02  -0.620
## HemisphereLNA218:as.factor(hours)58 -1.489e-01  1.491e-01  5.090e+02  -0.999
## HemisphereLNA218:as.factor(hours)59 -1.318e-01  1.491e-01  5.090e+02  -0.884
## HemisphereLNA218:as.factor(hours)60 -1.682e-01  1.491e-01  5.090e+02  -1.128
## HemisphereLNA218:as.factor(hours)61  8.945e-03  2.697e-01  5.090e+02   0.033
## HemisphereLNA218:as.factor(hours)62 -2.036e-01  2.697e-01  5.090e+02  -0.755
## HemisphereLNA218:as.factor(hours)63 -1.298e-01  1.798e-01  5.090e+02  -0.722
## HemisphereLNA218:as.factor(hours)64 -1.249e-01  1.652e-01  5.090e+02  -0.756
## HemisphereLNA218:as.factor(hours)65 -1.739e-01  1.652e-01  5.090e+02  -1.053
## HemisphereLNA218:as.factor(hours)66 -2.304e-01  1.798e-01  5.090e+02  -1.281
## HemisphereLNA218:as.factor(hours)67 -1.541e-01  1.491e-01  5.090e+02  -1.034
## HemisphereLNA218:as.factor(hours)68 -1.558e-01  1.491e-01  5.090e+02  -1.045
## HemisphereLNA218:as.factor(hours)69 -1.570e-01  1.491e-01  5.090e+02  -1.053
## HemisphereLNA218:as.factor(hours)70 -9.872e-02  1.491e-01  5.090e+02  -0.662
## HemisphereLNA218:as.factor(hours)71 -2.279e-01  2.060e-01  5.090e+02  -1.106
## HemisphereLNA218:as.factor(hours)72 -1.393e-01  2.060e-01  5.090e+02  -0.676
##                                     Pr(>|t|)
## (Intercept)                          < 2e-16 ***
## HemisphereLNA218                    0.505013
## as.factor(hours)2                   0.005125 **
## as.factor(hours)3                   0.000459 ***
## as.factor(hours)4                   5.71e-05 ***
## as.factor(hours)5                   4.45e-05 ***
## as.factor(hours)6                   3.00e-06 ***
## as.factor(hours)7                   2.43e-05 ***
## as.factor(hours)8                   4.15e-07 ***
## as.factor(hours)9                   4.47e-06 ***
## as.factor(hours)10                  6.44e-06 ***
## as.factor(hours)11                  0.000230 ***
## as.factor(hours)12                  0.001039 **
## as.factor(hours)14                  0.783523
## as.factor(hours)15                  0.404045
## as.factor(hours)16                  0.237256
## as.factor(hours)17                  0.113421
## as.factor(hours)18                  0.005983 **
## as.factor(hours)19                  0.074018 .
## as.factor(hours)20                  0.037890 *
## as.factor(hours)21                  0.000754 ***
## as.factor(hours)22                  0.061599 .
## as.factor(hours)23                  0.010364 *
## as.factor(hours)25                  0.433386
## as.factor(hours)26                  0.004787 **
## as.factor(hours)27                  0.001472 **
## as.factor(hours)28                  7.21e-07 ***
## as.factor(hours)29                  2.31e-05 ***
## as.factor(hours)30                  2.69e-07 ***
## as.factor(hours)31                  3.89e-07 ***
```

```
## as.factor(hours)32                   1.30e-07 ***
## as.factor(hours)33                   3.66e-07 ***
## as.factor(hours)34                   1.57e-07 ***
## as.factor(hours)35                   2.76e-07 ***
## as.factor(hours)36                   0.000428 ***
## as.factor(hours)37                   0.075825 .
## as.factor(hours)38                   0.345234
## as.factor(hours)39                   0.816254
## as.factor(hours)40                   0.275351
## as.factor(hours)41                   0.067273 .
## as.factor(hours)42                   0.247637
## as.factor(hours)43                   0.005869 **
## as.factor(hours)44                   0.016660 *
## as.factor(hours)45                   0.001340 **
## as.factor(hours)46                   0.002383 **
## as.factor(hours)47                   0.460793
## as.factor(hours)48                   0.034214 *
## as.factor(hours)49                   0.000846 ***
## as.factor(hours)50                   0.427349
## as.factor(hours)51                   0.005272 **
## as.factor(hours)52                   0.002775 **
## as.factor(hours)53                   9.36e-05 ***
## as.factor(hours)54                   0.000189 ***
## as.factor(hours)55                   1.92e-06 ***
## as.factor(hours)56                   6.79e-06 ***
## as.factor(hours)57                   5.91e-08 ***
## as.factor(hours)58                   3.04e-07 ***
## as.factor(hours)59                   3.04e-06 ***
## as.factor(hours)60                   0.000632 ***
## as.factor(hours)61                   0.209170
## as.factor(hours)62                   0.563734
## as.factor(hours)63                   0.250947
## as.factor(hours)64                   0.929546
## as.factor(hours)65                   0.757079
## as.factor(hours)66                   0.373847
## as.factor(hours)67                   0.243639
## as.factor(hours)68                   0.083942 .
## as.factor(hours)69                   0.032751 *
## as.factor(hours)70                   0.001031 **
## as.factor(hours)71                   0.875058
## as.factor(hours)72                   0.151750
## HemisphereLNA218:as.factor(hours)2  0.372392
## HemisphereLNA218:as.factor(hours)3  0.504077
## HemisphereLNA218:as.factor(hours)4  0.553833
## HemisphereLNA218:as.factor(hours)5  0.584063
## HemisphereLNA218:as.factor(hours)6  0.355947
## HemisphereLNA218:as.factor(hours)7  0.663044
## HemisphereLNA218:as.factor(hours)8  0.646620
## HemisphereLNA218:as.factor(hours)9  0.746433
## HemisphereLNA218:as.factor(hours)10 0.742322
## HemisphereLNA218:as.factor(hours)11 0.749938
## HemisphereLNA218:as.factor(hours)12 0.872970
## HemisphereLNA218:as.factor(hours)14 0.624815
## HemisphereLNA218:as.factor(hours)15 0.358304
```

```
## HemisphereLNA218:as.factor(hours)16 0.349937
## HemisphereLNA218:as.factor(hours)17 0.456574
## HemisphereLNA218:as.factor(hours)18 0.268493
## HemisphereLNA218:as.factor(hours)19 0.688385
## HemisphereLNA218:as.factor(hours)20 0.585981
## HemisphereLNA218:as.factor(hours)21 0.410821
## HemisphereLNA218:as.factor(hours)22 0.513061
## HemisphereLNA218:as.factor(hours)23 0.631038
## HemisphereLNA218:as.factor(hours)25 0.948342
## HemisphereLNA218:as.factor(hours)26 0.999783
## HemisphereLNA218:as.factor(hours)27 0.968642
## HemisphereLNA218:as.factor(hours)28 0.698951
## HemisphereLNA218:as.factor(hours)29 0.969596
## HemisphereLNA218:as.factor(hours)30 0.903029
## HemisphereLNA218:as.factor(hours)31 0.812531
## HemisphereLNA218:as.factor(hours)32 0.877445
## HemisphereLNA218:as.factor(hours)33 0.987884
## HemisphereLNA218:as.factor(hours)34 0.937736
## HemisphereLNA218:as.factor(hours)35 0.922135
## HemisphereLNA218:as.factor(hours)36 0.932861
## HemisphereLNA218:as.factor(hours)37 0.576873
## HemisphereLNA218:as.factor(hours)38 0.886654
## HemisphereLNA218:as.factor(hours)39 0.912071
## HemisphereLNA218:as.factor(hours)40 0.490646
## HemisphereLNA218:as.factor(hours)41 0.881341
## HemisphereLNA218:as.factor(hours)42 0.765415
## HemisphereLNA218:as.factor(hours)43 0.767034
## HemisphereLNA218:as.factor(hours)44 0.932868
## HemisphereLNA218:as.factor(hours)45 0.866959
## HemisphereLNA218:as.factor(hours)46 0.998102
## HemisphereLNA218:as.factor(hours)47 0.397104
## HemisphereLNA218:as.factor(hours)48 0.857505
## HemisphereLNA218:as.factor(hours)49 0.013761 *
## HemisphereLNA218:as.factor(hours)50 0.110770
## HemisphereLNA218:as.factor(hours)51 0.257917
## HemisphereLNA218:as.factor(hours)52 0.188491
## HemisphereLNA218:as.factor(hours)53 0.245225
## HemisphereLNA218:as.factor(hours)54 0.299722
## HemisphereLNA218:as.factor(hours)55 0.463941
## HemisphereLNA218:as.factor(hours)56 0.303985
## HemisphereLNA218:as.factor(hours)57 0.535460
## HemisphereLNA218:as.factor(hours)58 0.318310
## HemisphereLNA218:as.factor(hours)59 0.377112
## HemisphereLNA218:as.factor(hours)60 0.259909
## HemisphereLNA218:as.factor(hours)61 0.973560
## HemisphereLNA218:as.factor(hours)62 0.450776
## HemisphereLNA218:as.factor(hours)63 0.470730
## HemisphereLNA218:as.factor(hours)64 0.449976
## HemisphereLNA218:as.factor(hours)65 0.292961
## HemisphereLNA218:as.factor(hours)66 0.200668
## HemisphereLNA218:as.factor(hours)67 0.301838
## HemisphereLNA218:as.factor(hours)68 0.296561
## HemisphereLNA218:as.factor(hours)69 0.292855
## HemisphereLNA218:as.factor(hours)70 0.508226
```

```
## HemisphereLNA218:as.factor(hours)71 0.269249
## HemisphereLNA218:as.factor(hours)72 0.499282
## ---
## Signif. codes:  0 '***' 0.001 '**' 0.01 '*' 0.05 '.' 0.1 ' ' 1
##
## Correlation matrix not shown by default, as p = 140 > 12.
## Use print(x, correlation=TRUE)  or
##     vcov(x)        if you need it
```

```
emmeans(mod, trt.vs.ctrl ~ Hemisphere | hours, ref="NC")
```

```
## $emmeans
## hours =  1:
##  Hemisphere emmean     SE    df lower.CL upper.CL
##  NC          1.377 0.0871 103.5    1.204    1.550
##  LNA218      1.304 0.0871 103.5    1.131    1.476
##
## hours =  2:
##  Hemisphere emmean     SE    df lower.CL upper.CL
##  NC          1.081 0.0810  80.7    0.919    1.242
##  LNA218      1.140 0.0810  80.7    0.979    1.301
##
## hours =  3:
##  Hemisphere emmean     SE    df lower.CL upper.CL
##  NC          1.005 0.0810  80.7    0.844    1.166
##  LNA218      1.031 0.0810  80.7    0.870    1.192
##
## hours =  4:
##  Hemisphere emmean     SE    df lower.CL upper.CL
##  NC          0.930 0.0871 103.5    0.757    1.102
##  LNA218      0.948 0.0871 103.5    0.776    1.121
##
## hours =  5:
##  Hemisphere emmean     SE    df lower.CL upper.CL
##  NC          0.923 0.0871 103.5    0.750    1.096
##  LNA218      0.935 0.0871 103.5    0.762    1.108
##
## hours =  6:
##  Hemisphere emmean     SE    df lower.CL upper.CL
##  NC          0.879 0.0810  80.7    0.718    1.040
##  LNA218      0.943 0.0810  80.7    0.782    1.104
##
## hours =  7:
##  Hemisphere emmean     SE    df lower.CL upper.CL
##  NC          0.907 0.0871 103.5    0.735    1.080
##  LNA218      0.902 0.0871 103.5    0.729    1.075
##
## hours =  8:
##  Hemisphere emmean     SE    df lower.CL upper.CL
##  NC          0.836 0.0810  80.7    0.675    0.997
##  LNA218      0.831 0.0810  80.7    0.670    0.992
##
## hours =  9:
##  Hemisphere emmean     SE    df lower.CL upper.CL
```

```
##  NC           0.866 0.0871 103.5    0.693    1.038
##  LNA218       0.843 0.0871 103.5    0.670    1.015
##
## hours = 10:
##  Hemisphere emmean     SE    df lower.CL upper.CL
##  NC           0.874 0.0871 103.5    0.702    1.047
##  LNA218       0.852 0.0871 103.5    0.680    1.025
##
## hours = 11:
##  Hemisphere emmean     SE    df lower.CL upper.CL
##  NC           0.986 0.0810  80.7    0.825    1.147
##  LNA218       0.960 0.0810  80.7    0.799    1.121
##
## hours = 12:
##  Hemisphere emmean     SE    df lower.CL upper.CL
##  NC           1.029 0.0810  80.7    0.868    1.190
##  LNA218       0.979 0.0810  80.7    0.818    1.141
##
## hours = 14:
##  Hemisphere emmean     SE    df lower.CL upper.CL
##  NC           1.324 0.1790 443.9    0.973    1.676
##  LNA218       1.383 0.1790 443.9    1.031    1.735
##
## hours = 15:
##  Hemisphere emmean     SE    df lower.CL upper.CL
##  NC           1.279 0.0954 139.2    1.091    1.468
##  LNA218       1.358 0.0954 139.2    1.169    1.547
##
## hours = 16:
##  Hemisphere emmean     SE    df lower.CL upper.CL
##  NC           1.226 0.1080 198.4    1.013    1.439
##  LNA218       1.321 0.1080 198.4    1.108    1.534
##
## hours = 17:
##  Hemisphere emmean     SE    df lower.CL upper.CL
##  NC           1.146 0.1295 298.9    0.891    1.400
##  LNA218       1.226 0.1295 298.9    0.971    1.480
##
## hours = 18:
##  Hemisphere emmean     SE    df lower.CL upper.CL
##  NC           1.054 0.0954 139.2    0.865    1.243
##  LNA218       1.164 0.0954 139.2    0.975    1.352
##
## hours = 19:
##  Hemisphere emmean     SE    df lower.CL upper.CL
##  NC           1.149 0.1080 198.4    0.936    1.362
##  LNA218       1.148 0.1080 198.4    0.935    1.361
##
## hours = 20:
##  Hemisphere emmean     SE    df lower.CL upper.CL
##  NC           1.158 0.0810  80.7    0.996    1.319
##  LNA218       1.165 0.0810  80.7    1.004    1.326
##
## hours = 21:
```

```
##  Hemisphere emmean     SE    df lower.CL upper.CL
##  NC          1.020 0.0810  80.7    0.858    1.181
##  LNA218      1.069 0.0810  80.7    0.908    1.230
##
## hours = 22:
##  Hemisphere emmean     SE    df lower.CL upper.CL
##  NC          1.139 0.1080 198.4    0.926    1.352
##  LNA218      1.183 0.1080 198.4    0.970    1.396
##
## hours = 23:
##  Hemisphere emmean     SE    df lower.CL upper.CL
##  NC          1.049 0.1080 198.4    0.837    1.262
##  LNA218      1.062 0.1080 198.4    0.850    1.275
##
## hours = 25:
##  Hemisphere emmean     SE    df lower.CL upper.CL
##  NC          1.294 0.0810  80.7    1.133    1.456
##  LNA218      1.231 0.0810  80.7    1.069    1.392
##
## hours = 26:
##  Hemisphere emmean     SE    df lower.CL upper.CL
##  NC          1.078 0.0810  80.7    0.917    1.239
##  LNA218      1.005 0.0810  80.7    0.844    1.166
##
## hours = 27:
##  Hemisphere emmean     SE    df lower.CL upper.CL
##  NC          1.040 0.0810  80.7    0.879    1.201
##  LNA218      0.960 0.0810  80.7    0.799    1.122
##
## hours = 28:
##  Hemisphere emmean     SE    df lower.CL upper.CL
##  NC          0.848 0.0810  80.7    0.687    1.009
##  LNA218      0.832 0.0810  80.7    0.671    0.993
##
## hours = 29:
##  Hemisphere emmean     SE    df lower.CL upper.CL
##  NC          0.926 0.0810  80.7    0.765    1.088
##  LNA218      0.859 0.0810  80.7    0.698    1.020
##
## hours = 30:
##  Hemisphere emmean     SE    df lower.CL upper.CL
##  NC          0.827 0.0810  80.7    0.666    0.988
##  LNA218      0.772 0.0810  80.7    0.611    0.933
##
## hours = 31:
##  Hemisphere emmean     SE    df lower.CL upper.CL
##  NC          0.835 0.0810  80.7    0.674    0.996
##  LNA218      0.797 0.0810  80.7    0.636    0.958
##
## hours = 32:
##  Hemisphere emmean     SE    df lower.CL upper.CL
##  NC          0.812 0.0810  80.7    0.651    0.973
##  LNA218      0.762 0.0810  80.7    0.601    0.923
##
```

```
## hours = 33:
##  Hemisphere emmean     SE    df lower.CL upper.CL
##  NC          0.833 0.0810  80.7    0.672    0.995
##  LNA218      0.762 0.0810  80.7    0.601    0.923
##
## hours = 34:
##  Hemisphere emmean     SE    df lower.CL upper.CL
##  NC          0.816 0.0810  80.7    0.655    0.977
##  LNA218      0.754 0.0810  80.7    0.593    0.915
##
## hours = 35:
##  Hemisphere emmean     SE    df lower.CL upper.CL
##  NC          0.828 0.0810  80.7    0.666    0.989
##  LNA218      0.769 0.0810  80.7    0.608    0.930
##
## hours = 36:
##  Hemisphere emmean     SE    df lower.CL upper.CL
##  NC          0.986 0.0871 103.5    0.814    1.159
##  LNA218      0.926 0.0871 103.5    0.753    1.099
##
## hours = 37:
##  Hemisphere emmean     SE    df lower.CL upper.CL
##  NC          1.037 0.1791 443.8    0.685    1.389
##  LNA218      0.813 0.1791 443.8    0.461    1.165
##
## hours = 38:
##  Hemisphere emmean     SE    df lower.CL upper.CL
##  NC          1.239 0.1295 298.9    0.984    1.494
##  LNA218      1.195 0.1295 298.9    0.940    1.450
##
## hours = 39:
##  Hemisphere emmean     SE    df lower.CL upper.CL
##  NC          1.347 0.1080 198.4    1.135    1.560
##  LNA218      1.254 0.1080 198.4    1.041    1.467
##
## hours = 40:
##  Hemisphere emmean     SE    df lower.CL upper.CL
##  NC          1.238 0.1080 198.4    1.025    1.451
##  LNA218      1.289 0.1080 198.4    1.076    1.502
##
## hours = 41:
##  Hemisphere emmean     SE    df lower.CL upper.CL
##  NC          1.144 0.1080 198.4    0.931    1.357
##  LNA218      1.097 0.1080 198.4    0.884    1.310
##
## hours = 42:
##  Hemisphere emmean     SE    df lower.CL upper.CL
##  NC          1.242 0.0954 139.1    1.053    1.430
##  LNA218      1.119 0.0954 139.1    0.930    1.308
##
## hours = 43:
##  Hemisphere emmean     SE    df lower.CL upper.CL
##  NC          1.053 0.0954 139.1    0.865    1.242
##  LNA218      1.029 0.0954 139.1    0.840    1.218
```

```
##
## hours = 44:
##  Hemisphere emmean     SE    df lower.CL upper.CL
##  NC          1.124 0.0810  80.7    0.963    1.285
##  LNA218      1.063 0.0810  80.7    0.902    1.224
##
## hours = 45:
##  Hemisphere emmean     SE    df lower.CL upper.CL
##  NC          1.037 0.0810  80.7    0.876    1.198
##  LNA218      0.988 0.0810  80.7    0.827    1.150
##
## hours = 46:
##  Hemisphere emmean     SE    df lower.CL upper.CL
##  NC          1.055 0.0810  80.7    0.894    1.216
##  LNA218      0.981 0.0810  80.7    0.820    1.142
##
## hours = 47:
##  Hemisphere emmean     SE    df lower.CL upper.CL
##  NC          1.269 0.1295 298.9    1.015    1.524
##  LNA218      1.370 0.1295 298.9    1.116    1.625
##
## hours = 48:
##  Hemisphere emmean     SE    df lower.CL upper.CL
##  NC          0.971 0.1791 443.8    0.619    1.323
##  LNA218      0.946 0.1791 443.8    0.594    1.298
##
## hours = 49:
##  Hemisphere emmean     SE    df lower.CL upper.CL
##  NC          1.747 0.0871 103.5    1.574    1.920
##  LNA218      1.289 0.0871 103.5    1.116    1.461
##
## hours = 50:
##  Hemisphere emmean     SE    df lower.CL upper.CL
##  NC          1.290 0.0871 103.5    1.117    1.462
##  LNA218      0.967 0.0871 103.5    0.795    1.140
##
## hours = 51:
##  Hemisphere emmean     SE    df lower.CL upper.CL
##  NC          1.082 0.0810  80.7    0.920    1.243
##  LNA218      0.839 0.0810  80.7    0.678    1.000
##
## hours = 52:
##  Hemisphere emmean     SE    df lower.CL upper.CL
##  NC          1.060 0.0810  80.7    0.899    1.221
##  LNA218      0.790 0.0810  80.7    0.629    0.951
##
## hours = 53:
##  Hemisphere emmean     SE    df lower.CL upper.CL
##  NC          0.962 0.0810  80.7    0.801    1.123
##  LNA218      0.715 0.0810  80.7    0.554    0.876
##
## hours = 54:
##  Hemisphere emmean     SE    df lower.CL upper.CL
##  NC          0.980 0.0810  80.7    0.819    1.142
```

```
##  LNA218      0.752 0.0810  80.7    0.591    0.913
##
## hours = 55:
##  Hemisphere emmean     SE    df lower.CL upper.CL
##  NC          0.869 0.0810  80.7    0.708    1.030
##  LNA218      0.686 0.0810  80.7    0.525    0.847
##
## hours = 56:
##  Hemisphere emmean     SE    df lower.CL upper.CL
##  NC          0.897 0.0810  80.7    0.736    1.059
##  LNA218      0.671 0.0810  80.7    0.509    0.832
##
## hours = 57:
##  Hemisphere emmean     SE    df lower.CL upper.CL
##  NC          0.797 0.0810  80.7    0.635    0.958
##  LNA218      0.631 0.0810  80.7    0.469    0.792
##
## hours = 58:
##  Hemisphere emmean     SE    df lower.CL upper.CL
##  NC          0.830 0.0810  80.7    0.668    0.991
##  LNA218      0.607 0.0810  80.7    0.446    0.768
##
## hours = 59:
##  Hemisphere emmean     SE    df lower.CL upper.CL
##  NC          0.879 0.0810  80.7    0.718    1.040
##  LNA218      0.674 0.0810  80.7    0.513    0.835
##
## hours = 60:
##  Hemisphere emmean     SE    df lower.CL upper.CL
##  NC          1.014 0.0810  80.7    0.853    1.175
##  LNA218      0.773 0.0810  80.7    0.612    0.934
##
## hours = 61:
##  Hemisphere emmean     SE    df lower.CL upper.CL
##  NC          1.137 0.1790 443.9    0.785    1.488
##  LNA218      1.072 0.1790 443.9    0.720    1.424
##
## hours = 62:
##  Hemisphere emmean     SE    df lower.CL upper.CL
##  NC          1.488 0.1790 443.9    1.136    1.839
##  LNA218      1.211 0.1790 443.9    0.859    1.562
##
## hours = 63:
##  Hemisphere emmean     SE    df lower.CL upper.CL
##  NC          1.231 0.1080 198.4    1.018    1.444
##  LNA218      1.027 0.1080 198.4    0.815    1.240
##
## hours = 64:
##  Hemisphere emmean     SE    df lower.CL upper.CL
##  NC          1.367 0.0954 139.1    1.178    1.555
##  LNA218      1.168 0.0954 139.1    0.980    1.357
##
## hours = 65:
##  Hemisphere emmean     SE    df lower.CL upper.CL
```

```
##  NC          1.341 0.0954 139.1    1.152    1.530
##  LNA218      1.094 0.0954 139.1    0.905    1.282
##
## hours = 66:
##  Hemisphere emmean     SE    df lower.CL upper.CL
##  NC          1.491 0.1080 198.4    1.278    1.703
##  LNA218      1.187 0.1080 198.4    0.974    1.400
##
## hours = 67:
##  Hemisphere emmean     SE    df lower.CL upper.CL
##  NC          1.254 0.0810  80.7    1.093    1.415
##  LNA218      1.026 0.0810  80.7    0.865    1.188
##
## hours = 68:
##  Hemisphere emmean     SE    df lower.CL upper.CL
##  NC          1.194 0.0810  80.7    1.033    1.356
##  LNA218      0.965 0.0810  80.7    0.804    1.126
##
## hours = 69:
##  Hemisphere emmean     SE    df lower.CL upper.CL
##  NC          1.151 0.0810  80.7    0.990    1.312
##  LNA218      0.921 0.0810  80.7    0.760    1.082
##
## hours = 70:
##  Hemisphere emmean     SE    df lower.CL upper.CL
##  NC          1.029 0.0810  80.7    0.868    1.190
##  LNA218      0.857 0.0810  80.7    0.696    1.018
##
## hours = 71:
##  Hemisphere emmean     SE    df lower.CL upper.CL
##  NC          1.354 0.1295 298.9    1.099    1.609
##  LNA218      1.053 0.1295 298.9    0.798    1.308
##
## hours = 72:
##  Hemisphere emmean     SE    df lower.CL upper.CL
##  NC          1.587 0.1295 298.9    1.332    1.841
##  LNA218      1.374 0.1295 298.9    1.119    1.629
##
## Degrees-of-freedom method: kenward-roger
## Confidence level used: 0.95
##
## $contrasts
## hours =  1:
##  contrast    estimate    SE  df t.ratio p.value
##  LNA218 - NC -0.07346 0.110 509  -0.667  0.5050
##
## hours =  2:
##  contrast    estimate    SE  df t.ratio p.value
##  LNA218 - NC  0.05966 0.101 509   0.593  0.5531
##
## hours =  3:
##  contrast    estimate    SE  df t.ratio p.value
##  LNA218 - NC  0.02622 0.101 509   0.261  0.7943
##
```

```
## hours =  4:
##  contrast    estimate    SE  df t.ratio p.value
##  LNA218 - NC  0.01880 0.110 509   0.171  0.8645
##
## hours =  5:
##  contrast    estimate    SE  df t.ratio p.value
##  LNA218 - NC  0.01185 0.110 509   0.108  0.9143
##
## hours =  6:
##  contrast    estimate    SE  df t.ratio p.value
##  LNA218 - NC  0.06430 0.101 509   0.640  0.5227
##
## hours =  7:
##  contrast    estimate    SE  df t.ratio p.value
##  LNA218 - NC -0.00557 0.110 509  -0.051  0.9597
##
## hours =  8:
##  contrast    estimate    SE  df t.ratio p.value
##  LNA218 - NC -0.00506 0.101 509  -0.050  0.9599
##
## hours =  9:
##  contrast    estimate    SE  df t.ratio p.value
##  LNA218 - NC -0.02308 0.110 509  -0.210  0.8341
##
## hours = 10:
##  contrast    estimate    SE  df t.ratio p.value
##  LNA218 - NC -0.02223 0.110 509  -0.202  0.8401
##
## hours = 11:
##  contrast    estimate    SE  df t.ratio p.value
##  LNA218 - NC -0.02591 0.101 509  -0.258  0.7967
##
## hours = 12:
##  contrast    estimate    SE  df t.ratio p.value
##  LNA218 - NC -0.04961 0.101 509  -0.493  0.6219
##
## hours = 14:
##  contrast    estimate    SE  df t.ratio p.value
##  LNA218 - NC  0.05853 0.246 509   0.238  0.8122
##
## hours = 15:
##  contrast    estimate    SE  df t.ratio p.value
##  LNA218 - NC  0.07841 0.123 509   0.637  0.5245
##
## hours = 16:
##  contrast    estimate    SE  df t.ratio p.value
##  LNA218 - NC  0.09478 0.142 509   0.667  0.5053
##
## hours = 17:
##  contrast    estimate    SE  df t.ratio p.value
##  LNA218 - NC  0.08004 0.174 509   0.460  0.6460
##
## hours = 18:
##  contrast    estimate    SE  df t.ratio p.value
```

```
##  LNA218 - NC  0.10952 0.123 509   0.890  0.3741
##
## hours = 19:
##  contrast    estimate    SE  df t.ratio p.value
##  LNA218 - NC -0.00130 0.142 509  -0.009  0.9927
##
## hours = 20:
##  contrast    estimate    SE  df t.ratio p.value
##  LNA218 - NC  0.00780 0.101 509   0.078  0.9382
##
## hours = 21:
##  contrast    estimate    SE  df t.ratio p.value
##  LNA218 - NC  0.04927 0.101 509   0.490  0.6243
##
## hours = 22:
##  contrast    estimate    SE  df t.ratio p.value
##  LNA218 - NC  0.04424 0.142 509   0.311  0.7558
##
## hours = 23:
##  contrast    estimate    SE  df t.ratio p.value
##  LNA218 - NC  0.01296 0.142 509   0.091  0.9274
##
## hours = 25:
##  contrast    estimate    SE  df t.ratio p.value
##  LNA218 - NC -0.06380 0.101 509  -0.635  0.5260
##
## hours = 26:
##  contrast    estimate    SE  df t.ratio p.value
##  LNA218 - NC -0.07342 0.101 509  -0.730  0.4655
##
## hours = 27:
##  contrast    estimate    SE  df t.ratio p.value
##  LNA218 - NC -0.07933 0.101 509  -0.789  0.4304
##
## hours = 28:
##  contrast    estimate    SE  df t.ratio p.value
##  LNA218 - NC -0.01576 0.101 509  -0.157  0.8754
##
## hours = 29:
##  contrast    estimate    SE  df t.ratio p.value
##  LNA218 - NC -0.06778 0.101 509  -0.674  0.5005
##
## hours = 30:
##  contrast    estimate    SE  df t.ratio p.value
##  LNA218 - NC -0.05529 0.101 509  -0.550  0.5826
##
## hours = 31:
##  contrast    estimate    SE  df t.ratio p.value
##  LNA218 - NC -0.03808 0.101 509  -0.379  0.7050
##
## hours = 32:
##  contrast    estimate    SE  df t.ratio p.value
##  LNA218 - NC -0.05046 0.101 509  -0.502  0.6159
##
```

```
## hours = 33:
##  contrast    estimate    SE  df t.ratio p.value
##  LNA218 - NC -0.07120 0.101 509  -0.708  0.4791
##
## hours = 34:
##  contrast    estimate    SE  df t.ratio p.value
##  LNA218 - NC -0.06181 0.101 509  -0.615  0.5389
##
## hours = 35:
##  contrast    estimate    SE  df t.ratio p.value
##  LNA218 - NC -0.05888 0.101 509  -0.586  0.5583
##
## hours = 36:
##  contrast    estimate    SE  df t.ratio p.value
##  LNA218 - NC -0.06033 0.110 509  -0.548  0.5840
##
## hours = 37:
##  contrast    estimate    SE  df t.ratio p.value
##  LNA218 - NC -0.22406 0.246 509  -0.910  0.3633
##
## hours = 38:
##  contrast    estimate    SE  df t.ratio p.value
##  LNA218 - NC -0.04408 0.174 509  -0.253  0.8002
##
## hours = 39:
##  contrast    estimate    SE  df t.ratio p.value
##  LNA218 - NC -0.09333 0.142 509  -0.656  0.5118
##
## hours = 40:
##  contrast    estimate    SE  df t.ratio p.value
##  LNA218 - NC  0.05058 0.142 509   0.356  0.7222
##
## hours = 41:
##  contrast    estimate    SE  df t.ratio p.value
##  LNA218 - NC -0.04660 0.142 509  -0.328  0.7432
##
## hours = 42:
##  contrast    estimate    SE  df t.ratio p.value
##  LNA218 - NC -0.12277 0.123 509  -0.997  0.3191
##
## hours = 43:
##  contrast    estimate    SE  df t.ratio p.value
##  LNA218 - NC -0.02450 0.123 509  -0.199  0.8424
##
## hours = 44:
##  contrast    estimate    SE  df t.ratio p.value
##  LNA218 - NC -0.06089 0.101 509  -0.606  0.5449
##
## hours = 45:
##  contrast    estimate    SE  df t.ratio p.value
##  LNA218 - NC -0.04847 0.101 509  -0.482  0.6299
##
## hours = 46:
##  contrast    estimate    SE  df t.ratio p.value
```

```
##  LNA218 - NC -0.07382 0.101 509  -0.734  0.4631
##
## hours = 47:
##  contrast    estimate    SE  df t.ratio p.value
##  LNA218 - NC  0.10114 0.174 509   0.581  0.5616
##
## hours = 48:
##  contrast    estimate    SE  df t.ratio p.value
##  LNA218 - NC -0.02500 0.246 509  -0.102  0.9192
##
## hours = 49:
##  contrast    estimate    SE  df t.ratio p.value
##  LNA218 - NC -0.45844 0.110 509  -4.163  <.0001
##
## hours = 50:
##  contrast    estimate    SE  df t.ratio p.value
##  LNA218 - NC -0.32225 0.110 509  -2.926  0.0036
##
## hours = 51:
##  contrast    estimate    SE  df t.ratio p.value
##  LNA218 - NC -0.24234 0.101 509  -2.411  0.0163
##
## hours = 52:
##  contrast    estimate    SE  df t.ratio p.value
##  LNA218 - NC -0.26980 0.101 509  -2.684  0.0075
##
## hours = 53:
##  contrast    estimate    SE  df t.ratio p.value
##  LNA218 - NC -0.24692 0.101 509  -2.456  0.0144
##
## hours = 54:
##  contrast    estimate    SE  df t.ratio p.value
##  LNA218 - NC -0.22824 0.101 509  -2.271  0.0236
##
## hours = 55:
##  contrast    estimate    SE  df t.ratio p.value
##  LNA218 - NC -0.18274 0.101 509  -1.818  0.0697
##
## hours = 56:
##  contrast    estimate    SE  df t.ratio p.value
##  LNA218 - NC -0.22688 0.101 509  -2.257  0.0244
##
## hours = 57:
##  contrast    estimate    SE  df t.ratio p.value
##  LNA218 - NC -0.16592 0.101 509  -1.651  0.0994
##
## hours = 58:
##  contrast    estimate    SE  df t.ratio p.value
##  LNA218 - NC -0.22240 0.101 509  -2.212  0.0274
##
## hours = 59:
##  contrast    estimate    SE  df t.ratio p.value
##  LNA218 - NC -0.20527 0.101 509  -2.042  0.0417
##
```

```
## hours = 60:
##  contrast    estimate    SE  df t.ratio p.value
##  LNA218 - NC -0.24163 0.101 509  -2.404  0.0166
##
## hours = 61:
##  contrast    estimate    SE  df t.ratio p.value
##  LNA218 - NC -0.06452 0.246 509  -0.262  0.7934
##
## hours = 62:
##  contrast    estimate    SE  df t.ratio p.value
##  LNA218 - NC -0.27703 0.246 509  -1.125  0.2611
##
## hours = 63:
##  contrast    estimate    SE  df t.ratio p.value
##  LNA218 - NC -0.20326 0.142 509  -1.430  0.1534
##
## hours = 64:
##  contrast    estimate    SE  df t.ratio p.value
##  LNA218 - NC -0.19834 0.123 509  -1.611  0.1078
##
## hours = 65:
##  contrast    estimate    SE  df t.ratio p.value
##  LNA218 - NC -0.24735 0.123 509  -2.009  0.0451
##
## hours = 66:
##  contrast    estimate    SE  df t.ratio p.value
##  LNA218 - NC -0.30387 0.142 509  -2.137  0.0330
##
## hours = 67:
##  contrast    estimate    SE  df t.ratio p.value
##  LNA218 - NC -0.22757 0.101 509  -2.264  0.0240
##
## hours = 68:
##  contrast    estimate    SE  df t.ratio p.value
##  LNA218 - NC -0.22926 0.101 509  -2.281  0.0230
##
## hours = 69:
##  contrast    estimate    SE  df t.ratio p.value
##  LNA218 - NC -0.23046 0.101 509  -2.293  0.0223
##
## hours = 70:
##  contrast    estimate    SE  df t.ratio p.value
##  LNA218 - NC -0.17218 0.101 509  -1.713  0.0874
##
## hours = 71:
##  contrast    estimate    SE  df t.ratio p.value
##  LNA218 - NC -0.30132 0.174 509  -1.731  0.0841
##
## hours = 72:
##  contrast    estimate    SE  df t.ratio p.value
##  LNA218 - NC -0.21275 0.174 509  -1.222  0.2223
##
## Degrees-of-freedom method: kenward-roger
```

**Figure 6G: Binned SWA**

```
swa.ag <- allData$swa.ag

mod <- lmer(corrected ~ Hemisphere*Days*Phase + (1|Mouse), swa.ag)
summary(mod)
```

```
## Linear mixed model fit by REML. t-tests use Satterthwaite's method [
## lmerModLmerTest]
## Formula: corrected ~ Hemisphere * Days * Phase + (1 | Mouse)
##    Data: swa.ag
##
## REML criterion at convergence: -45.3
##
## Scaled residuals:
##      Min       1Q   Median       3Q      Max
## -1.98870 -0.63130  0.06924  0.51838  2.99392
##
## Random effects:
##  Groups   Name        Variance Std.Dev.
##  Mouse    (Intercept) 0.009096 0.09537
##  Residual             0.016228 0.12739
## Number of obs: 72, groups:  Mouse, 6
##
## Fixed effects:
##                                       Estimate Std. Error       df t value
## (Intercept)                            1.06307    0.06497 24.80260  16.363
## HemisphereLNA218                       0.02928    0.07355 55.00000   0.398
## DaysPID 3                             -0.00325    0.07355 55.00000  -0.044
## DaysPID 9                              0.15602    0.07355 55.00000   2.121
## PhaseDark                             -0.09488    0.07355 55.00000  -1.290
## HemisphereLNA218:DaysPID 3            -0.08617    0.10401 55.00000  -0.828
## HemisphereLNA218:DaysPID 9            -0.29069    0.10401 55.00000  -2.795
## HemisphereLNA218:PhaseDark            -0.02368    0.10401 55.00000  -0.228
## DaysPID 3:PhaseDark                   -0.02191    0.10401 55.00000  -0.211
## DaysPID 9:PhaseDark                   -0.12477    0.10401 55.00000  -1.200
## HemisphereLNA218:DaysPID 3:PhaseDark   0.02653    0.14710 55.00000   0.180
## HemisphereLNA218:DaysPID 9:PhaseDark   0.07793    0.14710 55.00000   0.530
##                                       Pr(>|t|)
## (Intercept)                           8.46e-15 ***
## HemisphereLNA218                       0.69211
## DaysPID 3                              0.96492
## DaysPID 9                              0.03842 *
## PhaseDark                              0.20246
## HemisphereLNA218:DaysPID 3             0.41103
## HemisphereLNA218:DaysPID 9             0.00714 **
## HemisphereLNA218:PhaseDark             0.82078
## DaysPID 3:PhaseDark                    0.83396
## DaysPID 9:PhaseDark                    0.23547
## HemisphereLNA218:DaysPID 3:PhaseDark   0.85754
## HemisphereLNA218:DaysPID 9:PhaseDark   0.59840
## ---
## Signif. codes:  0 '***' 0.001 '**' 0.01 '*' 0.05 '.' 0.1 ' ' 1
##
```

```
## Correlation of Fixed Effects:
##                (Intr) HmLNA218 DyPID3 DyPID9 PhsDrk HmLNA218:DPID3
## HmsphLNA218    -0.566
## DaysPID 3      -0.566  0.500
## DaysPID 9      -0.566  0.500    0.500
## PhaseDark      -0.566  0.500    0.500  0.500
## HmLNA218:DPID3  0.400 -0.707   -0.707 -0.354 -0.354
## HmLNA218:DPID9  0.400 -0.707   -0.354 -0.707 -0.354  0.500
## HmLNA218:PD     0.400 -0.707   -0.354 -0.354 -0.707  0.500
## DysPID3:PhD     0.400 -0.354   -0.707 -0.354 -0.707  0.500
## DysPID9:PhD     0.400 -0.354   -0.354 -0.707 -0.707  0.250
## HLNA218:DPID3: -0.283  0.500    0.500  0.250  0.500 -0.707
## HLNA218:DPID9: -0.283  0.500    0.250  0.500  0.500 -0.354
##                HmLNA218:DPID9 HLNA218: DPID3: DPID9: HLNA218:DPID3:
## HmsphLNA218
## DaysPID 3
## DaysPID 9
## PhaseDark
## HmLNA218:DPID3
## HmLNA218:DPID9
## HmLNA218:PD     0.500
## DysPID3:PhD     0.250          0.500
## DysPID9:PhD     0.500          0.500    0.500
## HLNA218:DPID3: -0.354         -0.707   -0.707 -0.354
## HLNA218:DPID9: -0.707         -0.707   -0.354 -0.707  0.500
```

```
emmeans(mod, trt.vs.ctrl ~ Hemisphere | Phase | Days, ref="NC")
```

```
## $emmeans
## Phase = Light, Days = Baseline:
##  Hemisphere emmean    SE   df lower.CL upper.CL
##  NC          1.063 0.065 24.8    0.929    1.197
##  LNA218      1.092 0.065 24.8    0.958    1.226
##
## Phase = Dark, Days = Baseline:
##  Hemisphere emmean    SE   df lower.CL upper.CL
##  NC          0.968 0.065 24.8    0.834    1.102
##  LNA218      0.974 0.065 24.8    0.840    1.108
##
## Phase = Light, Days = PID 3:
##  Hemisphere emmean    SE   df lower.CL upper.CL
##  NC          1.060 0.065 24.8    0.926    1.194
##  LNA218      1.003 0.065 24.8    0.869    1.137
##
## Phase = Dark, Days = PID 3:
##  Hemisphere emmean    SE   df lower.CL upper.CL
##  NC          0.943 0.065 24.8    0.809    1.077
##  LNA218      0.889 0.065 24.8    0.755    1.023
##
## Phase = Light, Days = PID 9:
##  Hemisphere emmean    SE   df lower.CL upper.CL
##  NC          1.219 0.065 24.8    1.085    1.353
##  LNA218      0.958 0.065 24.8    0.824    1.092
##
## Phase = Dark, Days = PID 9:
```

```
##  Hemisphere emmean    SE   df lower.CL upper.CL
##  NC          0.999 0.065 24.8    0.866    1.133
##  LNA218      0.792 0.065 24.8    0.658    0.926
##
## Degrees-of-freedom method: kenward-roger
## Confidence level used: 0.95
##
## $contrasts
## Phase = Light, Days = Baseline:
##  contrast    estimate     SE df t.ratio p.value
##  LNA218 - NC   0.0293 0.0735 55   0.398  0.6921
##
## Phase = Dark, Days = Baseline:
##  contrast    estimate     SE df t.ratio p.value
##  LNA218 - NC   0.0056 0.0735 55   0.076  0.9396
##
## Phase = Light, Days = PID 3:
##  contrast    estimate     SE df t.ratio p.value
##  LNA218 - NC  -0.0569 0.0735 55  -0.773  0.4426
##
## Phase = Dark, Days = PID 3:
##  contrast    estimate     SE df t.ratio p.value
##  LNA218 - NC  -0.0540 0.0735 55  -0.735  0.4657
##
## Phase = Light, Days = PID 9:
##  contrast    estimate     SE df t.ratio p.value
##  LNA218 - NC  -0.2614 0.0735 55  -3.554  0.0008
##
## Phase = Dark, Days = PID 9:
##  contrast    estimate     SE df t.ratio p.value
##  LNA218 - NC  -0.2072 0.0735 55  -2.817  0.0067
##
## Degrees-of-freedom method: kenward-roger
```

```
emmeans(mod, pairwise ~ Days| Hemisphere | Phase)
```

```
## $emmeans
## Hemisphere = NC, Phase = Light:
##  Days     emmean    SE   df lower.CL upper.CL
##  Baseline  1.063 0.065 24.8    0.929    1.197
##  PID 3     1.060 0.065 24.8    0.926    1.194
##  PID 9     1.219 0.065 24.8    1.085    1.353
##
## Hemisphere = LNA218, Phase = Light:
##  Days     emmean    SE   df lower.CL upper.CL
##  Baseline  1.092 0.065 24.8    0.958    1.226
##  PID 3     1.003 0.065 24.8    0.869    1.137
##  PID 9     0.958 0.065 24.8    0.824    1.092
##
## Hemisphere = NC, Phase = Dark:
##  Days     emmean    SE   df lower.CL upper.CL
##  Baseline  0.968 0.065 24.8    0.834    1.102
##  PID 3     0.943 0.065 24.8    0.809    1.077
##  PID 9     0.999 0.065 24.8    0.866    1.133
##
```

```
## Hemisphere = LNA218, Phase = Dark:
##  Days     emmean    SE   df lower.CL upper.CL
##  Baseline  0.974 0.065 24.8    0.840    1.108
##  PID 3     0.889 0.065 24.8    0.755    1.023
##  PID 9     0.792 0.065 24.8    0.658    0.926
##
## Degrees-of-freedom method: kenward-roger
## Confidence level used: 0.95
##
## $contrasts
## Hemisphere = NC, Phase = Light:
##  contrast         estimate     SE df t.ratio p.value
##  Baseline - PID 3  0.00325 0.0735 55   0.044  0.9989
##  Baseline - PID 9 -0.15602 0.0735 55  -2.121  0.0948
##  PID 3 - PID 9    -0.15927 0.0735 55  -2.166  0.0863
##
## Hemisphere = LNA218, Phase = Light:
##  contrast         estimate     SE df t.ratio p.value
##  Baseline - PID 3  0.08942 0.0735 55   1.216  0.4491
##  Baseline - PID 9  0.13467 0.0735 55   1.831  0.1691
##  PID 3 - PID 9     0.04525 0.0735 55   0.615  0.8124
##
## Hemisphere = NC, Phase = Dark:
##  contrast         estimate     SE df t.ratio p.value
##  Baseline - PID 3  0.02516 0.0735 55   0.342  0.9376
##  Baseline - PID 9 -0.03125 0.0735 55  -0.425  0.9054
##  PID 3 - PID 9    -0.05641 0.0735 55  -0.767  0.7247
##
## Hemisphere = LNA218, Phase = Dark:
##  contrast         estimate     SE df t.ratio p.value
##  Baseline - PID 3  0.08479 0.0735 55   1.153  0.4862
##  Baseline - PID 9  0.18151 0.0735 55   2.468  0.0435
##  PID 3 - PID 9     0.09671 0.0735 55   1.315  0.3930
##
## Degrees-of-freedom method: kenward-roger
## P value adjustment: tukey method for comparing a family of 3 estimates
```

**Figure 6I: Sleep Amounts**

```
sleep.amount.melt <- allData$sleep.amount.melt

mod <- lmer(Time ~ ZT*Day*Vigilance + (1|Mouse), subset(sleep.amount.melt))
```

```
## boundary (singular) fit: see help('isSingular')
```

```
summary(mod)
```

```
## Linear mixed model fit by REML. t-tests use Satterthwaite's method [
## lmerModLmerTest]
## Formula: Time ~ ZT * Day * Vigilance + (1 | Mouse)
##    Data: subset(sleep.amount.melt)
##
## REML criterion at convergence: -443.9
##
```

```
## Scaled residuals:
##     Min      1Q  Median      3Q     Max
## -3.7370 -0.2832  0.0000  0.3089  4.4052
##
## Random effects:
##  Groups   Name        Variance  Std.Dev.
##  Mouse    (Intercept) 6.974e-35 8.351e-18
##  Residual             1.192e-02 1.092e-01
## Number of obs: 432, groups:  Mouse, 6
##
## Fixed effects:
##                                  Estimate Std. Error         df t value
## (Intercept)                     4.707e-01  4.458e-02  3.600e+02  10.560
## ZTZT2-4                         4.157e-02  6.304e-02  3.600e+02   0.659
## ZTZT4-6                         3.065e-02  6.304e-02  3.600e+02   0.486
## ZTZT6-8                        -3.241e-03  6.304e-02  3.600e+02  -0.051
## ZTZT8-10                       -6.898e-02  6.304e-02  3.600e+02  -1.094
## ZTZT10-12                      -1.180e-01  6.304e-02  3.600e+02  -1.871
## ZTZT12-14                      -4.497e-01  6.304e-02  3.600e+02  -7.134
## ZTZT14-16                      -3.097e-01  6.304e-02  3.600e+02  -4.913
## ZTZT16-18                      -2.323e-01  6.304e-02  3.600e+02  -3.685
## ZTZT18-20                      -1.681e-01  6.304e-02  3.600e+02  -2.666
## ZTZT20-22                      -1.016e-01  6.304e-02  3.600e+02  -1.611
## ZTZT22-24                      -4.035e-01  6.304e-02  3.600e+02  -6.400
## DayDay9                        -6.741e-02  6.304e-02  3.600e+02  -1.069
## VigilanceREM                   -4.006e-01  6.304e-02  3.600e+02  -6.354
## VigilanceWake                  -1.167e-02  6.304e-02  3.600e+02  -0.185
## ZTZT2-4:DayDay9                 9.907e-02  8.916e-02  3.600e+02   1.111
## ZTZT4-6:DayDay9                 1.433e-01  8.916e-02  3.600e+02   1.608
## ZTZT6-8:DayDay9                 1.715e-01  8.916e-02  3.600e+02   1.923
## ZTZT8-10:DayDay9                1.900e-01  8.916e-02  3.600e+02   2.131
## ZTZT10-12:DayDay9               1.852e-01  8.916e-02  3.600e+02   2.077
## ZTZT12-14:DayDay9               6.852e-02  8.916e-02  3.600e+02   0.769
## ZTZT14-16:DayDay9               5.444e-02  8.916e-02  3.600e+02   0.611
## ZTZT16-18:DayDay9               5.370e-03  8.916e-02  3.600e+02   0.060
## ZTZT18-20:DayDay9               2.245e-01  8.916e-02  3.600e+02   2.518
## ZTZT20-22:DayDay9               1.556e-01  8.916e-02  3.600e+02   1.746
## ZTZT22-24:DayDay9               6.052e-02  8.916e-02  3.600e+02   0.679
## ZTZT2-4:VigilanceREM           -2.778e-04  8.916e-02  3.600e+02  -0.003
## ZTZT4-6:VigilanceREM           -6.944e-03  8.916e-02  3.600e+02  -0.078
## ZTZT6-8:VigilanceREM            3.269e-02  8.916e-02  3.600e+02   0.367
## ZTZT8-10:VigilanceREM           8.639e-02  8.916e-02  3.600e+02   0.969
## ZTZT10-12:VigilanceREM          1.187e-01  8.916e-02  3.600e+02   1.331
## ZTZT12-14:VigilanceREM          3.828e-01  8.916e-02  3.600e+02   4.293
## ZTZT14-16:VigilanceREM          2.490e-01  8.916e-02  3.600e+02   2.793
## ZTZT16-18:VigilanceREM          1.898e-01  8.916e-02  3.600e+02   2.129
## ZTZT18-20:VigilanceREM          1.226e-01  8.916e-02  3.600e+02   1.375
## ZTZT20-22:VigilanceREM          6.676e-02  8.916e-02  3.600e+02   0.749
## ZTZT22-24:VigilanceREM          3.361e-01  8.916e-02  3.600e+02   3.769
## ZTZT2-4:VigilanceWake          -1.244e-01  8.916e-02  3.600e+02  -1.396
## ZTZT4-6:VigilanceWake          -8.500e-02  8.916e-02  3.600e+02  -0.953
## ZTZT6-8:VigilanceWake          -2.296e-02  8.916e-02  3.600e+02  -0.258
## ZTZT8-10:VigilanceWake          1.206e-01  8.916e-02  3.600e+02   1.352
## ZTZT10-12:VigilanceWake         2.352e-01  8.916e-02  3.600e+02   2.638
```

```
## ZTZT12-14:VigilanceWake          9.664e-01  8.916e-02  3.600e+02  10.839
## ZTZT14-16:VigilanceWake          6.802e-01  8.916e-02  3.600e+02   7.629
## ZTZT16-18:VigilanceWake          5.071e-01  8.916e-02  3.600e+02   5.688
## ZTZT18-20:VigilanceWake          3.816e-01  8.916e-02  3.600e+02   4.280
## ZTZT20-22:VigilanceWake          2.380e-01  8.916e-02  3.600e+02   2.669
## ZTZT22-24:VigilanceWake          8.743e-01  8.916e-02  3.600e+02   9.807
## DayDay9:VigilanceREM             5.574e-02  8.916e-02  3.600e+02   0.625
## DayDay9:VigilanceWake            1.465e-01  8.916e-02  3.600e+02   1.643
## ZTZT2-4:DayDay9:VigilanceREM    -9.861e-02  1.261e-01  3.600e+02  -0.782
## ZTZT4-6:DayDay9:VigilanceREM    -1.277e-01  1.261e-01  3.600e+02  -1.013
## ZTZT6-8:DayDay9:VigilanceREM    -1.562e-01  1.261e-01  3.600e+02  -1.239
## ZTZT8-10:DayDay9:VigilanceREM   -1.673e-01  1.261e-01  3.600e+02  -1.327
## ZTZT10-12:DayDay9:VigilanceREM  -1.698e-01  1.261e-01  3.600e+02  -1.347
## ZTZT12-14:DayDay9:VigilanceREM  -6.009e-02  1.261e-01  3.600e+02  -0.477
## ZTZT14-16:DayDay9:VigilanceREM  -4.926e-02  1.261e-01  3.600e+02  -0.391
## ZTZT16-18:DayDay9:VigilanceREM  -1.407e-02  1.261e-01  3.600e+02  -0.112
## ZTZT18-20:DayDay9:VigilanceREM  -1.977e-01  1.261e-01  3.600e+02  -1.568
## ZTZT20-22:DayDay9:VigilanceREM  -1.419e-01  1.261e-01  3.600e+02  -1.125
## ZTZT22-24:DayDay9:VigilanceREM  -5.165e-02  1.261e-01  3.600e+02  -0.410
## ZTZT2-4:DayDay9:VigilanceWake   -1.986e-01  1.261e-01  3.600e+02  -1.575
## ZTZT4-6:DayDay9:VigilanceWake   -3.023e-01  1.261e-01  3.600e+02  -2.398
## ZTZT6-8:DayDay9:VigilanceWake   -3.582e-01  1.261e-01  3.600e+02  -2.841
## ZTZT8-10:DayDay9:VigilanceWake  -4.027e-01  1.261e-01  3.600e+02  -3.194
## ZTZT10-12:DayDay9:VigilanceWake -3.857e-01  1.261e-01  3.600e+02  -3.059
## ZTZT12-14:DayDay9:VigilanceWake -1.455e-01  1.261e-01  3.600e+02  -1.154
## ZTZT14-16:DayDay9:VigilanceWake -1.141e-01  1.261e-01  3.600e+02  -0.905
## ZTZT16-18:DayDay9:VigilanceWake -2.037e-03  1.261e-01  3.600e+02  -0.016
## ZTZT18-20:DayDay9:VigilanceWake -4.759e-01  1.261e-01  3.600e+02  -3.775
## ZTZT20-22:DayDay9:VigilanceWake -3.251e-01  1.261e-01  3.600e+02  -2.578
## ZTZT22-24:DayDay9:VigilanceWake -1.299e-01  1.261e-01  3.600e+02  -1.030
##                                 Pr(>|t|)
## (Intercept)                      < 2e-16 ***
## ZTZT2-4                         0.510023
## ZTZT4-6                         0.627158
## ZTZT6-8                         0.959031
## ZTZT8-10                        0.274598
## ZTZT10-12                       0.062134 .
## ZTZT12-14                       5.41e-12 ***
## ZTZT14-16                       1.36e-06 ***
## ZTZT16-18                       0.000264 ***
## ZTZT18-20                       0.008028 **
## ZTZT20-22                       0.108014
## ZTZT22-24                       4.85e-10 ***
## DayDay9                         0.285681
## VigilanceREM                    6.35e-10 ***
## VigilanceWake                   0.853286
## ZTZT2-4:DayDay9                 0.267204
## ZTZT4-6:DayDay9                 0.108784
## ZTZT6-8:DayDay9                 0.055220 .
## ZTZT8-10:DayDay9                0.033758 *
## ZTZT10-12:DayDay9               0.038502 *
## ZTZT12-14:DayDay9               0.442679
## ZTZT14-16:DayDay9               0.541806
## ZTZT16-18:DayDay9               0.952001
```

```
## ZTZT18-20:DayDay9               0.012219 *
## ZTZT20-22:DayDay9               0.081699 .
## ZTZT22-24:DayDay9               0.497676
## ZTZT2-4:VigilanceREM            0.997516
## ZTZT4-6:VigilanceREM            0.937958
## ZTZT6-8:VigilanceREM            0.714128
## ZTZT8-10:VigilanceREM           0.333214
## ZTZT10-12:VigilanceREM          0.183894
## ZTZT12-14:VigilanceREM          2.27e-05 ***
## ZTZT14-16:VigilanceREM          0.005507 **
## ZTZT16-18:VigilanceREM          0.033931 *
## ZTZT18-20:VigilanceREM          0.169975
## ZTZT20-22:VigilanceREM          0.454471
## ZTZT22-24:VigilanceREM          0.000191 ***
## ZTZT2-4:VigilanceWake           0.163633
## ZTZT4-6:VigilanceWake           0.341034
## ZTZT6-8:VigilanceWake           0.796894
## ZTZT8-10:VigilanceWake          0.177164
## ZTZT10-12:VigilanceWake         0.008704 **
## ZTZT12-14:VigilanceWake          < 2e-16 ***
## ZTZT14-16:VigilanceWake         2.13e-13 ***
## ZTZT16-18:VigilanceWake         2.66e-08 ***
## ZTZT18-20:VigilanceWake         2.40e-05 ***
## ZTZT20-22:VigilanceWake         0.007951 **
## ZTZT22-24:VigilanceWake          < 2e-16 ***
## DayDay9:VigilanceREM            0.532232
## DayDay9:VigilanceWake           0.101260
## ZTZT2-4:DayDay9:VigilanceREM    0.434672
## ZTZT4-6:DayDay9:VigilanceREM    0.311890
## ZTZT6-8:DayDay9:VigilanceREM    0.216200
## ZTZT8-10:DayDay9:VigilanceREM   0.185351
## ZTZT10-12:DayDay9:VigilanceREM  0.178885
## ZTZT12-14:DayDay9:VigilanceREM  0.633935
## ZTZT14-16:DayDay9:VigilanceREM  0.696264
## ZTZT16-18:DayDay9:VigilanceREM  0.911184
## ZTZT18-20:DayDay9:VigilanceREM  0.117791
## ZTZT20-22:DayDay9:VigilanceREM  0.261320
## ZTZT22-24:DayDay9:VigilanceREM  0.682326
## ZTZT2-4:DayDay9:VigilanceWake   0.116086
## ZTZT4-6:DayDay9:VigilanceWake   0.017007 *
## ZTZT6-8:DayDay9:VigilanceWake   0.004749 **
## ZTZT8-10:DayDay9:VigilanceWake  0.001528 **
## ZTZT10-12:DayDay9:VigilanceWake 0.002384 **
## ZTZT12-14:DayDay9:VigilanceWake 0.249395
## ZTZT14-16:DayDay9:VigilanceWake 0.366210
## ZTZT16-18:DayDay9:VigilanceWake 0.987119
## ZTZT18-20:DayDay9:VigilanceWake 0.000187 ***
## ZTZT20-22:DayDay9:VigilanceWake 0.010324 *
## ZTZT22-24:DayDay9:VigilanceWake 0.303509
## ---
## Signif. codes:  0 '***' 0.001 '**' 0.01 '*' 0.05 '.' 0.1 ' ' 1
##
## Correlation matrix not shown by default, as p = 72 > 12.
```

```
## Use print(x, correlation=TRUE)  or
##     vcov(x)        if you need it

## optimizer (nloptwrap) convergence code: 0 (OK)
## boundary (singular) fit: see help('isSingular')
```

```r
emmeans(mod, trt.vs.ctrl ~ Day | ZT | Vigilance, ref="Baseline")
```

```
## $emmeans
## ZT = ZT0-2, Vigilance = NREM:
##  Day        emmean     SE  df  lower.CL upper.CL
##  Baseline 0.47074 0.0446 360  3.83e-01   0.5584
##  Day9     0.40333 0.0446 360  3.16e-01   0.4910
##
## ZT = ZT2-4, Vigilance = NREM:
##  Day        emmean     SE  df  lower.CL upper.CL
##  Baseline 0.51231 0.0446 360  4.25e-01   0.6000
##  Day9     0.54398 0.0446 360  4.56e-01   0.6316
##
## ZT = ZT4-6, Vigilance = NREM:
##  Day        emmean     SE  df  lower.CL upper.CL
##  Baseline 0.50139 0.0446 360  4.14e-01   0.5891
##  Day9     0.57731 0.0446 360  4.90e-01   0.6650
##
## ZT = ZT6-8, Vigilance = NREM:
##  Day        emmean     SE  df  lower.CL upper.CL
##  Baseline 0.46750 0.0446 360  3.80e-01   0.5552
##  Day9     0.57157 0.0446 360  4.84e-01   0.6592
##
## ZT = ZT8-10, Vigilance = NREM:
##  Day        emmean     SE  df  lower.CL upper.CL
##  Baseline 0.40176 0.0446 360  3.14e-01   0.4894
##  Day9     0.52435 0.0446 360  4.37e-01   0.6120
##
## ZT = ZT10-12, Vigilance = NREM:
##  Day        emmean     SE  df  lower.CL upper.CL
##  Baseline 0.35278 0.0446 360  2.65e-01   0.4404
##  Day9     0.47056 0.0446 360  3.83e-01   0.5582
##
## ZT = ZT12-14, Vigilance = NREM:
##  Day        emmean     SE  df  lower.CL upper.CL
##  Baseline 0.02102 0.0446 360 -6.66e-02   0.1087
##  Day9     0.02213 0.0446 360 -6.55e-02   0.1098
##
## ZT = ZT14-16, Vigilance = NREM:
##  Day        emmean     SE  df  lower.CL upper.CL
##  Baseline 0.16102 0.0446 360  7.34e-02   0.2487
##  Day9     0.14806 0.0446 360  6.04e-02   0.2357
##
## ZT = ZT16-18, Vigilance = NREM:
##  Day        emmean     SE  df  lower.CL upper.CL
##  Baseline 0.23843 0.0446 360  1.51e-01   0.3261
##  Day9     0.17639 0.0446 360  8.87e-02   0.2641
##
## ZT = ZT18-20, Vigilance = NREM:
```

```
##  Day       emmean     SE  df  lower.CL upper.CL
##  Baseline 0.30269 0.0446 360  2.15e-01   0.3904
##  Day9     0.45981 0.0446 360  3.72e-01   0.5475
##
## ZT = ZT20-22, Vigilance = NREM:
##  Day       emmean     SE  df  lower.CL upper.CL
##  Baseline 0.36917 0.0446 360  2.82e-01   0.4568
##  Day9     0.45741 0.0446 360  3.70e-01   0.5451
##
## ZT = ZT22-24, Vigilance = NREM:
##  Day       emmean     SE  df  lower.CL upper.CL
##  Baseline 0.06728 0.0446 360 -2.04e-02   0.1549
##  Day9     0.06039 0.0446 360 -2.73e-02   0.1481
##
## ZT = ZT0-2, Vigilance = REM:
##  Day       emmean     SE  df  lower.CL upper.CL
##  Baseline 0.07019 0.0446 360 -1.75e-02   0.1579
##  Day9     0.05852 0.0446 360 -2.91e-02   0.1462
##
## ZT = ZT2-4, Vigilance = REM:
##  Day       emmean     SE  df  lower.CL upper.CL
##  Baseline 0.11148 0.0446 360  2.38e-02   0.1991
##  Day9     0.10028 0.0446 360  1.26e-02   0.1879
##
## ZT = ZT4-6, Vigilance = REM:
##  Day       emmean     SE  df  lower.CL upper.CL
##  Baseline 0.09389 0.0446 360  6.22e-03   0.1816
##  Day9     0.09787 0.0446 360  1.02e-02   0.1855
##
## ZT = ZT6-8, Vigilance = REM:
##  Day       emmean     SE  df  lower.CL upper.CL
##  Baseline 0.09963 0.0446 360  1.20e-02   0.1873
##  Day9     0.10324 0.0446 360  1.56e-02   0.1909
##
## ZT = ZT8-10, Vigilance = REM:
##  Day       emmean     SE  df  lower.CL upper.CL
##  Baseline 0.08759 0.0446 360 -7.33e-05   0.1753
##  Day9     0.09861 0.0446 360  1.09e-02   0.1863
##
## ZT = ZT10-12, Vigilance = REM:
##  Day       emmean     SE  df  lower.CL upper.CL
##  Baseline 0.07093 0.0446 360 -1.67e-02   0.1586
##  Day9     0.07463 0.0446 360 -1.30e-02   0.1623
##
## ZT = ZT12-14, Vigilance = REM:
##  Day       emmean     SE  df  lower.CL upper.CL
##  Baseline 0.00324 0.0446 360 -8.44e-02   0.0909
##  Day9     0.00000 0.0446 360 -8.77e-02   0.0877
##
## ZT = ZT14-16, Vigilance = REM:
##  Day       emmean     SE  df  lower.CL upper.CL
##  Baseline 0.00944 0.0446 360 -7.82e-02   0.0971
##  Day9     0.00296 0.0446 360 -8.47e-02   0.0906
##
```

```
## ZT = ZT16-18, Vigilance = REM:
##  Day       emmean     SE  df  lower.CL upper.CL
##  Baseline 0.02769 0.0446 360 -6.00e-02   0.1154
##  Day9     0.00731 0.0446 360 -8.04e-02   0.0950
##
## ZT = ZT18-20, Vigilance = REM:
##  Day       emmean     SE  df  lower.CL upper.CL
##  Baseline 0.02472 0.0446 360 -6.29e-02   0.1124
##  Day9     0.03991 0.0446 360 -4.78e-02   0.1276
##
## ZT = ZT20-22, Vigilance = REM:
##  Day       emmean     SE  df  lower.CL upper.CL
##  Baseline 0.03537 0.0446 360 -5.23e-02   0.1230
##  Day9     0.03750 0.0446 360 -5.02e-02   0.1252
##
## ZT = ZT22-24, Vigilance = REM:
##  Day       emmean     SE  df  lower.CL upper.CL
##  Baseline 0.00279 0.0446 360 -8.49e-02   0.0905
##  Day9     0.00000 0.0446 360 -8.77e-02   0.0877
##
## ZT = ZT0-2, Vigilance = Wake:
##  Day       emmean     SE  df  lower.CL upper.CL
##  Baseline 0.45907 0.0446 360  3.71e-01   0.5467
##  Day9     0.53815 0.0446 360  4.50e-01   0.6258
##
## ZT = ZT2-4, Vigilance = Wake:
##  Day       emmean     SE  df  lower.CL upper.CL
##  Baseline 0.37620 0.0446 360  2.89e-01   0.4639
##  Day9     0.35574 0.0446 360  2.68e-01   0.4434
##
## ZT = ZT4-6, Vigilance = Wake:
##  Day       emmean     SE  df  lower.CL upper.CL
##  Baseline 0.40472 0.0446 360  3.17e-01   0.4924
##  Day9     0.32481 0.0446 360  2.37e-01   0.4125
##
## ZT = ZT6-8, Vigilance = Wake:
##  Day       emmean     SE  df  lower.CL upper.CL
##  Baseline 0.43287 0.0446 360  3.45e-01   0.5205
##  Day9     0.32519 0.0446 360  2.38e-01   0.4129
##
## ZT = ZT8-10, Vigilance = Wake:
##  Day       emmean     SE  df  lower.CL upper.CL
##  Baseline 0.51065 0.0446 360  4.23e-01   0.5983
##  Day9     0.37704 0.0446 360  2.89e-01   0.4647
##
## ZT = ZT10-12, Vigilance = Wake:
##  Day       emmean     SE  df  lower.CL upper.CL
##  Baseline 0.57630 0.0446 360  4.89e-01   0.6640
##  Day9     0.45481 0.0446 360  3.67e-01   0.5425
##
## ZT = ZT12-14, Vigilance = Wake:
##  Day       emmean     SE  df  lower.CL upper.CL
##  Baseline 0.97574 0.0446 360  8.88e-01   1.0634
##  Day9     0.97787 0.0446 360  8.90e-01   1.0655
```

```
##
## ZT = ZT14-16, Vigilance = Wake:
##  Day       emmean     SE  df  lower.CL upper.CL
##  Baseline 0.82954 0.0446 360  7.42e-01   0.9172
##  Day9     0.84898 0.0446 360  7.61e-01   0.9366
##
## ZT = ZT16-18, Vigilance = Wake:
##  Day       emmean     SE  df  lower.CL upper.CL
##  Baseline 0.73389 0.0446 360  6.46e-01   0.8216
##  Day9     0.81630 0.0446 360  7.29e-01   0.9040
##
## ZT = ZT18-20, Vigilance = Wake:
##  Day       emmean     SE  df  lower.CL upper.CL
##  Baseline 0.67259 0.0446 360  5.85e-01   0.7603
##  Day9     0.50028 0.0446 360  4.13e-01   0.5879
##
## ZT = ZT20-22, Vigilance = Wake:
##  Day       emmean     SE  df  lower.CL upper.CL
##  Baseline 0.59546 0.0446 360  5.08e-01   0.6831
##  Day9     0.50509 0.0446 360  4.17e-01   0.5928
##
## ZT = ZT22-24, Vigilance = Wake:
##  Day       emmean     SE  df  lower.CL upper.CL
##  Baseline 0.92993 0.0446 360  8.42e-01   1.0176
##  Day9     0.93961 0.0446 360  8.52e-01   1.0273
##
## Degrees-of-freedom method: kenward-roger
## Confidence level used: 0.95
##
## $contrasts
## ZT = ZT0-2, Vigilance = NREM:
##  contrast        estimate    SE  df t.ratio p.value
##  Day9 - Baseline -0.06741 0.063 355  -1.069  0.2857
##
## ZT = ZT2-4, Vigilance = NREM:
##  contrast        estimate    SE  df t.ratio p.value
##  Day9 - Baseline  0.03167 0.063 355   0.502  0.6158
##
## ZT = ZT4-6, Vigilance = NREM:
##  contrast        estimate    SE  df t.ratio p.value
##  Day9 - Baseline  0.07593 0.063 355   1.204  0.2293
##
## ZT = ZT6-8, Vigilance = NREM:
##  contrast        estimate    SE  df t.ratio p.value
##  Day9 - Baseline  0.10407 0.063 355   1.651  0.0997
##
## ZT = ZT8-10, Vigilance = NREM:
##  contrast        estimate    SE  df t.ratio p.value
##  Day9 - Baseline  0.12259 0.063 355   1.945  0.0526
##
## ZT = ZT10-12, Vigilance = NREM:
##  contrast        estimate    SE  df t.ratio p.value
##  Day9 - Baseline  0.11778 0.063 355   1.868  0.0626
##
```

```
## ZT = ZT12-14, Vigilance = NREM:
##  contrast        estimate    SE  df t.ratio p.value
##  Day9 - Baseline  0.00111 0.063 355   0.018  0.9859
##
## ZT = ZT14-16, Vigilance = NREM:
##  contrast        estimate    SE  df t.ratio p.value
##  Day9 - Baseline -0.01296 0.063 355  -0.206  0.8372
##
## ZT = ZT16-18, Vigilance = NREM:
##  contrast        estimate    SE  df t.ratio p.value
##  Day9 - Baseline -0.06204 0.063 355  -0.984  0.3258
##
## ZT = ZT18-20, Vigilance = NREM:
##  contrast        estimate    SE  df t.ratio p.value
##  Day9 - Baseline  0.15713 0.063 355   2.492  0.0131
##
## ZT = ZT20-22, Vigilance = NREM:
##  contrast        estimate    SE  df t.ratio p.value
##  Day9 - Baseline  0.08824 0.063 355   1.400  0.1625
##
## ZT = ZT22-24, Vigilance = NREM:
##  contrast        estimate    SE  df t.ratio p.value
##  Day9 - Baseline -0.00688 0.063 355  -0.109  0.9131
##
## ZT = ZT0-2, Vigilance = REM:
##  contrast        estimate    SE  df t.ratio p.value
##  Day9 - Baseline -0.01167 0.063 355  -0.185  0.8533
##
## ZT = ZT2-4, Vigilance = REM:
##  contrast        estimate    SE  df t.ratio p.value
##  Day9 - Baseline -0.01120 0.063 355  -0.178  0.8590
##
## ZT = ZT4-6, Vigilance = REM:
##  contrast        estimate    SE  df t.ratio p.value
##  Day9 - Baseline  0.00398 0.063 355   0.063  0.9497
##
## ZT = ZT6-8, Vigilance = REM:
##  contrast        estimate    SE  df t.ratio p.value
##  Day9 - Baseline  0.00361 0.063 355   0.057  0.9544
##
## ZT = ZT8-10, Vigilance = REM:
##  contrast        estimate    SE  df t.ratio p.value
##  Day9 - Baseline  0.01102 0.063 355   0.175  0.8614
##
## ZT = ZT10-12, Vigilance = REM:
##  contrast        estimate    SE  df t.ratio p.value
##  Day9 - Baseline  0.00370 0.063 355   0.059  0.9532
##
## ZT = ZT12-14, Vigilance = REM:
##  contrast        estimate    SE  df t.ratio p.value
##  Day9 - Baseline -0.00324 0.063 355  -0.051  0.9590
##
## ZT = ZT14-16, Vigilance = REM:
##  contrast        estimate    SE  df t.ratio p.value
```

```
##  Day9 - Baseline -0.00648 0.063 355  -0.103  0.9182
##
## ZT = ZT16-18, Vigilance = REM:
##  contrast        estimate    SE  df t.ratio p.value
##  Day9 - Baseline -0.02037 0.063 355  -0.323  0.7468
##
## ZT = ZT18-20, Vigilance = REM:
##  contrast        estimate    SE  df t.ratio p.value
##  Day9 - Baseline  0.01519 0.063 355   0.241  0.8098
##
## ZT = ZT20-22, Vigilance = REM:
##  contrast        estimate    SE  df t.ratio p.value
##  Day9 - Baseline  0.00213 0.063 355   0.034  0.9731
##
## ZT = ZT22-24, Vigilance = REM:
##  contrast        estimate    SE  df t.ratio p.value
##  Day9 - Baseline -0.00279 0.063 355  -0.044  0.9647
##
## ZT = ZT0-2, Vigilance = Wake:
##  contrast        estimate    SE  df t.ratio p.value
##  Day9 - Baseline  0.07907 0.063 355   1.254  0.2106
##
## ZT = ZT2-4, Vigilance = Wake:
##  contrast        estimate    SE  df t.ratio p.value
##  Day9 - Baseline -0.02046 0.063 355  -0.325  0.7457
##
## ZT = ZT4-6, Vigilance = Wake:
##  contrast        estimate    SE  df t.ratio p.value
##  Day9 - Baseline -0.07991 0.063 355  -1.268  0.2058
##
## ZT = ZT6-8, Vigilance = Wake:
##  contrast        estimate    SE  df t.ratio p.value
##  Day9 - Baseline -0.10769 0.063 355  -1.708  0.0885
##
## ZT = ZT8-10, Vigilance = Wake:
##  contrast        estimate    SE  df t.ratio p.value
##  Day9 - Baseline -0.13361 0.063 355  -2.119  0.0348
##
## ZT = ZT10-12, Vigilance = Wake:
##  contrast        estimate    SE  df t.ratio p.value
##  Day9 - Baseline -0.12148 0.063 355  -1.927  0.0548
##
## ZT = ZT12-14, Vigilance = Wake:
##  contrast        estimate    SE  df t.ratio p.value
##  Day9 - Baseline  0.00213 0.063 355   0.034  0.9731
##
## ZT = ZT14-16, Vigilance = Wake:
##  contrast        estimate    SE  df t.ratio p.value
##  Day9 - Baseline  0.01944 0.063 355   0.308  0.7579
##
## ZT = ZT16-18, Vigilance = Wake:
##  contrast        estimate    SE  df t.ratio p.value
##  Day9 - Baseline  0.08241 0.063 355   1.307  0.1920
##
```

```
## ZT = ZT18-20, Vigilance = Wake:
##  contrast        estimate    SE  df t.ratio p.value
##  Day9 - Baseline -0.17231 0.063 355  -2.733  0.0066
##
## ZT = ZT20-22, Vigilance = Wake:
##  contrast        estimate    SE  df t.ratio p.value
##  Day9 - Baseline -0.09037 0.063 355  -1.433  0.1526
##
## ZT = ZT22-24, Vigilance = Wake:
##  contrast        estimate    SE  df t.ratio p.value
##  Day9 - Baseline  0.00968 0.063 355   0.153  0.8781
##
## Degrees-of-freedom method: kenward-roger
```

### Session Information

```
sessionInfo()
```

```
## R version 4.0.3 (2020-10-10)
## Platform: x86_64-pc-linux-gnu (64-bit)
## Running under: Ubuntu 18.04 LTS
##
## Matrix products: default
## BLAS:   /usr/lib/x86_64-linux-gnu/openblas/libblas.so.3
## LAPACK: /usr/lib/x86_64-linux-gnu/libopenblasp-r0.2.20.so
##
## locale:
##  [1] LC_CTYPE=C.UTF-8          LC_NUMERIC=C
##  [3] LC_TIME=C.UTF-8           LC_COLLATE=C.UTF-8
##  [5] LC_MONETARY=C.UTF-8       LC_MESSAGES=C.UTF-8
##  [7] LC_PAPER=C.UTF-8          LC_NAME=C.UTF-8
##  [9] LC_ADDRESS=C.UTF-8        LC_TELEPHONE=C.UTF-8
## [11] LC_MEASUREMENT=C.UTF-8    LC_IDENTIFICATION=C.UTF-8
##
## attached base packages:
##  [1] grid      parallel  stats4    stats     graphics  grDevices utils
##  [8] datasets  methods   base
##
## other attached packages:
##  [1] readxl_1.4.0                xlsx_0.6.5
##  [3] rstatix_0.7.0               org.Rn.eg.db_3.12.0
##  [5] topGO_2.42.0                SparseM_1.81
##  [7] GO.db_3.12.1                AnnotationDbi_1.52.0
##  [9] graph_1.68.0                emmeans_1.7.3
## [11] lmerTest_3.1-3              lme4_1.1-28
## [13] Matrix_1.5-3                sechm_1.9.4
## [15] ComplexHeatmap_2.15.1       stringr_1.5.0
## [17] ggrepel_0.9.3               scales_1.2.1
## [19] SEtools_1.9.3               plgINS_0.1.5
## [21] SummarizedExperiment_1.20.0 Biobase_2.50.0
## [23] GenomicRanges_1.42.0        GenomeInfoDb_1.26.7
## [25] IRanges_2.24.1              S4Vectors_0.28.1
```

```
## [27] BiocGenerics_0.36.1         MatrixGenerics_1.2.1
## [29] matrixStats_0.63.0          ggpubr_0.4.0
## [31] ggsci_3.0.0                 ggplot2_3.4.2
##
## loaded via a namespace (and not attached):
##   [1] backports_1.4.1        circlize_0.4.15        plyr_1.8.8
##   [4] splines_4.0.3          BiocParallel_1.24.1    TH.data_1.1-0
##   [7] sva_3.38.0             digest_0.6.31          foreach_1.5.2
##  [10] htmltools_0.5.5        fansi_1.0.4            magrittr_2.0.3
##  [13] memoise_2.0.1          cluster_2.1.4          doParallel_1.0.17
##  [16] tzdb_0.3.0             openxlsx_4.2.5         limma_3.46.0
##  [19] Biostrings_2.58.0      readr_2.1.2            annotate_1.68.0
##  [22] sandwich_3.0-1         colorspace_2.1-0       blob_1.2.2
##  [25] xfun_0.38              dplyr_1.1.1            crayon_1.5.2
##  [28] RCurl_1.98-1.6         jsonlite_1.8.4         genefilter_1.72.1
##  [31] GEOquery_2.58.0        zoo_1.8-9              survival_3.3-1
##  [34] iterators_1.0.14       glue_1.6.2             registry_0.5-1
##  [37] SRAdb_1.52.0           gtable_0.3.3           zlibbioc_1.36.0
##  [40] XVector_0.30.0         GetoptLong_1.0.5       DelayedArray_0.16.3
##  [43] V8_4.1.0               car_3.0-12             shape_1.4.6
##  [46] abind_1.4-5            mvtnorm_1.1-3          DBI_1.1.3
##  [49] edgeR_3.32.1           randomcoloR_1.1.0.1    Rcpp_1.0.10
##  [52] xtable_1.8-4           clue_0.3-60            bit_4.0.5
##  [55] httr_1.4.2             RColorBrewer_1.1-3     ellipsis_0.3.2
##  [58] rJava_1.0-6            pkgconfig_2.0.3        XML_3.99-0.9
##  [61] locfit_1.5-9.4         utf8_1.2.3             tidyselect_1.2.0
##  [64] rlang_1.1.0            cellranger_1.1.0       munsell_0.5.0
##  [67] tools_4.0.3            cachem_1.0.7           cli_3.6.1
##  [70] generics_0.1.3         RSQLite_2.2.11         broom_0.7.12
##  [73] evaluate_0.20          fastmap_1.1.1          yaml_2.3.7
##  [76] knitr_1.42             bit64_4.0.5            zip_2.2.0
##  [79] purrr_1.0.1            nlme_3.1-157           xml2_1.3.3
##  [82] pbkrtest_0.5.1         compiler_4.0.3         rstudioapi_0.13
##  [85] curl_5.0.0             png_0.1-7              ggsignif_0.6.4
##  [88] tibble_3.2.1           geneplotter_1.68.0     stringi_1.7.12
##  [91] lattice_0.21-8         nloptr_1.2.2.3         vctrs_0.6.1
##  [94] pillar_1.9.0           lifecycle_1.0.3        GlobalOptions_0.1.2
##  [97] estimability_1.4.1     data.table_1.14.8      bitops_1.0-7
## [100] seriation_1.3.4        R6_2.5.1               TSP_1.2-0
## [103] codetools_0.2-19       boot_1.3-28.1          MASS_7.3-56
## [106] xlsxjars_0.6.1         DESeq2_1.30.1          rjson_0.2.21
## [109] withr_2.5.0            multcomp_1.4-18        GenomeInfoDbData_1.2.4
## [112] mgcv_1.8-39            hms_1.1.1              coda_0.19-4
## [115] tidyr_1.2.0            minqa_1.2.4            rmarkdown_2.13
## [118] carData_3.0-5          Rtsne_0.16             numDeriv_2016.8-1.1
```
